# Sorting & Sequencing Flies by Size: Identification of novel TOR regulators and Parameters for Successful Sorting

**DOI:** 10.1101/119719

**Authors:** Katrin Strassburger, Tanja Zöller, Thomas Sandmann, Svenja Leible, Grainne Kerr, Michael Boutros, Aurelio A. Teleman

## Abstract

As DNA sequencing throughput increases, novel strategies for discovering genes that affect traits of interest become available. One strategy starts with a population of animals and selects individuals over multiple generations for a particular trait. Subsequent whole genome sequencing should identify loci affecting this trait. We apply this strategy by sorting flies for wing length over 18 generations, obtaining two populations that differ in wing length by 20%. Flies with longer wings had increased overall body sizes and elevated TOR activity, suggesting that genetic variation targets TOR signaling to influence body size. High-throughput sequencing of big and small flies identified thousands of single nucleotide polymorphisms that differed between the two populations, leading us to identify five novel regulators of TOR signaling. Surprisingly, stochastic simulations of the process show that large fractions of the genetic differences between the big and small flies are probably biological false positives, selected by chance by random drift. We employ these computer simulations to identify experimental setup parameters to improve the signal-to-noise ratio for successfully running sort-and-sequence experiments – a resource which will hopefully be useful for the community.

## Introduction

Forward-genetics has successfully been employed to discover genes affecting a particular biological process of interest in a variety of model organisms. Typically, as few genes as possible are perturbed per animal, and then large numbers of individuals are screened for the trait of interest. This couples a genomic perturbation to the phenotypic output. This approach is successful at identifying genes with strong roles in the biological process of interest, but is less well suited towards identifying genes with redundant roles, or for studying complex traits that are modulated via interactions amongst large numbers of genes. The reduced cost of whole genome sequencing has enabled a novel approach leveraging the broad genetic variability present in nature, rather than experimentally-induced mutations, by combining laboratory-based evolution with complete-genome sequencing. Conceptually, a selective pressure is applied to the population to enrich for variants affecting the selected trait. One implementation of this approach is to start with one population of animals and to select for a trait of interest in the laboratory over several generations to generate two differing end-populations, followed by whole-genome sequencing to identify the genetic differences between the two populations (“sort-and-sequence”). Other implementations can also be envisaged - such as performing a single round of selection followed by sequencing – which can also be applied to a human population in the form of a genome-wide association study. Unlike the forward genetics approach, where each experimental animal has only a few mutations with strong phenotypic consequences, in the ‘sort-and-sequence’ approach each individual animal in a population may have many genetic differences compared to the other animals. Therefore, one needs to sequence a large number of individuals to find correlations and to pinpoint relevant genetic loci.

Animal size is a complex trait, affected by many genetic loci (Gockel *et al.* 2002; Lango Allen *et al.* 2010; Lettre 2011; Perola 2011). Despite strong interest in uncovering the molecular mechanisms by which developing tissues and organs can measure their size and regulate their growth, we do not yet have a good understanding of these processes. Forward and reverse genetic approaches, in particular in Drosophila, have identified genes affecting organ size (Stocker and Hafen 2000; Johnston and Gallant 2002; Mirth and Riddiford 2007; Shingleton 2010). In particular, components of two major signaling pathways have been found: the insulin/insulin-like growth factor signaling (IIS) pathway and the Hippo/Yorkie pathway (Oldham and Hafen 2003; Pan 2007; Tumaneng *et al.* 2012). The IIS pathway consists of a number of kinases including PI3K, Akt, PDK1 and TOR, which become activated in response to hormonal signaling and environmental cues. These kinases in turn activate a plethora of anabolic processes and repress catabolic processes in the cell (Loewith and Hall 2011; Laplante and Sabatini 2012). One interpretation is that the IIS pathway coordinately scales the size of all tissues in the organism in response to nutrient availability, thereby generating well-proportioned animals of different sizes based on environmental conditions. A second signaling pathway, the Hippo/Yorkie pathway, also has the capacity to powerfully regulate tissue growth. This pathway consists of several kinases including Hippo and Warts, which act in concert to block activation of a transcriptional co-activator Yorkie. When active, Yorkie potently drives cell growth and proliferation by regulating gene transcription (Pan 2007; Genevet and Tapon 2011; Staley and Irvine 2012; Tumaneng *et al.* 2012). The upstream regulatory inputs into the Hippo/Yorkie pathway have been a subject of much recent interest (Grusche *et al.* 2010; Yu and Guan 2013). This pathway promotes tissue growth in response to wounding (Yu and Guan 2013). Whether Yorkie activity also drops at the end of normal animal development to control normal body size is not yet fully understood.

As a test-of-concept for the “sort-and-sequence” approach, we started with one population of Drosophila melanogaster and selected for wing length over 18 generations, yielding a population of ‘big’ and ‘small’ flies. Molecular analysis revealed that the ‘big’ and ‘small’ flies differed in their level of activation of the TOR pathway, but not in activation of the Hippo/Yorkie pathway, suggesting that the selective pressure mainly manipulated the IIS pathway to evolve body size. By whole-genome sequencing we identified almost 7000 Single Nucleotide Polymorphisms (SNPs) whose frequency was significantly different between these two populations, including five novel regulators of TOR. Based on the surprising result that only few of the perturbed genes had an effect on TOR activity, we asked whether the experimental setup can be optimized to improve the frequency of biologically relevant hits. We generated an in silico simulation of the sort-and-sequence approach which identifies parameter combinations that significantly reduce the false-positive rate. This will hopefully be a useful resource for the community to successfully set up sort-and-sequence types of experiments.

## Results

### Sorting flies for wing length successfully generates populations of differing wing length within few generations

To identify genes affecting tissue size, we sorted flies according to wing length (Figure 1A). To obtain a starting population with sufficient genetic variability, we intercrossed two inbred fly strains, w^1118^ and Oregon-R, for 3 generations. We then grew these animals in density-controlled conditions, quantified the wing length of 150 male and 150 female flies, and divided the population in half – flies with ‘big’ wings (length greater than the population mean) and flies with ‘small’ wings (smaller than the population mean). We repeated this process for 18 generations, consistently retaining either the larger 50% of the population or the smaller 50% of the population. As a consequence of this selection, average wing length in the two populations consistently diverged (Figure 1B) starting from the first generation. The difference in the average wing length between the two populations increased almost linearly, yielding two populations with an average wing length difference of 20% (Figure 1B’). Interestingly, compared to the distribution of wing-lengths in the starting population (Figure 1C, black trace), the distribution of wing-lengths in the final ‘big’ population displayed a smaller standard deviation (Figure 1C, green trace) whereas the distribution of wing-length in the final ‘small’ population was increased (Figure 1C, red trace). This lack of homogeneity within the small population could potentially occur if some SNPs causing reduced wing length represented a competitive disadvantage, e.g. display lethality when homozygous, not allowing them to become ‘fixed’ in the population. Flies with wing lengths either smaller or larger than the extremes observed in the initial population were easily observed after a few rounds of sorting (Figure 1C), indicating that the wing lengths in the apparently homogeneous starting population actually reflect the activities of multiple variants counteracting each other.

**Figure 1:**
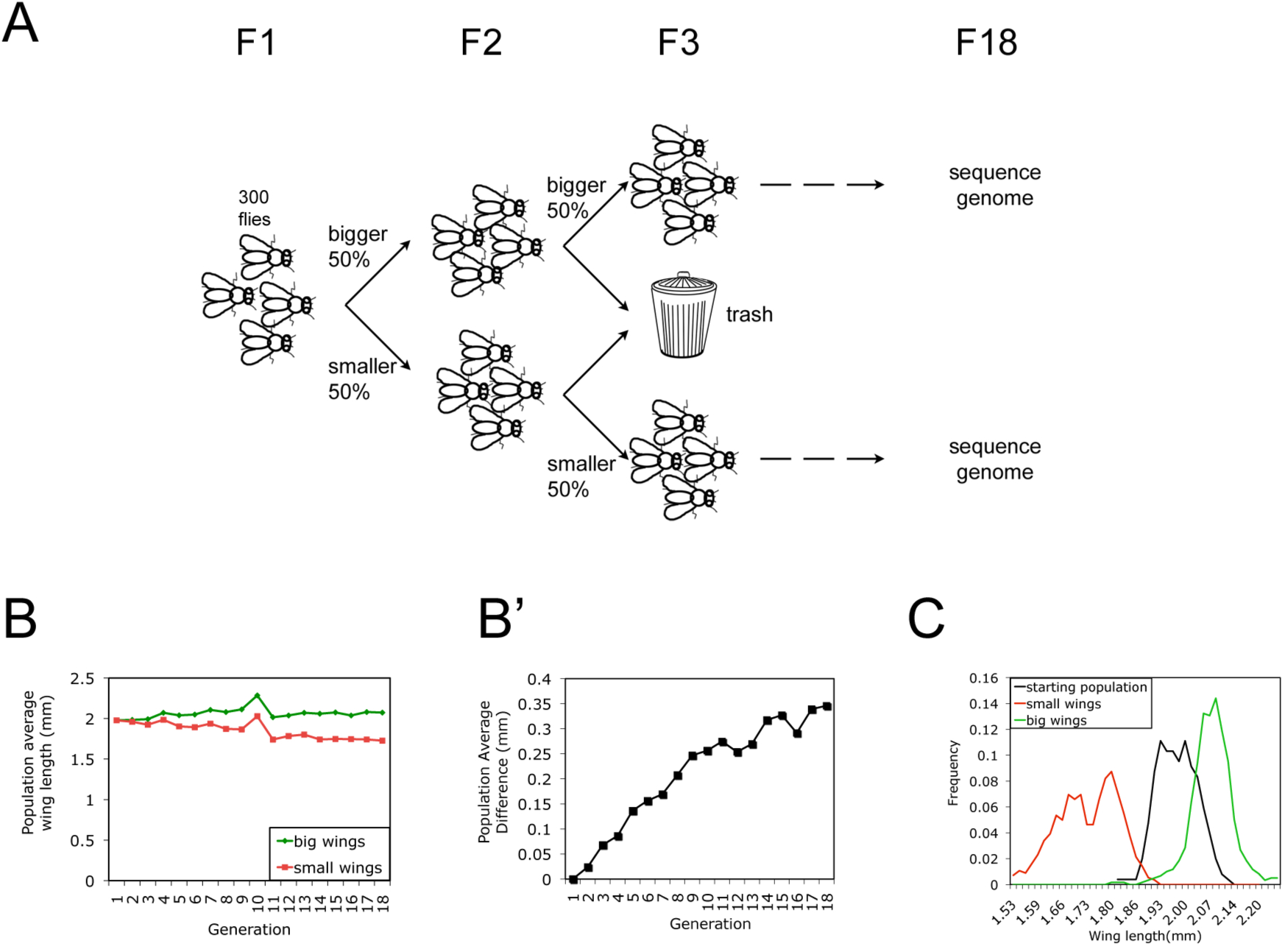
Sorting flies for wing length over 18 generations yields 20% difference in size. (A) Schematic representation of the sorting approach to select flies for differing wing sizes. From a single population of flies, the upper and lower 50% of flies were selected for wing length over 18 generations, after which their genomic DNA was sequenced. (B-B’) Average wing length for the “big” and “small” flies (B) and the difference between the two population averages (B’) over 18 generations of selection. (At generation 10, flies were reared at 19° C to slow down their development and span holidays, which leads to increased body size.) (C) Histogram showing the distribution of wing lengths for the starting fly population (black) and the final big (green) and small (red) populations after 18 generations of selection.

### Selection for wing length caused concomitant selection for total body size

Next, we examined whether modulation of wing size was correlated with a change in overall body size, e.g. if these traits can be selected for independently. Flies from the ‘small’ population displayed reduced wing size (Figure 2A), lower total body weight (Figure 2B), and reduced overall body size (Figure 2C) compared to flies from the ‘big’ population. The difference in body size was already visible at the end of the larval phases when animals formed pupae (Figure 2D). These results suggest that in our starting population, genetic variants affecting total body size were more common than genetic variants specifically affecting wing length. In contrast, embryo size was not significantly different for big and small flies (Figure 2E), indicating that different processes control adult size and egg size.

**Figure 2:**
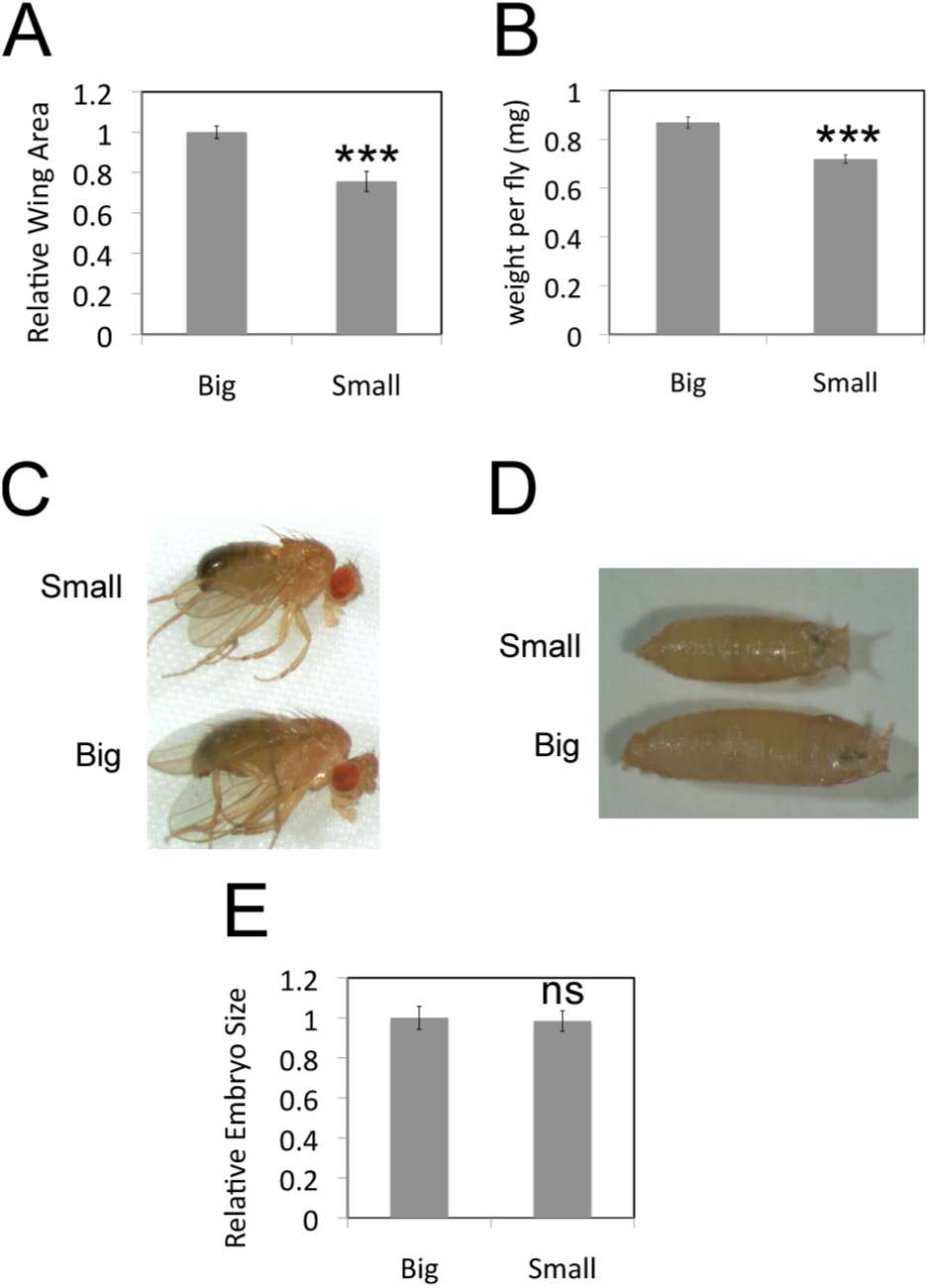
Selection for wing length yields animals of different sizes. (A) Relative wing area of flies from the ‘big’ and ‘small’ populations. (B) Weight of flies from the ‘big’ and ‘small’ populations. (C-D) Representative pictures of adult males (C) and pupae (D) from the ‘small and ‘big’ populations. (E) Unlike total body size, embryo size was not significantly different in the two populations selected for wing length. Embryo size, quantified as area occupied on a picture, from flies of the ‘big’ and ‘small’ populations. Error bars: Std. Dev. *** t-test < 0.001.

### ‘Big’ and ‘small’ flies were selected for differing levels of TOR activation during growth stages

We next aimed to characterize the underlying molecular changes leading to the different body sizes of the two populations. One way tissues size can be modulated is through changes in cell size. We quantified the cell size in wings of ‘big’ and ‘small’ flies and found that cells from the ‘small’ flies were roughly 10% smaller than cells from the ‘big’ flies (Figure 3A). Since the wing area of ‘small’ flies was 25% smaller than that of ‘big’ flies (Figure 2A), this indicates that the wings of ‘small’ flies contained both smaller cells and fewer cells. Signaling via TOR complex 1 (TOR-C1) is known to regulate tissue size in part via altered cell size (Stocker and Hafen 2000; Zhang *et al.* 2000), raising the possibility that TOR-C1 signaling might be altered in ‘small’ versus ‘big’ flies. Since TOR-C1 also regulates organismal metabolism, we asked whether ‘big’ and ‘small’ flies had altered metabolic profiles. Indeed, ‘small’ flies had disproportionately lower levels of triglycerides than ‘big’ flies, both when normalized to total body protein or body weight (Figure 3B and data not shown). A priori, it was unexpected that selecting flies based on wing length should also lead to two populations with strikingly different triglyceride levels. The fact that TOR-C1 regulates both tissue size and organismal metabolism provides a possible underlying mechanistic link to this coupling.

**Figure 3:**
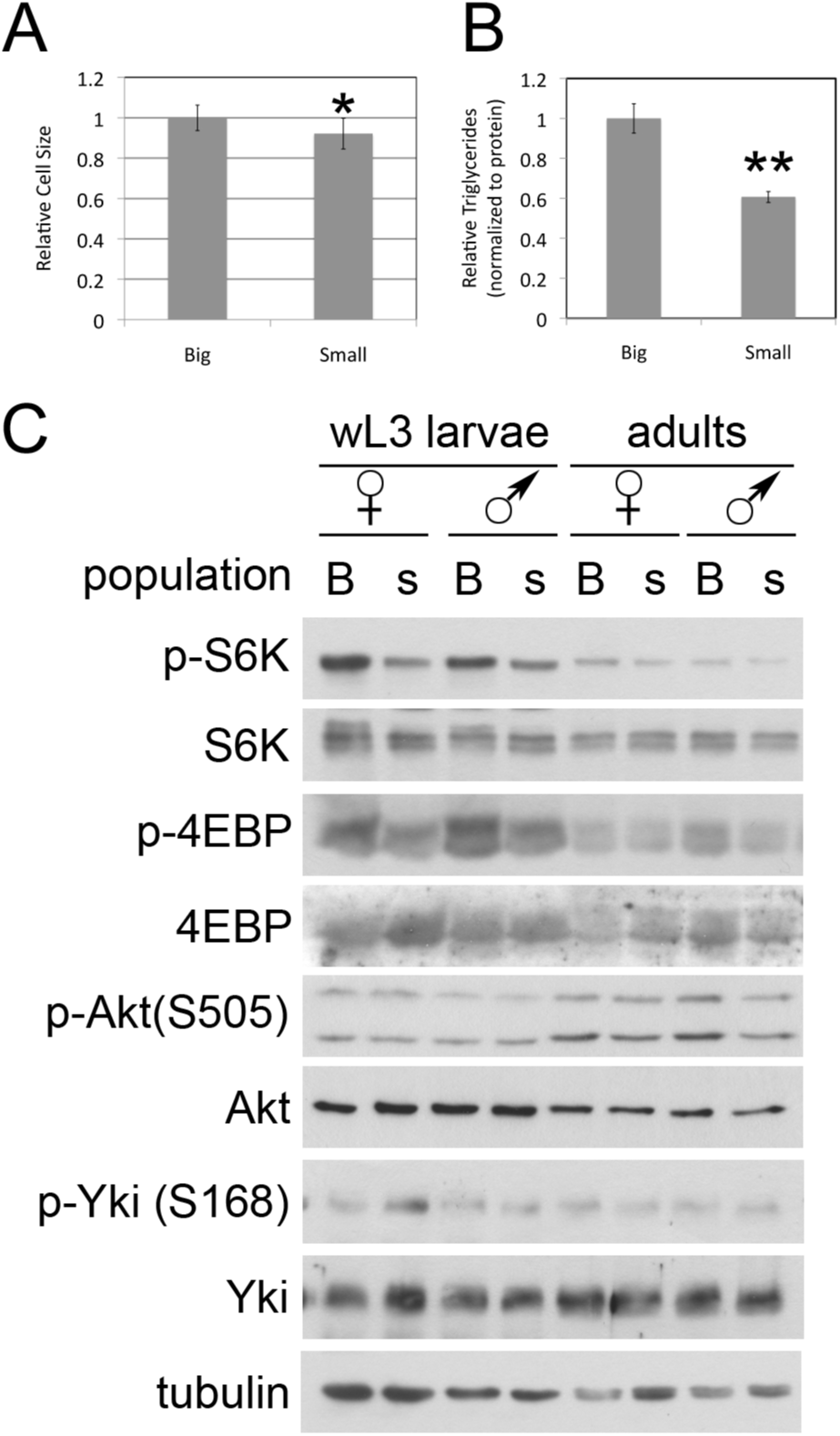
TOR activity altered in flies selected for size. (A) Wings of flies selected to be ‘small’ are composed of cells of smaller size, compared to wings from ‘big’ flies. Cell size quantified by counting trichomes and calculating area per trichome. n=10 for each population. (B) ‘Big’ flies have increased total body triglycerides compared to ‘small’ flies. n= 3 × 8 flies for each population. (C) Western blot analysis reveals that selecting for size causes TOR activity, but not Yorkie activity, to be upregulated during the larval (growth) stages of development, but not in adulthood. TOR activity quantified by phosphorylation of S6 Kinase and 4E-BP. Big “B”, small “s”. Error bars: Std dev. *ttest<0.05; **ttest<0.01.

To assay TOR-C1 signaling directly, we tested the level of phosphorylation of the canonical TOR-C1 targets S6K and 4EBP in extracts of ‘big’ and ‘small’ larvae and adults (Figure 3C). Phosphorylation of S6K and 4E-BP were increased in lysates from ‘big’ larvae compared to ‘small’ larvae (Figure 3C), indicating increased TOR-C1 activation in the ‘big’ larvae. Interestingly, TOR-C1 activity was not as strongly affected in ‘big’ versus ‘small’ adults (Figure 3C), indicating that the genetic differences selected in the two populations specifically affected TOR-C1 activity during the larval phases of development when organismal growth takes place. In contrast to TOR-C1 activity, phosphorylation of Akt on Ser505, a readout for TOR-C2 activity, was not increased in ‘big’ versus ‘small’ larvae (Figure 3C). Furthermore, activation of Yorkie, assessed via a standard inhibitory phosphorylation on Ser168 which reflects activation of the Hippo pathway, was similar in big and small animals (Figure 3C). In sum,these data suggest that the genetic variation present in our flies affected body size at least in part via regulation of TOR-C1 activity during juvenile phases, but not through modulation of the Hippo/Yorkie pathway.

### Whole-genome sequencing reveals many SNPs differentially enriched in the ‘big’ versus ‘small’ populations

To identify genetic differences underpinning the altered body size in the ‘big’ versus ‘small’ populations, we performed whole genome sequencing on genomic DNA extracted from the two populations. We sequenced genomic DNA pooled from either 50 females (i.e. 100 alleles) from the ‘big’ population or 50 females from the ‘small’ population at 30-fold coverage. The number of chromosomes assayed was chosen to be higher than the average sequencing depth, so that the data effectively samples the population (i.e. reads obtained for a given genomic position are probably from different chromosomes within the population). We then asked which SNPs were differentially represented in the two populations. We quantified two parameters for each position in the genome: 1) a measure for the difference in nucleotide frequency between the two populations and 2) a statistical score for the significance of this difference (see Materials & Methods for SNP calling). Selecting only SNPs that were completely polarized in the two populations and applying a stringent p-value cutoff identified 6984 SNPs with differential frequencies between the two populations (Supplemental Table 1, and Experimental Procedures for details), 368 of which led to changes in the amino acid sequences of 279 different genes (Supplemental Figure 2). None of the 279 genes was a known component of the IIS pathway, suggesting that some of the SNPs might affect novel regulators of IIS/TOR signaling.

To assess if these candidate genes can affect TOR-C1 activity, we performed RNAi in S2 cells on all 279 genes and assessed phosphorylation of S6K at Thr398. Knockdown of five candidate genes reduced TOR-C1 activity (Figure 4), validating a functional role in TOR-C1 regulation for these loci. Interestingly, mutation of one of these genes, mip130, yields viable adults with small body sizes, confirming that this gene can affect body size (Beall *et al*. 2004).

**Figure 4:**
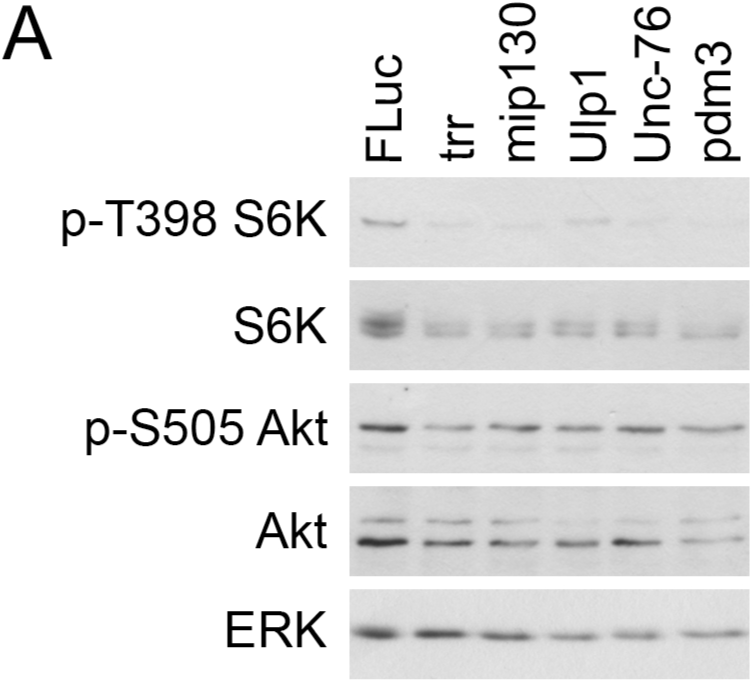
Identification of TOR regulators amongst genes affected by SNPs. (A) Identification of five genes whose knockdown causes reduced TOR-C1 activity in S2 cells, assayed via phosphorylation of S6K on Thr398, a well-established TOR-C1 target.

### Generation of an in silico simulation of fly sorting

Of the 279 genes whose protein coding was differentially affected by SNPs in the big and small populations, five affected TOR-C1 in S2 cells (Figure 4), representing roughly a 2% confirmation rate. We wondered whether the remaining 98% might represent false-positives, or true hits that affect different aspects of size determination that were not assayed using our TOR readout in S2 cells. To assess if and how the specificity for the experiment could be improved, we modeled the selection process using a stochastic Monte Carlo-like simulation of the ‘sort-and-sequence’ process (see Supplemental Material for computer code in the C programming language). Briefly, the simulation randomly generates SNPs throughout the genome (Figure 5A), selects a subset of them as SNPs that affect the trait of interest (in this case body size) in a dominant or recessive fashion, generates a random initial population of flies with random genotype, and then executes the phenotyping, sorting, and breeding over multiple generations, allowing for meiotic cross-overs in the female germline. Finally, the simulation records the genetic makeup of the final population thereby identifying the SNPs that were differentially represented in the two populations. (See Experimental Procedures for details). The model was constructed with a large number of parameters that can be adjusted to fit what we know about fly genetics and to test different experimental setups. These include the number of SNPs present genome-wide, how many of these affect animal size, the number of meiotic cross-overs per chromosome in the female germline, the number of flies selected in each generation, and the number of generations the simulation is allowed to proceed. Importantly, as the identity of the simulated SNPs affecting size are specified by the simulation, their frequency can be monitored during the selection process and compared to SNPs that are differentially enriched simply by chance.

**Figure 5:**
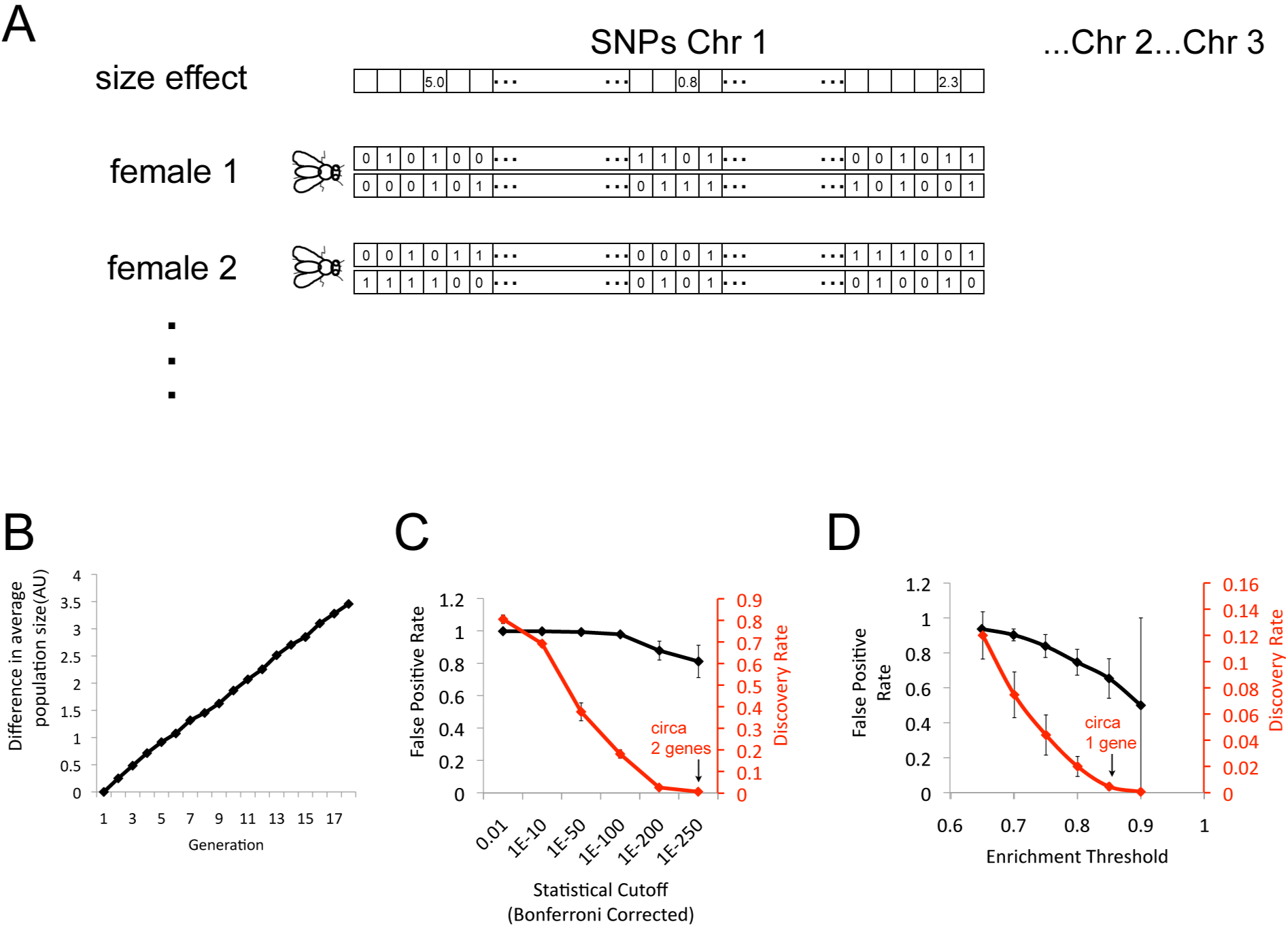
In silico simulation of sorting reveals that the vast majority of differential SNPs between the two populations are ‘false positives’ or ‘passenger mutations’. (A) Schematic representation of the genetic architecture used for the stochastic modeling. (B) In silico simulation of sorting of 300 ‘big’ and 300 ‘small’ flies per generation, assuming 300 SNPs affect size genome-wide, reproduces a gradual divergence in animal size as observed experimentally. (C-D) In silico simulation reveals that the vast majority of SNPs differentially selected for in the two populations are false positives. The false positive rate (black) and the discovery rate (red) indicated for different statistical threshold cut-offs (C). Even the highest possible Bonferroni-corrected statistical cut-off (that yields a non-zero discovery rate) of 10^-250^ leads to an 81% false positive rate. If instead an enrichment cut-off is used (D), representing how strongly a particular SNP was differentially selected for in the two populations, the most stringent cut-off has a false-positive rate of 65%. All simulations were run 3 times with identical parameters, and the mean is shown. Error bars: Std Dev.

We first modeled the experiment by choosing parameters matching the original experimental setup, e.g. sorting 300 flies (150 males and 150 females) per population per generation for 18 generations. We allowed 7 meiotic cross-overs genome-wide per generation in the female germline (Miller *et al.* 2012). From sequencing our ‘big’ versus ‘small’ flies and by comparing them to a reference genome, we estimated that our initial population, generated by combining two inbred laboratory stocks, contained roughly 175,000 SNPs. We estimated that a standard signaling pathway such as the IIS pathway has roughly 30 components, that each component might have two SNPs (as observed in our sequencing), and that 5 different pathways, both signaling and metabolic, might be affecting size, yielding 300 size-affecting SNPs. With these parameters, the model predicted two populations of flies that diverged in average size in a manner similar to what we observed experimentally (Figure 5B). We then asked how many SNPs were differentially selected in the two populations using a statistical measure that was Bonferroni corrected for multiple hypothesis testing. A p-score cut-off of 10^-50^ yielded roughly 15,000 differential SNPs whereas a p-score cut-off of 10^-100^ yielded 2,600 SNPs that were differentially represented in the two populations, on the same order of magnitude as what we observed experimentally. Surprisingly, at these two cut-offs, > 97% of the differentially enriched SNPs did not affect the selected trait (Figure 5C). This means that even though these SNPs were truly differentially selected in the two populations, they were not selected due to their size effects, but through genetic drift.

We examined whether applying an even more stringent statistical cut-off would decrease the false positive rate, however increasing it to the maximal possible score of 10^-250^, which only identifies on average 2 of the 300 size-affecting SNPs, still had a false positive rate of 81% (Figure 5C). Alternatively, we selected SNPs based on an ‘enrichment score’ cut-off, which describes how different the allele frequencies are in the two populations (0 if it is the same in the two populations and 1 if the SNP has become fully polarized in the two populations). Using an enrichment cut-off improved the outcome slightly however a stringent cut-off of 0.8 which identified on average 7 of the 300 size-affecting SNPs still had a false-positive rate of 74% (Figure 5D). In sum, the results from the simulation suggest that many of the SNPs that were differentially selected in the ‘big’ and ‘small’ fly populations from our experimental sorting might be unrelated to animal size determination, raising the question how the experimental setup can be improved to increase the true hit rate.

### Identification of parameters for optimizing a sort-and-sequence approach

Next we varied the model parameters to identify experimental conditions with a low false-positive rate. The analysis above suggested that the false positive rate depends strongly on the relative selection pressure on the size-SNPs versus random drift of the neutral SNPs. Selection pressure on the size-SNPs depends on how many SNPs affect size genome-wide. In the extreme example where only one SNP regulates a phenotype, that SNP will immediately become fixed within a few generations. Instead, when many SNPs affect a phenotype, the selection pressure on any one individual SNP is low. For this reason, we varied the number of SNPs affecting size from 20 to 300 (Figure 6A). This parameter will depend on the phenotype being studied, with simple traits regulated by few SNPs and complex phenotypes regulated by many. The random drift in the frequencies of the neutral SNPs is inversely correlated with the number of flies sorted at every generation. Therefore, we varied the number of selected flies per population from 200 to 1600 flies (Figure 6A). For simple phenotypes affected by only 20 SNPs genome-wide, selecting 400 flies per population per generation (i.e. 800 flies total) is enough to give a threshold with a false positive rate below 5% and a discovery rate of 100% (i.e. all 20 SNPs). In contrast, for complex phenotypes affected by 300 SNPs genome-wide, at least 1600 flies per population (3200 total) need to be sorted to yield a region where the false positive rate is below 5% yet the discovery rate is non-zero (e.g. 0.5 enrichment threshold gives a 3% false-positive rate and a 24% discovery rate, yielding 70 trait-related SNPs).

**Figure 6:**
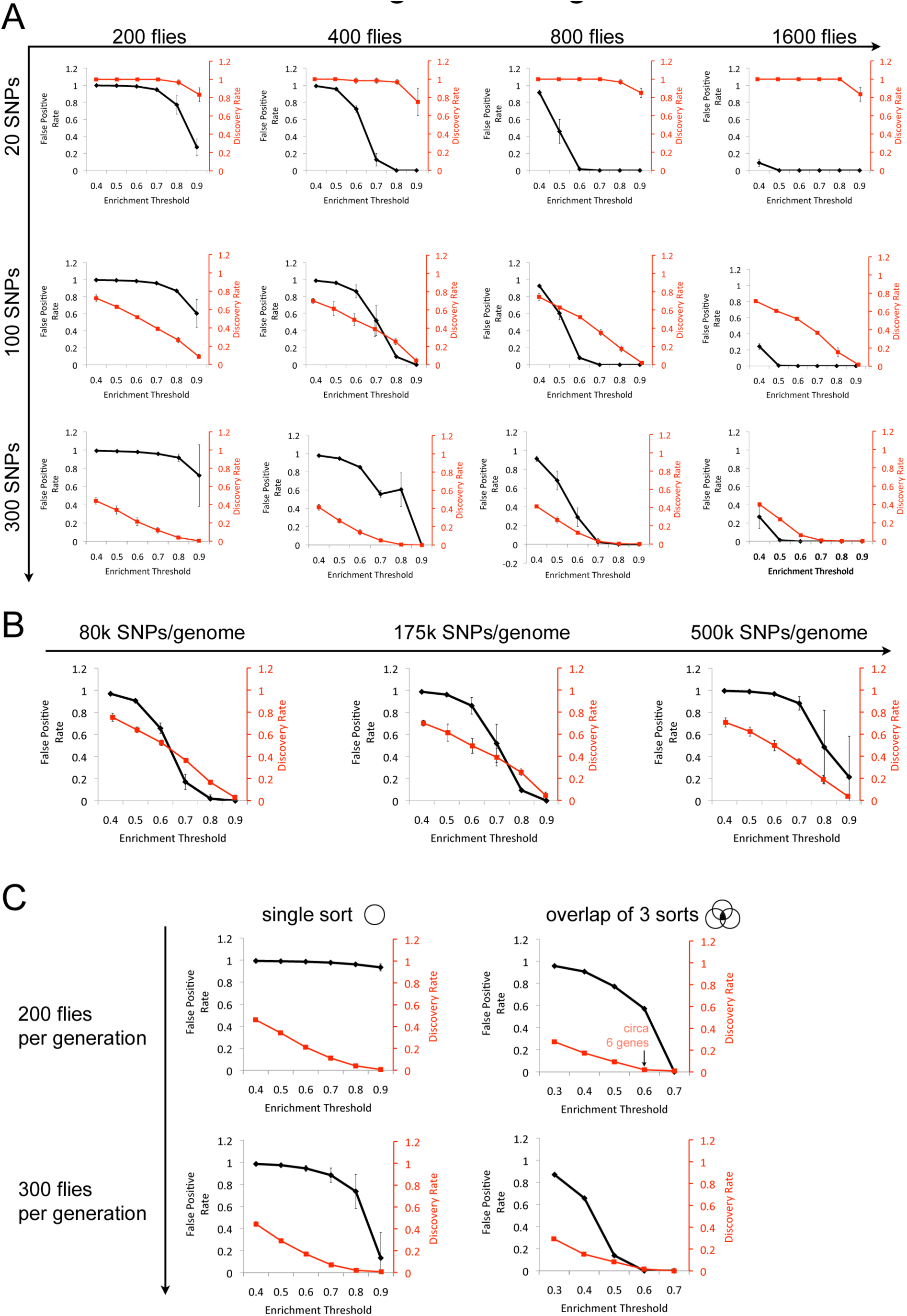
In silico simulation of sorting identifies parameters required for successfully setting up a “sort-and-sequence” approach. (A) False-positive and discovery rates resulting from in silico simulations in which two parameters are varied: the number of SNPs genome-wide affecting the trait of interest, as well as the number of flies sorted per generation in each population. As the number of flies sorted increases, the false-positive rate drops due to reduced chances of differentially selecting neutral SNPs by chance. (B) False-positive rates resulting from in silico simulations with starting populations of differing initial complexity, quantified as the number of floating SNPs genome-wide in the starting population. Simulations were run for 200 flies sorted per population per generation, and 100 SNPs affecting size. (C) Repeating the “sort-and-sequence’ experiment in 3 biological replicates can improve the false positive rate, however the number of animals sorted per generation is more important. All simulations were run 3 times with identical parameters, and the mean is shown. Error bars: Std Dev.

Another parameter that can be varied is the genetic complexity of the starting population: how many SNPs are present in the starting flies. This can be done by selecting in- or out-bred animals as a starting point. Although this does not change the rate of drift of any one individual SNP, it reduces the total number of neutral SNPs that have a chance to be enriched by the end of the experiment, and hence the false-positive rate. As shown in Figure 6B for simulations with 100 SNPs affecting the trait of interest and 200 animals sorted per population per generation, reducing the number of SNPs genome-wide to 80,000 significantly improves the false-positive rate, whereas increasing the SNPs to 500,000 genome-wide renders the experimental results unusable. Recent estimates from the D. melanogaster reference panel identified over 4.5 million SNPs in their populations (Mackay *et al*. 2012), indicating that 500,000 SNPs can easily occur in one starting population.

Lastly, we asked whether performing a sort-and-sequence experiment in biological triplicates, starting with the same initial population of flies, can improve the false-positive rate by looking at only the SNPs that overlap in all three replicates. One could imagine that this would remove all the ‘noise’. However, as shown in Figure 6C for simulations with 300 trait-affecting SNPs, this only significantly improves the outcome under some conditions. For the simulations performed with 200 flies being sorted (top row Figure 6C), repeating the experiment in triplicate still yields false-positive rates that are at best 60%. In contrast, if the number of flies sorted is increased to 300, looking only at the overlap of three biological replicates does reduce the false-positive rate (bottom row, Figure 6C). That said, this type of setup does not perform better than simply sorting 900 flies per generation (compare Figure 6A 300 SNPs and 800 flies with Figure 6C lower right panel). In sum, these stochastic simulations provide a panel of starting parameters that are necessary for performing sort-and-sequence experiments with improved false-positive rates.

## Discussion

We aimed in this study to employ a new screening strategy for uncovering genetic loci affecting animal size in Drosophila, based on the selection of flies for wing length over multiple generations followed by whole-genome sequencing. Interestingly, the average size of flies in the population could easily be shifted both towards increased size and reduced size, indicating that many naturally occurring genetic variants affect animal size. This observation implies that although the observed size of flies within one population may appear homogeneous, it actually reflects the balancing action of many counteracting SNPs and large genetic diversity within the population. We find that one mechanism by which these naturally occurring SNPs affect animal size is via regulation of TOR activity. Astoundingly, the fly sorting was able to select for very specific traits, because TOR activity was increased in big flies compared to small flies specifically during larval stages of development when animal growth is taking place. Furthermore, we found little evidence for modulation of Yorkie activity in big versus small flies, suggesting that modulation of TOR activity might be a more common mechanism by which animal size can vary in nature, compared to modulation of Yorkie activity.

We performed whole-genome sequencing of the big and small fly populations and identified a very large number of SNPs that were differentially enriched in the two populations – circa 7000 using stringent cut-offs – including non-synonymous changes in the coding sequence of 279 genes. By screening these for effects on TOR-C1 signaling in cell culture, we identified five novel regulators of TOR-C1 which will likely be interesting genes to study in the future. For the remaining 274 genes, a very large number of assays would need to be performed to test if they are size regulators. They could be affecting a number of different signaling pathways and metabolic networks, in a number of different tissues in the animal. Therefore it is difficult to formally exclude them as size regulators. For this reason, we turned to a stochastic computer simulation of the process to suggest how many of them are likely to be true positives. This work revealed that the majority of these differentially selected SNPs can be expected to be unrelated to size regulation, selected by chance due to random drift within the population. This is an important insight, especially since it is difficult to exclude these SNPs experimentally, and it is tempting to believe they are all true.

An important output of the simulations is an understanding of what experimental setup conditions are necessary for such an approach. We hope this information will constitute a useful resource for the community. One parameter that is clearly important is the number of organisms sorted at each generation. At least for a genome size of Drosophila, a minimum of 800 flies per population per generation is necessary (Figure 6), meaning that novel high throughput methods for selecting flies will be necessary. We sorted 600 flies at each generation based on wing length (150 flies of each sex for each big and small population), which was roughly the maximum of what was feasible by hand, indicating that automated size determination and sorting will be required to improve this setup for the future. An additional parameter that can be manipulated is the complexity of the starting population. Reducing the number of SNPs in the starting population increases the chances of obtaining meaningful data at the end. This can be done, for instance, by some in-breeding, as long as enough phenotypic variation remains for subsequent selection. Lastly, performing the experiment in biological replicates and looking at the data that overlaps does not seem a superior strategy than simply increasing the numbers of animals sorted at each generation.

Although these simulations were done for a particular sort-and-sequence setup - sorting over multiple generations for a given trait - similar logic will likely apply to other studies. For instance, cancer develops through clonal expansion of cells which accumulate mutations over time, some of which influence cancer development, and some of which do not. Sequencing the end population of cells does a good job at identifying the differences compared to matched non-cancer tissue, however identifying which of these mutations is biologically meaningful is a problem well known to members of the field. Likewise genome-wide association studies (GWAS) are similar to the setup employed in this study, except that only one round of selection is performed and individuals are sequenced separately. (Since in our case we selected SNPs that were entirely polarized in the two populations, sequencing the population or sequencing individuals separately yields the same result.) Also for GWAS studies, due to similar types of issues, statistical thresholds are placed very stringently so that only the SNPs with the largest magnitude effects can be identified. Perhaps, similar types of simulations as the ones described here might help, although the parameters for the simulations, such as how many cells are being selected that lead to a tumor, and what the mutation rate is, are difficult to obtain. We hope the data presented here will help to set-up sort-and-sequence strategies to successfully exploit this promising technology in the future.

## Experimental Procedures

### Fly stocks and antibodies

w^1118^ and Oregon-R were obtained from the Bloomington stock collection. Anti phospho-S6K(Thr398), phospho-4E-BP, phospho-Akt, and total-Akt were from Cell Signaling. Anti total-S6K (Hahn *et al*. 2010), total-4EBP (Teleman *et al*. 2005), phospho-Yki and total Yki (Dong *et al.* 2007), anti-tubulin (Developmental Studies Hybridoma Bank).

### Physiological measurements

All animals were grown under controlled density conditions of 60 animals per vial. Size and lipid quantifications were performed as previously described (Teleman 2010). Adult wing cell size was quantified by measuring area per trichome.

### Cell culture and dsRNA treatment

S2 cells were cultured in Express-Five serum-free medium (Invitrogen). After 5 days of treatment with 12μg/mL dsRNA, immunoblotting was performed on the cell lysates to detect phospho-S6K(Thr398) and total-S6K using the antibodies described above.

### Sequencing, bioinformatics and data analysis

Libraries were generated using the Applied Biosystems SOLiD 3 system standard fragment protocol and sequenced on the SOLiD 3 system. Base calling and quality scoring were done during SOLiD primary analysis. csfasta and qual files of sequencing reads were retrieved from the sequencing machine. Reads were mapped to the Drosophila melanogaster genome using the MapReads algorithm, part of the Corona Lite Pipeline supplied by Applied Biosystems (CoronaLite_v4.0 release 2.0). The genome multi-fasta file was downloaded from the UCSC genome browser (April 2006 Assembly [dm3, BDGP Release 5]). The reference genomes were validated and double encoded as part of the corona lite SNP calling pipeline. The genome coverage for each sample was calculated and a consensus sequence determined using Corona Lite pipeline (McKernan *et al*. 2009). Averaging across all samples, 51 million reads were sequenced per sample, with 14 million reads mapping to unique starting positions, totaling 63% of the genome coverage, with an accuracy of 99%. Only “valid adjacent color changes” where considered for SNP analysis.

For SNP calling, two measures were calculated for each chromosomal position in the ‘big’ and ‘small’ fly populations: 1) the difference in nucleotide frequency between the two populations quantified as the geometric distance in 4D space: (%A_big_-%A_small_)^2^+(%C_big_-%C_small_)^2^+(%G_big_-%G_small_)^2^+(%T_big_-%T_small_)^2^. 2) A p-score for the significance of this difference calculated using a binomial distribution probability, bonferroni-adjusted for multiple testing. First, the nucleotide frequencies observed in the ‘small’ fly population were used to derive an underlying frequency distribution that was used to calculate the probability of sampling the nucleotide frequencies observed in the ‘big’ population using a binomial distribution. To be conservative and to correct for low sequence coverage in the ‘small’ population, the underlying nucleotide distribution at any one position was initially assumed to be 25%A/25%C/25%G/25%T, with each sequence read skewing this distribution: %A_underlying distribution_=(0.25+A_small_)/(1+A_small_+C_small_+G_small_+T_small_) where A_small_ is the number of times A was read at that position in the small population, and equivalently for the other nucleotides. Hence, high sequence coverage leads to an underlying distribution almost equal to the one observed by sequencing, whereas low sequence coverage is skewed towards equal representation of the four nucleotides,thereby reducing the significance of observing any particular nucleotide distribution in the big flies. Secondly, the equivalent probability of observing the ‘small’ frequencies based on an underlying ‘big’ distribution was also calculated. Third, to be conservative, the less-significant of the two p-scores was retained, and Bonferroni-corrected for the number of nucleotides in the genome. For this study, a stringent cut-off of difference=2 and p-score=0 was used.

### Stochastic Simulation

The full simulation code in the C programming language is available in Supplemental Material. The in silico simulation starts with an initial population of flies (Figure 5A). Each fly contains 3 chromosomes as in *Drosophila melanogaster.* (Chromosome 4 is very small and was therefore ignored for simplicity). Along the chromosomes, SNP positions were randomly specified. Each SNP position could be filled with one of two possible nucleotides (for instance, ‘A’ or ‘C’), to reflect that most SNPs in a population are binary (i.e. one usually finds 2 of the 4 possible nucleotides at that position), and for the sake of simplicity. A random subset of these SNPs was chosen to quantitatively affect animal size (or any quantitative trait of interest) whereas the majority of SNP positions were considered neutral. The SNP positions affecting size were randomly chosen to be either recessive (i.e. they only affect animal size when homozygous in the animal) or dominant (in which case each of the two alleles contributes additively to the quantitative trait, so that heterozygous animals experience half the effect of homozygous animals for that position). The quantitative strength of the contribution towards total animal size for each ‘size SNP’ was chosen randomly within a 10-fold dynamic range, reflecting that not all SNPs are expected to have equally strong effects on animal size. Once this genetic landscape was established, each starting fly within the population was given a random genotype. The size of each animal was calculated, and the larger 50% or smaller 50% of the populations retained for breeding to create the next generation. The genotype of each individual in the next generation was compiled by randomly choosing one father and one mother from the breeding animals. Meiotic cross-overs between chromosomes were allowed to occur at random positions within the female germline but not the male germline. The size of each resulting animal was calculated and the selection process was reiterated for a given number of generations. Finally, the genetic makeup of the final population was recorded, including the allele frequency for each SNP in the ‘big’ and ‘small’ populations, thereby identifying the SNPs that were differentially represented in the two populations. The simulation allows for all of the above-mentioned parameters to be varied.

## Acknowledgements

This work was supported by a Helmholtz Young Investigator Grant and an ERC Starting Grant to A.A.T. and an ERC Advanced Grant to MB. Antibodies were obtained from the Developmental Studies Hybridoma Bank developed under the auspices of the NICHD and maintained by The University of Iowa.

## Supplemental Material

### C code of computer simulation

~~~
#include <Carbon/Carbon.h>
#include <Stdio.h>
#include <Stdlib.h>
#include <Math.h>

int main (void){

       #define NUM_BREAK 7 // number of meiotic recombination breakpoints
genome-wide (ie 5x value per chr arm)
       #define SIZE_GENES 300 // number of genes affecting size in the
whole genome
       #define SNPS_TOT 35000 // number of SNPs per chromosome arm
       #define TOTAL_GENERATIONS 18
       #define FLIES 150

       FILE *out, *SNPs_out, *pop_avg_out, *fly_sizes_out,
*flies_selected_out, *sequencing_out, *initial_pop_out,
*historic_SNP_scores_out;
       int i, j, k, l, generations, parent_num, rand_chr, temp_int, dominant;
       int num_m_select, num_f_select, num_aa, num_het, num_hom;
       float pop_f_size_avg, pop_m_size_avg, rand;
       char filename[27]=“sequencing_out_gen_00.txt”;
       int a_big;
       float *hist_snp_score;
       CFAbsoluteTime time;

       //To pick flies randomly
       int num_list[FLIES/2];
       int flag;

       int breaks[NUM_BREAK];

       struct SNP{
              int pos;
              float het;
              float hom;
       };

       struct animal{
              char chr1[SNPS_TOT*5];
              char chr2[SNPS_TOT*5];
              float size;
       };

       struct SNP SNPs[SNPS_TOT*5];
       struct animal *pop_f[FLIES], *pop_m[FLIES], *pop_s_f[FLIES],
*pop_s_m[FLIES], *pop_b_f[FLIES], *pop_b_m[FLIES];

       /* OPEN FILES & ALLOCATE MEMORY */

       if(NULL == (out = fopen(“output.txt”, “w”)) ){
       fprintf(out, “Error opening output file: \“output.txt\”! \n”);
       exit(1);
       }

       if(NULL == (SNPs_out = fopen(“SNPs_output.txt”, “w”)) ){
       fprintf(out,“Error opening output file: \“output.txt\”! \n”);
       exit(1);
       }
       if(NULL == (pop_avg_out = fopen(“pop_size_average_output.txt”,“w”)) ){
       fprintf(out,“Error opening output file: \“output.txt\”! \n”);
       exit(1);
       }
       if(NULL == (fly_sizes_out = fopen(“fly_sizes_output.txt”,“w”)) ){
       fprintf(out,“Error opening output file: \“output.txt\”! \n”);
       exit(1);
       }
       if(NULL == (flies_selected_out =
fopen(“flies_selected_output.txt”,“w”)) ){
       fprintf(out,“Error opening output file: \“output.txt\”! \n”);
       exit(1);
       }
       if(NULL == (historic_SNP_scores_out=
fopen(“historic_SNP_scores_out.txt”,“w”)) ){
       fprintf(out,“Error opening output file: \“output.txt\”! \n”);
       exit(1);
       }

       if((hist_snp_score =
malloc(sizeof(float)*SNPS_TOT*5*(TOTAL_GENERATIONS+1))) == NULL){
       fprintf(out,“Error allocating memory for protein! \n”);
              exit(1);
       }

       for(i=0; i<FLIES; i++){
              if((pop_f[i] = malloc(sizeof(struct animal))) == NULL){
                     fprintf(out,“Error allocating memory for pop_f\n”);
                     exit(1);
              }
              if((pop_m[i] = malloc(sizeof(struct animal))) == NULL){
                     fprintf(out,“Error allocating memory for pop_m\n”);
                     exit(1);
              }
              if((pop_s_f[i] = malloc(sizeof(struct animal))) == NULL){
                     fprintf(out,“Error allocating memory for pop_s_f\n”);
                     exit(1);
              }
              if((pop_s_m[i] = malloc(sizeof(struct animal))) == NULL){
                     fprintf(out,“Error allocating memory for pop_s_m\n”);
                     exit(1);
              }
              if((pop_b_f[i] = malloc(sizeof(struct animal))) == NULL){
                     fprintf(out,“Error allocating memory for pop_b_f\n”);
                     exit(1);
              }
              if((pop_b_m[i] = malloc(sizeof(struct animal))) == NULL){
                     fprintf(out,“Error allocating memory for pop_b_m\n”);
                     exit(1);
              }
       }

       /* INITIALIZE */

       // Random seed for random() generator
       time=CFAbsoluteTimeGetCurrent();
       srandom(time);

       // Set positions of SNPs

       for(i=0; i<(SNPS_TOT*5); i++){
              SNPs[i].pos = random()/50;
              SNPs[i].het=0;
              SNPs[i].hom=0;
       }

       // Sort SNP positions in ascending order
       for(k=0; k<(SNPS_TOT*5); k++){
              for(l=k+1; l<(SNPS_TOT*5); l++){
                     if(SNPs[l].pos < SNPs[k].pos){
                            temp_int = SNPs[k].pos;
                            SNPs[k].pos = SNPs[l].pos;
                            SNPs[l].pos = temp_int;
                     }
              }
       }

       /* Set effect of each SNP on animal size. Assume two alleles are
possible: a and A. If the SNP is dominant,
        AA is larger, Aa is also larger (but not as much as AA, assuming it’s
a quantitative trait and it sums) and aa is smaller.
        If the SNP is recessive, AA is larger whereas Aa and aa are smaller.
        */

       i=0;
       while(i<SIZE_GENES){
              j=( ((float)random()) /2147483648 * SNPS_TOT * 5);
              if(SNPs[j].hom == 0){
                     i +=1;
                     dominant = random()/1073741824;
                     // Allow 10-fold differences in size effect from 0.005
to 0.05
                     SNPs[j].hom=0.005+(((float)random())/2147483647)*0.045;
                     if(dominant == 1){
                           SNPs[j].het = SNPs[j].hom/2;
                     }
              }
       }

       // Print out SNPs and their size effects

       fprintf(SNPs_out,“SNP_numb\tposition\thet_score\thom_score\n”);
       for(i=0; i<SNPS_TOT; i++){
              fprintf(SNPs_out,“ChrX-%i\t%i\t%f\t%f\n”,i, SNPs[i].pos,
SNPs[i].het, SNPs[i].hom);
       }
       for(i=SNPS_TOT; i<(SNPS_TOT*3); i++){
              fprintf(SNPs_out, “Chr2-%i\t%i\t%f\t%f\n”, i, SNPs[i].pos,
SNPs[i].het, SNPs[i].hom);
       }
       for(i=(SNPS_TOT*3); i<(SNPS_TOT*5); i++){
              fprintf(SNPs_out,“Chr3-%i\t%i\t%f\t%f\n”, i, SNPs[i].pos,
SNPs[i].het, SNPs[i].hom);
       }

       /* SET STARTING POPULATION */
       for(i=0; i<FLIES; i++){

             // Set genotype of the animal
             for(j=0; j<(SNPS_TOT*5); j++){
                    pop_s_f[i]->chr1[j]=pop_b_f[i]-
>chr1[j]=random()/1073741824;
                    pop_s_f[i]->chr2[j]=pop_b_f[i]-
>chr2[j]=random()/1073741824;
                    pop_s_m[i]->chr1[j]=pop_b_m[i]-
>chr1[j]=random()/1073741824;
                    pop_s_m[i]->chr2[j]=pop_b_m[i]-
>chr2[j]=random()/1073741824;
             }

             // Calculate size of the animals
             pop_s_f[i]->size = pop_b_f[i]->size = 0;
             pop_s_m[i]->size = pop_b_m[i]->size = 0;

             for(j=0; j<(SNPS_TOT*5); j++){
                    if(SNPs[j].hom != 0){
                           if( (pop_s_f[i]->chr1[j]+pop_s_f[i]->chr2[j])==2
){
                                  pop_s_f[i]->size += SNPs[j].hom;
                                  pop_b_f[i]->size += SNPs[j].hom;
                           } else if((pop_s_f[i]->chr1[j]+pop_s_f[i]-
>chr2[j])==1 ){
                                  pop_s_f[i]->size +=SNPs[j].het;
                                  pop_b_f[i]->size +=SNPs[j].het;
                           }
                           if(SNPs[j].hom != 0){
                                 if( (pop_s_m[i]->chr1[j]+pop_s_m[i]-
>chr2[j]) ==2 ){
                                        pop_s_m[i]->size += SNPs[j].hom;
                                        pop_b_m[i]->size += SNPs[j].hom;
                                 } else if((pop_s_m[i]->chr1[j]+pop_s_m[i]-
>chr2[j]) ==1 ){
                                        pop_s_m[i]->size +=SNPs[j].het;                                     

                                        pop_b_m[i]->size +=SNPs[j].het;                                     

                                 }
                           }
                    }
              }
       }

       /*****OUTPUT GENOTYPES OF INITIAL POPULATION*****/
       if(NULL == (initial_pop_out =
fopen(“initial_population_genotypes_output.txt”,“w”)) ){
       fprintf(out,“Error opening output file: \“output.txt\”! \n”);
       exit(1);
       }

       fprintf(initial_pop_out,“SNP\t”);
       for(i=0; i<FLIES; i++){

       fprintf(initial_pop_out,“Female_%i_chr1\tFemale_%i_chr2\t”,i,i);
       }
       fprintf(initial_pop_out,“\n”);

       for(j=0; j<SNPS_TOT; j++){
             if(SNPs[j].hom != 0) fprintf(initial_pop_out,“*”);
             fprintf(initial_pop_out, “ChrX-%i\t”,j);
             for(i=0; i<FLIES; i++){
                    fprintf(initial_pop_out,“%c\t%c\t”, ‘0’ +pop_b_f[i]-
>chr1[j],‘0’+pop_b_f[i]->chr2[j]);
             }
             fprintf(initial_pop_out,“\n”);
       }
       for(j=SNPS_TOT; j<(3*SNPS_TOT); j++){
              if(SNPs[j].hom != 0) fprintf(initial_pop_out,“*”);
              fprintf(initial_pop_out, “Chr2-%i\t”,j);

              for(i=0; i<FLIES; i++){
                     fprintf(initial_pop_out, “%c\t%c\t”, ‘0’+pop_b_f[i]-
>chr1[j],‘0’+pop_b_f[i]->chr2[j]);
              }
              fprintf(initial_pop_out,“\n”);
       }
       for(j=SNPS_TOT;(3*SNPS_TOT); j<(5*SNPS_TOT); j++){
              if(SNPs[j].hom != 0) fprintf(initial_pop_out, “*”);
              fprintf(initial_pop_out,“Chr5-%i\t”,j);
              for(i=0; i<FLIES; i++){
                     fprintf(initial_pop_out,“%c\t%c\t”,‘0’+pop_b_f[i]-
>chr1[j],‘0’+pop_b_f[i]->chr2[j]);
              }
              fprintf(initial_pop_out,“\n”);
       }

       fclose(initial_pop_out);
       /*****OUTPUT GENOTYPES OF INITIAL POPULATION*****/

      /***** OUTPUT SEQUENCING RESULTS INITIAL POPULATION*****/

      if(NULL == (sequencing_out =
fopen(“sequencing_out_initial_pop.txt”,“w”)) ){
             fprintf(out,“Error opening output file: \“output.txt\”! \n”);
             exit(1);
       }

       fprintf(sequencing_out,“SNP\tBig a/a\tBig a/A\tBig A/A\tBig total a\tBig total A\tSmall a/a\tSmall a/A\tSmall A/A\tSmall total a\tSmall total A\tSNP score\n”);

       for(j=0; j<(5*SNPS_TOT); j++){
              num_aa=num_het=num_hom=0;
              for(i=0; i<FLIES; i++){
                     if( (pop_b_f[i]->chr1[j]+pop_b_f[i]->chr2[j])==2 ){
                            num_hom += 1;
                     } else if( (pop_b_f[i]->chr1[j]+pop_b_f[i]->chr2[j])==1
){
                            num_het += 1;
                     } else {
                            num_aa += 1;
                     }
              }

              fprintf(sequencing_out,“%i\t%i\t%i\t%i\t%i\t%i\t”,j,num_aa, num_het, num_hom,2*num_aa+num_het,num_het+2*num_hom);

              a_big = num_het + 2*num_hom;

              num_aa=num_het=num_hom=0;
              for(i=0; i<FLIES; i++){
                     if( (pop_s_f[i]->chr1[j]+pop_s_f[i]->chr2[j])==2 ){
                            num_hom += 1;
                     } else if( (pop_s_f[i]->chr1[j]+pop_s_f[i]->chr2[j])==1
){
                            num_het += 1;
                     } else {
                            num_aa += 1;
                     }
              }
              fprintf(sequencing_out,“%i\t%i\t%i\t%i\t%i\t%f\n”,num_aa,
num_het, num_hom,2*num_aa+num_het,num_het+2*num_hom, ((float)(a_big-
(num_het+2*num_hom)))/FLIES/2);
              // SNP score = # chromosomes in big popoulation that have the A
allele, minus # chr from small pop, divided by (2x # of flies- ie max
possible chromosomes)
              hist_snp_score[j]= ((float)(a_big-
(num_het+2*num_hom)))/FLIES/2;
       }
       fclose(sequencing_out);

       /***** OUTPUT SEQUENCING RESULTS INITIAL POPULATION*****/

       /*****OUTPUT FLY SIZES*****/
       fprintf(fly_sizes_out,“Initial females\n”);
       for(i=0; i<FLIES; i++){
              fprintf(fly_sizes_out,“%f\n”,pop_b_f[i]->size);
       }

       /*****OUTPUT FLY SIZES*****/

       /* EXECUTE SORTing*/

       srandom(time);

       fprintf(flies_selected_out,“Generation\tBig females\tBig males\tSmall
females\tSmall males\n”);

       for(generations=0; generations<TOTAL_GENERATIONS; generations++){

              /*‐‐‐‐‐‐‐‐‐‐‐‐‐‐‐‐‐‐‐‐‐‐‐‐‐‐‐‐‐‐‐‐‐‐‐‐‐‐‐‐‐‐BIG FLIES‐‐‐‐‐‐-
‐‐‐‐‐‐‐‐‐‐‐‐‐‐‐‐‐‐‐‐‐‐‐‐*/
              //Calc population average BIG FLIES
              pop_f_size_avg=0;
              pop_m_size_avg=0;

              for(i=0; i<FLIES; i++){
                     pop_f_size_avg += pop_b_f[i]->size;
                     pop_m_size_avg += pop_b_m[i]->size;
              }
              pop_f_size_avg = pop_f_size_avg / FLIES;
              pop_m_size_avg = pop_m_size_avg / FLIES;

              /*****OUTPUT POPULATION AVERAGE SIZE*****/
              fprintf(pop_avg_out,“%i\t%f\t”,generations,pop_f_size_avg);
              /*****OUTPUT POPULATION AVERAGE SIZE*****/

              //Sort BIG FLIES

              num_m_select = 0;
              num_f_select = 0;

              for(i=0; i<FLIES; i++){
                     if((pop_b_f[i]->size >= pop_f_size_avg) && (num_f_select
< FLIES) ){
                           for(j=0; j<(5*SNPS_TOT); j++) pop_f[num_f_select]-
>chr1[j]=pop_b_f[i]->chr1[j];
                           for(j=0; j<(5*SNPS_TOT); j++) pop_f[num_f_select]-
>chr2[j]=pop_b_f[i]->chr2[j];
                           for(j=0; j<(5*SNPS_TOT); j++) pop_f[num_f_select]-
>size=pop_b_f[i]->size;
                           num_f_select += 1;
                     }
              }
              for(i=0; i<FLIES; i++){
                     if( (pop_b_m[i]->size >= pop_m_size_avg) &&
(num_m_select < FLIES) ){
                            for(j=0; j<(5*SNPS_TOT); j++) pop_m[num_m_select]-
>chr1[j]=pop_b_m[i]->chr1[j];
                            for(j=0; j<(5*SNPS_TOT); j++) pop_m[num_m_select]-
>chr2[j]=pop_b_m[i]->chr2[j];
                            for(j=0; j<(5*SNPS_TOT); j++) pop_m[num_m_select]-
>size=pop_b_m[i]->size;
                            num_m_select += 1;
                     }

              }

              /*****OUTPUT FLY SELECTION NUMBERS*****/

       fprintf(flies_selected_out,“%i\t%i\t%i\t”,generations,num_f_select,num
_m_select);
              /*****OUTPUT FLY SELECTION NUMBERS*****/

              //Generate next generation BIG FLIES

              //Pick all male chromosomes (no recombination) -> chr1
              for(i=0; i<FLIES; i++){
                     parent_num=( ((float)random())/2147483648
)*num_m_select;
                     if(parent_num==num_m_select) fprintf(out,“ERROR !!!
parent_num=num_m_select at Generation %i probably due to population reaching
homogeneity\n”,generations);

                     // Chr X
                     rand_chr = random()/1073741824;
                     for(j=0; j<SNPS_TOT; j++){
                            if(rand_chr){
                                   pop_b_f[i]->chr1[j] = pop_m[parent_num]-
>chr1[j];
                            } else {
                                   pop_b_f[i]->chr1[j] = pop_m[parent_num]-
>chr2[j];
                            }
                     }

                     // Chr 2
                     rand_chr = random()/1073741824;
                     for(j=SNPS_TOT; j<(3*SNPS_TOT); j++){
                            if(rand_chr){
                                   pop_b_f[i]->chr1[j] = pop_m[parent_num]-
>chr1[j];
                            } else {
                                   pop_b_f[i]->chr1[j] = pop_m[parent_num]-
>chr2[j];
                            }
                     }

                     //Chr 3
                     rand_chr = random()/1073741824;
                     for(j=(3*SNPS_TOT); j<(5*SNPS_TOT); j++){
                            if(rand_chr){
                                   pop_b_f[i]->chr1[j] = pop_m[parent_num]-
>chr1[j];
                            } else {
                                   pop_b_f[i]->chr1[j] = pop_m[parent_num]-
>chr2[j];
                            }
                     }
              }

              for(i=0; i<FLIES; i++){
                     parent_num=( ((float)random()) /2147483648
)*num_m_select;
                     if(parent_num==num_m_select) fprintf(out,“ERROR !!!
parent_num=num_m_select at Generation %i probably due to population reaching
homogeneity %i\n”,generations);

                     // Chr X
                     rand_chr = random()/1073741824;

                     for(j=0; j<SNPS_TOT; j++){
                            if(rand_chr){
                                   pop_b_m[i]->chr1[j] = pop_m[parent_num]-
>chr1[j];
                            } else {
                                   pop_b_m[i]->chr1[j] = pop_m[parent_num]-
>chr2[j];
                            }
                     }

                     // Chr 2
                     rand_chr = random()/1073741824;
                     for(j=SNPS_TOT; j<(3*SNPS_TOT); j++){
                            if(rand_chr){
                                   pop_b_m[i]->chr1[j] = pop_m[parent_num]-
>chr1[j];
                            } else {
                                   pop_b_m[i]->chr1[j] = pop_m[parent_num]-
>chr2[j];
                            }
                     }

                     //Chr 3
                     rand_chr = random()/1073741824;
                     for(j=(3*SNPS_TOT); j<(5*SNPS_TOT); j++){
                            if(rand_chr){
                                   pop_b_m[i]->chr1[j] = pop_m[parent_num]-
>chr1[j];
                            } else {
                                   pop_b_m[i]->chr1[j] = pop_m[parent_num]-
>chr2[j];
                            }
                     }
              }

              //Pick all female chromosomes -> chr2
              for(i=0; i<FLIES; i++){
                     for(k=0; k<NUM_BREAK; k++){
                            breaks[k]=random()/50;
                            // check by chance that the same breakpoint wasn’t
already selected on this chromosome (ie want NUM_BREAK different
breakpoints)
                            for(l=0; l<k; l++){
                                   if(breaks[k]==breaks[l]){
                                          k -= 1;
                                          l=k;
                                   }
                            }
                     }
                     // sort the breakpoints in numberical order
                     for(k=0; k<NUM_BREAK; k++){
                            for(l=k+1; l<NUM_BREAK; l++){
                                   if(breaks[l] < breaks[k]){
                                          temp_int = breaks[k];
                                          breaks[k] = breaks[l];
                                          breaks[l] = temp_int;
                                   }
                            }
                     }
                     parent_num=( ((float)random()) /2147483648
)*num_f_select;
                     if(parent_num==num_f_select) fprintf(out,“ERROR !!!
parent_num=num_f_select at generation %i\n”,generations);

                     k=0;

                     // Chr X
                     rand_chr = random()/1073741824;
                     for(j=0; j<SNPS_TOT; j++){
                            // switch chromosome after passing a break
                            if(SNPs[j].pos > breaks[k]){
                                   rand_chr = 1-rand_chr;
                                   k++;
                            }
                            if(rand_chr){
                                   pop_b_f[i]->chr2[j] = pop_f[parent_num]-
>chr1[j];
                            } else {
                                   pop_b_f[i]->chr2[j] = pop_f[parent_num]-
>chr2[j];
                            }
                     }
                     // Chr 2
                     rand_chr = random()/1073741824;
                     for(j=SNPS_TOT; j<(3*SNPS_TOT); j++){
                            // switch chromosome after passing a break                                 
                            if(SNPs[j].pos > breaks[k]){                                        
                                   rand_chr = 1-rand_chr;
                                   k++;
                            }
                            if(rand_chr){
                                   pop_b_f[i]->chr2[j] = pop_f[parent_num]-
>chr1[j];
                            } else {
                                   pop_b_f[i]->chr2[j] = pop_f[parent_num]-
>chr2[j];
                            }
                     }
                     // Chr 3
                     rand_chr = random()/1073741824;
                     for(j=(3*SNPS_TOT); j<(5*SNPS_TOT); j++){
                            // switch chromosome after passing a break
                            if(SNPs[j].pos > breaks[k]){
                                   rand_chr = 1-rand_chr;
                                   k++;
                            }
                            if(rand_chr){
                                   pop_b_f[i]->chr2[j] = pop_f[parent_num]-
>chr1[j];
                            } else {
                                   pop_b_f[i]->chr2[j] = pop_f[parent_num]-
>chr2[j];
                            }
                     }
              }
              for(i=0; i<FLIES; i++){
                     for(k=0; k<NUM_BREAK; k++){
                            breaks[k]=random()/50;
                            // check by chance that the same breakpoint wasn’t
already selected on this chromosome (ie want NUM_BREAK different
breakpoints)
                            for(l=0; l<k; l++){
                                   if(breaks[k]==breaks[l]){
                                          k -= 1;
                                          l=k;
                                   }
                            }
                     }
                     // sort the breakpoints in numberical order
                     for(k=0; k<NUM_BREAK; k++){
                            for(l=k+1; l<NUM_BREAK; l++){
                                   if(breaks[l] < breaks[k]){
                                         temp_int = breaks[k];
                                         breaks[k]= breaks[l];
                                         breaks[l]=temp_int;
                                   }
                            }
                     }
                     parent_num=( ((float)random()) /2147483648
)*num_f_select;
                     if(parent_num==num_f_select) fprintf(out,“ERROR !!!
parent_num=num_f_select at generation %i\n”,generations);

                     k=0;

                     // Chr X
                     rand_chr = random()/1073741824;
                     for(j=0; j<SNPS_TOT; j++){
                            // switch chromosome after passing a break
                            if(SNPs[j].pos > breaks[k]){
                                   rand_chr = 1-rand_chr;
                                   k++;
                            }
                            if(rand_chr){
                                   pop_b_m[i]->chr2[j] = pop_f[parent_num]-
>chr1[j];
                            } else {
                                   pop_b_m[i]->chr2[j] = pop_f[parent_num]-
>chr2[j];
                            }
                     }
                     // Chr 2
                     rand_chr = random()/1073741824;
                     for(j=SNPS_TOT; j<(3*SNPS_TOT); j++){
                            // switch chromosome after passing a break
                            if(SNPs[j].pos > breaks[k]){
                                   rand_chr = 1-rand_chr;
                                   k++;
                            }
                            if(rand_chr){
                                   pop_b_m[i]->chr2[j] = pop_f[parent_num]-
>chr1[j];
                            } else {
                                   pop_b_m[i]->chr2[j] = pop_f[parent_num]-
>chr2[j];
                            }
                     }
                     // Chr 3
                     rand_chr = random()/1073741824;
                     for(j=(3*SNPS_TOT); j<(5*SNPS_TOT); j++){
                            // switch chromosome after passing a break
                            if(SNPs[j].pos > breaks[k]){
                                   rand_chr = 1-rand_chr;
                                   k++;
                            }
                            if(rand_chr){
                                   pop_b_m[i]->chr2[j] = pop_f[parent_num]-
>chr1[j];
                            } else {
                                   pop_b_m[i]->chr2[j] = pop_f[parent_num]-
>chr2[j];
                            }
                     }
              }

              /*‐‐‐‐‐‐‐‐‐‐‐‐‐‐‐‐‐‐‐‐‐‐‐‐‐‐‐‐‐‐‐‐‐‐‐‐‐‐‐‐ SMALL FLIES ‐‐‐‐‐‐‐‐‐‐
‐‐‐‐‐‐‐‐‐‐‐‐‐‐‐‐‐‐‐‐‐‐-*/
              //Calc population average SMALL FLIES
              pop_f_size_avg=0;
              pop_m_size_avg=0;

              for(i=0; i<FLIES; i++){
                      pop_f_size_avg += pop_s_f[i]->size;
                      pop_m_size_avg += pop_s_m[i]->size;
              }
              pop_f_size_avg = pop_f_size_avg / FLIES;
              pop_m_size_avg = pop_m_size_avg / FLIES;

              /*****OUTPUT POPULATION AVERAGE SIZE*****/
              fprintf(pop_avg_out,“%f\n”,pop_f_size_avg);
              /*****OUTPUT POPULATION AVERAGE SIZE*****/

              //Sort SMALL FLIES
        
              num_m_select = 0;
              num_f_select = 0;

              for(i=0; i<FLIES; i++){
                     if((pop_s_f[i]->size <= pop_f_size_avg) && (num_f_select
< FLIES) ){
                            for(j=0; j<(5*SNPS_TOT); j++) pop_f[num_f_select]-
>chr1[j]=pop_s_f[i]->chr1[j];
                            for(j=0; j<(5*SNPS_TOT); j++) pop_f[num_f_select]-
>chr2[j]=pop_s_f[i]->chr2[j];
                            for(j=0; j<(5*SNPS_TOT); j++) pop_f[num_f_select]-
>size=pop_s_f[i]->size;
                            num_f_select += 1;
                     }
              }
              for(i=0; i<FLIES; i++){
                     if( (pop_s_m[i]->size <= pop_m_size_avg) &&
(num_m_select < FLIES) ){
                             for(j=0; j<(5*SNPS_TOT); j++) pop_m[num_m_select]-
>chr1[j]=pop_s_m[i]->chr1[j];
                             for(j=0; j<(5*SNPS_TOT); j++) pop_m[num_m_select]-
>chr2[j]=pop_s_m[i]->chr2[j];
                             for(j=0; j<(5*SNPS_TOT); j++) pop_m[num_m_select]-
>size=pop_s_m[i]->size;
                             num_m_select += 1;
                     }
             }

             /*****OUTPUT FLY SELECTION NUMBERS*****/

       fprintf(flies_selected_out,“%i\t%i\n”,num_f_select,num_m_select);
             /*****OUTPUT FLY SELECTION NUMBERS*****/

             //Generate next generation SMALL FLIES

             //Pick all male chromosomes (no recombination) -> chr1
             for(i=0; i<FLIES; i++){
                    parent_num=( ((float)random())/2147483648
)*num_m_select;
                    if(parent_num==num_m_select) fprintf(out,“ERROR !!!
parent_num=num_m_select at Generation %i probably due to population reaching
homogeneity\n”,generations);

                    // Chr X
                    rand_chr = random()/1073741824;
                    for(j=0; j<SNPS_TOT; j++){
                           if(rand_chr){
                                  pop_s_f[i]->chr1[j] = pop_m[parent_num]-
>chr1[j];
                           } else {

                                  pop_s_f[i]->chr1[j] = pop_m[parent_num]-
>chr2[j];
                           }
                    }

                    // Chr 2
                    rand_chr = random()/1073741824;
                    for(j=SNPS_TOT; j<(3*SNPS_TOT); j++){
                           if(rand_chr){
                                  pop_s_f[i]->chr1[j] = pop_m[parent_num]-
>chr1[j];
                           } else {
                                  pop_s_f[i]->chr1[j] = pop_m[parent_num]-
>chr2[j];
                           }
                    }

                    //Chr 3
                    rand_chr = random()/1073741824;
                    for(j=(3*SNPS_TOT); j<(5*SNPS_TOT); j++){
                           if(rand_chr){
                                  pop_s_f[i]->chr1[j] = pop_m[parent_num]-
>chr1[j];
                           } else {
                                  pop_s_f[i]->chr1[j] = pop_m[parent_num]-
>chr2[j];
                           }
                    }
             }

             for(i=0; i<FLIES; i++){
                    parent_num=( ((float)random()) /2147483648
)*num_m_select;
                    if(parent_num==num_m_select) fprintf(out,“ERROR !!!
parent_num=num_m_select at Generation %i probably due to population reaching
homogeneity%i\n”,generations);

                    // Chr X
                    rand_chr = random()/1073741824;
                    for(j=0; j<SNPS_TOT; j++){
                           if(rand_chr){
                                  pop_s_m[i]->chr1[j] = pop_m[parent_num]-
>chr1[j];
                           } else {
                                  pop_s_m[i]->chr1[j] = pop_m[parent_num]-
>chr2[j];
                           }
                    }

                    // Chr 2
                    rand_chr = random()/1073741824;
                    for(j=SNPS_TOT; j<(3*SNPS_TOT); j++){
                           if(rand_chr){
                                  pop_s_m[i]->chr1[j] = pop_m[parent_num]-
>chr1[j];
                           } else {
                                  pop_s_m[i]->chr1[j] = pop_m[parent_num]-
>chr2[j];
                           }
                    }

                    //Chr 3
                    rand_chr = random()/1073741824;
                    for(j=(3*SNPS_TOT); j<(5*SNPS_TOT); j++){
                           if(rand_chr){
                                  pop_s_m[i]->chr1[j] = pop_m[parent_num]-
> chr1[j];
                           } else {

                                  pop_s_m[i]->chr1[j] = pop_m[parent_num]-
>chr2[j];
                           }
                    }
             }

             //Pick all female chromosomes -> chr2
             for(i=0; i<FLIES; i++){
                    for(k=0; k<NUM_BREAK; k++){
                           breaks[k]=random()/50;
                           // check by chance that the same breakpoint wasn’t
already selected on this chromosome (ie want NUM_BREAK different
breakpoints)
                           for(l=0; l<k; l++){
                                  if(breaks[k]==breaks[l]){
                                         k -= 1;
                                         l=k;
                                  }
                           }
                    }
                    // sort the breakpoints in numberical order
                    for(k=0; k<NUM_BREAK; k++){
                           for(l=k+1; l<NUM_BREAK; l++){
                                  if(breaks[l] < breaks[k]){
                                         temp_int = breaks[k];
                                         breaks[k] = breaks[l];
                                         breaks[l] = temp_int;
                                  }
                           }
                    }
                    parent_num=( ((float)random()) /2147483648
)*num_f_select;
                    if(parent_num==num_f_select) fprintf(out,“ERROR !!!
parent_num=num_f_select at generation %i\n“,generations);

                    k=0;

                    // Chr X
                    rand_chr = random()/1073741824;
                    for(j=0; j<SNPS_TOT; j++){
                           // switch chromosome after passing a break
                           if(SNPs[j].pos > breaks[k]){
                                  rand_chr = 1-rand_chr;
                                  k++;
                           }
                           if(rand_chr){
                                  pop_s_f[i]->chr2[j] = pop_f[parent_num]-
>chr1[j];
                           } else {
                                  pop_s_f[i]->chr2[j] = pop_f[parent_num]-
>chr2[j];
                           }
                    }
                    // Chr 2
                    rand_chr = random()/1073741824;
                    for(j=SNPS_TOT; j<(3*SNPS_TOT); j++){
                           // switch chromosome after passing a break
                           if(SNPs[j].pos > breaks[k]){
                                  rand_chr =1-rand_chr;
                                  k++;
                           }
                           if(rand_chr){
                                  pop_s_f[i]->chr2[j] = pop_f[parent_num]-
>chr1[j];
                           } else {

                                  pop_s_f[i]->chr2[j] = pop_f[parent_num]-
>chr2[j];
                           }
                    }
                    // Chr 3
                    rand_chr = random()/1073741824;
                    for(j=(3*SNPS_TOT); j<(5*SNPS_TOT); j++){
                           // switch chromosome after passing a break
                           if(SNPs[j].pos > breaks[k]){
                                  rand_chr = 1-rand_chr;
                                  k++;
                           }
                           if(rand_chr){
                                  pop_s_f[i]->chr2[j] = pop_f[parent_num]-
>chr1[j];
                           } else {
                                  pop_s_f[i]->chr2[j] = pop_f[parent_num]-
>chr2[j];
                           }
                     }
              }

              for(i=0; i<FLIES; i++){
                     for(k=0; k<NUM_BREAK; k++){
                            breaks[k]=random()/50;
                            // check by chance that the same breakpoint wasn’t
already selected on this chromosome (ie want NUM_BREAK different
breakpoints)
                            for(l=0; l<k; l++){
                                   if(breaks[k]==breaks[l]){
                                          k -= 1;
                                          l=k;
                                   }
                            }
                     }
                     // sort the breakpoints in numberical order
                     for(k=0; k<NUM_BREAK; k++){
                            for(l=k+1; l<NUM_BREAK; l++){
                                   if(breaks[l] < breaks[k]){
                                          temp_int = breaks[k];
                                          breaks[k] = breaks[l];
                                          breaks[l] = temp_int;
                                   }
                            }
                     }

                     parent_num=( ((float)random()) /2147483648
)*num_f_select;
                     if(parent_num==num_f_select) fprintf(out,“ERROR !!!
parent_num=num_f_select at generation %i\n”,generations);

                     k=0;

                     // Chr X
                     rand_chr = random()/1073741824;
                     for(j=0; j<SNPS_TOT; j++){
                            // switch chromosome after passing a break
                            if(SNPs[j].pos > breaks[k]){
                                   rand_chr = 1-rand_chr;
                                   k++;
                            }
                            if(rand_chr){
                                   pop_s_m[i]->chr2[j] = pop_f[parent_num]-
>chr1[j];
                            } else {
                                   pop_s_m[i]->chr2[j] = pop_f[parent_num]-
>chr2[j];
                            }
                     }
                     // Chr 2
                     rand_chr = random()/1073741824;
                     for(j=SNPS_TOT; j<(3*SNPS_TOT); j++){
                            // switch chromosome after passing a break
                            if(SNPs[j].pos > breaks[k]){
                                   rand_chr = 1-rand_chr;
                                   k++;
                             }
                             if(rand_chr){
                                    pop_s_m[i]->chr2[j] = pop_f[parent_num]-
>chr1[j];
                             } else {
                                    pop_s_m[i]->chr2[j] = pop_f[parent_num]-
>chr2[j];
                             }
                      }
                      // Chr 3
                      rand_chr = random()/1073741824;
                      for(j=(3*SNPS_TOT); j<(5*SNPS_TOT); j++){
                             // switch chromosome after passing a break
                             if(SNPs[j].pos > breaks[k]){
                                    rand_chr = 1-rand_chr;
                                    k++;
                             }
                             if(rand_chr){
                                    pop_s_m[i]->chr2[j] = pop_f[parent_num]-
>chr1[j];
                             } else {
                                    pop_s_m[i]->chr2[j] = pop_f[parent_num]-
>chr2[j];
                             }
                     }
              }

              /*‐‐‐‐‐‐‐‐‐‐‐‐‐‐‐‐‐‐‐‐‐‐‐‐‐‐‐‐‐‐‐‐‐‐‐‐  Calculate size of the
animals‐‐‐‐‐‐‐‐‐‐‐‐‐‐‐‐‐‐*/

              for(i=0; i<FLIES; i++){
                     pop_b_f[i]->size =0;
                     pop_b_m[i]->size =0;
                     pop_s_f[i]->size =0;
                     pop_s_m[i]->size =0;

                     for(j=0; j<(5*SNPS_TOT); j++){
                            if(SNPs[j].hom != 0){
                                   //Big females
                                   if( (pop_b_f[i]->chr1[j]+pop_b_f[i]-
>chr2[j])==2 ){
                                          pop_b_f[i]->size += SNPs[j].hom;
                                   } else if((pop_b_f[i]->chr1[j]+pop_b_f[i]-
>chr2[j])==1 ){
                                          pop_b_f[i]->size +=SNPs[j].het;
                                   }
                                   //Big males
                                   if( (pop_b_m[i]->chr1[j]+pop_b_m[i]-
>chr2[j])==2 ){
                                          pop_b_m[i]->size += SNPs[j].hom;
                                   } else if((pop_b_m[i]->chr1[j]+pop_b_m[i]-
>chr2[j])==1 ){
                                          pop_b_m[i]->size +=SNPs[j].het;
                                   }
                                   //Small females
                                   if( (pop_s_f[i]->chr1[j]+pop_s_f[i]-
>chr2[j])==2 ){
                                          pop_s_f[i]->size += SNPs[j].hom;
                                   } else if((pop_s_f[i]->chr1[j]+pop_s_f[i]-
>chr2[j])==1 ){
                                          pop_s_f[i]->size +=SNPs[j].het;
                                   }

                                   //Small males
                                   if( (pop_s_m[i]->chr1[j]+pop_s_m[i]-
>chr2[j])==2 ){
                                          pop_s_m[i]->size += SNPs[j].hom;
                                   } else if((pop_s_m[i]->chr1[j]+pop_s_m[i]-
>chr2[j])==1 ){
                                            pop_s_m[i]->size +=SNPs[j].het;
                                   }
                            }
                     }
              }

        /*****OUTPUT FLY SIZES*****/
        fprintf(fly_sizes_out,“Generation %i\n”,generations);
        fprintf(fly_sizes_out,“Big females\tSmall females\n”);
        for(i=0; i<FLIES; i++){
               fprintf(fly_sizes_out,“%f\t%f\n”,pop_b_f[i]-
>size,pop_s_f[i]->size);
        }
        /*****OUTPUT FLY SIZES*****/

        /***** OUTPUT SEQUENCING RESULTS *****/
        filename[19]=‘0’+(generations/10);
        filename[20]=‘0’+(generations%10);
        if(NULL == (sequencing_out = fopen(filename,“w”)) ){
               fprintf(out,“Error opening output file: \“output.txt\”!
\n”);
               exit(1);
        }

        fprintf(sequencing_out,“SNP\tBig a/a\tBig a/A\tBig A/A\tBig
total a\tBig total A\tSmall a/a\tSmall a/A\tSmall A/A\tSmall total a\tSmall
total A\tSNP score\n”);
        for(j=0; j<(5*SNPS_TOT); j++){
               num_aa=num_het=num_hom=0;
               for(i=0; i<FLIES; i++){
                      if( (pop_b_f[i]->chr1[j]+pop_b_f[i]->chr2[j])==2
){
                              num_hom += 1;
                      } else if( (pop_b_f[i]->chr1[j]+pop_b_f[i]-
>chr2[j])==1 ){
                              num_het += 1;
                      } else {
                              num_aa += 1;
                      }
               }
               if(SNPs[j].hom != 0) fprintf(sequencing_out,“*”);

     fprintf(sequencing_out,“%i\t%i\t%i\t%i\t%i\t%i\t”,j,num_aa, num_het,
num_hom,2*num_aa+num_het,num_het+2*num_hom);

               a_big = num_het + 2*num_hom;

               num_aa=num_het=num_hom=0;
               for(i=0; i<FLIES; i++){
                      if( (pop_s_f[i]->chr1[j]+pop_s_f[i]->chr2[j])==2
){
                              num_hom += 1;
                      } else if( (pop_s_f[i]->chr1[j]+pop_s_f[i]-
>chr2[j])==1 ){
                              num_het += 1;
                      } else {
                              num_aa += 1;
                      }
               }

     fprintf(sequencing_out,“%i\t%i\t%i\t%i\t%i\t%f\n”,num_aa, num_het,
num_hom,2*num_aa+num_het,num_het+2*num_hom, ((float)(a_big-
(num_het+2*num_hom)))/FLIES/2);
     hist_snp_score[j+(5*SNPS_TOT)*(generations+1)]=((float)(a_big-
(num_het+2*num_hom)))/FLIES/2;
               }
               fclose(sequencing_out);
               /***** OUTPUT SEQUENCING RESULTS *****/

     }

         /***** OUTPUT HISTORIC SNP SCORES *****/
         fprintf(historic_SNP_scores_out,“SNP\t”);
         for(k=0; k<(generations+1); k++) fprintf(historic_SNP_scores_out,“Generation %i\t”,k);
         fprintf(historic_SNP_scores_out,“\n”);
         for(j=0; j<(5*SNPS_TOT); j++){
                     if(SNPs[j].hom != 0) fprintf(historic_SNP_scores_out,“*”);
                     fprintf(historic_SNP_scores_out,“%i\t”,j);
                     for(k=0; k<(generations+1); k++){

       fprintf(historic_SNP_scores_out,“%f\t”,hist_snp_score[j+(5*SNPS_TOT)*k
]);
                     }
                     fprintf(historic_SNP_scores_out,“\n”);
       }

       /***** OUTPUT HISTORIC SNP SCORES *****/

       /* CLOSE AND EXIT */
       free(hist_snp_score);
       fclose(out);
       fclose(SNPs_out);
       fclose(pop_avg_out);
       fclose(fly_sizes_out);
       fclose(flies_selected_out);
       fclose(historic_SNP_scores_out);
       for(i=0; i<FLIES; i++){
                    free(pop_f[i]);
                    free(pop_m[i]);
                    free(pop_s_f[i]);
                    free(pop_s_m[i]);
                    free(pop_b_f[i]);
                    free(pop_b_m[i]);
       }

}
~~~

**Supplemental Table 1:** Positions of Differential SNPs.

| Chromosome | position | difference | BIG coverage | small coverage | p-score |
| --- | --- | --- | --- | --- | --- |
| 2L | 372959 | 2 | 5 | 19 | 0 |
| 2L | 529663 | 2 | 12 | 4 | 0 |
| 2L | 555027 | 2 | 5 | 12 | 0 |
| 2L | 674684 | 2 | 4 | 9 | 0 |
| 2L | 747998 | 2 | 5 | 15 | 0 |
| 2L | 760040 | 2 | 13 | 15 | 0 |
| 2L | 761871 | 2 | 6 | 16 | 0 |
| 2L | 907606 | 2 | 10 | 4 | 0 |
| 2L | 1316977 | 2 | 11 | 7 | 0 |
| 2L | 1633751 | 2 | 5 | 5 | 0 |
| 2L | 1852691 | 2 | 5 | 7 | 0 |
| 2L | 1955402 | 2 | 6 | 6 | 0 |
| 2L | 2126661 | 2 | 7 | 6 | 0 |
| 2L | 2204937 | 2 | 10 | 9 | 0 |
| 2L | 2254612 | 2 | 12 | 7 | 0 |
| 2L | 2273379 | 2 | 11 | 10 | 0 |
| 2L | 2379679 | 2 | 6 | 9 | 0 |
| 2L | 2381128 | 2 | 8 | 7 | 0 |
| 2L | 2406752 | 2 | 7 | 10 | 0 |
| 2L | 2420284 | 2 | 7 | 9 | 0 |
| 2L | 2421314 | 2 | 4 | 9 | 0 |
| 2L | 2422426 | 2 | 4 | 10 | 0 |
| 2L | 2537011 | 2 | 7 | 8 | 0 |
| 2L | 2539673 | 2 | 5 | 8 | 0 |
| 2L | 3113130 | 2 | 6 | 9 | 0 |
| 2L | 3121293 | 2 | 6 | 11 | 0 |
| 2L | 3125826 | 2 | 5 | 5 | 0 |
| 2L | 3125841 | 2 | 6 | 6 | 0 |
| 2L | 3126036 | 2 | 5 | 8 | 0 |
| 2L | 3141089 | 2 | 4 | 11 | 0 |
| 2L | 3145302 | 2 | 5 | 5 | 0 |
| 2L | 3184743 | 2 | 16 | 11 | 0 |
| 2L | 3240836 | 2 | 5 | 5 | 0 |
| 2L | 3276018 | 2 | 6 | 5 | 0 |
| 2L | 3407409 | 2 | 6 | 5 | 0 |
| 2L | 3472150 | 2 | 5 | 6 | 0 |
| 2L | 3585490 | 2 | 8 | 5 | 0 |
| 2L | 3611433 | 2 | 5 | 7 | 0 |
| 2L | 3724848 | 2 | 5 | 6 | 0 |
| 2L | 3729928 | 2 | 5 | 8 | 0 |
| 2L | 3777454 | 2 | 5 | 12 | 0 |
| 2L | 3820880 | 2 | 4 | 11 | 0 |
| 2L | 3873893 | 2 | 11 | 5 | 0 |
| 2L | 4209604 | 2 | 6 | 9 | 0 |
| 2L | 4231126 | 2 | 9 | 5 | 0 |
| 2L | 4292861 | 2 | 6 | 6 | 0 |
| 2L | 4301661 | 2 | 6 | 9 | 0 |
| 2L | 4320671 | 2 | 5 | 12 | 0 |
| 2L | 4339602 | 2 | 5 | 5 | 0 |
| 2L | 4403288 | 2 | 5 | 6 | 0 |
| 2L | 4417855 | 2 | 4 | 9 | 0 |
| 2L | 4486067 | 2 | 7 | 10 | 0 |
| 2L | 4706540 | 2 | 5 | 5 | 0 |
| 2L | 4730160 | 2 | 4 | 9 | 0 |
| 2L | 7194640 | 2 | 5 | 5 | 0 |
| 2L | 7288005 | 2 | 9 | 8 | 0 |
| 2L | 7443361 | 2 | 8 | 11 | 0 |
| 2L | 7922249 | 2 | 6 | 5 | 0 |
| 2L | 7974791 | 2 | 10 | 8 | 0 |
| 2L | 8467435 | 2 | 5 | 7 | 0 |
| 2L | 8508568 | 2 | 7 | 6 | 0 |
| 2L | 8580810 | 2 | 5 | 5 | 0 |
| 2L | 8587578 | 2 | 7 | 7 | 0 |
| 2L | 8614811 | 2 | 5 | 11 | 0 |
| 2L | 8624367 | 2 | 4 | 9 | 0 |
| 2L | 8634207 | 2 | 4 | 9 | 0 |
| 2L | 8762613 | 2 | 5 | 7 | 0 |
| 2L | 8765945 | 2 | 5 | 12 | 0 |
| 2L | 8779463 | 2 | 17 | 7 | 0 |
| 2L | 8780836 | 2 | 6 | 5 | 0 |
| 2L | 8810549 | 2 | 5 | 8 | 0 |
| 2L | 8810575 | 2 | 14 | 4 | 0 |
| 2L | 8824047 | 2 | 6 | 7 | 0 |
| 2L | 8871626 | 2 | 4 | 9 | 0 |
| 2L | 8927320 | 2 | 7 | 13 | 0 |
| 2L | 8967452 | 2 | 5 | 11 | 0 |
| 2L | 8981467 | 2 | 4 | 10 | 0 |
| 2L | 8989508 | 2 | 7 | 5 | 0 |
| 2L | 8994799 | 2 | 4 | 9 | 0 |
| 2L | 8999857 | 2 | 5 | 24 | 0 |
| 2L | 9048671 | 2 | 5 | 13 | 0 |
| 2L | 9064559 | 2 | 7 | 5 | 0 |
| 2L | 9098696 | 2 | 8 | 7 | 0 |
| 2L | 9101135 | 2 | 11 | 10 | 0 |
| 2L | 9114595 | 2 | 6 | 9 | 0 |
| 2L | 9150191 | 2 | 8 | 6 | 0 |
| 2L | 9209937 | 2 | 8 | 11 | 0 |
| 2L | 9214460 | 2 | 6 | 8 | 0 |
| 2L | 9235267 | 2 | 7 | 7 | 0 |
| 2L | 9261590 | 2 | 9 | 17 | 0 |
| 2L | 9302319 | 2 | 5 | 15 | 0 |
| 2L | 9321925 | 2 | 5 | 12 | 0 |
| 2L | 9329616 | 2 | 7 | 24 | 0 |
| 2L | 9333483 | 2 | 5 | 6 | 0 |
| 2L | 9344218 | 2 | 9 | 5 | 0 |
| 2L | 9364788 | 2 | 8 | 9 | 0 |
| 2L | 9387371 | 2 | 15 | 4 | 0 |
| 2L | 9457474 | 2 | 6 | 8 | 0 |
| 2L | 9504991 | 2 | 5 | 5 | 0 |
| 2L | 9527431 | 2 | 4 | 9 | 0 |
| 2L | 9600530 | 2 | 10 | 16 | 0 |
| 2L | 9608716 | 2 | 4 | 11 | 0 |
| 2L | 9611121 | 2 | 8 | 18 | 0 |
| 2L | 9633396 | 2 | 10 | 4 | 0 |
| 2L | 9779551 | 2 | 4 | 10 | 0 |
| 2L | 9784086 | 2 | 11 | 10 | 0 |
| 2L | 9882920 | 2 | 5 | 5 | 0 |
| 2L | 9911954 | 2 | 6 | 21 | 0 |
| 2L | 9957014 | 2 | 5 | 6 | 0 |
| 2L | 10011711 | 2 | 10 | 13 | 0 |
| 2L | 10015466 | 2 | 4 | 19 | 0 |
| 2L | 10017656 | 2 | 4 | 14 | 0 |
| 2L | 10020574 | 2 | 7 | 6 | 0 |
| 2L | 10020602 | 2 | 6 | 10 | 0 |
| 2L | 10049066 | 2 | 8 | 12 | 0 |
| 2L | 10136914 | 2 | 9 | 5 | 0 |
| 2L | 10151359 | 2 | 4 | 9 | 0 |
| 2L | 10152808 | 2 | 6 | 13 | 0 |
| 2L | 10165520 | 2 | 5 | 7 | 0 |
| 2L | 10178968 | 2 | 4 | 11 | 0 |
| 2L | 10199877 | 2 | 4 | 9 | 0 |
| 2L | 10212301 | 2 | 4 | 11 | 0 |
| 2L | 10212872 | 2 | 5 | 5 | 0 |
| 2L | 10250259 | 2 | 6 | 10 | 0 |
| 2L | 10252260 | 2 | 5 | 5 | 0 |
| 2L | 10307866 | 2 | 8 | 7 | 0 |
| 2L | 10838949 | 2 | 5 | 6 | 0 |
| 2L | 13695737 | 2 | 5 | 11 | 0 |
| 2L | 18748989 | 2 | 6 | 11 | 0 |
| 2R | 883056 | 2 | 6 | 5 | 0 |
| 2R | 3915796 | 2 | 8 | 8 | 0 |
| 2R | 3921584 | 2 | 7 | 5 | 0 |
| 2R | 3926376 | 2 | 5 | 28 | 0 |
| 2R | 3927715 | 2 | 4 | 13 | 0 |
| 2R | 3928319 | 2 | 32 | 13 | 0 |
| 2R | 3928457 | 2 | 5 | 7 | 0 |
| 2R | 3931643 | 2 | 5 | 18 | 0 |
| 2R | 3932881 | 2 | 23 | 19 | 0 |
| 2R | 3935024 | 2 | 8 | 32 | 0 |
| 2R | 3935141 | 2 | 4 | 22 | 0 |
| 2R | 3935573 | 2 | 21 | 15 | 0 |
| 2R | 3936788 | 2 | 9 | 5 | 0 |
| 2R | 3944833 | 2 | 13 | 14 | 0 |
| 2R | 3948365 | 2 | 8 | 8 | 0 |
| 2R | 3948894 | 2 | 9 | 8 | 0 |
| 2R | 3949215 | 2 | 17 | 5 | 0 |
| 2R | 3949671 | 2 | 11 | 11 | 0 |
| 2R | 3950717 | 2 | 6 | 8 | 0 |
| 2R | 3954596 | 2 | 10 | 5 | 0 |
| 2R | 3956824 | 2 | 12 | 5 | 0 |
| 2R | 3959542 | 2 | 9 | 9 | 0 |
| 2R | 3968849 | 2 | 18 | 6 | 0 |
| 2R | 3969167 | 2 | 11 | 4 | 0 |
| 2R | 3969297 | 2 | 9 | 4 | 0 |
| 2R | 3969573 | 2 | 6 | 10 | 0 |
| 2R | 3971783 | 2 | 7 | 6 | 0 |
| 2R | 3972864 | 2 | 11 | 6 | 0 |
| 2R | 3974155 | 2 | 20 | 8 | 0 |
| 2R | 3975602 | 2 | 16 | 11 | 0 |
| 2R | 4001944 | 2 | 7 | 5 | 0 |
| 2R | 4005880 | 2 | 10 | 12 | 0 |
| 2R | 4010646 | 2 | 29 | 22 | 0 |
| 2R | 4016702 | 2 | 15 | 4 | 0 |
| 2R | 4027965 | 2 | 19 | 18 | 0 |
| 2R | 4030731 | 2 | 27 | 9 | 0 |
| 2R | 4038931 | 2 | 8 | 5 | 0 |
| 2R | 4039111 | 2 | 9 | 8 | 0 |
| 2R | 4039991 | 2 | 22 | 15 | 0 |
| 2R | 4040628 | 2 | 17 | 14 | 0 |
| 2R | 4041476 | 2 | 17 | 4 | 0 |
| 2R | 4041948 | 2 | 17 | 11 | 0 |
| 2R | 4042093 | 2 | 14 | 4 | 0 |
| 2R | 4043058 | 2 | 26 | 16 | 0 |
| 2R | 4045370 | 2 | 16 | 31 | 0 |
| 2R | 4051169 | 2 | 9 | 4 | 0 |
| 2R | 4053115 | 2 | 16 | 4 | 0 |
| 2R | 4058718 | 2 | 26 | 9 | 0 |
| 2R | 4063232 | 2 | 4 | 15 | 0 |
| 2R | 4063447 | 2 | 17 | 25 | 0 |
| 2R | 4063546 | 2 | 4 | 12 | 0 |
| 2R | 4066186 | 2 | 9 | 14 | 0 |
| 2R | 4069126 | 2 | 12 | 5 | 0 |
| 2R | 4069389 | 2 | 19 | 12 | 0 |
| 2R | 4071886 | 2 | 5 | 6 | 0 |
| 2R | 4072483 | 2 | 9 | 16 | 0 |
| 2R | 4072681 | 2 | 16 | 9 | 0 |
| 2R | 4074894 | 2 | 6 | 6 | 0 |
| 2R | 4075154 | 2 | 5 | 16 | 0 |
| 2R | 4075479 | 2 | 21 | 26 | 0 |
| 2R | 4076676 | 2 | 12 | 5 | 0 |
| 2R | 4080366 | 2 | 14 | 6 | 0 |
| 2R | 4080711 | 2 | 10 | 10 | 0 |
| 2R | 4082191 | 2 | 8 | 6 | 0 |
| 2R | 4083436 | 2 | 14 | 6 | 0 |
| 2R | 4086408 | 2 | 4 | 23 | 0 |
| 2R | 4087268 | 2 | 9 | 4 | 0 |
| 2R | 4087390 | 2 | 8 | 12 | 0 |
| 2R | 4089530 | 2 | 11 | 7 | 0 |
| 2R | 4089915 | 2 | 15 | 5 | 0 |
| 2R | 4092182 | 2 | 11 | 17 | 0 |
| 2R | 4092387 | 2 | 20 | 7 | 0 |
| 2R | 4095130 | 2 | 5 | 8 | 0 |
| 2R | 4099635 | 2 | 26 | 9 | 0 |
| 2R | 4099688 | 2 | 31 | 8 | 0 |
| 2R | 4099819 | 2 | 11 | 8 | 0 |
| 2R | 4103337 | 2 | 24 | 10 | 0 |
| 2R | 4104635 | 2 | 29 | 25 | 0 |
| 2R | 4110358 | 2 | 7 | 13 | 0 |
| 2R | 4112484 | 2 | 10 | 24 | 0 |
| 2R | 4112833 | 2 | 10 | 6 | 0 |
| 2R | 4112996 | 2 | 37 | 24 | 0 |
| 2R | 4114645 | 2 | 9 | 34 | 0 |
| 2R | 4115014 | 2 | 9 | 17 | 0 |
| 2R | 4116407 | 2 | 25 | 16 | 0 |
| 2R | 4116851 | 2 | 9 | 10 | 0 |
| 2R | 4119375 | 2 | 13 | 6 | 0 |
| 2R | 4121667 | 2 | 24 | 4 | 0 |
| 2R | 4122424 | 2 | 13 | 5 | 0 |
| 2R | 4123001 | 2 | 10 | 4 | 0 |
| 2R | 4123485 | 2 | 15 | 7 | 0 |
| 2R | 4125976 | 2 | 19 | 9 | 0 |
| 2R | 4128568 | 2 | 16 | 13 | 0 |
| 2R | 4129353 | 2 | 22 | 13 | 0 |
| 2R | 4131123 | 2 | 4 | 16 | 0 |
| 2R | 4134149 | 2 | 27 | 8 | 0 |
| 2R | 4134924 | 2 | 17 | 8 | 0 |
| 2R | 4135038 | 2 | 20 | 5 | 0 |
| 2R | 4135687 | 2 | 26 | 8 | 0 |
| 2R | 4135787 | 2 | 6 | 19 | 0 |
| 2R | 4136152 | 2 | 23 | 4 | 0 |
| 2R | 4137681 | 2 | 11 | 5 | 0 |
| 2R | 4139607 | 2 | 9 | 4 | 0 |
| 2R | 4140177 | 2 | 14 | 9 | 0 |
| 2R | 4141187 | 2 | 14 | 7 | 0 |
| 2R | 4142597 | 2 | 19 | 4 | 0 |
| 2R | 4143233 | 2 | 26 | 6 | 0 |
| 2R | 4143388 | 2 | 11 | 12 | 0 |
| 2R | 4145741 | 2 | 4 | 21 | 0 |
| 2R | 4146304 | 2 | 21 | 9 | 0 |
| 2R | 4146345 | 2 | 16 | 8 | 0 |
| 2R | 4146450 | 2 | 10 | 7 | 0 |
| 2R | 4146906 | 2 | 14 | 5 | 0 |
| 2R | 4148155 | 2 | 14 | 7 | 0 |
| 2R | 4148268 | 2 | 9 | 18 | 0 |
| 2R | 4148737 | 2 | 5 | 31 | 0 |
| 2R | 4148763 | 2 | 9 | 28 | 0 |
| 2R | 4155357 | 2 | 6 | 7 | 0 |
| 2R | 4156587 | 2 | 8 | 35 | 0 |
| 2R | 4157198 | 2 | 15 | 5 | 0 |
| 2R | 4157261 | 2 | 7 | 6 | 0 |
| 2R | 4158233 | 2 | 13 | 5 | 0 |
| 2R | 4158282 | 2 | 14 | 8 | 0 |
| 2R | 4158298 | 2 | 13 | 6 | 0 |
| 2R | 4158391 | 2 | 5 | 7 | 0 |
| 2R | 4158540 | 2 | 9 | 4 | 0 |
| 2R | 4159196 | 2 | 20 | 7 | 0 |
| 2R | 4162197 | 2 | 16 | 9 | 0 |
| 2R | 4162785 | 2 | 8 | 20 | 0 |
| 2R | 4163113 | 2 | 7 | 12 | 0 |
| 2R | 4163585 | 2 | 9 | 20 | 0 |
| 2R | 4163760 | 2 | 5 | 15 | 0 |
| 2R | 4164645 | 2 | 4 | 13 | 0 |
| 2R | 4164790 | 2 | 5 | 19 | 0 |
| 2R | 4165617 | 2 | 15 | 21 | 0 |
| 2R | 4166846 | 2 | 5 | 12 | 0 |
| 2R | 4168365 | 2 | 4 | 15 | 0 |
| 2R | 4168631 | 2 | 18 | 4 | 0 |
| 2R | 4170063 | 2 | 26 | 14 | 0 |
| 2R | 4170400 | 2 | 7 | 6 | 0 |
| 2R | 4170571 | 2 | 6 | 7 | 0 |
| 2R | 4172682 | 2 | 5 | 17 | 0 |
| 2R | 4173193 | 2 | 7 | 17 | 0 |
| 2R | 4173770 | 2 | 17 | 15 | 0 |
| 2R | 4173842 | 2 | 4 | 11 | 0 |
| 2R | 4174889 | 2 | 23 | 16 | 0 |
| 2R | 4175715 | 2 | 4 | 20 | 0 |
| 2R | 4179139 | 2 | 13 | 4 | 0 |
| 2R | 4182470 | 2 | 10 | 13 | 0 |
| 2R | 4182667 | 2 | 6 | 7 | 0 |
| 2R | 4183215 | 2 | 15 | 13 | 0 |
| 2R | 4184462 | 2 | 14 | 7 | 0 |
| 2R | 4185763 | 2 | 10 | 4 | 0 |
| 2R | 4186222 | 2 | 18 | 14 | 0 |
| 2R | 4186688 | 2 | 11 | 5 | 0 |
| 2R | 4188324 | 2 | 17 | 12 | 0 |
| 2R | 4188735 | 2 | 15 | 5 | 0 |
| 2R | 4190686 | 2 | 10 | 9 | 0 |
| 2R | 4191653 | 2 | 5 | 12 | 0 |
| 2R | 4195164 | 2 | 9 | 8 | 0 |
| 2R | 4195690 | 2 | 20 | 16 | 0 |
| 2R | 4197605 | 2 | 23 | 8 | 0 |
| 2R | 4199104 | 2 | 26 | 20 | 0 |
| 2R | 4199743 | 2 | 20 | 8 | 0 |
| 2R | 4200562 | 2 | 15 | 6 | 0 |
| 2R | 4203868 | 2 | 11 | 7 | 0 |
| 2R | 4203938 | 2 | 12 | 6 | 0 |
| 2R | 4204995 | 2 | 9 | 17 | 0 |
| 2R | 4205387 | 2 | 21 | 11 | 0 |
| 2R | 4205442 | 2 | 28 | 10 | 0 |
| 2R | 4205470 | 2 | 21 | 4 | 0 |
| 2R | 4207953 | 2 | 16 | 7 | 0 |
| 2R | 4208733 | 2 | 13 | 9 | 0 |
| 2R | 4208830 | 2 | 20 | 10 | 0 |
| 2R | 4210279 | 2 | 17 | 8 | 0 |
| 2R | 4211023 | 2 | 6 | 10 | 0 |
| 2R | 4211314 | 2 | 10 | 7 | 0 |
| 2R | 4211620 | 2 | 18 | 4 | 0 |
| 2R | 4213615 | 2 | 17 | 10 | 0 |
| 2R | 4214322 | 2 | 17 | 4 | 0 |
| 2R | 4215077 | 2 | 13 | 17 | 0 |
| 2R | 4215389 | 2 | 38 | 7 | 0 |
| 2R | 4215504 | 2 | 17 | 16 | 0 |
| 2R | 4218387 | 2 | 5 | 8 | 0 |
| 2R | 4219099 | 2 | 10 | 7 | 0 |
| 2R | 4219788 | 2 | 8 | 6 | 0 |
| 2R | 4220930 | 2 | 17 | 17 | 0 |
| 2R | 4221453 | 2 | 5 | 13 | 0 |
| 2R | 4221753 | 2 | 4 | 13 | 0 |
| 2R | 4222886 | 2 | 16 | 14 | 0 |
| 2R | 4223461 | 2 | 8 | 10 | 0 |
| 2R | 4223523 | 2 | 14 | 5 | 0 |
| 2R | 4224092 | 2 | 28 | 8 | 0 |
| 2R | 4224767 | 2 | 27 | 12 | 0 |
| 2R | 4225282 | 2 | 22 | 14 | 0 |
| 2R | 4225544 | 2 | 13 | 5 | 0 |
| 2R | 4226815 | 2 | 24 | 9 | 0 |
| 2R | 4227321 | 2 | 20 | 14 | 0 |
| 2R | 4232365 | 2 | 15 | 9 | 0 |
| 2R | 4234161 | 2 | 18 | 13 | 0 |
| 2R | 4234588 | 2 | 12 | 11 | 0 |
| 2R | 4234703 | 2 | 21 | 11 | 0 |
| 2R | 4234736 | 2 | 15 | 7 | 0 |
| 2R | 4235896 | 2 | 11 | 5 | 0 |
| 2R | 4236027 | 2 | 8 | 12 | 0 |
| 2R | 4236095 | 2 | 26 | 14 | 0 |
| 2R | 4237670 | 2 | 8 | 5 | 0 |
| 2R | 4237726 | 2 | 7 | 25 | 0 |
| 2R | 4239446 | 2 | 16 | 4 | 0 |
| 2R | 4244308 | 2 | 6 | 8 | 0 |
| 2R | 4244350 | 2 | 8 | 9 | 0 |
| 2R | 4244633 | 2 | 18 | 12 | 0 |
| 2R | 4245566 | 2 | 7 | 16 | 0 |
| 2R | 4245598 | 2 | 20 | 10 | 0 |
| 2R | 4246799 | 2 | 22 | 4 | 0 |
| 2R | 4246951 | 2 | 7 | 7 | 0 |
| 2R | 4247063 | 2 | 27 | 17 | 0 |
| 2R | 4248042 | 2 | 22 | 8 | 0 |
| 2R | 4249339 | 2 | 18 | 23 | 0 |
| 2R | 4249551 | 2 | 22 | 9 | 0 |
| 2R | 4249951 | 2 | 6 | 5 | 0 |
| 2R | 4251702 | 2 | 17 | 5 | 0 |
| 2R | 4251918 | 2 | 11 | 19 | 0 |
| 2R | 4252458 | 2 | 20 | 12 | 0 |
| 2R | 4252645 | 2 | 16 | 12 | 0 |
| 2R | 4255996 | 2 | 26 | 12 | 0 |
| 2R | 4257639 | 2 | 9 | 7 | 0 |
| 2R | 4258194 | 2 | 18 | 6 | 0 |
| 2R | 4259712 | 2 | 13 | 4 | 0 |
| 2R | 4259964 | 2 | 11 | 28 | 0 |
| 2R | 4260531 | 2 | 13 | 4 | 0 |
| 2R | 4261270 | 2 | 7 | 7 | 0 |
| 2R | 4261322 | 2 | 14 | 5 | 0 |
| 2R | 4262914 | 2 | 26 | 15 | 0 |
| 2R | 4263967 | 2 | 7 | 19 | 0 |
| 2R | 4266435 | 2 | 11 | 12 | 0 |
| 2R | 4266933 | 2 | 16 | 9 | 0 |
| 2R | 4267339 | 2 | 6 | 19 | 0 |
| 2R | 4268342 | 2 | 15 | 9 | 0 |
| 2R | 4268661 | 2 | 16 | 9 | 0 |
| 2R | 4268903 | 2 | 14 | 13 | 0 |
| 2R | 4269378 | 2 | 19 | 12 | 0 |
| 2R | 4270499 | 2 | 13 | 13 | 0 |
| 2R | 4270664 | 2 | 8 | 21 | 0 |
| 2R | 4271121 | 2 | 10 | 4 | 0 |
| 2R | 4272000 | 2 | 15 | 16 | 0 |
| 2R | 4272895 | 2 | 7 | 6 | 0 |
| 2R | 4274188 | 2 | 6 | 9 | 0 |
| 2R | 4274586 | 2 | 17 | 6 | 0 |
| 2R | 4275982 | 2 | 10 | 9 | 0 |
| 2R | 4276047 | 2 | 10 | 6 | 0 |
| 2R | 4276249 | 2 | 25 | 15 | 0 |
| 2R | 4277752 | 2 | 15 | 7 | 0 |
| 2R | 4278507 | 2 | 10 | 5 | 0 |
| 2R | 4278916 | 2 | 13 | 9 | 0 |
| 2R | 4278994 | 2 | 30 | 12 | 0 |
| 2R | 4279470 | 2 | 15 | 7 | 0 |
| 2R | 4280298 | 2 | 5 | 13 | 0 |
| 2R | 4280555 | 2 | 13 | 4 | 0 |
| 2R | 4282905 | 2 | 15 | 10 | 0 |
| 2R | 4283460 | 2 | 5 | 11 | 0 |
| 2R | 4291213 | 2 | 6 | 35 | 0 |
| 2R | 4291249 | 2 | 20 | 18 | 0 |
| 2R | 4291588 | 2 | 8 | 7 | 0 |
| 2R | 4291670 | 2 | 7 | 23 | 0 |
| 2R | 4291904 | 2 | 16 | 8 | 0 |
| 2R | 4291983 | 2 | 23 | 18 | 0 |
| 2R | 4292157 | 2 | 6 | 17 | 0 |
| 2R | 4292859 | 2 | 19 | 5 | 0 |
| 2R | 4294068 | 2 | 11 | 6 | 0 |
| 2R | 4294782 | 2 | 5 | 7 | 0 |
| 2R | 4295031 | 2 | 13 | 8 | 0 |
| 2R | 4295083 | 2 | 5 | 6 | 0 |
| 2R | 4299950 | 2 | 12 | 22 | 0 |
| 2R | 4300847 | 2 | 25 | 22 | 0 |
| 2R | 4301052 | 2 | 15 | 4 | 0 |
| 2R | 4303514 | 2 | 25 | 6 | 0 |
| 2R | 4303672 | 2 | 12 | 7 | 0 |
| 2R | 4304901 | 2 | 14 | 12 | 0 |
| 2R | 4313631 | 2 | 14 | 7 | 0 |
| 2R | 4313669 | 2 | 9 | 6 | 0 |
| 2R | 4315669 | 2 | 19 | 19 | 0 |
| 2R | 4315698 | 2 | 24 | 15 | 0 |
| 2R | 4318646 | 2 | 14 | 8 | 0 |
| 2R | 4318938 | 2 | 10 | 5 | 0 |
| 2R | 4320603 | 2 | 11 | 4 | 0 |
| 2R | 4320861 | 2 | 24 | 21 | 0 |
| 2R | 4321121 | 2 | 14 | 8 | 0 |
| 2R | 4326644 | 2 | 15 | 16 | 0 |
| 2R | 4329032 | 2 | 9 | 8 | 0 |
| 2R | 4329217 | 2 | 8 | 12 | 0 |
| 2R | 4329647 | 2 | 6 | 14 | 0 |
| 2R | 4338772 | 2 | 18 | 17 | 0 |
| 2R | 4356187 | 2 | 4 | 26 | 0 |
| 2R | 4357856 | 2 | 20 | 20 | 0 |
| 2R | 4358018 | 2 | 21 | 4 | 0 |
| 2R | 4358664 | 2 | 20 | 15 | 0 |
| 2R | 4358729 | 2 | 24 | 4 | 0 |
| 2R | 4358816 | 2 | 21 | 14 | 0 |
| 2R | 4358944 | 2 | 15 | 16 | 0 |
| 2R | 4359074 | 2 | 18 | 27 | 0 |
| 2R | 4360693 | 2 | 10 | 25 | 0 |
| 2R | 4360804 | 2 | 6 | 15 | 0 |
| 2R | 4360921 | 2 | 13 | 24 | 0 |
| 2R | 4365120 | 2 | 4 | 16 | 0 |
| 2R | 4365241 | 2 | 7 | 8 | 0 |
| 2R | 4365765 | 2 | 5 | 10 | 0 |
| 2R | 4367758 | 2 | 17 | 5 | 0 |
| 2R | 4368084 | 2 | 28 | 16 | 0 |
| 2R | 4368756 | 2 | 17 | 7 | 0 |
| 2R | 4369748 | 2 | 17 | 9 | 0 |
| 2R | 4369865 | 2 | 30 | 13 | 0 |
| 2R | 4370507 | 2 | 27 | 13 | 0 |
| 2R | 4374658 | 2 | 6 | 7 | 0 |
| 2R | 4374847 | 2 | 22 | 9 | 0 |
| 2R | 4376121 | 2 | 19 | 6 | 0 |
| 2R | 4376417 | 2 | 10 | 26 | 0 |
| 2R | 4376604 | 2 | 5 | 5 | 0 |
| 2R | 4377077 | 2 | 8 | 6 | 0 |
| 2R | 4381024 | 2 | 13 | 4 | 0 |
| 2R | 4382219 | 2 | 12 | 5 | 0 |
| 2R | 4383623 | 2 | 10 | 15 | 0 |
| 2R | 4386050 | 2 | 18 | 5 | 0 |
| 2R | 4387334 | 2 | 8 | 10 | 0 |
| 2R | 4389781 | 2 | 20 | 8 | 0 |
| 2R | 4389884 | 2 | 14 | 10 | 0 |
| 2R | 4389948 | 2 | 28 | 15 | 0 |
| 2R | 4390166 | 2 | 10 | 4 | 0 |
| 2R | 4390239 | 2 | 27 | 7 | 0 |
| 2R | 4392165 | 2 | 16 | 4 | 0 |
| 2R | 4397170 | 2 | 9 | 10 | 0 |
| 2R | 4397362 | 2 | 4 | 9 | 0 |
| 2R | 4397978 | 2 | 24 | 13 | 0 |
| 2R | 4398644 | 2 | 7 | 6 | 0 |
| 2R | 4398703 | 2 | 6 | 19 | 0 |
| 2R | 4398850 | 2 | 6 | 11 | 0 |
| 2R | 4399028 | 2 | 7 | 25 | 0 |
| 2R | 4399970 | 2 | 12 | 15 | 0 |
| 2R | 4402029 | 2 | 11 | 5 | 0 |
| 2R | 4402272 | 2 | 7 | 8 | 0 |
| 2R | 4402631 | 2 | 14 | 6 | 0 |
| 2R | 4402761 | 2 | 28 | 5 | 0 |
| 2R | 4403000 | 2 | 17 | 12 | 0 |
| 2R | 4403863 | 2 | 5 | 5 | 0 |
| 2R | 4403963 | 2 | 16 | 12 | 0 |
| 2R | 4405604 | 2 | 5 | 14 | 0 |
| 2R | 4405668 | 2 | 15 | 12 | 0 |
| 2R | 4405904 | 2 | 10 | 17 | 0 |
| 2R | 4405979 | 2 | 9 | 4 | 0 |
| 2R | 4409390 | 2 | 30 | 13 | 0 |
| 2R | 4409473 | 2 | 28 | 14 | 0 |
| 2R | 4410332 | 2 | 11 | 6 | 0 |
| 2R | 4410877 | 2 | 17 | 4 | 0 |
| 2R | 4413067 | 2 | 16 | 13 | 0 |
| 2R | 4413562 | 2 | 10 | 5 | 0 |
| 2R | 4415201 | 2 | 25 | 12 | 0 |
| 2R | 4416203 | 2 | 20 | 13 | 0 |
| 2R | 4417899 | 2 | 14 | 8 | 0 |
| 2R | 4418458 | 2 | 16 | 7 | 0 |
| 2R | 4419448 | 2 | 4 | 16 | 0 |
| 2R | 4421098 | 2 | 10 | 7 | 0 |
| 2R | 4421475 | 2 | 20 | 11 | 0 |
| 2R | 4422261 | 2 | 7 | 5 | 0 |
| 2R | 4422560 | 2 | 9 | 4 | 0 |
| 2R | 4423288 | 2 | 5 | 7 | 0 |
| 2R | 4423402 | 2 | 12 | 11 | 0 |
| 2R | 4424124 | 2 | 22 | 8 | 0 |
| 2R | 4424188 | 2 | 14 | 5 | 0 |
| 2R | 4424300 | 2 | 15 | 9 | 0 |
| 2R | 4424402 | 2 | 5 | 5 | 0 |
| 2R | 4425054 | 2 | 9 | 12 | 0 |
| 2R | 4426412 | 2 | 5 | 6 | 0 |
| 2R | 4427009 | 2 | 4 | 12 | 0 |
| 2R | 4427867 | 2 | 17 | 13 | 0 |
| 2R | 4428794 | 2 | 5 | 5 | 0 |
| 2R | 4429171 | 2 | 6 | 7 | 0 |
| 2R | 4429346 | 2 | 11 | 4 | 0 |
| 2R | 4430567 | 2 | 7 | 17 | 0 |
| 2R | 4432041 | 2 | 13 | 4 | 0 |
| 2R | 4432149 | 2 | 7 | 8 | 0 |
| 2R | 4432245 | 2 | 4 | 16 | 0 |
| 2R | 4433400 | 2 | 28 | 17 | 0 |
| 2R | 4434415 | 2 | 7 | 5 | 0 |
| 2R | 4434552 | 2 | 8 | 26 | 0 |
| 2R | 4435101 | 2 | 13 | 11 | 0 |
| 2R | 4435231 | 2 | 14 | 39 | 0 |
| 2R | 4437053 | 2 | 18 | 16 | 0 |
| 2R | 4437978 | 2 | 11 | 7 | 0 |
| 2R | 4439624 | 2 | 24 | 6 | 0 |
| 2R | 4440610 | 2 | 19 | 7 | 0 |
| 2R | 4441184 | 2 | 4 | 9 | 0 |
| 2R | 4441246 | 2 | 14 | 6 | 0 |
| 2R | 4441649 | 2 | 5 | 6 | 0 |
| 2R | 4441762 | 2 | 14 | 11 | 0 |
| 2R | 4441816 | 2 | 15 | 9 | 0 |
| 2R | 4442979 | 2 | 10 | 7 | 0 |
| 2R | 4443156 | 2 | 19 | 7 | 0 |
| 2R | 4443420 | 2 | 7 | 5 | 0 |
| 2R | 4444608 | 2 | 19 | 10 | 0 |
| 2R | 4445736 | 2 | 16 | 4 | 0 |
| 2R | 4445942 | 2 | 7 | 9 | 0 |
| 2R | 4447049 | 2 | 15 | 9 | 0 |
| 2R | 4447366 | 2 | 11 | 8 | 0 |
| 2R | 4447463 | 2 | 34 | 21 | 0 |
| 2R | 4447545 | 2 | 5 | 5 | 0 |
| 2R | 4448556 | 2 | 37 | 10 | 0 |
| 2R | 4448624 | 2 | 18 | 5 | 0 |
| 2R | 4449350 | 2 | 5 | 13 | 0 |
| 2R | 4449601 | 2 | 15 | 22 | 0 |
| 2R | 4450353 | 2 | 12 | 14 | 0 |
| 2R | 4450588 | 2 | 22 | 12 | 0 |
| 2R | 4452794 | 2 | 24 | 4 | 0 |
| 2R | 4454918 | 2 | 5 | 21 | 0 |
| 2R | 4455744 | 2 | 16 | 6 | 0 |
| 2R | 4455772 | 2 | 12 | 9 | 0 |
| 2R | 4457383 | 2 | 23 | 7 | 0 |
| 2R | 4458819 | 2 | 20 | 12 | 0 |
| 2R | 4459026 | 2 | 23 | 10 | 0 |
| 2R | 4459463 | 2 | 22 | 16 | 0 |
| 2R | 4461012 | 2 | 14 | 9 | 0 |
| 2R | 4462673 | 2 | 7 | 25 | 0 |
| 2R | 4463292 | 2 | 12 | 13 | 0 |
| 2R | 4464789 | 2 | 25 | 15 | 0 |
| 2R | 4465936 | 2 | 15 | 7 | 0 |
| 2R | 4466099 | 2 | 9 | 6 | 0 |
| 2R | 4466406 | 2 | 9 | 4 | 0 |
| 2R | 4466790 | 2 | 4 | 19 | 0 |
| 2R | 4467087 | 2 | 13 | 22 | 0 |
| 2R | 4467244 | 2 | 19 | 20 | 0 |
| 2R | 4468329 | 2 | 24 | 4 | 0 |
| 2R | 4468681 | 2 | 14 | 10 | 0 |
| 2R | 4469108 | 2 | 15 | 9 | 0 |
| 2R | 4469156 | 2 | 11 | 5 | 0 |
| 2R | 4469848 | 2 | 9 | 9 | 0 |
| 2R | 4469878 | 2 | 10 | 12 | 0 |
| 2R | 4470182 | 2 | 19 | 5 | 0 |
| 2R | 4471074 | 2 | 11 | 8 | 0 |
| 2R | 4473215 | 2 | 13 | 4 | 0 |
| 2R | 4473388 | 2 | 16 | 5 | 0 |
| 2R | 4473478 | 2 | 11 | 4 | 0 |
| 2R | 4474082 | 2 | 10 | 9 | 0 |
| 2R | 4474367 | 2 | 12 | 7 | 0 |
| 2R | 4474715 | 2 | 13 | 9 | 0 |
| 2R | 4475140 | 2 | 17 | 8 | 0 |
| 2R | 4475411 | 2 | 8 | 9 | 0 |
| 2R | 4476017 | 2 | 29 | 14 | 0 |
| 2R | 4476822 | 2 | 10 | 4 | 0 |
| 2R | 4476988 | 2 | 15 | 5 | 0 |
| 2R | 4477423 | 2 | 24 | 16 | 0 |
| 2R | 4478034 | 2 | 17 | 11 | 0 |
| 2R | 4478961 | 2 | 7 | 8 | 0 |
| 2R | 4479939 | 2 | 12 | 4 | 0 |
| 2R | 4480327 | 2 | 15 | 11 | 0 |
| 2R | 4482057 | 2 | 9 | 6 | 0 |
| 2R | 4482332 | 2 | 26 | 8 | 0 |
| 2R | 4482835 | 2 | 21 | 11 | 0 |
| 2R | 4483796 | 2 | 15 | 6 | 0 |
| 2R | 4485560 | 2 | 9 | 23 | 0 |
| 2R | 4486918 | 2 | 7 | 13 | 0 |
| 2R | 4488793 | 2 | 9 | 18 | 0 |
| 2R | 4489118 | 2 | 12 | 5 | 0 |
| 2R | 4489949 | 2 | 16 | 28 | 0 |
| 2R | 4490392 | 2 | 5 | 20 | 0 |
| 2R | 4491725 | 2 | 24 | 20 | 0 |
| 2R | 4492048 | 2 | 18 | 11 | 0 |
| 2R | 4492536 | 2 | 18 | 13 | 0 |
| 2R | 4493444 | 2 | 8 | 5 | 0 |
| 2R | 4494466 | 2 | 21 | 10 | 0 |
| 2R | 4494761 | 2 | 11 | 6 | 0 |
| 2R | 4495689 | 2 | 23 | 7 | 0 |
| 2R | 4496560 | 2 | 10 | 4 | 0 |
| 2R | 4496707 | 2 | 4 | 15 | 0 |
| 2R | 4498120 | 2 | 9 | 4 | 0 |
| 2R | 4498800 | 2 | 15 | 15 | 0 |
| 2R | 4498871 | 2 | 20 | 9 | 0 |
| 2R | 4499274 | 2 | 4 | 15 | 0 |
| 2R | 4499456 | 2 | 11 | 6 | 0 |
| 2R | 4500032 | 2 | 11 | 5 | 0 |
| 2R | 4500268 | 2 | 4 | 17 | 0 |
| 2R | 4500424 | 2 | 8 | 13 | 0 |
| 2R | 4501379 | 2 | 22 | 9 | 0 |
| 2R | 4501840 | 2 | 13 | 7 | 0 |
| 2R | 4502911 | 2 | 10 | 21 | 0 |
| 2R | 4504543 | 2 | 5 | 6 | 0 |
| 2R | 4505736 | 2 | 5 | 18 | 0 |
| 2R | 4506260 | 2 | 9 | 11 | 0 |
| 2R | 4506551 | 2 | 31 | 14 | 0 |
| 2R | 4507772 | 2 | 13 | 7 | 0 |
| 2R | 4508682 | 2 | 29 | 21 | 0 |
| 2R | 4509076 | 2 | 18 | 22 | 0 |
| 2R | 4511754 | 2 | 18 | 4 | 0 |
| 2R | 4512580 | 2 | 28 | 16 | 0 |
| 2R | 4513500 | 2 | 8 | 8 | 0 |
| 2R | 4513616 | 2 | 18 | 9 | 0 |
| 2R | 4513814 | 2 | 12 | 13 | 0 |
| 2R | 4513915 | 2 | 19 | 8 | 0 |
| 2R | 4514328 | 2 | 20 | 10 | 0 |
| 2R | 4514451 | 2 | 21 | 15 | 0 |
| 2R | 4514544 | 2 | 23 | 5 | 0 |
| 2R | 4514607 | 2 | 32 | 7 | 0 |
| 2R | 4514753 | 2 | 8 | 8 | 0 |
| 2R | 4514904 | 2 | 12 | 4 | 0 |
| 2R | 4514974 | 2 | 15 | 17 | 0 |
| 2R | 4515517 | 2 | 12 | 5 | 0 |
| 2R | 4516141 | 2 | 15 | 5 | 0 |
| 2R | 4516450 | 2 | 15 | 10 | 0 |
| 2R | 4516643 | 2 | 18 | 4 | 0 |
| 2R | 4517151 | 2 | 23 | 14 | 0 |
| 2R | 4517636 | 2 | 24 | 9 | 0 |
| 2R | 4518784 | 2 | 5 | 8 | 0 |
| 2R | 4518916 | 2 | 10 | 9 | 0 |
| 2R | 4519324 | 2 | 30 | 21 | 0 |
| 2R | 4519617 | 2 | 12 | 5 | 0 |
| 2R | 4519663 | 2 | 20 | 8 | 0 |
| 2R | 4519766 | 2 | 13 | 4 | 0 |
| 2R | 4519805 | 2 | 21 | 13 | 0 |
| 2R | 4519870 | 2 | 19 | 10 | 0 |
| 2R | 4520489 | 2 | 22 | 6 | 0 |
| 2R | 4521369 | 2 | 14 | 6 | 0 |
| 2R | 4521972 | 2 | 19 | 7 | 0 |
| 2R | 4522690 | 2 | 21 | 6 | 0 |
| 2R | 4524375 | 2 | 14 | 7 | 0 |
| 2R | 4525356 | 2 | 11 | 7 | 0 |
| 2R | 4525987 | 2 | 14 | 5 | 0 |
| 2R | 4526229 | 2 | 27 | 7 | 0 |
| 2R | 4526400 | 2 | 16 | 4 | 0 |
| 2R | 4526508 | 2 | 7 | 24 | 0 |
| 2R | 4526554 | 2 | 10 | 22 | 0 |
| 2R | 4526611 | 2 | 13 | 4 | 0 |
| 2R | 4527180 | 2 | 4 | 9 | 0 |
| 2R | 4527836 | 2 | 12 | 4 | 0 |
| 2R | 4528077 | 2 | 34 | 16 | 0 |
| 2R | 4530115 | 2 | 7 | 8 | 0 |
| 2R | 4531617 | 2 | 6 | 5 | 0 |
| 2R | 4532231 | 2 | 5 | 15 | 0 |
| 2R | 4534783 | 2 | 10 | 5 | 0 |
| 2R | 4535391 | 2 | 7 | 23 | 0 |
| 2R | 4535845 | 2 | 7 | 10 | 0 |
| 2R | 4537537 | 2 | 15 | 8 | 0 |
| 2R | 4537605 | 2 | 21 | 23 | 0 |
| 2R | 4538980 | 2 | 9 | 9 | 0 |
| 2R | 4539075 | 2 | 6 | 6 | 0 |
| 2R | 4540734 | 2 | 19 | 7 | 0 |
| 2R | 4541136 | 2 | 12 | 5 | 0 |
| 2R | 4541449 | 2 | 9 | 9 | 0 |
| 2R | 4541642 | 2 | 23 | 10 | 0 |
| 2R | 4541689 | 2 | 19 | 11 | 0 |
| 2R | 4547173 | 2 | 23 | 9 | 0 |
| 2R | 4548040 | 2 | 9 | 17 | 0 |
| 2R | 4548813 | 2 | 4 | 9 | 0 |
| 2R | 4549695 | 2 | 5 | 21 | 0 |
| 2R | 4549769 | 2 | 11 | 25 | 0 |
| 2R | 4551268 | 2 | 19 | 7 | 0 |
| 2R | 4552498 | 2 | 15 | 5 | 0 |
| 2R | 4553865 | 2 | 9 | 7 | 0 |
| 2R | 4554060 | 2 | 13 | 12 | 0 |
| 2R | 4554716 | 2 | 18 | 22 | 0 |
| 2R | 4554948 | 2 | 11 | 8 | 0 |
| 2R | 4555046 | 2 | 20 | 4 | 0 |
| 2R | 4555392 | 2 | 21 | 9 | 0 |
| 2R | 4555931 | 2 | 12 | 4 | 0 |
| 2R | 4556063 | 2 | 19 | 13 | 0 |
| 2R | 4556284 | 2 | 9 | 9 | 0 |
| 2R | 4556399 | 2 | 14 | 10 | 0 |
| 2R | 4556477 | 2 | 9 | 13 | 0 |
| 2R | 4556679 | 2 | 12 | 8 | 0 |
| 2R | 4561209 | 2 | 4 | 16 | 0 |
| 2R | 4561223 | 2 | 4 | 12 | 0 |
| 2R | 4561244 | 2 | 11 | 10 | 0 |
| 2R | 4562020 | 2 | 26 | 13 | 0 |
| 2R | 4563016 | 2 | 14 | 9 | 0 |
| 2R | 4563903 | 2 | 10 | 8 | 0 |
| 2R | 4564402 | 2 | 23 | 16 | 0 |
| 2R | 4564849 | 2 | 11 | 13 | 0 |
| 2R | 4564857 | 2 | 9 | 13 | 0 |
| 2R | 4565364 | 2 | 6 | 6 | 0 |
| 2R | 4565482 | 2 | 19 | 17 | 0 |
| 2R | 4565547 | 2 | 6 | 10 | 0 |
| 2R | 4565952 | 2 | 17 | 8 | 0 |
| 2R | 4566336 | 2 | 17 | 6 | 0 |
| 2R | 4566403 | 2 | 24 | 15 | 0 |
| 2R | 4566458 | 2 | 23 | 21 | 0 |
| 2R | 4566836 | 2 | 12 | 5 | 0 |
| 2R | 4567125 | 2 | 18 | 12 | 0 |
| 2R | 4567989 | 2 | 7 | 14 | 0 |
| 2R | 4568694 | 2 | 26 | 23 | 0 |
| 2R | 4568757 | 2 | 19 | 9 | 0 |
| 2R | 4571260 | 2 | 6 | 13 | 0 |
| 2R | 4571770 | 2 | 19 | 5 | 0 |
| 2R | 4571912 | 2 | 16 | 13 | 0 |
| 2R | 4572342 | 2 | 5 | 6 | 0 |
| 2R | 4572597 | 2 | 10 | 4 | 0 |
| 2R | 4572726 | 2 | 12 | 5 | 0 |
| 2R | 4572744 | 2 | 15 | 4 | 0 |
| 2R | 4573764 | 2 | 12 | 10 | 0 |
| 2R | 4573997 | 2 | 15 | 4 | 0 |
| 2R | 4574721 | 2 | 26 | 14 | 0 |
| 2R | 4575995 | 2 | 14 | 4 | 0 |
| 2R | 4576261 | 2 | 16 | 6 | 0 |
| 2R | 4579896 | 2 | 24 | 4 | 0 |
| 2R | 4579910 | 2 | 21 | 6 | 0 |
| 2R | 4580509 | 2 | 21 | 15 | 0 |
| 2R | 4582865 | 2 | 19 | 8 | 0 |
| 2R | 4583351 | 2 | 30 | 4 | 0 |
| 2R | 4583439 | 2 | 15 | 12 | 0 |
| 2R | 4584729 | 2 | 25 | 7 | 0 |
| 2R | 4585667 | 2 | 24 | 10 | 0 |
| 2R | 4585781 | 2 | 13 | 7 | 0 |
| 2R | 4586026 | 2 | 20 | 9 | 0 |
| 2R | 4586949 | 2 | 12 | 6 | 0 |
| 2R | 4588768 | 2 | 23 | 7 | 0 |
| 2R | 4588861 | 2 | 15 | 6 | 0 |
| 2R | 4589005 | 2 | 23 | 7 | 0 |
| 2R | 4589635 | 2 | 30 | 10 | 0 |
| 2R | 4591107 | 2 | 16 | 7 | 0 |
| 2R | 4595176 | 2 | 24 | 6 | 0 |
| 2R | 4595697 | 2 | 24 | 4 | 0 |
| 2R | 4596778 | 2 | 12 | 10 | 0 |
| 2R | 4596972 | 2 | 20 | 4 | 0 |
| 2R | 4597969 | 2 | 15 | 10 | 0 |
| 2R | 4599586 | 2 | 23 | 12 | 0 |
| 2R | 4599762 | 2 | 14 | 17 | 0 |
| 2R | 4600037 | 2 | 26 | 16 | 0 |
| 2R | 4600348 | 2 | 15 | 10 | 0 |
| 2R | 4600368 | 2 | 9 | 13 | 0 |
| 2R | 4600547 | 2 | 6 | 10 | 0 |
| 2R | 4601637 | 2 | 11 | 10 | 0 |
| 2R | 4603063 | 2 | 14 | 5 | 0 |
| 2R | 4603537 | 2 | 21 | 6 | 0 |
| 2R | 4603878 | 2 | 10 | 7 | 0 |
| 2R | 4604173 | 2 | 14 | 8 | 0 |
| 2R | 4604184 | 2 | 16 | 10 | 0 |
| 2R | 4604541 | 2 | 11 | 16 | 0 |
| 2R | 4609188 | 2 | 17 | 15 | 0 |
| 2R | 4611189 | 2 | 12 | 5 | 0 |
| 2R | 4614055 | 2 | 9 | 4 | 0 |
| 2R | 4619937 | 2 | 17 | 5 | 0 |
| 2R | 4620784 | 2 | 14 | 5 | 0 |
| 2R | 4624018 | 2 | 10 | 13 | 0 |
| 2R | 4624246 | 2 | 18 | 5 | 0 |
| 2R | 4624435 | 2 | 5 | 6 | 0 |
| 2R | 4624689 | 2 | 16 | 7 | 0 |
| 2R | 4624810 | 2 | 24 | 16 | 0 |
| 2R | 4626449 | 2 | 5 | 16 | 0 |
| 2R | 4626598 | 2 | 5 | 9 | 0 |
| 2R | 4627460 | 2 | 5 | 27 | 0 |
| 2R | 4628923 | 2 | 21 | 15 | 0 |
| 2R | 4629251 | 2 | 28 | 16 | 0 |
| 2R | 4631880 | 2 | 12 | 9 | 0 |
| 2R | 4632562 | 2 | 4 | 11 | 0 |
| 2R | 4633675 | 2 | 5 | 15 | 0 |
| 2R | 4637476 | 2 | 9 | 5 | 0 |
| 2R | 4646288 | 2 | 10 | 6 | 0 |
| 2R | 4646330 | 2 | 25 | 23 | 0 |
| 2R | 4646430 | 2 | 6 | 15 | 0 |
| 2R | 4649172 | 2 | 12 | 11 | 0 |
| 2R | 4652266 | 2 | 7 | 5 | 0 |
| 2R | 4652371 | 2 | 10 | 6 | 0 |
| 2R | 4655402 | 2 | 16 | 4 | 0 |
| 2R | 4659690 | 2 | 13 | 14 | 0 |
| 2R | 4661109 | 2 | 12 | 21 | 0 |
| 2R | 4661363 | 2 | 17 | 33 | 0 |
| 2R | 4663676 | 2 | 30 | 7 | 0 |
| 2R | 4664609 | 2 | 10 | 6 | 0 |
| 2R | 4672005 | 2 | 7 | 7 | 0 |
| 2R | 4673634 | 2 | 9 | 24 | 0 |
| 2R | 4675760 | 2 | 11 | 15 | 0 |
| 2R | 4679536 | 2 | 30 | 10 | 0 |
| 2R | 4679720 | 2 | 9 | 7 | 0 |
| 2R | 4680279 | 2 | 19 | 17 | 0 |
| 2R | 4681554 | 2 | 13 | 8 | 0 |
| 2R | 4681981 | 2 | 15 | 10 | 0 |
| 2R | 4682067 | 2 | 8 | 19 | 0 |
| 2R | 4682158 | 2 | 19 | 16 | 0 |
| 2R | 4682773 | 2 | 22 | 8 | 0 |
| 2R | 4683142 | 2 | 6 | 16 | 0 |
| 2R | 4684134 | 2 | 4 | 20 | 0 |
| 2R | 4684993 | 2 | 24 | 6 | 0 |
| 2R | 4685885 | 2 | 10 | 22 | 0 |
| 2R | 4686547 | 2 | 27 | 8 | 0 |
| 2R | 4686760 | 2 | 18 | 8 | 0 |
| 2R | 4687065 | 2 | 7 | 29 | 0 |
| 2R | 4687891 | 2 | 28 | 12 | 0 |
| 2R | 4688825 | 2 | 19 | 33 | 0 |
| 2R | 4688910 | 2 | 14 | 16 | 0 |
| 2R | 4689320 | 2 | 18 | 5 | 0 |
| 2R | 4691041 | 2 | 7 | 5 | 0 |
| 2R | 4691257 | 2 | 6 | 14 | 0 |
| 2R | 4691285 | 2 | 7 | 6 | 0 |
| 2R | 4691733 | 2 | 18 | 7 | 0 |
| 2R | 4692383 | 2 | 11 | 23 | 0 |
| 2R | 4693513 | 2 | 16 | 10 | 0 |
| 2R | 4693622 | 2 | 5 | 7 | 0 |
| 2R | 4694343 | 2 | 4 | 19 | 0 |
| 2R | 4694498 | 2 | 13 | 4 | 0 |
| 2R | 4695076 | 2 | 5 | 5 | 0 |
| 2R | 4695137 | 2 | 12 | 6 | 0 |
| 2R | 4695783 | 2 | 14 | 5 | 0 |
| 2R | 4695853 | 2 | 13 | 5 | 0 |
| 2R | 4695970 | 2 | 18 | 12 | 0 |
| 2R | 4696570 | 2 | 21 | 11 | 0 |
| 2R | 4698344 | 2 | 16 | 6 | 0 |
| 2R | 4698706 | 2 | 22 | 20 | 0 |
| 2R | 4700759 | 2 | 8 | 8 | 0 |
| 2R | 4701420 | 2 | 15 | 8 | 0 |
| 2R | 4701501 | 2 | 11 | 4 | 0 |
| 2R | 4702167 | 2 | 16 | 7 | 0 |
| 2R | 4702185 | 2 | 11 | 13 | 0 |
| 2R | 4702264 | 2 | 14 | 7 | 0 |
| 2R | 4702931 | 2 | 17 | 16 | 0 |
| 2R | 4703523 | 2 | 4 | 17 | 0 |
| 2R | 4704635 | 2 | 15 | 4 | 0 |
| 2R | 4704709 | 2 | 21 | 5 | 0 |
| 2R | 4705135 | 2 | 16 | 6 | 0 |
| 2R | 4705185 | 2 | 23 | 13 | 0 |
| 2R | 4707598 | 2 | 4 | 11 | 0 |
| 2R | 4709566 | 2 | 19 | 8 | 0 |
| 2R | 4709760 | 2 | 18 | 6 | 0 |
| 2R | 4719279 | 2 | 20 | 9 | 0 |
| 2R | 4724439 | 2 | 5 | 23 | 0 |
| 2R | 4724574 | 2 | 18 | 19 | 0 |
| 2R | 4726147 | 2 | 14 | 18 | 0 |
| 2R | 4728597 | 2 | 7 | 15 | 0 |
| 2R | 4730573 | 2 | 9 | 4 | 0 |
| 2R | 4731077 | 2 | 4 | 11 | 0 |
| 2R | 4739280 | 2 | 7 | 5 | 0 |
| 2R | 4740262 | 2 | 26 | 16 | 0 |
| 2R | 4740683 | 2 | 12 | 4 | 0 |
| 2R | 4753256 | 2 | 9 | 4 | 0 |
| 2R | 4758668 | 2 | 28 | 9 | 0 |
| 2R | 4766145 | 2 | 6 | 17 | 0 |
| 2R | 4779025 | 2 | 14 | 11 | 0 |
| 2R | 4794423 | 2 | 22 | 15 | 0 |
| 2R | 4796295 | 2 | 15 | 5 | 0 |
| 2R | 4799432 | 2 | 20 | 10 | 0 |
| 2R | 4801550 | 2 | 14 | 6 | 0 |
| 2R | 4809725 | 2 | 6 | 7 | 0 |
| 2R | 4814797 | 2 | 6 | 24 | 0 |
| 2R | 4820309 | 2 | 7 | 9 | 0 |
| 2R | 4821067 | 2 | 28 | 12 | 0 |
| 2R | 4823787 | 2 | 12 | 9 | 0 |
| 2R | 4826061 | 2 | 10 | 6 | 0 |
| 2R | 4826290 | 2 | 16 | 26 | 0 |
| 2R | 4827093 | 2 | 10 | 26 | 0 |
| 2R | 4828580 | 2 | 6 | 6 | 0 |
| 2R | 4829149 | 2 | 18 | 16 | 0 |
| 2R | 4831141 | 2 | 17 | 12 | 0 |
| 2R | 4831273 | 2 | 12 | 15 | 0 |
| 2R | 4832198 | 2 | 8 | 7 | 0 |
| 2R | 4835014 | 2 | 12 | 4 | 0 |
| 2R | 4837720 | 2 | 34 | 10 | 0 |
| 2R | 4837993 | 2 | 13 | 7 | 0 |
| 2R | 4839146 | 2 | 6 | 9 | 0 |
| 2R | 4839406 | 2 | 20 | 6 | 0 |
| 2R | 4840963 | 2 | 12 | 15 | 0 |
| 2R | 4851707 | 2 | 10 | 6 | 0 |
| 2R | 4852559 | 2 | 7 | 6 | 0 |
| 2R | 4855675 | 2 | 19 | 16 | 0 |
| 2R | 4856828 | 2 | 17 | 16 | 0 |
| 2R | 4864063 | 2 | 4 | 9 | 0 |
| 2R | 4868835 | 2 | 6 | 19 | 0 |
| 2R | 4872015 | 2 | 16 | 8 | 0 |
| 2R | 4874699 | 2 | 6 | 5 | 0 |
| 2R | 4876987 | 2 | 8 | 14 | 0 |
| 2R | 4877490 | 2 | 5 | 27 | 0 |
| 2R | 4878209 | 2 | 21 | 8 | 0 |
| 2R | 4881718 | 2 | 6 | 17 | 0 |
| 2R | 4883905 | 2 | 10 | 10 | 0 |
| 2R | 4884769 | 2 | 14 | 4 | 0 |
| 2R | 4886001 | 2 | 9 | 8 | 0 |
| 2R | 4886886 | 2 | 22 | 10 | 0 |
| 2R | 4887321 | 2 | 16 | 11 | 0 |
| 2R | 4891833 | 2 | 16 | 10 | 0 |
| 2R | 4893649 | 2 | 16 | 13 | 0 |
| 2R | 4893736 | 2 | 7 | 9 | 0 |
| 2R | 4901452 | 2 | 17 | 9 | 0 |
| 2R | 4903685 | 2 | 17 | 5 | 0 |
| 2R | 4907246 | 2 | 15 | 8 | 0 |
| 2R | 4907567 | 2 | 19 | 9 | 0 |
| 2R | 4908337 | 2 | 20 | 6 | 0 |
| 2R | 4911303 | 2 | 4 | 16 | 0 |
| 2R | 4912301 | 2 | 13 | 6 | 0 |
| 2R | 4912531 | 2 | 8 | 16 | 0 |
| 2R | 4915257 | 2 | 9 | 14 | 0 |
| 2R | 4917380 | 2 | 8 | 8 | 0 |
| 2R | 4922139 | 2 | 11 | 19 | 0 |
| 2R | 4922988 | 2 | 21 | 16 | 0 |
| 2R | 4926591 | 2 | 14 | 17 | 0 |
| 2R | 4926653 | 2 | 25 | 12 | 0 |
| 2R | 4926750 | 2 | 8 | 5 | 0 |
| 2R | 4928195 | 2 | 16 | 9 | 0 |
| 2R | 4928336 | 2 | 10 | 5 | 0 |
| 2R | 4929760 | 2 | 15 | 4 | 0 |
| 2R | 4931400 | 2 | 12 | 11 | 0 |
| 2R | 4933910 | 2 | 11 | 60 | 0 |
| 2R | 4936572 | 2 | 5 | 7 | 0 |
| 2R | 4937612 | 2 | 21 | 17 | 0 |
| 2R | 4938888 | 2 | 10 | 6 | 0 |
| 2R | 4940486 | 2 | 19 | 7 | 0 |
| 2R | 4942391 | 2 | 21 | 15 | 0 |
| 2R | 4942420 | 2 | 20 | 10 | 0 |
| 2R | 4942733 | 2 | 5 | 11 | 0 |
| 2R | 4943713 | 2 | 16 | 5 | 0 |
| 2R | 4943844 | 2 | 14 | 10 | 0 |
| 2R | 4944686 | 2 | 24 | 22 | 0 |
| 2R | 4944816 | 2 | 16 | 10 | 0 |
| 2R | 4945452 | 2 | 16 | 4 | 0 |
| 2R | 4948040 | 2 | 6 | 7 | 0 |
| 2R | 4949324 | 2 | 32 | 9 | 0 |
| 2R | 4950597 | 2 | 4 | 22 | 0 |
| 2R | 4952590 | 2 | 7 | 7 | 0 |
| 2R | 4955426 | 2 | 5 | 10 | 0 |
| 2R | 4964342 | 2 | 15 | 7 | 0 |
| 2R | 4964909 | 2 | 5 | 29 | 0 |
| 2R | 4965351 | 2 | 7 | 19 | 0 |
| 2R | 4966352 | 2 | 13 | 8 | 0 |
| 2R | 4976191 | 2 | 4 | 21 | 0 |
| 2R | 4978623 | 2 | 21 | 6 | 0 |
| 2R | 4978640 | 2 | 16 | 7 | 0 |
| 2R | 4981322 | 2 | 16 | 23 | 0 |
| 2R | 4987386 | 2 | 4 | 23 | 0 |
| 2R | 4990312 | 2 | 28 | 18 | 0 |
| 2R | 4990425 | 2 | 17 | 10 | 0 |
| 2R | 4996709 | 2 | 20 | 4 | 0 |
| 2R | 4998116 | 2 | 5 | 20 | 0 |
| 2R | 5001176 | 2 | 5 | 5 | 0 |
| 2R | 5001406 | 2 | 5 | 6 | 0 |
| 2R | 5002838 | 2 | 17 | 40 | 0 |
| 2R | 5005210 | 2 | 13 | 4 | 0 |
| 2R | 5005386 | 2 | 11 | 18 | 0 |
| 2R | 5009262 | 2 | 14 | 12 | 0 |
| 2R | 5010734 | 2 | 17 | 10 | 0 |
| 2R | 5010868 | 2 | 6 | 27 | 0 |
| 2R | 5015154 | 2 | 7 | 24 | 0 |
| 2R | 5022116 | 2 | 9 | 5 | 0 |
| 2R | 5026158 | 2 | 19 | 12 | 0 |
| 2R | 5028193 | 2 | 12 | 25 | 0 |
| 2R | 5028929 | 2 | 17 | 12 | 0 |
| 2R | 5030388 | 2 | 11 | 7 | 0 |
| 2R | 5031437 | 2 | 10 | 5 | 0 |
| 2R | 5033518 | 2 | 8 | 6 | 0 |
| 2R | 5033590 | 2 | 10 | 5 | 0 |
| 2R | 5046236 | 2 | 14 | 9 | 0 |
| 2R | 5050119 | 2 | 18 | 11 | 0 |
| 2R | 5051742 | 2 | 17 | 9 | 0 |
| 2R | 5052534 | 2 | 9 | 20 | 0 |
| 2R | 5053536 | 2 | 5 | 21 | 0 |
| 2R | 5054362 | 2 | 6 | 9 | 0 |
| 2R | 5070728 | 2 | 22 | 8 | 0 |
| 2R | 5073857 | 2 | 10 | 4 | 0 |
| 2R | 5113193 | 2 | 9 | 35 | 0 |
| 2R | 5114267 | 2 | 11 | 4 | 0 |
| 2R | 5114714 | 2 | 7 | 12 | 0 |
| 2R | 5115020 | 2 | 14 | 4 | 0 |
| 2R | 5116246 | 2 | 4 | 14 | 0 |
| 2R | 5118618 | 2 | 26 | 13 | 0 |
| 2R | 5120475 | 2 | 14 | 11 | 0 |
| 2R | 5123641 | 2 | 8 | 13 | 0 |
| 2R | 5123739 | 2 | 5 | 9 | 0 |
| 2R | 5124002 | 2 | 16 | 5 | 0 |
| 2R | 5127108 | 2 | 13 | 4 | 0 |
| 2R | 5127659 | 2 | 11 | 8 | 0 |
| 2R | 5128268 | 2 | 7 | 14 | 0 |
| 2R | 5129245 | 2 | 19 | 9 | 0 |
| 2R | 5129382 | 2 | 20 | 7 | 0 |
| 2R | 5131086 | 2 | 11 | 5 | 0 |
| 2R | 5133306 | 2 | 14 | 12 | 0 |
| 2R | 5134686 | 2 | 9 | 7 | 0 |
| 2R | 5137019 | 2 | 5 | 12 | 0 |
| 2R | 5137395 | 2 | 7 | 13 | 0 |
| 2R | 5137592 | 2 | 5 | 30 | 0 |
| 2R | 5138887 | 2 | 5 | 14 | 0 |
| 2R | 5138937 | 2 | 7 | 16 | 0 |
| 2R | 5138967 | 2 | 6 | 8 | 0 |
| 2R | 5139013 | 2 | 19 | 10 | 0 |
| 2R | 5139145 | 2 | 10 | 19 | 0 |
| 2R | 5139179 | 2 | 10 | 27 | 0 |
| 2R | 5139221 | 2 | 6 | 15 | 0 |
| 2R | 5140577 | 2 | 21 | 14 | 0 |
| 2R | 5141234 | 2 | 26 | 31 | 0 |
| 2R | 5141338 | 2 | 17 | 7 | 0 |
| 2R | 5141614 | 2 | 7 | 15 | 0 |
| 2R | 5142918 | 2 | 10 | 10 | 0 |
| 2R | 5143146 | 2 | 13 | 14 | 0 |
| 2R | 5143402 | 2 | 4 | 18 | 0 |
| 2R | 5143472 | 2 | 4 | 14 | 0 |
| 2R | 5143796 | 2 | 9 | 4 | 0 |
| 2R | 5144773 | 2 | 11 | 7 | 0 |
| 2R | 5145506 | 2 | 6 | 5 | 0 |
| 2R | 5145553 | 2 | 15 | 14 | 0 |
| 2R | 5145595 | 2 | 37 | 17 | 0 |
| 2R | 5145734 | 2 | 19 | 9 | 0 |
| 2R | 5145775 | 2 | 17 | 13 | 0 |
| 2R | 5146768 | 2 | 32 | 9 | 0 |
| 2R | 5146978 | 2 | 12 | 8 | 0 |
| 2R | 5148101 | 2 | 16 | 8 | 0 |
| 2R | 5148182 | 2 | 16 | 7 | 0 |
| 2R | 5148425 | 2 | 21 | 14 | 0 |
| 2R | 5148541 | 2 | 24 | 5 | 0 |
| 2R | 5148728 | 2 | 14 | 12 | 0 |
| 2R | 5149987 | 2 | 31 | 19 | 0 |
| 2R | 5150228 | 2 | 12 | 15 | 0 |
| 2R | 5151432 | 2 | 10 | 6 | 0 |
| 2R | 5151438 | 2 | 11 | 9 | 0 |
| 2R | 5152092 | 2 | 4 | 20 | 0 |
| 2R | 5153685 | 2 | 4 | 20 | 0 |
| 2R | 5156807 | 2 | 5 | 6 | 0 |
| 2R | 5160577 | 2 | 7 | 25 | 0 |
| 2R | 5162573 | 2 | 9 | 21 | 0 |
| 2R | 5162656 | 2 | 5 | 19 | 0 |
| 2R | 5162728 | 2 | 7 | 9 | 0 |
| 2R | 5163361 | 2 | 8 | 18 | 0 |
| 2R | 5163621 | 2 | 7 | 14 | 0 |
| 2R | 5163777 | 2 | 7 | 15 | 0 |
| 2R | 5164522 | 2 | 6 | 8 | 0 |
| 2R | 5164839 | 2 | 20 | 5 | 0 |
| 2R | 5164919 | 2 | 17 | 18 | 0 |
| 2R | 5165020 | 2 | 19 | 20 | 0 |
| 2R | 5165093 | 2 | 20 | 17 | 0 |
| 2R | 5165208 | 2 | 33 | 12 | 0 |
| 2R | 5165634 | 2 | 6 | 24 | 0 |
| 2R | 5168349 | 2 | 14 | 9 | 0 |
| 2R | 5180257 | 2 | 4 | 9 | 0 |
| 2R | 5181681 | 2 | 15 | 6 | 0 |
| 2R | 5181730 | 2 | 4 | 18 | 0 |
| 2R | 5184428 | 2 | 18 | 14 | 0 |
| 2R | 5185649 | 2 | 8 | 11 | 0 |
| 2R | 5186712 | 2 | 26 | 5 | 0 |
| 2R | 5188816 | 2 | 15 | 4 | 0 |
| 2R | 5188909 | 2 | 9 | 4 | 0 |
| 2R | 5189210 | 2 | 15 | 9 | 0 |
| 2R | 5190616 | 2 | 12 | 20 | 0 |
| 2R | 5191382 | 2 | 23 | 9 | 0 |
| 2R | 5191429 | 2 | 12 | 5 | 0 |
| 2R | 5192535 | 2 | 8 | 30 | 0 |
| 2R | 5192953 | 2 | 24 | 18 | 0 |
| 2R | 5195197 | 2 | 7 | 10 | 0 |
| 2R | 5195251 | 2 | 14 | 10 | 0 |
| 2R | 5196359 | 2 | 4 | 19 | 0 |
| 2R | 5196501 | 2 | 7 | 8 | 0 |
| 2R | 5197420 | 2 | 20 | 14 | 0 |
| 2R | 5199316 | 2 | 4 | 9 | 0 |
| 2R | 5201139 | 2 | 5 | 21 | 0 |
| 2R | 5202830 | 2 | 8 | 18 | 0 |
| 2R | 5204212 | 2 | 11 | 16 | 0 |
| 2R | 5205292 | 2 | 9 | 18 | 0 |
| 2R | 5205475 | 2 | 6 | 7 | 0 |
| 2R | 5206138 | 2 | 9 | 12 | 0 |
| 2R | 5207041 | 2 | 11 | 6 | 0 |
| 2R | 5207191 | 2 | 13 | 10 | 0 |
| 2R | 5208094 | 2 | 10 | 5 | 0 |
| 2R | 5208096 | 2 | 10 | 6 | 0 |
| 2R | 5208569 | 2 | 4 | 11 | 0 |
| 2R | 5210965 | 2 | 14 | 7 | 0 |
| 2R | 5210986 | 2 | 7 | 9 | 0 |
| 2R | 5211612 | 2 | 12 | 9 | 0 |
| 2R | 5214835 | 2 | 5 | 19 | 0 |
| 2R | 5216784 | 2 | 8 | 14 | 0 |
| 2R | 5217532 | 2 | 18 | 10 | 0 |
| 2R | 5217623 | 2 | 18 | 9 | 0 |
| 2R | 5220015 | 2 | 4 | 14 | 0 |
| 2R | 5220267 | 2 | 17 | 10 | 0 |
| 2R | 5220702 | 2 | 16 | 15 | 0 |
| 2R | 5221869 | 2 | 8 | 9 | 0 |
| 2R | 5226277 | 2 | 9 | 4 | 0 |
| 2R | 5226617 | 2 | 14 | 5 | 0 |
| 2R | 5226933 | 2 | 8 | 5 | 0 |
| 2R | 5227342 | 2 | 11 | 9 | 0 |
| 2R | 5228435 | 2 | 20 | 16 | 0 |
| 2R | 5228495 | 2 | 21 | 4 | 0 |
| 2R | 5228590 | 2 | 18 | 21 | 0 |
| 2R | 5231430 | 2 | 14 | 21 | 0 |
| 2R | 5232122 | 2 | 14 | 4 | 0 |
| 2R | 5232576 | 2 | 26 | 9 | 0 |
| 2R | 5233297 | 2 | 20 | 4 | 0 |
| 2R | 5233621 | 2 | 9 | 18 | 0 |
| 2R | 5234075 | 2 | 18 | 16 | 0 |
| 2R | 5234520 | 2 | 10 | 31 | 0 |
| 2R | 5234795 | 2 | 17 | 8 | 0 |
| 2R | 5235538 | 2 | 7 | 16 | 0 |
| 2R | 5238453 | 2 | 10 | 7 | 0 |
| 2R | 5238875 | 2 | 9 | 9 | 0 |
| 2R | 5249347 | 2 | 14 | 31 | 0 |
| 2R | 5250140 | 2 | 6 | 14 | 0 |
| 2R | 5250444 | 2 | 13 | 21 | 0 |
| 2R | 5250698 | 2 | 6 | 10 | 0 |
| 2R | 5250948 | 2 | 8 | 16 | 0 |
| 2R | 5251301 | 2 | 11 | 23 | 0 |
| 2R | 5251977 | 2 | 5 | 29 | 0 |
| 2R | 5255093 | 2 | 7 | 11 | 0 |
| 2R | 5255110 | 2 | 11 | 13 | 0 |
| 2R | 5256600 | 2 | 12 | 6 | 0 |
| 2R | 5261900 | 2 | 8 | 31 | 0 |
| 2R | 5263332 | 2 | 11 | 12 | 0 |
| 2R | 5263473 | 2 | 8 | 14 | 0 |
| 2R | 5269108 | 2 | 10 | 9 | 0 |
| 2R | 5269601 | 2 | 4 | 19 | 0 |
| 2R | 5269897 | 2 | 4 | 21 | 0 |
| 2R | 5279323 | 2 | 17 | 17 | 0 |
| 2R | 5279421 | 2 | 11 | 8 | 0 |
| 2R | 5281621 | 2 | 26 | 10 | 0 |
| 2R | 5283327 | 2 | 14 | 17 | 0 |
| 2R | 5283768 | 2 | 9 | 6 | 0 |
| 2R | 5283990 | 2 | 6 | 10 | 0 |
| 2R | 5284647 | 2 | 25 | 8 | 0 |
| 2R | 5284851 | 2 | 10 | 20 | 0 |
| 2R | 5285898 | 2 | 12 | 4 | 0 |
| 2R | 5286023 | 2 | 4 | 13 | 0 |
| 2R | 5289394 | 2 | 7 | 14 | 0 |
| 2R | 5289856 | 2 | 9 | 4 | 0 |
| 2R | 5290126 | 2 | 18 | 4 | 0 |
| 2R | 5293103 | 2 | 5 | 9 | 0 |
| 2R | 5293456 | 2 | 4 | 21 | 0 |
| 2R | 5293749 | 2 | 22 | 8 | 0 |
| 2R | 5295110 | 2 | 10 | 10 | 0 |
| 2R | 5296246 | 2 | 10 | 5 | 0 |
| 2R | 5296632 | 2 | 19 | 4 | 0 |
| 2R | 5297539 | 2 | 15 | 4 | 0 |
| 2R | 5300112 | 2 | 7 | 23 | 0 |
| 2R | 5300241 | 2 | 9 | 10 | 0 |
| 2R | 5301420 | 2 | 27 | 6 | 0 |
| 2R | 5302638 | 2 | 28 | 14 | 0 |
| 2R | 5302709 | 2 | 28 | 5 | 0 |
| 2R | 5303635 | 2 | 18 | 4 | 0 |
| 2R | 5304129 | 2 | 5 | 20 | 0 |
| 2R | 5304331 | 2 | 4 | 15 | 0 |
| 2R | 5304539 | 2 | 24 | 18 | 0 |
| 2R | 5304732 | 2 | 6 | 15 | 0 |
| 2R | 5305954 | 2 | 12 | 39 | 0 |
| 2R | 5306423 | 2 | 5 | 13 | 0 |
| 2R | 5307444 | 2 | 8 | 5 | 0 |
| 2R | 5310214 | 2 | 7 | 6 | 0 |
| 2R | 5310983 | 2 | 25 | 12 | 0 |
| 2R | 5313287 | 2 | 7 | 11 | 0 |
| 2R | 5313925 | 2 | 7 | 16 | 0 |
| 2R | 5314063 | 2 | 9 | 21 | 0 |
| 2R | 5321762 | 2 | 7 | 22 | 0 |
| 2R | 5326019 | 2 | 7 | 14 | 0 |
| 2R | 5327128 | 2 | 12 | 10 | 0 |
| 2R | 5332789 | 2 | 10 | 21 | 0 |
| 2R | 5332906 | 2 | 11 | 6 | 0 |
| 2R | 5334246 | 2 | 7 | 12 | 0 |
| 2R | 5334281 | 2 | 5 | 20 | 0 |
| 2R | 5336853 | 2 | 4 | 21 | 0 |
| 2R | 5338611 | 2 | 20 | 6 | 0 |
| 2R | 5338730 | 2 | 10 | 11 | 0 |
| 2R | 5340023 | 2 | 15 | 12 | 0 |
| 2R | 5341512 | 2 | 19 | 11 | 0 |
| 2R | 5342249 | 2 | 18 | 7 | 0 |
| 2R | 5342322 | 2 | 23 | 16 | 0 |
| 2R | 5343999 | 2 | 12 | 6 | 0 |
| 2R | 5344045 | 2 | 19 | 13 | 0 |
| 2R | 5344316 | 2 | 15 | 10 | 0 |
| 2R | 5344401 | 2 | 26 | 8 | 0 |
| 2R | 5345020 | 2 | 4 | 13 | 0 |
| 2R | 5346498 | 2 | 6 | 25 | 0 |
| 2R | 5347396 | 2 | 12 | 11 | 0 |
| 2R | 5349423 | 2 | 7 | 10 | 0 |
| 2R | 5351601 | 2 | 22 | 19 | 0 |
| 2R | 5354366 | 2 | 4 | 12 | 0 |
| 2R | 5354892 | 2 | 5 | 8 | 0 |
| 2R | 5355492 | 2 | 20 | 6 | 0 |
| 2R | 5361718 | 2 | 29 | 4 | 0 |
| 2R | 5364333 | 2 | 5 | 17 | 0 |
| 2R | 5364734 | 2 | 18 | 17 | 0 |
| 2R | 5365591 | 2 | 13 | 4 | 0 |
| 2R | 5382073 | 2 | 20 | 11 | 0 |
| 2R | 5382247 | 2 | 13 | 10 | 0 |
| 2R | 5383153 | 2 | 12 | 7 | 0 |
| 2R | 5384034 | 2 | 20 | 5 | 0 |
| 2R | 5394057 | 2 | 23 | 4 | 0 |
| 2R | 5395594 | 2 | 16 | 6 | 0 |
| 2R | 7059535 | 2 | 7 | 6 | 0 |
| 2R | 7060850 | 2 | 13 | 4 | 0 |
| 2R | 7087689 | 2 | 14 | 6 | 0 |
| 2R | 7104715 | 2 | 11 | 8 | 0 |
| 2R | 7107191 | 2 | 13 | 5 | 0 |
| 2R | 7114493 | 2 | 28 | 9 | 0 |
| 2R | 7115670 | 2 | 19 | 8 | 0 |
| 2R | 7117748 | 2 | 17 | 8 | 0 |
| 2R | 7120049 | 2 | 5 | 5 | 0 |
| 2R | 7125347 | 2 | 14 | 20 | 0 |
| 2R | 7126328 | 2 | 11 | 11 | 0 |
| 2R | 7126386 | 2 | 14 | 4 | 0 |
| 2R | 7126889 | 2 | 8 | 5 | 0 |
| 2R | 7126912 | 2 | 5 | 7 | 0 |
| 2R | 7127614 | 2 | 11 | 4 | 0 |
| 2R | 7130991 | 2 | 20 | 7 | 0 |
| 2R | 7131627 | 2 | 12 | 7 | 0 |
| 2R | 7133769 | 2 | 11 | 4 | 0 |
| 2R | 7135452 | 2 | 8 | 5 | 0 |
| 2R | 7136293 | 2 | 6 | 6 | 0 |
| 2R | 7140393 | 2 | 12 | 5 | 0 |
| 2R | 7140780 | 2 | 4 | 18 | 0 |
| 2R | 7140872 | 2 | 13 | 12 | 0 |
| 2R | 7164834 | 2 | 10 | 4 | 0 |
| 2R | 7180142 | 2 | 12 | 6 | 0 |
| 2R | 7217607 | 2 | 19 | 8 | 0 |
| 2R | 7337511 | 2 | 6 | 14 | 0 |
| 2R | 7386442 | 2 | 5 | 5 | 0 |
| 2R | 7470544 | 2 | 6 | 5 | 0 |
| 2R | 7482127 | 2 | 6 | 6 | 0 |
| 2R | 7514871 | 2 | 5 | 6 | 0 |
| 2R | 7692059 | 2 | 5 | 5 | 0 |
| 2R | 8797425 | 2 | 10 | 5 | 0 |
| 2R | 10050925 | 2 | 6 | 5 | 0 |
| 2R | 10819403 | 2 | 5 | 6 | 0 |
| 2R | 11564762 | 2 | 5 | 15 | 0 |
| 2R | 12404110 | 2 | 8 | 16 | 0 |
| 2R | 12538064 | 2 | 11 | 4 | 0 |
| 2R | 12547208 | 2 | 14 | 7 | 0 |
| 2R | 12557923 | 2 | 7 | 5 | 0 |
| 2R | 12605941 | 2 | 11 | 5 | 0 |
| 2R | 12645466 | 2 | 6 | 8 | 0 |
| 2R | 12700340 | 2 | 11 | 4 | 0 |
| 2R | 12785102 | 2 | 6 | 7 | 0 |
| 2R | 12835342 | 2 | 4 | 10 | 0 |
| 2R | 13052695 | 2 | 4 | 9 | 0 |
| 2R | 13147264 | 2 | 6 | 5 | 0 |
| 2R | 13148363 | 2 | 15 | 5 | 0 |
| 2R | 13304982 | 2 | 7 | 7 | 0 |
| 2R | 13357074 | 2 | 6 | 6 | 0 |
| 2R | 13532631 | 2 | 6 | 5 | 0 |
| 2R | 13648614 | 2 | 12 | 6 | 0 |
| 2R | 13741244 | 2 | 9 | 5 | 0 |
| 2R | 13741253 | 2 | 7 | 6 | 0 |
| 2R | 13811347 | 2 | 4 | 13 | 0 |
| 2R | 14223065 | 2 | 5 | 5 | 0 |
| 2R | 15530974 | 2 | 6 | 5 | 0 |
| 2R | 15546399 | 2 | 7 | 5 | 0 |
| 2R | 16380626 | 2 | 5 | 6 | 0 |
| 2R | 17162490 | 2 | 9 | 4 | 0 |
| 2R | 17616067 | 2 | 5 | 5 | 0 |
| 2R | 20712666 | 2 | 9 | 7 | 0 |
| 2R | 20733085 | 2 | 14 | 4 | 0 |
| 2R | 20874975 | 2 | 4 | 10 | 0 |
| 2R | 21007206 | 2 | 4 | 11 | 0 |
| 2R | 21105286 | 2 | 10 | 5 | 0 |
| 2R | 21108822 | 2 | 5 | 9 | 0 |
| 2R | 21122080 | 2 | 9 | 5 | 0 |
| 3L | 108478 | 2 | 5 | 5 | 0 |
| 3L | 131690 | 2 | 4 | 9 | 0 |
| 3L | 438017 | 2 | 6 | 5 | 0 |
| 3L | 448575 | 2 | 5 | 11 | 0 |
| 3L | 558331 | 2 | 7 | 6 | 0 |
| 3L | 558644 | 2 | 4 | 9 | 0 |
| 3L | 576704 | 2 | 11 | 5 | 0 |
| 3L | 657947 | 2 | 12 | 9 | 0 |
| 3L | 659520 | 2 | 7 | 12 | 0 |
| 3L | 712066 | 2 | 4 | 10 | 0 |
| 3L | 713342 | 2 | 19 | 4 | 0 |
| 3L | 715761 | 2 | 5 | 10 | 0 |
| 3L | 716881 | 2 | 7 | 6 | 0 |
| 3L | 726572 | 2 | 8 | 7 | 0 |
| 3L | 739431 | 2 | 5 | 6 | 0 |
| 3L | 747303 | 2 | 16 | 9 | 0 |
| 3L | 750449 | 2 | 5 | 6 | 0 |
| 3L | 769090 | 2 | 15 | 11 | 0 |
| 3L | 771543 | 2 | 12 | 11 | 0 |
| 3L | 772783 | 2 | 17 | 5 | 0 |
| 3L | 774684 | 2 | 7 | 6 | 0 |
| 3L | 776866 | 2 | 15 | 9 | 0 |
| 3L | 785681 | 2 | 5 | 19 | 0 |
| 3L | 824769 | 2 | 4 | 16 | 0 |
| 3L | 832489 | 2 | 3 | 36 | 0 |
| 3L | 832809 | 2 | 5 | 13 | 0 |
| 3L | 849718 | 2 | 8 | 15 | 0 |
| 3L | 876013 | 2 | 6 | 10 | 0 |
| 3L | 876300 | 2 | 5 | 9 | 0 |
| 3L | 877143 | 2 | 5 | 7 | 0 |
| 3L | 884197 | 2 | 6 | 14 | 0 |
| 3L | 901662 | 2 | 9 | 4 | 0 |
| 3L | 904355 | 2 | 7 | 7 | 0 |
| 3L | 912395 | 2 | 9 | 7 | 0 |
| 3L | 919237 | 2 | 5 | 14 | 0 |
| 3L | 919251 | 2 | 4 | 10 | 0 |
| 3L | 941253 | 2 | 9 | 20 | 0 |
| 3L | 951541 | 2 | 9 | 4 | 0 |
| 3L | 970880 | 2 | 6 | 6 | 0 |
| 3L | 979868 | 2 | 10 | 6 | 0 |
| 3L | 989965 | 2 | 8 | 16 | 0 |
| 3L | 1009880 | 2 | 5 | 7 | 0 |
| 3L | 1013462 | 2 | 4 | 12 | 0 |
| 3L | 1018697 | 2 | 5 | 8 | 0 |
| 3L | 1021528 | 2 | 5 | 11 | 0 |
| 3L | 1057464 | 2 | 4 | 11 | 0 |
| 3L | 1059626 | 2 | 5 | 7 | 0 |
| 3L | 1093446 | 2 | 5 | 8 | 0 |
| 3L | 1104315 | 2 | 4 | 13 | 0 |
| 3L | 1175733 | 2 | 12 | 11 | 0 |
| 3L | 1186435 | 2 | 7 | 14 | 0 |
| 3L | 1188020 | 2 | 12 | 5 | 0 |
| 3L | 1194193 | 2 | 7 | 14 | 0 |
| 3L | 1194275 | 2 | 5 | 13 | 0 |
| 3L | 1203538 | 2 | 7 | 9 | 0 |
| 3L | 1218696 | 2 | 7 | 8 | 0 |
| 3L | 1229832 | 2 | 19 | 12 | 0 |
| 3L | 1265616 | 2 | 5 | 10 | 0 |
| 3L | 1269689 | 2 | 18 | 10 | 0 |
| 3L | 1281477 | 2 | 5 | 13 | 0 |
| 3L | 1284321 | 2 | 11 | 4 | 0 |
| 3L | 1286937 | 2 | 8 | 6 | 0 |
| 3L | 1293506 | 2 | 8 | 12 | 0 |
| 3L | 1302519 | 2 | 6 | 5 | 0 |
| 3L | 1308320 | 2 | 5 | 6 | 0 |
| 3L | 1314196 | 2 | 6 | 5 | 0 |
| 3L | 1317899 | 2 | 11 | 8 | 0 |
| 3L | 1357349 | 2 | 5 | 9 | 0 |
| 3L | 1378354 | 2 | 7 | 13 | 0 |
| 3L | 1400319 | 2 | 4 | 16 | 0 |
| 3L | 1405798 | 2 | 5 | 8 | 0 |
| 3L | 1417124 | 2 | 14 | 4 | 0 |
| 3L | 1448691 | 2 | 4 | 17 | 0 |
| 3L | 1453215 | 2 | 13 | 5 | 0 |
| 3L | 1455030 | 2 | 15 | 10 | 0 |
| 3L | 1483038 | 2 | 5 | 17 | 0 |
| 3L | 1503317 | 2 | 5 | 15 | 0 |
| 3L | 1503677 | 2 | 8 | 16 | 0 |
| 3L | 1522125 | 2 | 16 | 4 | 0 |
| 3L | 1523973 | 2 | 4 | 10 | 0 |
| 3L | 1528870 | 2 | 11 | 10 | 0 |
| 3L | 1528874 | 2 | 12 | 10 | 0 |
| 3L | 1533743 | 2 | 6 | 5 | 0 |
| 3L | 1535806 | 2 | 20 | 14 | 0 |
| 3L | 1541761 | 2 | 5 | 11 | 0 |
| 3L | 1543351 | 2 | 9 | 15 | 0 |
| 3L | 1553466 | 2 | 14 | 6 | 0 |
| 3L | 1567305 | 2 | 9 | 9 | 0 |
| 3L | 1574178 | 2 | 5 | 6 | 0 |
| 3L | 1577602 | 2 | 13 | 5 | 0 |
| 3L | 1590235 | 2 | 5 | 5 | 0 |
| 3L | 1594812 | 2 | 7 | 7 | 0 |
| 3L | 1598874 | 2 | 7 | 14 | 0 |
| 3L | 1602557 | 2 | 7 | 11 | 0 |
| 3L | 1605791 | 2 | 12 | 23 | 0 |
| 3L | 1610427 | 2 | 12 | 22 | 0 |
| 3L | 1617049 | 2 | 9 | 6 | 0 |
| 3L | 1617064 | 2 | 14 | 13 | 0 |
| 3L | 1621133 | 2 | 11 | 5 | 0 |
| 3L | 1622890 | 2 | 10 | 6 | 0 |
| 3L | 1627079 | 2 | 4 | 9 | 0 |
| 3L | 1627965 | 2 | 5 | 5 | 0 |
| 3L | 1629240 | 2 | 19 | 8 | 0 |
| 3L | 1631045 | 2 | 5 | 22 | 0 |
| 3L | 1633923 | 2 | 12 | 22 | 0 |
| 3L | 1636207 | 2 | 4 | 26 | 0 |
| 3L | 1640672 | 2 | 5 | 6 | 0 |
| 3L | 1643994 | 2 | 14 | 12 | 0 |
| 3L | 1647005 | 2 | 6 | 19 | 0 |
| 3L | 1647061 | 2 | 8 | 7 | 0 |
| 3L | 1649526 | 2 | 5 | 7 | 0 |
| 3L | 1654507 | 2 | 6 | 14 | 0 |
| 3L | 1657154 | 2 | 4 | 23 | 0 |
| 3L | 1657960 | 2 | 5 | 7 | 0 |
| 3L | 1661508 | 2 | 16 | 8 | 0 |
| 3L | 1661646 | 2 | 5 | 10 | 0 |
| 3L | 1662211 | 2 | 5 | 6 | 0 |
| 3L | 1664682 | 2 | 5 | 10 | 0 |
| 3L | 1666246 | 2 | 6 | 25 | 0 |
| 3L | 1669040 | 2 | 4 | 13 | 0 |
| 3L | 1670987 | 2 | 4 | 18 | 0 |
| 3L | 1673015 | 2 | 4 | 14 | 0 |
| 3L | 1673511 | 2 | 9 | 11 | 0 |
| 3L | 1674558 | 2 | 17 | 12 | 0 |
| 3L | 1675841 | 2 | 4 | 13 | 0 |
| 3L | 1676043 | 2 | 5 | 20 | 0 |
| 3L | 1678760 | 2 | 5 | 7 | 0 |
| 3L | 1679000 | 2 | 4 | 16 | 0 |
| 3L | 1679634 | 2 | 4 | 24 | 0 |
| 3L | 1681778 | 2 | 11 | 4 | 0 |
| 3L | 1687254 | 2 | 11 | 6 | 0 |
| 3L | 1688736 | 2 | 9 | 6 | 0 |
| 3L | 1688967 | 2 | 6 | 29 | 0 |
| 3L | 1695202 | 2 | 5 | 7 | 0 |
| 3L | 1698403 | 2 | 6 | 7 | 0 |
| 3L | 1699798 | 2 | 4 | 10 | 0 |
| 3L | 1700652 | 2 | 6 | 6 | 0 |
| 3L | 1701455 | 2 | 21 | 5 | 0 |
| 3L | 1701602 | 2 | 14 | 7 | 0 |
| 3L | 1701847 | 2 | 5 | 7 | 0 |
| 3L | 1702445 | 2 | 7 | 6 | 0 |
| 3L | 1705996 | 2 | 11 | 6 | 0 |
| 3L | 1707038 | 2 | 12 | 4 | 0 |
| 3L | 1708188 | 2 | 14 | 20 | 0 |
| 3L | 1708740 | 2 | 5 | 21 | 0 |
| 3L | 1710160 | 2 | 5 | 16 | 0 |
| 3L | 1713151 | 2 | 4 | 10 | 0 |
| 3L | 1716216 | 2 | 13 | 7 | 0 |
| 3L | 1717029 | 2 | 5 | 25 | 0 |
| 3L | 1717142 | 2 | 16 | 19 | 0 |
| 3L | 1717786 | 2 | 13 | 18 | 0 |
| 3L | 1718037 | 2 | 12 | 23 | 0 |
| 3L | 1719256 | 2 | 12 | 7 | 0 |
| 3L | 1720193 | 2 | 22 | 17 | 0 |
| 3L | 1720315 | 2 | 9 | 4 | 0 |
| 3L | 1720491 | 2 | 4 | 10 | 0 |
| 3L | 1722021 | 2 | 5 | 16 | 0 |
| 3L | 1722141 | 2 | 5 | 5 | 0 |
| 3L | 1722510 | 2 | 14 | 13 | 0 |
| 3L | 1723030 | 2 | 4 | 18 | 0 |
| 3L | 1723568 | 2 | 7 | 24 | 0 |
| 3L | 1724921 | 2 | 8 | 25 | 0 |
| 3L | 1729491 | 2 | 5 | 13 | 0 |
| 3L | 1729572 | 2 | 4 | 19 | 0 |
| 3L | 1729661 | 2 | 4 | 14 | 0 |
| 3L | 1731079 | 2 | 10 | 8 | 0 |
| 3L | 1731084 | 2 | 10 | 9 | 0 |
| 3L | 1731975 | 2 | 4 | 17 | 0 |
| 3L | 1733028 | 2 | 7 | 6 | 0 |
| 3L | 1733100 | 2 | 6 | 24 | 0 |
| 3L | 1734626 | 2 | 18 | 12 | 0 |
| 3L | 1734895 | 2 | 5 | 13 | 0 |
| 3L | 1735476 | 2 | 5 | 29 | 0 |
| 3L | 1737029 | 2 | 8 | 22 | 0 |
| 3L | 1737275 | 2 | 7 | 21 | 0 |
| 3L | 1738587 | 2 | 5 | 6 | 0 |
| 3L | 1739142 | 2 | 10 | 21 | 0 |
| 3L | 1739366 | 2 | 14 | 18 | 0 |
| 3L | 1739396 | 2 | 6 | 5 | 0 |
| 3L | 1742646 | 2 | 6 | 24 | 0 |
| 3L | 1742755 | 2 | 4 | 12 | 0 |
| 3L | 1744103 | 2 | 10 | 12 | 0 |
| 3L | 1759152 | 2 | 14 | 10 | 0 |
| 3L | 1759415 | 2 | 23 | 7 | 0 |
| 3L | 1763390 | 2 | 7 | 10 | 0 |
| 3L | 1764092 | 2 | 20 | 5 | 0 |
| 3L | 1765762 | 2 | 10 | 9 | 0 |
| 3L | 1766688 | 2 | 9 | 10 | 0 |
| 3L | 1767450 | 2 | 9 | 19 | 0 |
| 3L | 1768704 | 2 | 12 | 5 | 0 |
| 3L | 1771930 | 2 | 8 | 11 | 0 |
| 3L | 1775666 | 2 | 7 | 20 | 0 |
| 3L | 1775879 | 2 | 17 | 10 | 0 |
| 3L | 1775896 | 2 | 17 | 11 | 0 |
| 3L | 1776343 | 2 | 5 | 6 | 0 |
| 3L | 1777100 | 2 | 13 | 11 | 0 |
| 3L | 1777833 | 2 | 9 | 4 | 0 |
| 3L | 1779556 | 2 | 6 | 8 | 0 |
| 3L | 1780038 | 2 | 6 | 14 | 0 |
| 3L | 1783959 | 2 | 12 | 11 | 0 |
| 3L | 1784475 | 2 | 17 | 40 | 0 |
| 3L | 1784552 | 2 | 6 | 6 | 0 |
| 3L | 1784816 | 2 | 5 | 13 | 0 |
| 3L | 1784856 | 2 | 11 | 21 | 0 |
| 3L | 1786410 | 2 | 7 | 21 | 0 |
| 3L | 1787052 | 2 | 17 | 32 | 0 |
| 3L | 1787595 | 2 | 4 | 19 | 0 |
| 3L | 1789360 | 2 | 12 | 6 | 0 |
| 3L | 1790077 | 2 | 12 | 12 | 0 |
| 3L | 1790210 | 2 | 10 | 7 | 0 |
| 3L | 1791412 | 2 | 15 | 9 | 0 |
| 3L | 1794848 | 2 | 21 | 10 | 0 |
| 3L | 1798384 | 2 | 4 | 26 | 0 |
| 3L | 1798511 | 2 | 4 | 15 | 0 |
| 3L | 1799271 | 2 | 15 | 8 | 0 |
| 3L | 1803869 | 2 | 14 | 18 | 0 |
| 3L | 1805422 | 2 | 10 | 10 | 0 |
| 3L | 1809333 | 2 | 6 | 5 | 0 |
| 3L | 1812100 | 2 | 7 | 7 | 0 |
| 3L | 1813377 | 2 | 9 | 18 | 0 |
| 3L | 1817848 | 2 | 12 | 10 | 0 |
| 3L | 1818098 | 2 | 11 | 17 | 0 |
| 3L | 1828632 | 2 | 10 | 10 | 0 |
| 3L | 1833752 | 2 | 4 | 13 | 0 |
| 3L | 1834323 | 2 | 6 | 8 | 0 |
| 3L | 1834690 | 2 | 12 | 21 | 0 |
| 3L | 1834851 | 2 | 6 | 23 | 0 |
| 3L | 1839604 | 2 | 11 | 6 | 0 |
| 3L | 1840428 | 2 | 7 | 15 | 0 |
| 3L | 1840439 | 2 | 10 | 16 | 0 |
| 3L | 1840816 | 2 | 14 | 4 | 0 |
| 3L | 1841461 | 2 | 11 | 8 | 0 |
| 3L | 1843625 | 2 | 5 | 7 | 0 |
| 3L | 1844038 | 2 | 18 | 8 | 0 |
| 3L | 1851797 | 2 | 5 | 6 | 0 |
| 3L | 1852753 | 2 | 4 | 14 | 0 |
| 3L | 1854038 | 2 | 9 | 15 | 0 |
| 3L | 1855185 | 2 | 18 | 18 | 0 |
| 3L | 1855470 | 2 | 10 | 15 | 0 |
| 3L | 1855548 | 2 | 4 | 17 | 0 |
| 3L | 1855874 | 2 | 5 | 6 | 0 |
| 3L | 1856810 | 2 | 9 | 28 | 0 |
| 3L | 1857016 | 2 | 6 | 27 | 0 |
| 3L | 1860894 | 2 | 9 | 15 | 0 |
| 3L | 1862064 | 2 | 19 | 5 | 0 |
| 3L | 1863498 | 2 | 4 | 22 | 0 |
| 3L | 1863777 | 2 | 6 | 11 | 0 |
| 3L | 1867074 | 2 | 6 | 19 | 0 |
| 3L | 1867601 | 2 | 6 | 5 | 0 |
| 3L | 1870616 | 2 | 16 | 4 | 0 |
| 3L | 1873782 | 2 | 15 | 24 | 0 |
| 3L | 1874310 | 2 | 6 | 23 | 0 |
| 3L | 1874328 | 2 | 4 | 21 | 0 |
| 3L | 1875230 | 2 | 4 | 15 | 0 |
| 3L | 1880384 | 2 | 13 | 10 | 0 |
| 3L | 1884972 | 2 | 20 | 15 | 0 |
| 3L | 1892105 | 2 | 5 | 6 | 0 |
| 3L | 1893597 | 2 | 12 | 6 | 0 |
| 3L | 1894998 | 2 | 13 | 7 | 0 |
| 3L | 1906516 | 2 | 7 | 7 | 0 |
| 3L | 1912194 | 2 | 8 | 17 | 0 |
| 3L | 1913582 | 2 | 5 | 12 | 0 |
| 3L | 1921175 | 2 | 4 | 9 | 0 |
| 3L | 1922176 | 2 | 7 | 7 | 0 |
| 3L | 1922334 | 2 | 5 | 5 | 0 |
| 3L | 1927379 | 2 | 12 | 4 | 0 |
| 3L | 1932199 | 2 | 8 | 7 | 0 |
| 3L | 1932212 | 2 | 7 | 12 | 0 |
| 3L | 1934429 | 2 | 8 | 10 | 0 |
| 3L | 1934562 | 2 | 43 | 13 | 0 |
| 3L | 1935039 | 2 | 5 | 24 | 0 |
| 3L | 1935903 | 2 | 6 | 9 | 0 |
| 3L | 1935909 | 2 | 6 | 8 | 0 |
| 3L | 1936174 | 2 | 17 | 5 | 0 |
| 3L | 1936376 | 2 | 12 | 15 | 0 |
| 3L | 1942844 | 2 | 10 | 8 | 0 |
| 3L | 1943883 | 2 | 5 | 16 | 0 |
| 3L | 1944019 | 2 | 19 | 15 | 0 |
| 3L | 1944635 | 2 | 21 | 9 | 0 |
| 3L | 1944925 | 2 | 23 | 17 | 0 |
| 3L | 1959012 | 2 | 9 | 19 | 0 |
| 3L | 1959096 | 2 | 11 | 30 | 0 |
| 3L | 1959317 | 2 | 10 | 6 | 0 |
| 3L | 1959379 | 2 | 4 | 16 | 0 |
| 3L | 1960649 | 2 | 7 | 13 | 0 |
| 3L | 1961754 | 2 | 6 | 9 | 0 |
| 3L | 1969287 | 2 | 20 | 5 | 0 |
| 3L | 1973191 | 2 | 12 | 10 | 0 |
| 3L | 1973811 | 2 | 18 | 4 | 0 |
| 3L | 1976088 | 2 | 11 | 17 | 0 |
| 3L | 1980375 | 2 | 8 | 14 | 0 |
| 3L | 1982308 | 2 | 11 | 11 | 0 |
| 3L | 1989211 | 2 | 6 | 14 | 0 |
| 3L | 1991394 | 2 | 11 | 4 | 0 |
| 3L | 2006300 | 2 | 5 | 17 | 0 |
| 3L | 2013370 | 2 | 6 | 5 | 0 |
| 3L | 2014683 | 2 | 9 | 5 | 0 |
| 3L | 2015888 | 2 | 7 | 6 | 0 |
| 3L | 2017309 | 2 | 10 | 8 | 0 |
| 3L | 2021527 | 2 | 8 | 13 | 0 |
| 3L | 2021543 | 2 | 11 | 9 | 0 |
| 3L | 2022424 | 2 | 21 | 12 | 0 |
| 3L | 2022519 | 2 | 26 | 7 | 0 |
| 3L | 2039711 | 2 | 5 | 12 | 0 |
| 3L | 2056108 | 2 | 7 | 5 | 0 |
| 3L | 2058503 | 2 | 12 | 8 | 0 |
| 3L | 2073490 | 2 | 10 | 22 | 0 |
| 3L | 2080795 | 2 | 11 | 4 | 0 |
| 3L | 2082431 | 2 | 13 | 9 | 0 |
| 3L | 2088408 | 2 | 19 | 7 | 0 |
| 3L | 2096181 | 2 | 5 | 8 | 0 |
| 3L | 2099493 | 2 | 6 | 12 | 0 |
| 3L | 2106318 | 2 | 7 | 8 | 0 |
| 3L | 2123859 | 2 | 14 | 8 | 0 |
| 3L | 2135028 | 2 | 6 | 10 | 0 |
| 3L | 2135037 | 2 | 6 | 16 | 0 |
| 3L | 2135154 | 2 | 10 | 26 | 0 |
| 3L | 2135529 | 2 | 7 | 7 | 0 |
| 3L | 2135532 | 2 | 5 | 7 | 0 |
| 3L | 2141551 | 2 | 5 | 14 | 0 |
| 3L | 2147142 | 2 | 5 | 15 | 0 |
| 3L | 2150102 | 2 | 6 | 6 | 0 |
| 3L | 2153903 | 2 | 4 | 18 | 0 |
| 3L | 2161344 | 2 | 8 | 5 | 0 |
| 3L | 2172619 | 2 | 33 | 9 | 0 |
| 3L | 2174292 | 2 | 4 | 10 | 0 |
| 3L | 2201825 | 2 | 5 | 7 | 0 |
| 3L | 2211459 | 2 | 14 | 6 | 0 |
| 3L | 2230053 | 2 | 6 | 6 | 0 |
| 3L | 2232586 | 2 | 6 | 7 | 0 |
| 3L | 2233165 | 2 | 16 | 7 | 0 |
| 3L | 2234266 | 2 | 9 | 8 | 0 |
| 3L | 2235774 | 2 | 12 | 7 | 0 |
| 3L | 2253640 | 2 | 5 | 22 | 0 |
| 3L | 2254539 | 2 | 4 | 10 | 0 |
| 3L | 2255756 | 2 | 15 | 14 | 0 |
| 3L | 2257694 | 2 | 13 | 6 | 0 |
| 3L | 2259387 | 2 | 10 | 4 | 0 |
| 3L | 2259626 | 2 | 26 | 7 | 0 |
| 3L | 2283378 | 2 | 4 | 17 | 0 |
| 3L | 2284289 | 2 | 8 | 10 | 0 |
| 3L | 2310079 | 2 | 8 | 19 | 0 |
| 3L | 2313912 | 2 | 14 | 7 | 0 |
| 3L | 2319492 | 2 | 20 | 9 | 0 |
| 3L | 2334949 | 2 | 4 | 14 | 0 |
| 3L | 2343170 | 2 | 6 | 6 | 0 |
| 3L | 2352243 | 2 | 11 | 4 | 0 |
| 3L | 2354455 | 2 | 13 | 15 | 0 |
| 3L | 2355156 | 2 | 9 | 10 | 0 |
| 3L | 2358537 | 2 | 16 | 6 | 0 |
| 3L | 2360223 | 2 | 17 | 11 | 0 |
| 3L | 2365889 | 2 | 4 | 9 | 0 |
| 3L | 2374849 | 2 | 8 | 12 | 0 |
| 3L | 2376745 | 2 | 4 | 13 | 0 |
| 3L | 2378253 | 2 | 10 | 5 | 0 |
| 3L | 2379733 | 2 | 17 | 5 | 0 |
| 3L | 2385287 | 2 | 16 | 5 | 0 |
| 3L | 2396898 | 2 | 12 | 7 | 0 |
| 3L | 2399553 | 2 | 8 | 8 | 0 |
| 3L | 2400873 | 2 | 7 | 5 | 0 |
| 3L | 2403097 | 2 | 14 | 4 | 0 |
| 3L | 2403232 | 2 | 15 | 13 | 0 |
| 3L | 2403272 | 2 | 11 | 7 | 0 |
| 3L | 2406484 | 2 | 17 | 10 | 0 |
| 3L | 2410408 | 2 | 7 | 10 | 0 |
| 3L | 2415699 | 2 | 4 | 14 | 0 |
| 3L | 2422018 | 2 | 7 | 7 | 0 |
| 3L | 2422142 | 2 | 12 | 9 | 0 |
| 3L | 2423534 | 2 | 6 | 11 | 0 |
| 3L | 2428793 | 2 | 6 | 5 | 0 |
| 3L | 2434114 | 2 | 8 | 10 | 0 |
| 3L | 2435884 | 2 | 7 | 8 | 0 |
| 3L | 2435893 | 2 | 6 | 10 | 0 |
| 3L | 2437498 | 2 | 5 | 8 | 0 |
| 3L | 2438358 | 2 | 13 | 4 | 0 |
| 3L | 2438407 | 2 | 5 | 6 | 0 |
| 3L | 2439782 | 2 | 16 | 18 | 0 |
| 3L | 2458180 | 2 | 6 | 9 | 0 |
| 3L | 2463342 | 2 | 6 | 27 | 0 |
| 3L | 2466023 | 2 | 14 | 4 | 0 |
| 3L | 2473416 | 2 | 10 | 6 | 0 |
| 3L | 2486476 | 2 | 4 | 11 | 0 |
| 3L | 2488995 | 2 | 8 | 8 | 0 |
| 3L | 2492218 | 2 | 5 | 8 | 0 |
| 3L | 2492924 | 2 | 17 | 4 | 0 |
| 3L | 2493772 | 2 | 9 | 10 | 0 |
| 3L | 2496877 | 2 | 12 | 24 | 0 |
| 3L | 2496892 | 2 | 6 | 29 | 0 |
| 3L | 2503079 | 2 | 8 | 6 | 0 |
| 3L | 2503397 | 2 | 11 | 4 | 0 |
| 3L | 2505582 | 2 | 9 | 8 | 0 |
| 3L | 2505794 | 2 | 13 | 8 | 0 |
| 3L | 2520937 | 2 | 9 | 4 | 0 |
| 3L | 2522103 | 2 | 11 | 7 | 0 |
| 3L | 2540154 | 2 | 16 | 9 | 0 |
| 3L | 2549085 | 2 | 6 | 6 | 0 |
| 3L | 2562064 | 2 | 10 | 5 | 0 |
| 3L | 2562631 | 2 | 8 | 5 | 0 |
| 3L | 2567680 | 2 | 4 | 13 | 0 |
| 3L | 2569315 | 2 | 15 | 9 | 0 |
| 3L | 2570856 | 2 | 18 | 9 | 0 |
| 3L | 2574896 | 2 | 11 | 6 | 0 |
| 3L | 2575862 | 2 | 12 | 7 | 0 |
| 3L | 2578961 | 2 | 5 | 7 | 0 |
| 3L | 2583132 | 2 | 7 | 7 | 0 |
| 3L | 2588383 | 2 | 9 | 13 | 0 |
| 3L | 2595609 | 2 | 10 | 5 | 0 |
| 3L | 2598056 | 2 | 16 | 12 | 0 |
| 3L | 2599510 | 2 | 5 | 5 | 0 |
| 3L | 2600314 | 2 | 6 | 6 | 0 |
| 3L | 2606197 | 2 | 10 | 10 | 0 |
| 3L | 2606217 | 2 | 9 | 9 | 0 |
| 3L | 2608648 | 2 | 7 | 5 | 0 |
| 3L | 2619165 | 2 | 11 | 5 | 0 |
| 3L | 2619193 | 2 | 11 | 19 | 0 |
| 3L | 2631871 | 2 | 13 | 9 | 0 |
| 3L | 2633467 | 2 | 13 | 4 | 0 |
| 3L | 2633534 | 2 | 20 | 7 | 0 |
| 3L | 2634614 | 2 | 13 | 8 | 0 |
| 3L | 2635715 | 2 | 6 | 8 | 0 |
| 3L | 2638446 | 2 | 19 | 10 | 0 |
| 3L | 2639019 | 2 | 8 | 21 | 0 |
| 3L | 2641656 | 2 | 18 | 9 | 0 |
| 3L | 2642560 | 2 | 9 | 11 | 0 |
| 3L | 2660018 | 2 | 9 | 12 | 0 |
| 3L | 2660024 | 2 | 7 | 14 | 0 |
| 3L | 2660374 | 2 | 51 | 4 | 0 |
| 3L | 2662128 | 2 | 21 | 12 | 0 |
| 3L | 2666509 | 2 | 5 | 11 | 0 |
| 3L | 2675028 | 2 | 15 | 7 | 0 |
| 3L | 2685570 | 2 | 11 | 7 | 0 |
| 3L | 2688900 | 2 | 7 | 5 | 0 |
| 3L | 2696887 | 2 | 9 | 25 | 0 |
| 3L | 2710368 | 2 | 7 | 8 | 0 |
| 3L | 2711256 | 2 | 12 | 11 | 0 |
| 3L | 2712852 | 2 | 12 | 11 | 0 |
| 3L | 2713728 | 2 | 16 | 4 | 0 |
| 3L | 2714413 | 2 | 5 | 18 | 0 |
| 3L | 2715678 | 2 | 8 | 10 | 0 |
| 3L | 2716888 | 2 | 17 | 8 | 0 |
| 3L | 2725909 | 2 | 19 | 15 | 0 |
| 3L | 2726125 | 2 | 12 | 8 | 0 |
| 3L | 2727363 | 2 | 15 | 5 | 0 |
| 3L | 2741659 | 2 | 10 | 12 | 0 |
| 3L | 2743112 | 2 | 4 | 21 | 0 |
| 3L | 2747059 | 2 | 16 | 4 | 0 |
| 3L | 2750474 | 2 | 8 | 8 | 0 |
| 3L | 2750949 | 2 | 6 | 10 | 0 |
| 3L | 2760010 | 2 | 11 | 7 | 0 |
| 3L | 2763798 | 2 | 5 | 6 | 0 |
| 3L | 2765982 | 2 | 14 | 4 | 0 |
| 3L | 2766702 | 2 | 8 | 5 | 0 |
| 3L | 2767561 | 2 | 10 | 5 | 0 |
| 3L | 2767579 | 2 | 14 | 8 | 0 |
| 3L | 2770538 | 2 | 15 | 6 | 0 |
| 3L | 2772806 | 2 | 7 | 13 | 0 |
| 3L | 2773225 | 2 | 21 | 6 | 0 |
| 3L | 2774230 | 2 | 9 | 6 | 0 |
| 3L | 2778349 | 2 | 5 | 14 | 0 |
| 3L | 2785665 | 2 | 14 | 12 | 0 |
| 3L | 2788366 | 2 | 19 | 6 | 0 |
| 3L | 2789467 | 2 | 10 | 17 | 0 |
| 3L | 2790396 | 2 | 7 | 15 | 0 |
| 3L | 2793598 | 2 | 4 | 10 | 0 |
| 3L | 2794571 | 2 | 13 | 4 | 0 |
| 3L | 2799950 | 2 | 6 | 7 | 0 |
| 3L | 2813691 | 2 | 6 | 8 | 0 |
| 3L | 2817856 | 2 | 9 | 4 | 0 |
| 3L | 2822188 | 2 | 8 | 12 | 0 |
| 3L | 2828870 | 2 | 12 | 5 | 0 |
| 3L | 2830463 | 2 | 18 | 4 | 0 |
| 3L | 2834048 | 2 | 7 | 11 | 0 |
| 3L | 2834990 | 2 | 7 | 18 | 0 |
| 3L | 2835400 | 2 | 11 | 7 | 0 |
| 3L | 2838309 | 2 | 8 | 6 | 0 |
| 3L | 2848828 | 2 | 25 | 7 | 0 |
| 3L | 2856098 | 2 | 31 | 13 | 0 |
| 3L | 2859558 | 2 | 6 | 18 | 0 |
| 3L | 2867275 | 2 | 5 | 7 | 0 |
| 3L | 2867646 | 2 | 9 | 15 | 0 |
| 3L | 2868878 | 2 | 18 | 6 | 0 |
| 3L | 2873963 | 2 | 14 | 6 | 0 |
| 3L | 2874039 | 2 | 14 | 6 | 0 |
| 3L | 2882277 | 2 | 8 | 13 | 0 |
| 3L | 2882766 | 2 | 12 | 14 | 0 |
| 3L | 2882777 | 2 | 12 | 11 | 0 |
| 3L | 2882865 | 2 | 14 | 7 | 0 |
| 3L | 2886847 | 2 | 10 | 11 | 0 |
| 3L | 2889892 | 2 | 10 | 7 | 0 |
| 3L | 2902251 | 2 | 10 | 12 | 0 |
| 3L | 2902455 | 2 | 4 | 21 | 0 |
| 3L | 2905075 | 2 | 11 | 5 | 0 |
| 3L | 2908433 | 2 | 10 | 4 | 0 |
| 3L | 2915999 | 2 | 11 | 5 | 0 |
| 3L | 2917109 | 2 | 6 | 17 | 0 |
| 3L | 2920621 | 2 | 7 | 16 | 0 |
| 3L | 2921482 | 2 | 5 | 14 | 0 |
| 3L | 2924011 | 2 | 9 | 7 | 0 |
| 3L | 2938034 | 2 | 18 | 21 | 0 |
| 3L | 2942869 | 2 | 14 | 11 | 0 |
| 3L | 2945474 | 2 | 6 | 21 | 0 |
| 3L | 2955764 | 2 | 8 | 18 | 0 |
| 3L | 2957766 | 2 | 7 | 9 | 0 |
| 3L | 2958437 | 2 | 7 | 6 | 0 |
| 3L | 2959712 | 2 | 8 | 10 | 0 |
| 3L | 2963891 | 2 | 17 | 31 | 0 |
| 3L | 2964169 | 2 | 8 | 8 | 0 |
| 3L | 2979085 | 2 | 13 | 19 | 0 |
| 3L | 2981656 | 2 | 10 | 10 | 0 |
| 3L | 2984445 | 2 | 10 | 5 | 0 |
| 3L | 2984458 | 2 | 5 | 8 | 0 |
| 3L | 2987109 | 2 | 9 | 4 | 0 |
| 3L | 2988412 | 2 | 7 | 16 | 0 |
| 3L | 2989277 | 2 | 9 | 4 | 0 |
| 3L | 3004782 | 2 | 9 | 5 | 0 |
| 3L | 3006863 | 2 | 9 | 5 | 0 |
| 3L | 3006865 | 2 | 5 | 5 | 0 |
| 3L | 3007190 | 2 | 7 | 26 | 0 |
| 3L | 3008718 | 2 | 8 | 25 | 0 |
| 3L | 3008755 | 2 | 10 | 17 | 0 |
| 3L | 3008931 | 2 | 5 | 12 | 0 |
| 3L | 3009288 | 2 | 4 | 18 | 0 |
| 3L | 3022314 | 2 | 16 | 10 | 0 |
| 3L | 3032551 | 2 | 10 | 14 | 0 |
| 3L | 3048476 | 2 | 11 | 13 | 0 |
| 3L | 3048727 | 2 | 18 | 10 | 0 |
| 3L | 3051466 | 2 | 10 | 6 | 0 |
| 3L | 3055224 | 2 | 14 | 5 | 0 |
| 3L | 3057405 | 2 | 14 | 13 | 0 |
| 3L | 3057978 | 2 | 14 | 6 | 0 |
| 3L | 3079550 | 2 | 9 | 9 | 0 |
| 3L | 3081310 | 2 | 9 | 4 | 0 |
| 3L | 3082631 | 2 | 12 | 4 | 0 |
| 3L | 3083118 | 2 | 27 | 5 | 0 |
| 3L | 3101133 | 2 | 11 | 5 | 0 |
| 3L | 3102510 | 2 | 28 | 8 | 0 |
| 3L | 3102741 | 2 | 7 | 8 | 0 |
| 3L | 3107349 | 2 | 10 | 9 | 0 |
| 3L | 3107581 | 2 | 16 | 13 | 0 |
| 3L | 3110278 | 2 | 13 | 15 | 0 |
| 3L | 3115472 | 2 | 5 | 10 | 0 |
| 3L | 3124463 | 2 | 14 | 14 | 0 |
| 3L | 3125351 | 2 | 4 | 10 | 0 |
| 3L | 3126967 | 2 | 11 | 4 | 0 |
| 3L | 3132714 | 2 | 13 | 5 | 0 |
| 3L | 3133477 | 2 | 5 | 7 | 0 |
| 3L | 3135468 | 2 | 12 | 6 | 0 |
| 3L | 3136286 | 2 | 4 | 16 | 0 |
| 3L | 3139007 | 2 | 12 | 10 | 0 |
| 3L | 3140125 | 2 | 6 | 6 | 0 |
| 3L | 3144093 | 2 | 17 | 8 | 0 |
| 3L | 3145251 | 2 | 15 | 7 | 0 |
| 3L | 3145362 | 2 | 18 | 4 | 0 |
| 3L | 3148261 | 2 | 13 | 5 | 0 |
| 3L | 3154693 | 2 | 8 | 16 | 0 |
| 3L | 3165345 | 2 | 12 | 8 | 0 |
| 3L | 3171040 | 2 | 12 | 5 | 0 |
| 3L | 3190117 | 2 | 13 | 7 | 0 |
| 3L | 3190168 | 2 | 11 | 5 | 0 |
| 3L | 3192952 | 2 | 17 | 11 | 0 |
| 3L | 3197437 | 2 | 8 | 16 | 0 |
| 3L | 3200485 | 2 | 4 | 18 | 0 |
| 3L | 3200811 | 2 | 6 | 26 | 0 |
| 3L | 3205560 | 2 | 4 | 19 | 0 |
| 3L | 3205692 | 2 | 4 | 13 | 0 |
| 3L | 3207707 | 2 | 7 | 14 | 0 |
| 3L | 3211852 | 2 | 6 | 27 | 0 |
| 3L | 3237294 | 2 | 8 | 18 | 0 |
| 3L | 3248706 | 2 | 23 | 9 | 0 |
| 3L | 3271200 | 2 | 15 | 4 | 0 |
| 3L | 3273610 | 2 | 8 | 14 | 0 |
| 3L | 3276156 | 2 | 15 | 5 | 0 |
| 3L | 3279526 | 2 | 11 | 6 | 0 |
| 3L | 3287660 | 2 | 8 | 5 | 0 |
| 3L | 3287947 | 2 | 14 | 21 | 0 |
| 3L | 3288239 | 2 | 11 | 26 | 0 |
| 3L | 3294635 | 2 | 9 | 16 | 0 |
| 3L | 3294914 | 2 | 8 | 5 | 0 |
| 3L | 3296715 | 2 | 17 | 12 | 0 |
| 3L | 3307754 | 2 | 22 | 10 | 0 |
| 3L | 3311366 | 2 | 4 | 9 | 0 |
| 3L | 3313160 | 2 | 6 | 13 | 0 |
| 3L | 3316302 | 2 | 12 | 17 | 0 |
| 3L | 3319622 | 2 | 14 | 6 | 0 |
| 3L | 3323547 | 2 | 6 | 23 | 0 |
| 3L | 3327895 | 2 | 10 | 15 | 0 |
| 3L | 3332774 | 2 | 9 | 14 | 0 |
| 3L | 3335646 | 2 | 9 | 5 | 0 |
| 3L | 3336324 | 2 | 15 | 7 | 0 |
| 3L | 3347724 | 2 | 11 | 4 | 0 |
| 3L | 3349642 | 2 | 8 | 19 | 0 |
| 3L | 3365544 | 2 | 7 | 22 | 0 |
| 3L | 3376716 | 2 | 5 | 13 | 0 |
| 3L | 3411188 | 2 | 8 | 11 | 0 |
| 3L | 3413382 | 2 | 12 | 4 | 0 |
| 3L | 3423097 | 2 | 4 | 21 | 0 |
| 3L | 3454673 | 2 | 8 | 6 | 0 |
| 3L | 3469931 | 2 | 20 | 9 | 0 |
| 3L | 3492919 | 2 | 13 | 4 | 0 |
| 3L | 3495067 | 2 | 10 | 10 | 0 |
| 3L | 3495087 | 2 | 7 | 7 | 0 |
| 3L | 3497447 | 2 | 5 | 6 | 0 |
| 3L | 3501628 | 2 | 5 | 16 | 0 |
| 3L | 3501642 | 2 | 5 | 12 | 0 |
| 3L | 3508869 | 2 | 6 | 9 | 0 |
| 3L | 3513608 | 2 | 10 | 11 | 0 |
| 3L | 3516843 | 2 | 13 | 4 | 0 |
| 3L | 3518764 | 2 | 6 | 5 | 0 |
| 3L | 3519439 | 2 | 7 | 22 | 0 |
| 3L | 3525734 | 2 | 8 | 12 | 0 |
| 3L | 3532034 | 2 | 4 | 12 | 0 |
| 3L | 3532216 | 2 | 10 | 7 | 0 |
| 3L | 3532322 | 2 | 7 | 20 | 0 |
| 3L | 3534541 | 2 | 12 | 10 | 0 |
| 3L | 3538035 | 2 | 18 | 12 | 0 |
| 3L | 3540116 | 2 | 4 | 14 | 0 |
| 3L | 3543718 | 2 | 7 | 6 | 0 |
| 3L | 3547111 | 2 | 4 | 10 | 0 |
| 3L | 3550580 | 2 | 13 | 8 | 0 |
| 3L | 3552360 | 2 | 4 | 11 | 0 |
| 3L | 3556945 | 2 | 9 | 14 | 0 |
| 3L | 3557132 | 2 | 6 | 9 | 0 |
| 3L | 3558860 | 2 | 6 | 7 | 0 |
| 3L | 3571257 | 2 | 9 | 5 | 0 |
| 3L | 3571411 | 2 | 4 | 15 | 0 |
| 3L | 3575995 | 2 | 8 | 5 | 0 |
| 3L | 3579191 | 2 | 7 | 7 | 0 |
| 3L | 3585061 | 2 | 4 | 11 | 0 |
| 3L | 3586159 | 2 | 5 | 5 | 0 |
| 3L | 3586930 | 2 | 15 | 6 | 0 |
| 3L | 3589435 | 2 | 10 | 4 | 0 |
| 3L | 3596412 | 2 | 12 | 12 | 0 |
| 3L | 3599993 | 2 | 28 | 13 | 0 |
| 3L | 3609955 | 2 | 24 | 10 | 0 |
| 3L | 3611446 | 2 | 12 | 11 | 0 |
| 3L | 3613116 | 2 | 5 | 8 | 0 |
| 3L | 3615735 | 2 | 17 | 4 | 0 |
| 3L | 3616835 | 2 | 7 | 9 | 0 |
| 3L | 3618319 | 2 | 7 | 13 | 0 |
| 3L | 3632915 | 2 | 14 | 16 | 0 |
| 3L | 3647347 | 2 | 7 | 16 | 0 |
| 3L | 3647714 | 2 | 4 | 16 | 0 |
| 3L | 3662332 | 2 | 13 | 5 | 0 |
| 3L | 3670926 | 2 | 13 | 5 | 0 |
| 3L | 3673226 | 2 | 5 | 5 | 0 |
| 3L | 3680385 | 2 | 4 | 17 | 0 |
| 3L | 3683539 | 2 | 6 | 18 | 0 |
| 3L | 3690317 | 2 | 4 | 15 | 0 |
| 3L | 3690461 | 2 | 5 | 7 | 0 |
| 3L | 3693444 | 2 | 4 | 9 | 0 |
| 3L | 3706892 | 2 | 11 | 7 | 0 |
| 3L | 3718360 | 2 | 15 | 5 | 0 |
| 3L | 3719836 | 2 | 11 | 4 | 0 |
| 3L | 3721062 | 2 | 10 | 18 | 0 |
| 3L | 3721392 | 2 | 4 | 22 | 0 |
| 3L | 3722802 | 2 | 4 | 16 | 0 |
| 3L | 3727842 | 2 | 8 | 5 | 0 |
| 3L | 3728208 | 2 | 13 | 4 | 0 |
| 3L | 3728357 | 2 | 21 | 6 | 0 |
| 3L | 3729004 | 2 | 8 | 12 | 0 |
| 3L | 3732943 | 2 | 11 | 11 | 0 |
| 3L | 3733315 | 2 | 9 | 4 | 0 |
| 3L | 3735339 | 2 | 13 | 5 | 0 |
| 3L | 3756604 | 2 | 11 | 5 | 0 |
| 3L | 3756971 | 2 | 4 | 12 | 0 |
| 3L | 3762352 | 2 | 7 | 7 | 0 |
| 3L | 3773343 | 2 | 12 | 12 | 0 |
| 3L | 3775155 | 2 | 7 | 21 | 0 |
| 3L | 3786453 | 2 | 8 | 11 | 0 |
| 3L | 3787623 | 2 | 8 | 9 | 0 |
| 3L | 3788397 | 2 | 17 | 8 | 0 |
| 3L | 3788482 | 2 | 5 | 5 | 0 |
| 3L | 3788757 | 2 | 4 | 10 | 0 |
| 3L | 3791339 | 2 | 6 | 14 | 0 |
| 3L | 3792347 | 2 | 4 | 20 | 0 |
| 3L | 3793179 | 2 | 8 | 12 | 0 |
| 3L | 3793680 | 2 | 11 | 9 | 0 |
| 3L | 3804118 | 2 | 6 | 16 | 0 |
| 3L | 3812499 | 2 | 9 | 10 | 0 |
| 3L | 3812901 | 2 | 6 | 9 | 0 |
| 3L | 3827109 | 2 | 7 | 7 | 0 |
| 3L | 3844186 | 2 | 22 | 11 | 0 |
| 3L | 3845451 | 2 | 6 | 14 | 0 |
| 3L | 3846418 | 2 | 6 | 13 | 0 |
| 3L | 3848700 | 2 | 10 | 8 | 0 |
| 3L | 3851170 | 2 | 5 | 18 | 0 |
| 3L | 3852931 | 2 | 10 | 25 | 0 |
| 3L | 3867274 | 2 | 12 | 4 | 0 |
| 3L | 3872384 | 2 | 10 | 5 | 0 |
| 3L | 3872883 | 2 | 8 | 8 | 0 |
| 3L | 3886668 | 2 | 7 | 9 | 0 |
| 3L | 3893440 | 2 | 4 | 12 | 0 |
| 3L | 3898835 | 2 | 8 | 8 | 0 |
| 3L | 3903216 | 2 | 9 | 5 | 0 |
| 3L | 3905057 | 2 | 10 | 7 | 0 |
| 3L | 3909449 | 2 | 9 | 7 | 0 |
| 3L | 3924456 | 2 | 7 | 14 | 0 |
| 3L | 3930839 | 2 | 5 | 6 | 0 |
| 3L | 3938507 | 2 | 9 | 10 | 0 |
| 3L | 3944294 | 2 | 6 | 25 | 0 |
| 3L | 3946142 | 2 | 5 | 9 | 0 |
| 3L | 3947275 | 2 | 9 | 4 | 0 |
| 3L | 3953934 | 2 | 6 | 19 | 0 |
| 3L | 3975101 | 2 | 17 | 7 | 0 |
| 3L | 3976345 | 2 | 12 | 5 | 0 |
| 3L | 3981020 | 2 | 10 | 5 | 0 |
| 3L | 3987578 | 2 | 5 | 10 | 0 |
| 3L | 3997146 | 2 | 13 | 6 | 0 |
| 3L | 4007566 | 2 | 4 | 23 | 0 |
| 3L | 4008680 | 2 | 8 | 14 | 0 |
| 3L | 4008699 | 2 | 7 | 12 | 0 |
| 3L | 4020755 | 2 | 5 | 8 | 0 |
| 3L | 4030399 | 2 | 7 | 17 | 0 |
| 3L | 4032926 | 2 | 18 | 8 | 0 |
| 3L | 4035868 | 2 | 16 | 11 | 0 |
| 3L | 4035899 | 2 | 7 | 7 | 0 |
| 3L | 4043336 | 2 | 4 | 16 | 0 |
| 3L | 4045887 | 2 | 11 | 7 | 0 |
| 3L | 4050761 | 2 | 11 | 5 | 0 |
| 3L | 4055997 | 2 | 4 | 12 | 0 |
| 3L | 4060905 | 2 | 11 | 4 | 0 |
| 3L | 4061013 | 2 | 15 | 11 | 0 |
| 3L | 4064981 | 2 | 5 | 16 | 0 |
| 3L | 4065619 | 2 | 24 | 4 | 0 |
| 3L | 4067064 | 2 | 15 | 9 | 0 |
| 3L | 4079493 | 2 | 8 | 13 | 0 |
| 3L | 4080002 | 2 | 10 | 10 | 0 |
| 3L | 4093904 | 2 | 28 | 9 | 0 |
| 3L | 4113153 | 2 | 10 | 17 | 0 |
| 3L | 4114059 | 2 | 8 | 8 | 0 |
| 3L | 4116952 | 2 | 4 | 10 | 0 |
| 3L | 4117826 | 2 | 5 | 6 | 0 |
| 3L | 4130115 | 2 | 21 | 16 | 0 |
| 3L | 4131099 | 2 | 12 | 5 | 0 |
| 3L | 4143576 | 2 | 10 | 9 | 0 |
| 3L | 4173226 | 2 | 15 | 4 | 0 |
| 3L | 4173896 | 2 | 6 | 5 | 0 |
| 3L | 4185048 | 2 | 4 | 9 | 0 |
| 3L | 4185711 | 2 | 7 | 9 | 0 |
| 3L | 4215024 | 2 | 7 | 11 | 0 |
| 3L | 4224817 | 2 | 7 | 16 | 0 |
| 3L | 4235875 | 2 | 12 | 5 | 0 |
| 3L | 4268427 | 2 | 8 | 18 | 0 |
| 3L | 4268791 | 2 | 9 | 6 | 0 |
| 3L | 4283446 | 2 | 17 | 8 | 0 |
| 3L | 4289491 | 2 | 5 | 16 | 0 |
| 3L | 4289935 | 2 | 7 | 27 | 0 |
| 3L | 4290193 | 2 | 9 | 16 | 0 |
| 3L | 4290235 | 2 | 7 | 12 | 0 |
| 3L | 4290664 | 2 | 4 | 25 | 0 |
| 3L | 4314807 | 2 | 17 | 4 | 0 |
| 3L | 4315104 | 2 | 7 | 6 | 0 |
| 3L | 4315526 | 2 | 12 | 10 | 0 |
| 3L | 4325881 | 2 | 4 | 12 | 0 |
| 3L | 4343452 | 2 | 4 | 10 | 0 |
| 3L | 4348864 | 2 | 7 | 7 | 0 |
| 3L | 4349863 | 2 | 7 | 11 | 0 |
| 3L | 4352106 | 2 | 7 | 16 | 0 |
| 3L | 4355895 | 2 | 8 | 8 | 0 |
| 3L | 4359361 | 2 | 4 | 12 | 0 |
| 3L | 4359513 | 2 | 4 | 10 | 0 |
| 3L | 4365251 | 2 | 7 | 7 | 0 |
| 3L | 4366085 | 2 | 10 | 8 | 0 |
| 3L | 4368193 | 2 | 13 | 6 | 0 |
| 3L | 4368446 | 2 | 5 | 10 | 0 |
| 3L | 4377406 | 2 | 11 | 13 | 0 |
| 3L | 4383107 | 2 | 19 | 10 | 0 |
| 3L | 4389344 | 2 | 6 | 12 | 0 |
| 3L | 4391130 | 2 | 18 | 9 | 0 |
| 3L | 4392642 | 2 | 6 | 20 | 0 |
| 3L | 4400812 | 2 | 11 | 4 | 0 |
| 3L | 4404707 | 2 | 16 | 11 | 0 |
| 3L | 4432304 | 2 | 14 | 4 | 0 |
| 3L | 4462457 | 2 | 13 | 12 | 0 |
| 3L | 4469234 | 2 | 6 | 7 | 0 |
| 3L | 4471585 | 2 | 9 | 4 | 0 |
| 3L | 4487068 | 2 | 21 | 13 | 0 |
| 3L | 4500091 | 2 | 24 | 15 | 0 |
| 3L | 4506555 | 2 | 13 | 6 | 0 |
| 3L | 4512233 | 2 | 14 | 11 | 0 |
| 3L | 4512248 | 2 | 11 | 20 | 0 |
| 3L | 4520847 | 2 | 10 | 4 | 0 |
| 3L | 4528462 | 2 | 17 | 14 | 0 |
| 3L | 4529752 | 2 | 4 | 16 | 0 |
| 3L | 4534289 | 2 | 11 | 11 | 0 |
| 3L | 4564595 | 2 | 11 | 6 | 0 |
| 3L | 4566576 | 2 | 11 | 9 | 0 |
| 3L | 4566986 | 2 | 9 | 13 | 0 |
| 3L | 4588207 | 2 | 32 | 18 | 0 |
| 3L | 4595407 | 2 | 12 | 21 | 0 |
| 3L | 4599849 | 2 | 14 | 5 | 0 |
| 3L | 4601765 | 2 | 10 | 4 | 0 |
| 3L | 4609739 | 2 | 5 | 8 | 0 |
| 3L | 4610515 | 2 | 9 | 5 | 0 |
| 3L | 4617056 | 2 | 12 | 14 | 0 |
| 3L | 4621192 | 2 | 11 | 5 | 0 |
| 3L | 4634838 | 2 | 7 | 13 | 0 |
| 3L | 4644820 | 2 | 12 | 9 | 0 |
| 3L | 4646257 | 2 | 17 | 17 | 0 |
| 3L | 4647639 | 2 | 5 | 23 | 0 |
| 3L | 4647708 | 2 | 18 | 11 | 0 |
| 3L | 4650621 | 2 | 28 | 17 | 0 |
| 3L | 4653783 | 2 | 4 | 9 | 0 |
| 3L | 4662124 | 2 | 4 | 12 | 0 |
| 3L | 4673456 | 2 | 9 | 15 | 0 |
| 3L | 4676265 | 2 | 5 | 5 | 0 |
| 3L | 4691566 | 2 | 5 | 24 | 0 |
| 3L | 4691654 | 2 | 5 | 15 | 0 |
| 3L | 4703681 | 2 | 5 | 23 | 0 |
| 3L | 4715547 | 2 | 7 | 6 | 0 |
| 3L | 4716077 | 2 | 11 | 8 | 0 |
| 3L | 4726485 | 2 | 5 | 11 | 0 |
| 3L | 4737028 | 2 | 18 | 4 | 0 |
| 3L | 4737461 | 2 | 5 | 5 | 0 |
| 3L | 4737511 | 2 | 13 | 4 | 0 |
| 3L | 4738364 | 2 | 13 | 5 | 0 |
| 3L | 4740090 | 2 | 4 | 11 | 0 |
| 3L | 4740098 | 2 | 5 | 5 | 0 |
| 3L | 4741182 | 2 | 10 | 4 | 0 |
| 3L | 4750309 | 2 | 8 | 5 | 0 |
| 3L | 4757116 | 2 | 5 | 25 | 0 |
| 3L | 4768992 | 2 | 8 | 13 | 0 |
| 3L | 4781498 | 2 | 8 | 9 | 0 |
| 3L | 4785291 | 2 | 5 | 11 | 0 |
| 3L | 4789733 | 2 | 9 | 13 | 0 |
| 3L | 4799819 | 2 | 4 | 11 | 0 |
| 3L | 4803380 | 2 | 10 | 8 | 0 |
| 3L | 4806154 | 2 | 14 | 5 | 0 |
| 3L | 4823682 | 2 | 15 | 6 | 0 |
| 3L | 4827576 | 2 | 13 | 6 | 0 |
| 3L | 4830994 | 2 | 8 | 22 | 0 |
| 3L | 4840737 | 2 | 4 | 13 | 0 |
| 3L | 4854515 | 2 | 15 | 7 | 0 |
| 3L | 4856179 | 2 | 11 | 4 | 0 |
| 3L | 4856466 | 2 | 12 | 12 | 0 |
| 3L | 4885413 | 2 | 7 | 7 | 0 |
| 3L | 4916798 | 2 | 24 | 11 | 0 |
| 3L | 4918793 | 2 | 13 | 7 | 0 |
| 3L | 4918902 | 2 | 11 | 15 | 0 |
| 3L | 4922501 | 2 | 4 | 16 | 0 |
| 3L | 4950603 | 2 | 5 | 8 | 0 |
| 3L | 4955925 | 2 | 6 | 6 | 0 |
| 3L | 4957939 | 2 | 12 | 4 | 0 |
| 3L | 4962462 | 2 | 6 | 8 | 0 |
| 3L | 4972263 | 2 | 4 | 13 | 0 |
| 3L | 4974589 | 2 | 10 | 6 | 0 |
| 3L | 5082819 | 2 | 19 | 6 | 0 |
| 3L | 5093761 | 2 | 8 | 6 | 0 |
| 3L | 5097377 | 2 | 7 | 8 | 0 |
| 3L | 5101323 | 2 | 17 | 11 | 0 |
| 3L | 5104460 | 2 | 8 | 5 | 0 |
| 3L | 5172852 | 2 | 6 | 5 | 0 |
| 3L | 5187354 | 2 | 7 | 21 | 0 |
| 3L | 5195303 | 2 | 8 | 5 | 0 |
| 3L | 5404385 | 2 | 12 | 7 | 0 |
| 3L | 5413933 | 2 | 6 | 5 | 0 |
| 3L | 5481015 | 2 | 5 | 10 | 0 |
| 3L | 5492425 | 2 | 17 | 4 | 0 |
| 3L | 5496689 | 2 | 5 | 6 | 0 |
| 3L | 5511664 | 2 | 8 | 11 | 0 |
| 3L | 5521447 | 2 | 8 | 5 | 0 |
| 3L | 5543268 | 2 | 9 | 8 | 0 |
| 3L | 5587787 | 2 | 15 | 9 | 0 |
| 3L | 5638149 | 2 | 11 | 5 | 0 |
| 3L | 5645791 | 2 | 6 | 6 | 0 |
| 3L | 5675659 | 2 | 7 | 5 | 0 |
| 3L | 5752965 | 2 | 16 | 6 | 0 |
| 3L | 5786057 | 2 | 5 | 6 | 0 |
| 3L | 5823334 | 2 | 8 | 12 | 0 |
| 3L | 5885896 | 2 | 13 | 7 | 0 |
| 3L | 5886659 | 2 | 5 | 5 | 0 |
| 3L | 5899376 | 2 | 4 | 9 | 0 |
| 3L | 5910335 | 2 | 15 | 4 | 0 |
| 3L | 5917392 | 2 | 10 | 12 | 0 |
| 3L | 5922554 | 2 | 5 | 9 | 0 |
| 3L | 5939474 | 2 | 11 | 8 | 0 |
| 3L | 6011258 | 2 | 6 | 6 | 0 |
| 3L | 6016707 | 2 | 6 | 6 | 0 |
| 3L | 6064411 | 2 | 10 | 6 | 0 |
| 3L | 6066508 | 2 | 10 | 6 | 0 |
| 3L | 6092565 | 2 | 9 | 6 | 0 |
| 3L | 6119920 | 2 | 9 | 4 | 0 |
| 3L | 6155454 | 2 | 6 | 5 | 0 |
| 3L | 6259059 | 2 | 12 | 4 | 0 |
| 3L | 6259735 | 2 | 11 | 11 | 0 |
| 3L | 6282767 | 2 | 7 | 10 | 0 |
| 3L | 6300596 | 2 | 5 | 22 | 0 |
| 3L | 6312756 | 2 | 5 | 22 | 0 |
| 3L | 6338603 | 2 | 5 | 9 | 0 |
| 3L | 6448623 | 2 | 4 | 11 | 0 |
| 3L | 6536398 | 2 | 6 | 5 | 0 |
| 3L | 6566453 | 2 | 9 | 7 | 0 |
| 3L | 6576136 | 2 | 9 | 6 | 0 |
| 3L | 6619629 | 2 | 12 | 7 | 0 |
| 3L | 6634214 | 2 | 9 | 5 | 0 |
| 3L | 6644335 | 2 | 5 | 13 | 0 |
| 3L | 6711568 | 2 | 4 | 11 | 0 |
| 3L | 6749186 | 2 | 6 | 5 | 0 |
| 3L | 6753325 | 2 | 5 | 6 | 0 |
| 3L | 6794403 | 2 | 5 | 14 | 0 |
| 3L | 6820031 | 2 | 6 | 5 | 0 |
| 3L | 6835015 | 2 | 5 | 11 | 0 |
| 3L | 7140210 | 2 | 6 | 8 | 0 |
| 3L | 7501123 | 2 | 4 | 10 | 0 |
| 3L | 7591432 | 2 | 5 | 8 | 0 |
| 3L | 7837268 | 2 | 8 | 6 | 0 |
| 3L | 8003189 | 2 | 11 | 9 | 0 |
| 3L | 8396030 | 2 | 4 | 19 | 0 |
| 3L | 8639346 | 2 | 5 | 5 | 0 |
| 3L | 8803273 | 2 | 6 | 5 | 0 |
| 3L | 8830298 | 2 | 6 | 5 | 0 |
| 3L | 8912159 | 2 | 8 | 5 | 0 |
| 3L | 9000977 | 2 | 6 | 10 | 0 |
| 3L | 10624371 | 2 | 5 | 6 | 0 |
| 3L | 10794234 | 2 | 5 | 7 | 0 |
| 3L | 11627234 | 2 | 5 | 15 | 0 |
| 3L | 13017287 | 2 | 5 | 6 | 0 |
| 3L | 14951911 | 2 | 7 | 5 | 0 |
| 3L | 15821377 | 2 | 6 | 6 | 0 |
| 3L | 17478714 | 2 | 6 | 11 | 0 |
| 3L | 18360891 | 2 | 15 | 4 | 0 |
| 3L | 18924622 | 2 | 18 | 6 | 0 |
| 3L | 18928306 | 2 | 8 | 8 | 0 |
| 3L | 18955358 | 2 | 4 | 12 | 0 |
| 3L | 18958068 | 2 | 11 | 7 | 0 |
| 3L | 18959790 | 2 | 7 | 21 | 0 |
| 3L | 18959958 | 2 | 7 | 7 | 0 |
| 3L | 18960203 | 2 | 6 | 5 | 0 |
| 3L | 19011791 | 2 | 9 | 9 | 0 |
| 3L | 19020074 | 2 | 5 | 16 | 0 |
| 3L | 19032638 | 2 | 4 | 12 | 0 |
| 3L | 19037206 | 2 | 14 | 10 | 0 |
| 3L | 19038340 | 2 | 5 | 6 | 0 |
| 3L | 19062863 | 2 | 7 | 7 | 0 |
| 3L | 19065589 | 2 | 5 | 26 | 0 |
| 3L | 19066641 | 2 | 11 | 14 | 0 |
| 3L | 19084377 | 2 | 6 | 9 | 0 |
| 3L | 19097149 | 2 | 14 | 8 | 0 |
| 3L | 19119592 | 2 | 5 | 9 | 0 |
| 3L | 19123515 | 2 | 18 | 5 | 0 |
| 3L | 19132069 | 2 | 5 | 12 | 0 |
| 3L | 19157025 | 2 | 4 | 11 | 0 |
| 3L | 19174461 | 2 | 9 | 8 | 0 |
| 3L | 19178467 | 2 | 6 | 6 | 0 |
| 3L | 19182884 | 2 | 5 | 5 | 0 |
| 3L | 19185222 | 2 | 5 | 6 | 0 |
| 3L | 19208795 | 2 | 8 | 6 | 0 |
| 3L | 19235922 | 2 | 6 | 7 | 0 |
| 3L | 19249262 | 2 | 6 | 13 | 0 |
| 3L | 19258689 | 2 | 5 | 11 | 0 |
| 3L | 19282500 | 2 | 10 | 17 | 0 |
| 3L | 19285762 | 2 | 10 | 18 | 0 |
| 3L | 19326857 | 2 | 13 | 5 | 0 |
| 3L | 19337030 | 2 | 10 | 9 | 0 |
| 3L | 19429218 | 2 | 9 | 12 | 0 |
| 3L | 19447725 | 2 | 10 | 5 | 0 |
| 3L | 19457159 | 2 | 6 | 7 | 0 |
| 3L | 19492291 | 2 | 9 | 4 | 0 |
| 3L | 19658887 | 2 | 9 | 6 | 0 |
| 3L | 19850821 | 2 | 6 | 12 | 0 |
| 3L | 19916911 | 2 | 21 | 8 | 0 |
| 3L | 19952604 | 2 | 8 | 6 | 0 |
| 3L | 20046641 | 2 | 7 | 12 | 0 |
| 3L | 20049258 | 2 | 15 | 7 | 0 |
| 3L | 20049344 | 2 | 18 | 9 | 0 |
| 3L | 20057369 | 2 | 4 | 9 | 0 |
| 3L | 20061015 | 2 | 6 | 6 | 0 |
| 3L | 20077885 | 2 | 5 | 11 | 0 |
| 3L | 20088687 | 2 | 20 | 26 | 0 |
| 3L | 20095756 | 2 | 8 | 7 | 0 |
| 3L | 20142432 | 2 | 10 | 7 | 0 |
| 3L | 20163359 | 2 | 9 | 4 | 0 |
| 3L | 20204278 | 2 | 6 | 6 | 0 |
| 3L | 20212064 | 2 | 5 | 6 | 0 |
| 3L | 20231729 | 2 | 6 | 6 | 0 |
| 3L | 20255821 | 2 | 8 | 8 | 0 |
| 3L | 20286506 | 2 | 6 | 22 | 0 |
| 3L | 20306174 | 2 | 5 | 13 | 0 |
| 3L | 20339421 | 2 | 8 | 21 | 0 |
| 3L | 20406236 | 2 | 4 | 21 | 0 |
| 3L | 20407457 | 2 | 9 | 8 | 0 |
| 3L | 20408468 | 2 | 12 | 4 | 0 |
| 3L | 20449226 | 2 | 10 | 20 | 0 |
| 3L | 20467709 | 2 | 6 | 14 | 0 |
| 3L | 20478193 | 2 | 31 | 10 | 0 |
| 3L | 20661974 | 2 | 4 | 9 | 0 |
| 3L | 20669732 | 2 | 4 | 11 | 0 |
| 3L | 20710240 | 2 | 10 | 6 | 0 |
| 3L | 20718546 | 2 | 8 | 6 | 0 |
| 3L | 20858481 | 2 | 11 | 5 | 0 |
| 3L | 20900260 | 2 | 5 | 5 | 0 |
| 3L | 20900406 | 2 | 12 | 7 | 0 |
| 3L | 21045625 | 2 | 6 | 5 | 0 |
| 3L | 21054501 | 2 | 5 | 14 | 0 |
| 3L | 21063560 | 2 | 7 | 7 | 0 |
| 3L | 21185102 | 2 | 9 | 4 | 0 |
| 3L | 21200757 | 2 | 29 | 5 | 0 |
| 3L | 21204342 | 2 | 9 | 5 | 0 |
| 3L | 21222522 | 2 | 14 | 5 | 0 |
| 3L | 21229786 | 2 | 5 | 7 | 0 |
| 3L | 21249779 | 2 | 11 | 6 | 0 |
| 3L | 21282407 | 2 | 4 | 15 | 0 |
| 3L | 21365562 | 2 | 10 | 5 | 0 |
| 3L | 21409971 | 2 | 4 | 13 | 0 |
| 3L | 21456269 | 2 | 5 | 5 | 0 |
| 3L | 21569832 | 2 | 5 | 16 | 0 |
| 3L | 21583604 | 2 | 4 | 19 | 0 |
| 3L | 21622273 | 2 | 15 | 6 | 0 |
| 3L | 21625080 | 2 | 15 | 9 | 0 |
| 3L | 21627016 | 2 | 6 | 7 | 0 |
| 3L | 21629670 | 2 | 5 | 7 | 0 |
| 3L | 21631208 | 2 | 6 | 10 | 0 |
| 3L | 21632185 | 2 | 23 | 14 | 0 |
| 3L | 21657439 | 2 | 16 | 5 | 0 |
| 3L | 21663937 | 2 | 5 | 9 | 0 |
| 3L | 21668796 | 2 | 8 | 5 | 0 |
| 3L | 21772176 | 2 | 11 | 4 | 0 |
| 3L | 21773671 | 2 | 10 | 5 | 0 |
| 3L | 21797113 | 2 | 8 | 25 | 0 |
| 3L | 21850174 | 2 | 4 | 9 | 0 |
| 3L | 21883641 | 2 | 7 | 21 | 0 |
| 3L | 21884717 | 2 | 5 | 11 | 0 |
| 3L | 21920735 | 2 | 7 | 5 | 0 |
| 3L | 21926271 | 2 | 24 | 11 | 0 |
| 3L | 21928511 | 2 | 16 | 4 | 0 |
| 3L | 21973356 | 2 | 10 | 6 | 0 |
| 3L | 21996031 | 2 | 8 | 10 | 0 |
| 3L | 21998210 | 2 | 9 | 5 | 0 |
| 3L | 22056493 | 2 | 8 | 11 | 0 |
| 3L | 22115189 | 2 | 10 | 8 | 0 |
| 3L | 22115196 | 2 | 8 | 6 | 0 |
| 3L | 22115469 | 2 | 5 | 11 | 0 |
| 3L | 22158157 | 2 | 8 | 9 | 0 |
| 3L | 22170833 | 2 | 4 | 9 | 0 |
| 3L | 22187901 | 2 | 17 | 7 | 0 |
| 3L | 22205376 | 2 | 13 | 5 | 0 |
| 3L | 22207224 | 2 | 7 | 7 | 0 |
| 3L | 22234470 | 2 | 18 | 12 | 0 |
| 3L | 22248593 | 2 | 14 | 4 | 0 |
| 3L | 22273050 | 2 | 13 | 4 | 0 |
| 3L | 22304042 | 2 | 10 | 4 | 0 |
| 3L | 22307227 | 2 | 11 | 4 | 0 |
| 3L | 22314782 | 2 | 9 | 4 | 0 |
| 3L | 22346505 | 2 | 14 | 11 | 0 |
| 3L | 22358743 | 2 | 8 | 10 | 0 |
| 3L | 22378307 | 2 | 12 | 9 | 0 |
| 3L | 22393962 | 2 | 9 | 5 | 0 |
| 3L | 22454899 | 2 | 20 | 6 | 0 |
| 3L | 22463789 | 2 | 8 | 5 | 0 |
| 3L | 22488317 | 2 | 12 | 16 | 0 |
| 3L | 22586300 | 2 | 4 | 9 | 0 |
| 3L | 22606238 | 2 | 12 | 7 | 0 |
| 3L | 22609193 | 2 | 23 | 4 | 0 |
| 3L | 22627639 | 2 | 7 | 6 | 0 |
| 3L | 22628885 | 2 | 7 | 15 | 0 |
| 3L | 22631538 | 2 | 8 | 7 | 0 |
| 3L | 22631610 | 2 | 6 | 11 | 0 |
| 3L | 22631673 | 2 | 14 | 6 | 0 |
| 3L | 22645479 | 2 | 15 | 7 | 0 |
| 3L | 22650605 | 2 | 8 | 10 | 0 |
| 3L | 22667716 | 2 | 7 | 10 | 0 |
| 3L | 22677848 | 2 | 6 | 11 | 0 |
| 3L | 22682330 | 2 | 10 | 11 | 0 |
| 3L | 22746063 | 2 | 7 | 7 | 0 |
| 3L | 22839245 | 2 | 9 | 4 | 0 |
| 3L | 22881580 | 2 | 9 | 22 | 0 |
| 3L | 22899207 | 2 | 6 | 7 | 0 |
| 3L | 22899380 | 2 | 4 | 17 | 0 |
| 3L | 22920116 | 2 | 9 | 4 | 0 |
| 3L | 22940041 | 2 | 7 | 6 | 0 |
| 3L | 22960178 | 2 | 7 | 10 | 0 |
| 3L | 22962733 | 2 | 7 | 11 | 0 |
| 3L | 23051871 | 2 | 13 | 6 | 0 |
| 3L | 23061964 | 2 | 13 | 5 | 0 |
| 3L | 23146999 | 2 | 10 | 4 | 0 |
| 3L | 23224329 | 2 | 7 | 17 | 0 |
| 3L | 23256285 | 2 | 8 | 14 | 0 |
| 3L | 23393609 | 2 | 7 | 8 | 0 |
| 3L | 23448250 | 2 | 4 | 12 | 0 |
| 3L | 23463788 | 2 | 6 | 12 | 0 |
| 3L | 23471445 | 2 | 19 | 5 | 0 |
| 3L | 23496681 | 2 | 12 | 6 | 0 |
| 3L | 23605025 | 2 | 9 | 7 | 0 |
| 3L | 23713341 | 2 | 10 | 17 | 0 |
| 3L | 23735341 | 2 | 4 | 12 | 0 |
| 3L | 23841621 | 2 | 7 | 5 | 0 |
| 3L | 23958072 | 2 | 7 | 11 | 0 |
| 3L | 24178032 | 2 | 8 | 9 | 0 |
| 3L | 24329353 | 2 | 8 | 5 | 0 |
| 3L | 24520727 | 2 | 6 | 8 | 0 |
| 3L | 24530497 | 2 | 13 | 4 | 0 |
| 3R | 30845 | 2 | 15 | 7 | 0 |
| 3R | 129054 | 2 | 12 | 7 | 0 |
| 3R | 146485 | 2 | 5 | 13 | 0 |
| 3R | 173772 | 2 | 4 | 11 | 0 |
| 3R | 201358 | 2 | 11 | 23 | 0 |
| 3R | 223996 | 2 | 5 | 13 | 0 |
| 3R | 348786 | 2 | 14 | 19 | 0 |
| 3R | 348851 | 2 | 13 | 10 | 0 |
| 3R | 354810 | 2 | 11 | 10 | 0 |
| 3R | 367961 | 2 | 7 | 6 | 0 |
| 3R | 368261 | 2 | 16 | 26 | 0 |
| 3R | 370997 | 2 | 8 | 5 | 0 |
| 3R | 372921 | 2 | 12 | 8 | 0 |
| 3R | 377395 | 2 | 21 | 4 | 0 |
| 3R | 384261 | 2 | 5 | 5 | 0 |
| 3R | 386873 | 2 | 23 | 5 | 0 |
| 3R | 396766 | 2 | 9 | 10 | 0 |
| 3R | 397139 | 2 | 12 | 13 | 0 |
| 3R | 398931 | 2 | 7 | 5 | 0 |
| 3R | 400475 | 2 | 15 | 8 | 0 |
| 3R | 401025 | 2 | 8 | 12 | 0 |
| 3R | 402782 | 2 | 9 | 18 | 0 |
| 3R | 414074 | 2 | 12 | 9 | 0 |
| 3R | 416643 | 2 | 5 | 5 | 0 |
| 3R | 417713 | 2 | 18 | 7 | 0 |
| 3R | 417809 | 2 | 8 | 6 | 0 |
| 3R | 419144 | 2 | 14 | 5 | 0 |
| 3R | 420348 | 2 | 13 | 8 | 0 |
| 3R | 422867 | 2 | 13 | 9 | 0 |
| 3R | 432657 | 2 | 7 | 6 | 0 |
| 3R | 439292 | 2 | 8 | 9 | 0 |
| 3R | 441372 | 2 | 12 | 9 | 0 |
| 3R | 446388 | 2 | 10 | 5 | 0 |
| 3R | 446713 | 2 | 21 | 8 | 0 |
| 3R | 446750 | 2 | 9 | 4 | 0 |
| 3R | 456051 | 2 | 9 | 10 | 0 |
| 3R | 457008 | 2 | 9 | 5 | 0 |
| 3R | 458972 | 2 | 12 | 5 | 0 |
| 3R | 466382 | 2 | 14 | 6 | 0 |
| 3R | 468409 | 2 | 17 | 10 | 0 |
| 3R | 474358 | 2 | 8 | 5 | 0 |
| 3R | 475240 | 2 | 12 | 18 | 0 |
| 3R | 486393 | 2 | 18 | 12 | 0 |
| 3R | 490656 | 2 | 10 | 13 | 0 |
| 3R | 490795 | 2 | 14 | 4 | 0 |
| 3R | 498327 | 2 | 8 | 21 | 0 |
| 3R | 504941 | 2 | 7 | 12 | 0 |
| 3R | 506554 | 2 | 13 | 12 | 0 |
| 3R | 511618 | 2 | 4 | 28 | 0 |
| 3R | 529962 | 2 | 12 | 6 | 0 |
| 3R | 561354 | 2 | 8 | 5 | 0 |
| 3R | 570114 | 2 | 11 | 8 | 0 |
| 3R | 570979 | 2 | 11 | 6 | 0 |
| 3R | 590126 | 2 | 5 | 25 | 0 |
| 3R | 610453 | 2 | 11 | 5 | 0 |
| 3R | 625904 | 2 | 6 | 17 | 0 |
| 3R | 663695 | 2 | 18 | 4 | 0 |
| 3R | 675512 | 2 | 5 | 7 | 0 |
| 3R | 792829 | 2 | 4 | 13 | 0 |
| 3R | 802955 | 2 | 6 | 13 | 0 |
| 3R | 803647 | 2 | 8 | 12 | 0 |
| 3R | 821920 | 2 | 4 | 13 | 0 |
| 3R | 825322 | 2 | 9 | 7 | 0 |
| 3R | 838365 | 2 | 10 | 24 | 0 |
| 3R | 839321 | 2 | 9 | 5 | 0 |
| 3R | 852209 | 2 | 4 | 11 | 0 |
| 3R | 879490 | 2 | 14 | 9 | 0 |
| 3R | 880399 | 2 | 11 | 5 | 0 |
| 3R | 886023 | 2 | 9 | 5 | 0 |
| 3R | 886146 | 2 | 5 | 9 | 0 |
| 3R | 907147 | 2 | 5 | 5 | 0 |
| 3R | 932325 | 2 | 9 | 6 | 0 |
| 3R | 934374 | 2 | 11 | 21 | 0 |
| 3R | 951517 | 2 | 13 | 10 | 0 |
| 3R | 952339 | 2 | 5 | 8 | 0 |
| 3R | 960522 | 2 | 8 | 6 | 0 |
| 3R | 972526 | 2 | 11 | 11 | 0 |
| 3R | 976374 | 2 | 12 | 23 | 0 |
| 3R | 996022 | 2 | 6 | 7 | 0 |
| 3R | 999585 | 2 | 8 | 10 | 0 |
| 3R | 1011406 | 2 | 4 | 16 | 0 |
| 3R | 1026043 | 2 | 16 | 12 | 0 |
| 3R | 1036976 | 2 | 16 | 7 | 0 |
| 3R | 1046590 | 2 | 14 | 4 | 0 |
| 3R | 1057633 | 2 | 11 | 5 | 0 |
| 3R | 1070257 | 2 | 14 | 8 | 0 |
| 3R | 1074737 | 2 | 10 | 4 | 0 |
| 3R | 1083742 | 2 | 10 | 18 | 0 |
| 3R | 1109319 | 2 | 13 | 8 | 0 |
| 3R | 1116113 | 2 | 10 | 13 | 0 |
| 3R | 1116811 | 2 | 16 | 14 | 0 |
| 3R | 1125241 | 2 | 7 | 6 | 0 |
| 3R | 1130075 | 2 | 6 | 9 | 0 |
| 3R | 1132347 | 2 | 23 | 16 | 0 |
| 3R | 1132985 | 2 | 12 | 4 | 0 |
| 3R | 1141222 | 2 | 8 | 15 | 0 |
| 3R | 1151890 | 2 | 7 | 6 | 0 |
| 3R | 1189081 | 2 | 26 | 11 | 0 |
| 3R | 1199299 | 2 | 11 | 7 | 0 |
| 3R | 1210780 | 2 | 8 | 5 | 0 |
| 3R | 1252405 | 2 | 8 | 5 | 0 |
| 3R | 1255523 | 2 | 18 | 11 | 0 |
| 3R | 1256048 | 2 | 15 | 7 | 0 |
| 3R | 1261315 | 2 | 12 | 4 | 0 |
| 3R | 1265143 | 2 | 12 | 16 | 0 |
| 3R | 1266966 | 2 | 10 | 6 | 0 |
| 3R | 1270986 | 2 | 10 | 4 | 0 |
| 3R | 1271336 | 2 | 5 | 20 | 0 |
| 3R | 1278168 | 2 | 11 | 8 | 0 |
| 3R | 1279566 | 2 | 22 | 6 | 0 |
| 3R | 1279622 | 2 | 14 | 9 | 0 |
| 3R | 1283893 | 2 | 8 | 21 | 0 |
| 3R | 1289300 | 2 | 10 | 6 | 0 |
| 3R | 1289800 | 2 | 17 | 8 | 0 |
| 3R | 1292381 | 2 | 12 | 12 | 0 |
| 3R | 1295900 | 2 | 26 | 6 | 0 |
| 3R | 1300761 | 2 | 12 | 5 | 0 |
| 3R | 1310118 | 2 | 9 | 4 | 0 |
| 3R | 1313135 | 2 | 9 | 16 | 0 |
| 3R | 1322505 | 2 | 21 | 4 | 0 |
| 3R | 1327881 | 2 | 16 | 4 | 0 |
| 3R | 1329956 | 2 | 13 | 6 | 0 |
| 3R | 1332219 | 2 | 10 | 7 | 0 |
| 3R | 1332514 | 2 | 11 | 6 | 0 |
| 3R | 1337721 | 2 | 27 | 11 | 0 |
| 3R | 1343253 | 2 | 5 | 16 | 0 |
| 3R | 1404043 | 2 | 7 | 6 | 0 |
| 3R | 1409251 | 2 | 6 | 5 | 0 |
| 3R | 1409441 | 2 | 19 | 13 | 0 |
| 3R | 1443727 | 2 | 7 | 6 | 0 |
| 3R | 1470967 | 2 | 18 | 5 | 0 |
| 3R | 1470996 | 2 | 19 | 6 | 0 |
| 3R | 1473363 | 2 | 10 | 12 | 0 |
| 3R | 1485809 | 2 | 4 | 21 | 0 |
| 3R | 1508259 | 2 | 10 | 9 | 0 |
| 3R | 1525849 | 2 | 6 | 13 | 0 |
| 3R | 1531569 | 2 | 6 | 23 | 0 |
| 3R | 1532218 | 2 | 16 | 8 | 0 |
| 3R | 1554782 | 2 | 5 | 6 | 0 |
| 3R | 1556182 | 2 | 14 | 8 | 0 |
| 3R | 1558054 | 2 | 6 | 16 | 0 |
| 3R | 1572112 | 2 | 4 | 18 | 0 |
| 3R | 1583583 | 2 | 12 | 4 | 0 |
| 3R | 1590047 | 2 | 17 | 4 | 0 |
| 3R | 1600481 | 2 | 9 | 5 | 0 |
| 3R | 1603277 | 2 | 7 | 5 | 0 |
| 3R | 1614610 | 2 | 5 | 22 | 0 |
| 3R | 1626527 | 2 | 15 | 8 | 0 |
| 3R | 1694891 | 2 | 5 | 10 | 0 |
| 3R | 1705851 | 2 | 5 | 10 | 0 |
| 3R | 1706794 | 2 | 9 | 11 | 0 |
| 3R | 1710352 | 2 | 18 | 17 | 0 |
| 3R | 1713624 | 2 | 5 | 16 | 0 |
| 3R | 1723217 | 2 | 20 | 5 | 0 |
| 3R | 1728583 | 2 | 8 | 21 | 0 |
| 3R | 1732756 | 2 | 7 | 5 | 0 |
| 3R | 1744683 | 2 | 4 | 16 | 0 |
| 3R | 1744992 | 2 | 17 | 10 | 0 |
| 3R | 1755775 | 2 | 14 | 8 | 0 |
| 3R | 1761400 | 2 | 18 | 5 | 0 |
| 3R | 1761555 | 2 | 15 | 18 | 0 |
| 3R | 1775598 | 2 | 5 | 21 | 0 |
| 3R | 1778767 | 2 | 15 | 5 | 0 |
| 3R | 1779179 | 2 | 14 | 7 | 0 |
| 3R | 1779206 | 2 | 7 | 10 | 0 |
| 3R | 1779252 | 2 | 15 | 8 | 0 |
| 3R | 1791682 | 2 | 11 | 19 | 0 |
| 3R | 1795276 | 2 | 14 | 5 | 0 |
| 3R | 1813345 | 2 | 12 | 5 | 0 |
| 3R | 1830985 | 2 | 13 | 8 | 0 |
| 3R | 1835546 | 2 | 26 | 15 | 0 |
| 3R | 1837267 | 2 | 12 | 5 | 0 |
| 3R | 1839119 | 2 | 11 | 6 | 0 |
| 3R | 1840406 | 2 | 22 | 6 | 0 |
| 3R | 1853984 | 2 | 8 | 5 | 0 |
| 3R | 1854190 | 2 | 23 | 5 | 0 |
| 3R | 1856875 | 2 | 23 | 12 | 0 |
| 3R | 1858291 | 2 | 12 | 4 | 0 |
| 3R | 1864450 | 2 | 7 | 9 | 0 |
| 3R | 1873105 | 2 | 11 | 5 | 0 |
| 3R | 1874128 | 2 | 4 | 11 | 0 |
| 3R | 1886063 | 2 | 17 | 10 | 0 |
| 3R | 1889447 | 2 | 13 | 7 | 0 |
| 3R | 1889632 | 2 | 4 | 12 | 0 |
| 3R | 1900677 | 2 | 23 | 6 | 0 |
| 3R | 1900775 | 2 | 10 | 5 | 0 |
| 3R | 1901484 | 2 | 15 | 6 | 0 |
| 3R | 1904115 | 2 | 13 | 4 | 0 |
| 3R | 1913258 | 2 | 6 | 13 | 0 |
| 3R | 1919062 | 2 | 11 | 4 | 0 |
| 3R | 1921527 | 2 | 11 | 21 | 0 |
| 3R | 1922647 | 2 | 8 | 5 | 0 |
| 3R | 1946041 | 2 | 8 | 7 | 0 |
| 3R | 1954322 | 2 | 20 | 6 | 0 |
| 3R | 1956520 | 2 | 11 | 10 | 0 |
| 3R | 1957709 | 2 | 14 | 5 | 0 |
| 3R | 1959779 | 2 | 12 | 10 | 0 |
| 3R | 1974739 | 2 | 12 | 4 | 0 |
| 3R | 1978298 | 2 | 7 | 20 | 0 |
| 3R | 1979507 | 2 | 15 | 7 | 0 |
| 3R | 2016656 | 2 | 5 | 6 | 0 |
| 3R | 2018626 | 2 | 5 | 13 | 0 |
| 3R | 2019879 | 2 | 10 | 7 | 0 |
| 3R | 2026210 | 2 | 10 | 8 | 0 |
| 3R | 2027750 | 2 | 5 | 6 | 0 |
| 3R | 2028515 | 2 | 16 | 10 | 0 |
| 3R | 2030149 | 2 | 16 | 11 | 0 |
| 3R | 2032661 | 2 | 18 | 10 | 0 |
| 3R | 2044973 | 2 | 18 | 14 | 0 |
| 3R | 2052453 | 2 | 19 | 13 | 0 |
| 3R | 2076946 | 2 | 10 | 12 | 0 |
| 3R | 2102627 | 2 | 11 | 6 | 0 |
| 3R | 2110023 | 2 | 7 | 7 | 0 |
| 3R | 2111544 | 2 | 16 | 4 | 0 |
| 3R | 2139791 | 2 | 15 | 6 | 0 |
| 3R | 2142356 | 2 | 9 | 6 | 0 |
| 3R | 2142500 | 2 | 11 | 15 | 0 |
| 3R | 2143267 | 2 | 5 | 5 | 0 |
| 3R | 2153501 | 2 | 11 | 9 | 0 |
| 3R | 2154002 | 2 | 11 | 15 | 0 |
| 3R | 2162631 | 2 | 6 | 5 | 0 |
| 3R | 2168589 | 2 | 8 | 5 | 0 |
| 3R | 2172617 | 2 | 14 | 4 | 0 |
| 3R | 2181967 | 2 | 19 | 12 | 0 |
| 3R | 2183992 | 2 | 14 | 6 | 0 |
| 3R | 2186513 | 2 | 17 | 16 | 0 |
| 3R | 2192056 | 2 | 11 | 8 | 0 |
| 3R | 2195528 | 2 | 9 | 13 | 0 |
| 3R | 2195591 | 2 | 10 | 22 | 0 |
| 3R | 2197966 | 2 | 17 | 14 | 0 |
| 3R | 2200354 | 2 | 9 | 7 | 0 |
| 3R | 2209896 | 2 | 15 | 6 | 0 |
| 3R | 2221245 | 2 | 15 | 6 | 0 |
| 3R | 2232214 | 2 | 6 | 12 | 0 |
| 3R | 2240943 | 2 | 4 | 19 | 0 |
| 3R | 2257463 | 2 | 9 | 6 | 0 |
| 3R | 2266228 | 2 | 10 | 11 | 0 |
| 3R | 2266687 | 2 | 12 | 7 | 0 |
| 3R | 2280468 | 2 | 6 | 6 | 0 |
| 3R | 2285882 | 2 | 13 | 8 | 0 |
| 3R | 2305567 | 2 | 6 | 13 | 0 |
| 3R | 2310928 | 2 | 17 | 8 | 0 |
| 3R | 2312427 | 2 | 8 | 8 | 0 |
| 3R | 2314605 | 2 | 13 | 21 | 0 |
| 3R | 2325379 | 2 | 9 | 5 | 0 |
| 3R | 2329283 | 2 | 12 | 5 | 0 |
| 3R | 2344785 | 2 | 4 | 10 | 0 |
| 3R | 2346819 | 2 | 9 | 6 | 0 |
| 3R | 2347028 | 2 | 21 | 7 | 0 |
| 3R | 2350180 | 2 | 8 | 17 | 0 |
| 3R | 2351805 | 2 | 11 | 8 | 0 |
| 3R | 2352794 | 2 | 15 | 7 | 0 |
| 3R | 2357034 | 2 | 9 | 8 | 0 |
| 3R | 2357940 | 2 | 7 | 11 | 0 |
| 3R | 2371101 | 2 | 8 | 11 | 0 |
| 3R | 2372945 | 2 | 7 | 5 | 0 |
| 3R | 2374073 | 2 | 8 | 5 | 0 |
| 3R | 2380759 | 2 | 10 | 6 | 0 |
| 3R | 2393041 | 2 | 4 | 14 | 0 |
| 3R | 2397149 | 2 | 5 | 5 | 0 |
| 3R | 2413300 | 2 | 10 | 7 | 0 |
| 3R | 2424304 | 2 | 7 | 6 | 0 |
| 3R | 2426861 | 2 | 15 | 7 | 0 |
| 3R | 2431700 | 2 | 11 | 9 | 0 |
| 3R | 2453689 | 2 | 7 | 9 | 0 |
| 3R | 2455975 | 2 | 12 | 11 | 0 |
| 3R | 2459534 | 2 | 12 | 7 | 0 |
| 3R | 2466690 | 2 | 8 | 6 | 0 |
| 3R | 2466787 | 2 | 9 | 8 | 0 |
| 3R | 2472706 | 2 | 9 | 4 | 0 |
| 3R | 2478759 | 2 | 15 | 16 | 0 |
| 3R | 2500888 | 2 | 21 | 9 | 0 |
| 3R | 2501362 | 2 | 8 | 6 | 0 |
| 3R | 2537453 | 2 | 8 | 5 | 0 |
| 3R | 2551948 | 2 | 4 | 11 | 0 |
| 3R | 2552409 | 2 | 10 | 5 | 0 |
| 3R | 2568501 | 2 | 11 | 7 | 0 |
| 3R | 2568589 | 2 | 12 | 14 | 0 |
| 3R | 2573613 | 2 | 15 | 11 | 0 |
| 3R | 2577666 | 2 | 11 | 7 | 0 |
| 3R | 2580309 | 2 | 12 | 4 | 0 |
| 3R | 2587170 | 2 | 9 | 4 | 0 |
| 3R | 2587763 | 2 | 13 | 4 | 0 |
| 3R | 2589269 | 2 | 9 | 11 | 0 |
| 3R | 2591530 | 2 | 25 | 9 | 0 |
| 3R | 2603241 | 2 | 9 | 5 | 0 |
| 3R | 2674420 | 2 | 5 | 10 | 0 |
| 3R | 2680255 | 2 | 14 | 16 | 0 |
| 3R | 2683253 | 2 | 9 | 4 | 0 |
| 3R | 2686154 | 2 | 6 | 15 | 0 |
| 3R | 2686169 | 2 | 7 | 8 | 0 |
| 3R | 2688335 | 2 | 10 | 9 | 0 |
| 3R | 2692106 | 2 | 17 | 14 | 0 |
| 3R | 2692506 | 2 | 5 | 5 | 0 |
| 3R | 2693627 | 2 | 17 | 17 | 0 |
| 3R | 2694775 | 2 | 8 | 5 | 0 |
| 3R | 2697159 | 2 | 14 | 9 | 0 |
| 3R | 2697773 | 2 | 11 | 5 | 0 |
| 3R | 2700405 | 2 | 13 | 12 | 0 |
| 3R | 2701957 | 2 | 17 | 9 | 0 |
| 3R | 2702853 | 2 | 15 | 8 | 0 |
| 3R | 2703903 | 2 | 5 | 11 | 0 |
| 3R | 2705887 | 2 | 6 | 8 | 0 |
| 3R | 2707463 | 2 | 8 | 5 | 0 |
| 3R | 2708245 | 2 | 11 | 16 | 0 |
| 3R | 2709113 | 2 | 5 | 10 | 0 |
| 3R | 2710002 | 2 | 5 | 16 | 0 |
| 3R | 2716446 | 2 | 11 | 19 | 0 |
| 3R | 2716689 | 2 | 11 | 15 | 0 |
| 3R | 2720420 | 2 | 14 | 6 | 0 |
| 3R | 2731767 | 2 | 5 | 15 | 0 |
| 3R | 2748004 | 2 | 6 | 19 | 0 |
| 3R | 2748198 | 2 | 5 | 7 | 0 |
| 3R | 2764365 | 2 | 10 | 8 | 0 |
| 3R | 2767568 | 2 | 11 | 7 | 0 |
| 3R | 2770572 | 2 | 7 | 6 | 0 |
| 3R | 2773629 | 2 | 8 | 14 | 0 |
| 3R | 2798978 | 2 | 6 | 7 | 0 |
| 3R | 2805417 | 2 | 9 | 16 | 0 |
| 3R | 2813642 | 2 | 15 | 7 | 0 |
| 3R | 2816089 | 2 | 23 | 5 | 0 |
| 3R | 2832797 | 2 | 17 | 7 | 0 |
| 3R | 2840984 | 2 | 10 | 11 | 0 |
| 3R | 2862070 | 2 | 8 | 5 | 0 |
| 3R | 2862778 | 2 | 18 | 9 | 0 |
| 3R | 2876702 | 2 | 21 | 17 | 0 |
| 3R | 2888602 | 2 | 7 | 8 | 0 |
| 3R | 2891049 | 2 | 17 | 8 | 0 |
| 3R | 2894377 | 2 | 12 | 24 | 0 |
| 3R | 2897201 | 2 | 9 | 15 | 0 |
| 3R | 2907931 | 2 | 15 | 7 | 0 |
| 3R | 2920166 | 2 | 6 | 5 | 0 |
| 3R | 2923506 | 2 | 12 | 5 | 0 |
| 3R | 2965905 | 2 | 6 | 6 | 0 |
| 3R | 2983801 | 2 | 10 | 6 | 0 |
| 3R | 2985886 | 2 | 5 | 20 | 0 |
| 3R | 2988443 | 2 | 8 | 5 | 0 |
| 3R | 2999103 | 2 | 18 | 10 | 0 |
| 3R | 3014006 | 2 | 20 | 7 | 0 |
| 3R | 3036781 | 2 | 6 | 5 | 0 |
| 3R | 3040781 | 2 | 6 | 16 | 0 |
| 3R | 3060824 | 2 | 24 | 18 | 0 |
| 3R | 3061314 | 2 | 7 | 9 | 0 |
| 3R | 3077833 | 2 | 9 | 5 | 0 |
| 3R | 3078885 | 2 | 21 | 11 | 0 |
| 3R | 3097821 | 2 | 4 | 20 | 0 |
| 3R | 3120201 | 2 | 19 | 14 | 0 |
| 3R | 3127377 | 2 | 4 | 12 | 0 |
| 3R | 3136355 | 2 | 12 | 4 | 0 |
| 3R | 3156201 | 2 | 8 | 13 | 0 |
| 3R | 3157582 | 2 | 13 | 16 | 0 |
| 3R | 3161915 | 2 | 4 | 13 | 0 |
| 3R | 3167321 | 2 | 5 | 19 | 0 |
| 3R | 3168325 | 2 | 5 | 11 | 0 |
| 3R | 3195988 | 2 | 12 | 15 | 0 |
| 3R | 3219561 | 2 | 8 | 9 | 0 |
| 3R | 3220248 | 2 | 9 | 6 | 0 |
| 3R | 3247171 | 2 | 20 | 24 | 0 |
| 3R | 3248014 | 2 | 8 | 5 | 0 |
| 3R | 3248074 | 2 | 13 | 10 | 0 |
| 3R | 3248587 | 2 | 17 | 5 | 0 |
| 3R | 3261085 | 2 | 10 | 14 | 0 |
| 3R | 3264805 | 2 | 14 | 4 | 0 |
| 3R | 3265270 | 2 | 4 | 9 | 0 |
| 3R | 3267900 | 2 | 22 | 8 | 0 |
| 3R | 3270952 | 2 | 5 | 26 | 0 |
| 3R | 3271116 | 2 | 7 | 18 | 0 |
| 3R | 3273092 | 2 | 10 | 7 | 0 |
| 3R | 3281209 | 2 | 14 | 14 | 0 |
| 3R | 3290693 | 2 | 24 | 12 | 0 |
| 3R | 3304175 | 2 | 4 | 10 | 0 |
| 3R | 3321515 | 2 | 19 | 9 | 0 |
| 3R | 3324254 | 2 | 13 | 5 | 0 |
| 3R | 3327872 | 2 | 18 | 10 | 0 |
| 3R | 3330983 | 2 | 18 | 10 | 0 |
| 3R | 3334456 | 2 | 20 | 5 | 0 |
| 3R | 3344379 | 2 | 6 | 5 | 0 |
| 3R | 3359421 | 2 | 4 | 16 | 0 |
| 3R | 3366457 | 2 | 13 | 5 | 0 |
| 3R | 3371528 | 2 | 16 | 5 | 0 |
| 3R | 3375894 | 2 | 12 | 6 | 0 |
| 3R | 3377523 | 2 | 16 | 6 | 0 |
| 3R | 3399929 | 2 | 11 | 7 | 0 |
| 3R | 3402373 | 2 | 16 | 6 | 0 |
| 3R | 3409309 | 2 | 14 | 7 | 0 |
| 3R | 3414753 | 2 | 7 | 6 | 0 |
| 3R | 3418663 | 2 | 17 | 4 | 0 |
| 3R | 3422067 | 2 | 7 | 13 | 0 |
| 3R | 3425487 | 2 | 22 | 11 | 0 |
| 3R | 3449061 | 2 | 10 | 24 | 0 |
| 3R | 3453380 | 2 | 8 | 12 | 0 |
| 3R | 3454060 | 2 | 7 | 10 | 0 |
| 3R | 3455980 | 2 | 11 | 9 | 0 |
| 3R | 3461037 | 2 | 14 | 6 | 0 |
| 3R | 3468126 | 2 | 7 | 12 | 0 |
| 3R | 3479633 | 2 | 15 | 4 | 0 |
| 3R | 3482368 | 2 | 10 | 8 | 0 |
| 3R | 3489618 | 2 | 12 | 6 | 0 |
| 3R | 3491733 | 2 | 16 | 12 | 0 |
| 3R | 3492801 | 2 | 8 | 9 | 0 |
| 3R | 3510901 | 2 | 6 | 12 | 0 |
| 3R | 3511833 | 2 | 4 | 16 | 0 |
| 3R | 3543781 | 2 | 4 | 10 | 0 |
| 3R | 3545685 | 2 | 5 | 14 | 0 |
| 3R | 3548769 | 2 | 4 | 17 | 0 |
| 3R | 3550453 | 2 | 5 | 5 | 0 |
| 3R | 3565892 | 2 | 20 | 26 | 0 |
| 3R | 3567883 | 2 | 5 | 15 | 0 |
| 3R | 3601222 | 2 | 7 | 28 | 0 |
| 3R | 3625942 | 2 | 7 | 5 | 0 |
| 3R | 3626131 | 2 | 4 | 10 | 0 |
| 3R | 3669681 | 2 | 6 | 11 | 0 |
| 3R | 3755785 | 2 | 5 | 14 | 0 |
| 3R | 3764340 | 2 | 5 | 9 | 0 |
| 3R | 3779383 | 2 | 11 | 23 | 0 |
| 3R | 3800401 | 2 | 5 | 19 | 0 |
| 3R | 3800599 | 2 | 11 | 7 | 0 |
| 3R | 3803144 | 2 | 5 | 10 | 0 |
| 3R | 3812388 | 2 | 7 | 26 | 0 |
| 3R | 3817677 | 2 | 7 | 23 | 0 |
| 3R | 3856749 | 2 | 6 | 12 | 0 |
| 3R | 3875782 | 2 | 5 | 20 | 0 |
| 3R | 3879632 | 2 | 5 | 10 | 0 |
| 3R | 3883227 | 2 | 12 | 21 | 0 |
| 3R | 3890056 | 2 | 4 | 10 | 0 |
| 3R | 3892478 | 2 | 6 | 9 | 0 |
| 3R | 3896009 | 2 | 5 | 18 | 0 |
| 3R | 3911042 | 2 | 5 | 7 | 0 |
| 3R | 3969785 | 2 | 5 | 8 | 0 |
| 3R | 4010401 | 2 | 6 | 19 | 0 |
| 3R | 4023453 | 2 | 7 | 25 | 0 |
| 3R | 4024466 | 2 | 4 | 18 | 0 |
| 3R | 4047807 | 2 | 11 | 28 | 0 |
| 3R | 4067941 | 2 | 4 | 10 | 0 |
| 3R | 4085358 | 2 | 7 | 19 | 0 |
| 3R | 4103240 | 2 | 6 | 12 | 0 |
| 3R | 4105958 | 2 | 8 | 8 | 0 |
| 3R | 4159743 | 2 | 4 | 14 | 0 |
| 3R | 4212896 | 2 | 10 | 8 | 0 |
| 3R | 4225570 | 2 | 4 | 9 | 0 |
| 3R | 4271394 | 2 | 5 | 11 | 0 |
| 3R | 4277219 | 2 | 5 | 23 | 0 |
| 3R | 4301610 | 2 | 7 | 5 | 0 |
| 3R | 4339796 | 2 | 7 | 16 | 0 |
| 3R | 4411425 | 2 | 6 | 5 | 0 |
| 3R | 4412054 | 2 | 4 | 9 | 0 |
| 3R | 4412250 | 2 | 12 | 8 | 0 |
| 3R | 4419534 | 2 | 7 | 6 | 0 |
| 3R | 4447826 | 2 | 9 | 8 | 0 |
| 3R | 4464541 | 2 | 5 | 28 | 0 |
| 3R | 4472512 | 2 | 15 | 7 | 0 |
| 3R | 4512935 | 2 | 5 | 7 | 0 |
| 3R | 4541707 | 2 | 8 | 6 | 0 |
| 3R | 4550559 | 2 | 6 | 11 | 0 |
| 3R | 4579473 | 2 | 16 | 8 | 0 |
| 3R | 4611088 | 2 | 13 | 8 | 0 |
| 3R | 4615546 | 2 | 5 | 6 | 0 |
| 3R | 4619578 | 2 | 10 | 9 | 0 |
| 3R | 4649529 | 2 | 18 | 6 | 0 |
| 3R | 4661672 | 2 | 5 | 6 | 0 |
| 3R | 4673965 | 2 | 6 | 7 | 0 |
| 3R | 4678694 | 2 | 4 | 9 | 0 |
| 3R | 4703452 | 2 | 21 | 9 | 0 |
| 3R | 4703976 | 2 | 14 | 4 | 0 |
| 3R | 4711570 | 2 | 9 | 8 | 0 |
| 3R | 4718878 | 2 | 15 | 5 | 0 |
| 3R | 4764879 | 2 | 6 | 11 | 0 |
| 3R | 4790759 | 2 | 10 | 7 | 0 |
| 3R | 4793941 | 2 | 6 | 5 | 0 |
| 3R | 4801266 | 2 | 6 | 14 | 0 |
| 3R | 4802435 | 2 | 4 | 9 | 0 |
| 3R | 4804668 | 2 | 5 | 6 | 0 |
| 3R | 4822791 | 2 | 6 | 16 | 0 |
| 3R | 4823096 | 2 | 6 | 5 | 0 |
| 3R | 4850214 | 2 | 4 | 9 | 0 |
| 3R | 4854926 | 2 | 4 | 18 | 0 |
| 3R | 4893117 | 2 | 6 | 13 | 0 |
| 3R | 4895073 | 2 | 4 | 13 | 0 |
| 3R | 4906847 | 2 | 4 | 9 | 0 |
| 3R | 4916894 | 2 | 5 | 11 | 0 |
| 3R | 4917652 | 2 | 8 | 11 | 0 |
| 3R | 4931805 | 2 | 5 | 8 | 0 |
| 3R | 4948445 | 2 | 4 | 13 | 0 |
| 3R | 4951140 | 2 | 5 | 5 | 0 |
| 3R | 4965437 | 2 | 10 | 8 | 0 |
| 3R | 4965931 | 2 | 11 | 10 | 0 |
| 3R | 4985662 | 2 | 6 | 13 | 0 |
| 3R | 4997288 | 2 | 8 | 10 | 0 |
| 3R | 5015330 | 2 | 4 | 14 | 0 |
| 3R | 5038683 | 2 | 4 | 29 | 0 |
| 3R | 5072115 | 2 | 4 | 10 | 0 |
| 3R | 5103676 | 2 | 5 | 23 | 0 |
| 3R | 5129426 | 2 | 12 | 14 | 0 |
| 3R | 5129725 | 2 | 6 | 17 | 0 |
| 3R | 5154499 | 2 | 12 | 16 | 0 |
| 3R | 5154832 | 2 | 5 | 9 | 0 |
| 3R | 5163798 | 2 | 5 | 14 | 0 |
| 3R | 5168265 | 2 | 10 | 6 | 0 |
| 3R | 5204536 | 2 | 5 | 14 | 0 |
| 3R | 5209917 | 2 | 14 | 10 | 0 |
| 3R | 5212912 | 2 | 8 | 20 | 0 |
| 3R | 5301391 | 2 | 15 | 18 | 0 |
| 3R | 5301830 | 2 | 17 | 9 | 0 |
| 3R | 5304728 | 2 | 10 | 10 | 0 |
| 3R | 5306425 | 2 | 9 | 9 | 0 |
| 3R | 5318831 | 2 | 11 | 10 | 0 |
| 3R | 5319871 | 2 | 6 | 6 | 0 |
| 3R | 5368337 | 2 | 4 | 13 | 0 |
| 3R | 5377750 | 2 | 4 | 10 | 0 |
| 3R | 5408269 | 2 | 10 | 10 | 0 |
| 3R | 5410538 | 2 | 6 | 10 | 0 |
| 3R | 5411911 | 2 | 12 | 28 | 0 |
| 3R | 5449070 | 2 | 16 | 8 | 0 |
| 3R | 5452085 | 2 | 6 | 17 | 0 |
| 3R | 5458206 | 2 | 17 | 13 | 0 |
| 3R | 5474860 | 2 | 5 | 15 | 0 |
| 3R | 5652353 | 2 | 5 | 9 | 0 |
| 3R | 5688856 | 2 | 5 | 11 | 0 |
| 3R | 5707066 | 2 | 4 | 14 | 0 |
| 3R | 5722220 | 2 | 5 | 7 | 0 |
| 3R | 5729839 | 2 | 12 | 9 | 0 |
| 3R | 5730289 | 2 | 4 | 12 | 0 |
| 3R | 5742631 | 2 | 8 | 13 | 0 |
| 3R | 5744140 | 2 | 4 | 12 | 0 |
| 3R | 5785443 | 2 | 5 | 12 | 0 |
| 3R | 5819544 | 2 | 13 | 6 | 0 |
| 3R | 5825072 | 2 | 12 | 12 | 0 |
| 3R | 5838019 | 2 | 7 | 16 | 0 |
| 3R | 5866788 | 2 | 8 | 16 | 0 |
| 3R | 5867112 | 2 | 9 | 4 | 0 |
| 3R | 5935087 | 2 | 5 | 10 | 0 |
| 3R | 5948486 | 2 | 16 | 13 | 0 |
| 3R | 5968276 | 2 | 7 | 13 | 0 |
| 3R | 5984374 | 2 | 4 | 22 | 0 |
| 3R | 5985822 | 2 | 7 | 16 | 0 |
| 3R | 5987078 | 2 | 4 | 11 | 0 |
| 3R | 5993723 | 2 | 5 | 11 | 0 |
| 3R | 6014847 | 2 | 4 | 16 | 0 |
| 3R | 6032841 | 2 | 8 | 5 | 0 |
| 3R | 6039042 | 2 | 9 | 5 | 0 |
| 3R | 6050546 | 2 | 12 | 4 | 0 |
| 3R | 6051124 | 2 | 14 | 4 | 0 |
| 3R | 6090694 | 2 | 6 | 5 | 0 |
| 3R | 6152798 | 2 | 11 | 9 | 0 |
| 3R | 6180961 | 2 | 5 | 9 | 0 |
| 3R | 6200914 | 2 | 6 | 20 | 0 |
| 3R | 6219830 | 2 | 9 | 4 | 0 |
| 3R | 6221742 | 2 | 8 | 15 | 0 |
| 3R | 6234301 | 2 | 6 | 9 | 0 |
| 3R | 6288302 | 2 | 4 | 13 | 0 |
| 3R | 6291220 | 2 | 6 | 9 | 0 |
| 3R | 6293287 | 2 | 4 | 9 | 0 |
| 3R | 6297818 | 2 | 11 | 7 | 0 |
| 3R | 6319270 | 2 | 9 | 4 | 0 |
| 3R | 6336221 | 2 | 5 | 8 | 0 |
| 3R | 6353237 | 2 | 4 | 9 | 0 |
| 3R | 6359322 | 2 | 8 | 8 | 0 |
| 3R | 6398702 | 2 | 6 | 6 | 0 |
| 3R | 6422595 | 2 | 5 | 8 | 0 |
| 3R | 6433350 | 2 | 5 | 10 | 0 |
| 3R | 6468346 | 2 | 10 | 6 | 0 |
| 3R | 6471689 | 2 | 9 | 16 | 0 |
| 3R | 6480515 | 2 | 5 | 6 | 0 |
| 3R | 6501696 | 2 | 7 | 13 | 0 |
| 3R | 6507957 | 2 | 15 | 11 | 0 |
| 3R | 6530782 | 2 | 7 | 21 | 0 |
| 3R | 6540074 | 2 | 4 | 10 | 0 |
| 3R | 6562469 | 2 | 17 | 4 | 0 |
| 3R | 6582693 | 2 | 22 | 6 | 0 |
| 3R | 6601520 | 2 | 14 | 5 | 0 |
| 3R | 6616377 | 2 | 15 | 19 | 0 |
| 3R | 6617982 | 2 | 10 | 11 | 0 |
| 3R | 6622501 | 2 | 15 | 10 | 0 |
| 3R | 6628968 | 2 | 10 | 4 | 0 |
| 3R | 6644789 | 2 | 18 | 4 | 0 |
| 3R | 6654705 | 2 | 16 | 6 | 0 |
| 3R | 6662091 | 2 | 4 | 9 | 0 |
| 3R | 6684256 | 2 | 6 | 18 | 0 |
| 3R | 6690776 | 2 | 29 | 4 | 0 |
| 3R | 6691097 | 2 | 6 | 6 | 0 |
| 3R | 6692491 | 2 | 4 | 18 | 0 |
| 3R | 6694701 | 2 | 19 | 11 | 0 |
| 3R | 6696528 | 2 | 13 | 10 | 0 |
| 3R | 6696641 | 2 | 6 | 5 | 0 |
| 3R | 6698339 | 2 | 6 | 5 | 0 |
| 3R | 6720691 | 2 | 6 | 5 | 0 |
| 3R | 6740183 | 2 | 10 | 13 | 0 |
| 3R | 6742003 | 2 | 11 | 24 | 0 |
| 3R | 6757739 | 2 | 11 | 11 | 0 |
| 3R | 6766309 | 2 | 5 | 5 | 0 |
| 3R | 6767031 | 2 | 4 | 9 | 0 |
| 3R | 6771414 | 2 | 6 | 5 | 0 |
| 3R | 6796860 | 2 | 13 | 16 | 0 |
| 3R | 6816763 | 2 | 16 | 9 | 0 |
| 3R | 6836721 | 2 | 6 | 6 | 0 |
| 3R | 6845767 | 2 | 11 | 7 | 0 |
| 3R | 6850975 | 2 | 5 | 14 | 0 |
| 3R | 6856060 | 2 | 9 | 9 | 0 |
| 3R | 6868652 | 2 | 7 | 7 | 0 |
| 3R | 6878495 | 2 | 8 | 10 | 0 |
| 3R | 6889363 | 2 | 8 | 7 | 0 |
| 3R | 6907469 | 2 | 9 | 4 | 0 |
| 3R | 6922869 | 2 | 20 | 8 | 0 |
| 3R | 6928040 | 2 | 14 | 11 | 0 |
| 3R | 6929142 | 2 | 10 | 9 | 0 |
| 3R | 6934466 | 2 | 10 | 9 | 0 |
| 3R | 6942462 | 2 | 4 | 13 | 0 |
| 3R | 6955247 | 2 | 7 | 8 | 0 |
| 3R | 6958966 | 2 | 14 | 6 | 0 |
| 3R | 6968517 | 2 | 15 | 8 | 0 |
| 3R | 6973023 | 2 | 6 | 15 | 0 |
| 3R | 6990776 | 2 | 5 | 6 | 0 |
| 3R | 7072281 | 2 | 6 | 8 | 0 |
| 3R | 7075395 | 2 | 9 | 22 | 0 |
| 3R | 7081742 | 2 | 7 | 5 | 0 |
| 3R | 7082576 | 2 | 4 | 11 | 0 |
| 3R | 7089087 | 2 | 5 | 9 | 0 |
| 3R | 7114874 | 2 | 10 | 7 | 0 |
| 3R | 7137679 | 2 | 9 | 7 | 0 |
| 3R | 7139665 | 2 | 5 | 15 | 0 |
| 3R | 7143826 | 2 | 6 | 9 | 0 |
| 3R | 7154292 | 2 | 18 | 9 | 0 |
| 3R | 7166194 | 2 | 6 | 15 | 0 |
| 3R | 7192980 | 2 | 16 | 8 | 0 |
| 3R | 7278803 | 2 | 9 | 4 | 0 |
| 3R | 7307857 | 2 | 16 | 14 | 0 |
| 3R | 7312430 | 2 | 7 | 5 | 0 |
| 3R | 7321747 | 2 | 8 | 9 | 0 |
| 3R | 7355095 | 2 | 9 | 4 | 0 |
| 3R | 7360421 | 2 | 11 | 4 | 0 |
| 3R | 7388187 | 2 | 7 | 12 | 0 |
| 3R | 7411543 | 2 | 5 | 16 | 0 |
| 3R | 7471645 | 2 | 7 | 13 | 0 |
| 3R | 7492260 | 2 | 6 | 5 | 0 |
| 3R | 7497769 | 2 | 4 | 21 | 0 |
| 3R | 7503828 | 2 | 13 | 6 | 0 |
| 3R | 7505032 | 2 | 8 | 5 | 0 |
| 3R | 7506552 | 2 | 24 | 13 | 0 |
| 3R | 7508745 | 2 | 8 | 7 | 0 |
| 3R | 7535423 | 2 | 17 | 8 | 0 |
| 3R | 7538604 | 2 | 5 | 5 | 0 |
| 3R | 7542374 | 2 | 4 | 20 | 0 |
| 3R | 7544107 | 2 | 16 | 6 | 0 |
| 3R | 7576399 | 2 | 13 | 8 | 0 |
| 3R | 7579202 | 2 | 5 | 14 | 0 |
| 3R | 7592426 | 2 | 8 | 6 | 0 |
| 3R | 7641950 | 2 | 7 | 5 | 0 |
| 3R | 7657553 | 2 | 6 | 10 | 0 |
| 3R | 7674778 | 2 | 7 | 6 | 0 |
| 3R | 7686915 | 2 | 16 | 14 | 0 |
| 3R | 7701214 | 2 | 9 | 4 | 0 |
| 3R | 7710116 | 2 | 6 | 15 | 0 |
| 3R | 7710356 | 2 | 15 | 9 | 0 |
| 3R | 7711517 | 2 | 18 | 11 | 0 |
| 3R | 7719080 | 2 | 13 | 4 | 0 |
| 3R | 7727012 | 2 | 15 | 7 | 0 |
| 3R | 7739668 | 2 | 15 | 5 | 0 |
| 3R | 7743381 | 2 | 7 | 11 | 0 |
| 3R | 7758897 | 2 | 17 | 6 | 0 |
| 3R | 7761083 | 2 | 23 | 6 | 0 |
| 3R | 7777988 | 2 | 4 | 10 | 0 |
| 3R | 7797822 | 2 | 8 | 21 | 0 |
| 3R | 7800553 | 2 | 6 | 11 | 0 |
| 3R | 7802091 | 2 | 4 | 20 | 0 |
| 3R | 7804045 | 2 | 6 | 6 | 0 |
| 3R | 7804816 | 2 | 4 | 10 | 0 |
| 3R | 7816658 | 2 | 12 | 8 | 0 |
| 3R | 7817712 | 2 | 9 | 9 | 0 |
| 3R | 7818255 | 2 | 4 | 13 | 0 |
| 3R | 7829938 | 2 | 7 | 9 | 0 |
| 3R | 7832709 | 2 | 12 | 7 | 0 |
| 3R | 7851008 | 2 | 13 | 9 | 0 |
| 3R | 7852008 | 2 | 7 | 30 | 0 |
| 3R | 7859271 | 2 | 8 | 6 | 0 |
| 3R | 7864034 | 2 | 7 | 24 | 0 |
| 3R | 7873912 | 2 | 8 | 6 | 0 |
| 3R | 7874179 | 2 | 12 | 7 | 0 |
| 3R | 7880782 | 2 | 7 | 14 | 0 |
| 3R | 7884247 | 2 | 5 | 7 | 0 |
| 3R | 7886141 | 2 | 8 | 6 | 0 |
| 3R | 7892098 | 2 | 17 | 7 | 0 |
| 3R | 7893942 | 2 | 12 | 4 | 0 |
| 3R | 7895897 | 2 | 11 | 4 | 0 |
| 3R | 7899494 | 2 | 4 | 19 | 0 |
| 3R | 7914908 | 2 | 7 | 6 | 0 |
| 3R | 7916005 | 2 | 9 | 4 | 0 |
| 3R | 7927504 | 2 | 16 | 9 | 0 |
| 3R | 7942280 | 2 | 10 | 12 | 0 |
| 3R | 7962663 | 2 | 8 | 6 | 0 |
| 3R | 7970147 | 2 | 4 | 15 | 0 |
| 3R | 7972855 | 2 | 6 | 5 | 0 |
| 3R | 7977720 | 2 | 12 | 4 | 0 |
| 3R | 7979092 | 2 | 5 | 13 | 0 |
| 3R | 7990042 | 2 | 23 | 4 | 0 |
| 3R | 7998787 | 2 | 15 | 6 | 0 |
| 3R | 8003248 | 2 | 19 | 4 | 0 |
| 3R | 8027028 | 2 | 5 | 18 | 0 |
| 3R | 8061395 | 2 | 7 | 15 | 0 |
| 3R | 8070972 | 2 | 12 | 19 | 0 |
| 3R | 8077160 | 2 | 6 | 11 | 0 |
| 3R | 8104344 | 2 | 4 | 10 | 0 |
| 3R | 8108735 | 2 | 9 | 7 | 0 |
| 3R | 8117788 | 2 | 11 | 25 | 0 |
| 3R | 8118643 | 2 | 8 | 5 | 0 |
| 3R | 8131983 | 2 | 7 | 20 | 0 |
| 3R | 8133613 | 2 | 4 | 16 | 0 |
| 3R | 8147501 | 2 | 4 | 11 | 0 |
| 3R | 8148600 | 2 | 11 | 4 | 0 |
| 3R | 8148715 | 2 | 10 | 7 | 0 |
| 3R | 8152129 | 2 | 10 | 5 | 0 |
| 3R | 8155670 | 2 | 10 | 8 | 0 |
| 3R | 8173405 | 2 | 15 | 5 | 0 |
| 3R | 8173830 | 2 | 12 | 6 | 0 |
| 3R | 8184220 | 2 | 20 | 6 | 0 |
| 3R | 8190543 | 2 | 5 | 17 | 0 |
| 3R | 8190694 | 2 | 6 | 21 | 0 |
| 3R | 8237425 | 2 | 10 | 14 | 0 |
| 3R | 8247252 | 2 | 16 | 12 | 0 |
| 3R | 8249756 | 2 | 15 | 10 | 0 |
| 3R | 8271168 | 2 | 16 | 5 | 0 |
| 3R | 8286537 | 2 | 4 | 20 | 0 |
| 3R | 8343466 | 2 | 4 | 10 | 0 |
| 3R | 8344006 | 2 | 10 | 5 | 0 |
| 3R | 8344644 | 2 | 5 | 11 | 0 |
| 3R | 8364355 | 2 | 7 | 16 | 0 |
| 3R | 8365396 | 2 | 5 | 20 | 0 |
| 3R | 8366867 | 2 | 16 | 4 | 0 |
| 3R | 8378537 | 2 | 9 | 4 | 0 |
| 3R | 8410839 | 2 | 10 | 5 | 0 |
| 3R | 8416163 | 2 | 22 | 8 | 0 |
| 3R | 8426555 | 2 | 4 | 11 | 0 |
| 3R | 8428746 | 2 | 8 | 20 | 0 |
| 3R | 8430466 | 2 | 5 | 12 | 0 |
| 3R | 8435842 | 2 | 5 | 7 | 0 |
| 3R | 8440715 | 2 | 4 | 12 | 0 |
| 3R | 8448444 | 2 | 9 | 9 | 0 |
| 3R | 8472818 | 2 | 14 | 5 | 0 |
| 3R | 8487426 | 2 | 11 | 7 | 0 |
| 3R | 8492844 | 2 | 10 | 11 | 0 |
| 3R | 8493677 | 2 | 29 | 12 | 0 |
| 3R | 8496498 | 2 | 8 | 5 | 0 |
| 3R | 8504484 | 2 | 13 | 6 | 0 |
| 3R | 8523278 | 2 | 8 | 11 | 0 |
| 3R | 8552010 | 2 | 7 | 14 | 0 |
| 3R | 8555418 | 2 | 12 | 4 | 0 |
| 3R | 8560101 | 2 | 5 | 6 | 0 |
| 3R | 8569916 | 2 | 11 | 16 | 0 |
| 3R | 8572212 | 2 | 6 | 10 | 0 |
| 3R | 8576978 | 2 | 4 | 9 | 0 |
| 3R | 8585248 | 2 | 11 | 7 | 0 |
| 3R | 8612781 | 2 | 13 | 4 | 0 |
| 3R | 8613480 | 2 | 15 | 10 | 0 |
| 3R | 8615531 | 2 | 8 | 6 | 0 |
| 3R | 8615668 | 2 | 12 | 7 | 0 |
| 3R | 8616515 | 2 | 11 | 10 | 0 |
| 3R | 8621637 | 2 | 13 | 8 | 0 |
| 3R | 8624787 | 2 | 5 | 5 | 0 |
| 3R | 8627025 | 2 | 5 | 8 | 0 |
| 3R | 8640169 | 2 | 5 | 7 | 0 |
| 3R | 8640204 | 2 | 5 | 12 | 0 |
| 3R | 8645464 | 2 | 6 | 23 | 0 |
| 3R | 8650195 | 2 | 9 | 13 | 0 |
| 3R | 8650899 | 2 | 5 | 9 | 0 |
| 3R | 8656720 | 2 | 8 | 5 | 0 |
| 3R | 8683239 | 2 | 7 | 8 | 0 |
| 3R | 8718793 | 2 | 7 | 11 | 0 |
| 3R | 8720294 | 2 | 9 | 4 | 0 |
| 3R | 8733469 | 2 | 14 | 6 | 0 |
| 3R | 8733474 | 2 | 11 | 7 | 0 |
| 3R | 8752363 | 2 | 6 | 5 | 0 |
| 3R | 8754814 | 2 | 5 | 9 | 0 |
| 3R | 8761849 | 2 | 8 | 5 | 0 |
| 3R | 8763152 | 2 | 14 | 5 | 0 |
| 3R | 8765277 | 2 | 8 | 18 | 0 |
| 3R | 8771798 | 2 | 13 | 7 | 0 |
| 3R | 8776769 | 2 | 10 | 10 | 0 |
| 3R | 8780199 | 2 | 14 | 6 | 0 |
| 3R | 8793684 | 2 | 4 | 15 | 0 |
| 3R | 8809229 | 2 | 12 | 17 | 0 |
| 3R | 8833481 | 2 | 19 | 10 | 0 |
| 3R | 8834623 | 2 | 7 | 9 | 0 |
| 3R | 8839398 | 2 | 10 | 8 | 0 |
| 3R | 8841576 | 2 | 14 | 4 | 0 |
| 3R | 8856881 | 2 | 22 | 10 | 0 |
| 3R | 8858130 | 2 | 6 | 19 | 0 |
| 3R | 8867172 | 2 | 11 | 4 | 0 |
| 3R | 8897602 | 2 | 6 | 13 | 0 |
| 3R | 8897961 | 2 | 11 | 4 | 0 |
| 3R | 8908278 | 2 | 5 | 6 | 0 |
| 3R | 8925850 | 2 | 4 | 17 | 0 |
| 3R | 8994268 | 2 | 11 | 5 | 0 |
| 3R | 9057618 | 2 | 17 | 8 | 0 |
| 3R | 9078324 | 2 | 13 | 5 | 0 |
| 3R | 9081485 | 2 | 5 | 6 | 0 |
| 3R | 9099346 | 2 | 13 | 8 | 0 |
| 3R | 9206033 | 2 | 8 | 5 | 0 |
| 3R | 9236066 | 2 | 9 | 5 | 0 |
| 3R | 9245377 | 2 | 4 | 12 | 0 |
| 3R | 9258216 | 2 | 11 | 4 | 0 |
| 3R | 9277109 | 2 | 19 | 11 | 0 |
| 3R | 9278385 | 2 | 9 | 6 | 0 |
| 3R | 9285145 | 2 | 11 | 7 | 0 |
| 3R | 9285488 | 2 | 20 | 29 | 0 |
| 3R | 9296906 | 2 | 6 | 7 | 0 |
| 3R | 9306636 | 2 | 6 | 13 | 0 |
| 3R | 9319162 | 2 | 5 | 23 | 0 |
| 3R | 9324313 | 2 | 5 | 7 | 0 |
| 3R | 9324924 | 2 | 10 | 5 | 0 |
| 3R | 9381418 | 2 | 9 | 6 | 0 |
| 3R | 9385737 | 2 | 10 | 9 | 0 |
| 3R | 9388155 | 2 | 7 | 18 | 0 |
| 3R | 9419189 | 2 | 11 | 5 | 0 |
| 3R | 9502358 | 2 | 8 | 7 | 0 |
| 3R | 9548808 | 2 | 6 | 9 | 0 |
| 3R | 9585079 | 2 | 10 | 10 | 0 |
| 3R | 9621450 | 2 | 8 | 5 | 0 |
| 3R | 9623482 | 2 | 6 | 10 | 0 |
| 3R | 9646218 | 2 | 13 | 7 | 0 |
| 3R | 9650789 | 2 | 9 | 5 | 0 |
| 3R | 9650796 | 2 | 9 | 4 | 0 |
| 3R | 9675873 | 2 | 7 | 5 | 0 |
| 3R | 9679941 | 2 | 7 | 8 | 0 |
| 3R | 9716357 | 2 | 5 | 8 | 0 |
| 3R | 9716538 | 2 | 5 | 6 | 0 |
| 3R | 9718901 | 2 | 4 | 23 | 0 |
| 3R | 9734074 | 2 | 16 | 7 | 0 |
| 3R | 9735938 | 2 | 13 | 7 | 0 |
| 3R | 9745579 | 2 | 12 | 12 | 0 |
| 3R | 9770520 | 2 | 5 | 5 | 0 |
| 3R | 9776903 | 2 | 12 | 6 | 0 |
| 3R | 9794542 | 2 | 13 | 5 | 0 |
| 3R | 9835562 | 2 | 9 | 7 | 0 |
| 3R | 9861379 | 2 | 4 | 21 | 0 |
| 3R | 10130589 | 2 | 19 | 4 | 0 |
| 3R | 10137871 | 2 | 5 | 9 | 0 |
| 3R | 11390609 | 2 | 9 | 4 | 0 |
| 3R | 11631542 | 2 | 5 | 9 | 0 |
| 3R | 11671233 | 2 | 4 | 9 | 0 |
| 3R | 12784541 | 2 | 6 | 8 | 0 |
| 3R | 12842110 | 2 | 8 | 10 | 0 |
| 3R | 12843918 | 2 | 4 | 9 | 0 |
| 3R | 12887849 | 2 | 5 | 8 | 0 |
| 3R | 13066472 | 2 | 4 | 15 | 0 |
| 3R | 13093562 | 2 | 5 | 5 | 0 |
| 3R | 13211899 | 2 | 5 | 9 | 0 |
| 3R | 13234914 | 2 | 5 | 6 | 0 |
| 3R | 13308882 | 2 | 6 | 7 | 0 |
| 3R | 13426412 | 2 | 12 | 5 | 0 |
| 3R | 13593061 | 2 | 6 | 9 | 0 |
| 3R | 13696430 | 2 | 5 | 6 | 0 |
| 3R | 13734327 | 2 | 7 | 17 | 0 |
| 3R | 14037551 | 2 | 8 | 9 | 0 |
| 3R | 14138534 | 2 | 11 | 6 | 0 |
| 3R | 14339293 | 2 | 4 | 12 | 0 |
| 3R | 14357650 | 2 | 4 | 11 | 0 |
| 3R | 14377140 | 2 | 6 | 13 | 0 |
| 3R | 14381191 | 2 | 8 | 10 | 0 |
| 3R | 14413418 | 2 | 6 | 5 | 0 |
| 3R | 14415032 | 2 | 10 | 16 | 0 |
| 3R | 14488178 | 2 | 7 | 10 | 0 |
| 3R | 14499725 | 2 | 4 | 11 | 0 |
| 3R | 14504667 | 2 | 5 | 5 | 0 |
| 3R | 14523836 | 2 | 6 | 12 | 0 |
| 3R | 14945211 | 2 | 5 | 7 | 0 |
| 3R | 15061093 | 2 | 9 | 4 | 0 |
| 3R | 15141735 | 2 | 5 | 6 | 0 |
| 3R | 15344440 | 2 | 6 | 5 | 0 |
| 3R | 15802873 | 2 | 17 | 4 | 0 |
| 3R | 16092053 | 2 | 5 | 10 | 0 |
| 3R | 16173175 | 2 | 5 | 7 | 0 |
| 3R | 16692895 | 2 | 5 | 5 | 0 |
| 3R | 17201269 | 2 | 19 | 4 | 0 |
| 3R | 17420865 | 2 | 4 | 9 | 0 |
| 3R | 17782444 | 2 | 4 | 9 | 0 |
| 3R | 20636763 | 2 | 10 | 7 | 0 |
| 3R | 20691742 | 2 | 9 | 6 | 0 |
| 3R | 20703001 | 2 | 6 | 7 | 0 |
| 3R | 20738747 | 2 | 5 | 10 | 0 |
| 3R | 20747095 | 2 | 12 | 5 | 0 |
| 3R | 20930994 | 2 | 6 | 7 | 0 |
| 3R | 20931458 | 2 | 6 | 5 | 0 |
| 3R | 20932299 | 2 | 9 | 4 | 0 |
| 3R | 20957331 | 2 | 9 | 4 | 0 |
| 3R | 20993247 | 2 | 5 | 5 | 0 |
| 3R | 20997255 | 2 | 8 | 13 | 0 |
| 3R | 21013830 | 2 | 9 | 4 | 0 |
| 3R | 21021456 | 2 | 6 | 7 | 0 |
| 3R | 21034741 | 2 | 5 | 7 | 0 |
| 3R | 21035237 | 2 | 10 | 6 | 0 |
| 3R | 21040529 | 2 | 9 | 5 | 0 |
| 3R | 21045802 | 2 | 13 | 6 | 0 |
| 3R | 21072864 | 2 | 8 | 5 | 0 |
| 3R | 21094509 | 2 | 9 | 4 | 0 |
| 3R | 21096564 | 2 | 6 | 10 | 0 |
| 3R | 21097207 | 2 | 4 | 9 | 0 |
| 3R | 21118407 | 2 | 7 | 6 | 0 |
| 3R | 21122720 | 2 | 10 | 4 | 0 |
| 3R | 21127478 | 2 | 7 | 10 | 0 |
| 3R | 21127568 | 2 | 12 | 5 | 0 |
| 3R | 21131466 | 2 | 11 | 5 | 0 |
| 3R | 21278955 | 2 | 6 | 8 | 0 |
| 3R | 21354324 | 2 | 8 | 8 | 0 |
| 3R | 21373852 | 2 | 5 | 5 | 0 |
| 3R | 21429727 | 2 | 13 | 4 | 0 |
| 3R | 21453004 | 2 | 12 | 4 | 0 |
| 3R | 21455946 | 2 | 13 | 7 | 0 |
| 3R | 21527432 | 2 | 12 | 8 | 0 |
| 3R | 21567789 | 2 | 6 | 5 | 0 |
| 3R | 21567827 | 2 | 9 | 7 | 0 |
| 3R | 21643061 | 2 | 8 | 5 | 0 |
| 3R | 21680890 | 2 | 9 | 15 | 0 |
| 3R | 21724036 | 2 | 5 | 6 | 0 |
| 3R | 21740609 | 2 | 6 | 6 | 0 |
| 3R | 21747063 | 2 | 7 | 5 | 0 |
| 3R | 21910389 | 2 | 5 | 15 | 0 |
| 3R | 21925737 | 2 | 5 | 11 | 0 |
| 3R | 21977976 | 2 | 7 | 6 | 0 |
| 3R | 21995673 | 2 | 5 | 7 | 0 |
| 3R | 22017318 | 2 | 8 | 8 | 0 |
| 3R | 22026020 | 2 | 15 | 9 | 0 |
| 3R | 22037414 | 2 | 5 | 10 | 0 |
| 3R | 22091088 | 2 | 6 | 6 | 0 |
| 3R | 22157975 | 2 | 4 | 9 | 0 |
| 3R | 22175880 | 2 | 6 | 5 | 0 |
| 3R | 22187957 | 2 | 9 | 7 | 0 |
| 3R | 22759044 | 2 | 9 | 5 | 0 |
| 3R | 22899508 | 2 | 9 | 9 | 0 |
| 3R | 22900563 | 2 | 5 | 7 | 0 |
| 3R | 22911498 | 2 | 14 | 6 | 0 |
| 3R | 22927276 | 2 | 16 | 5 | 0 |
| 3R | 22959922 | 2 | 7 | 7 | 0 |
| 3R | 23230447 | 2 | 7 | 6 | 0 |
| 3R | 23299380 | 2 | 6 | 10 | 0 |
| 3R | 23425352 | 2 | 5 | 7 | 0 |
| 3R | 23459094 | 2 | 6 | 7 | 0 |
| 3R | 23463972 | 2 | 9 | 6 | 0 |
| 3R | 23490478 | 2 | 17 | 4 | 0 |
| 3R | 23505408 | 2 | 4 | 9 | 0 |
| 3R | 23608768 | 2 | 11 | 6 | 0 |
| 3R | 23624647 | 2 | 5 | 8 | 0 |
| 3R | 24116080 | 2 | 7 | 8 | 0 |
| 3R | 24438982 | 2 | 5 | 5 | 0 |
| 3R | 24580256 | 2 | 5 | 7 | 0 |
| 3R | 24681117 | 2 | 7 | 7 | 0 |
| 3R | 24845764 | 2 | 8 | 8 | 0 |
| X | 36653 | 2 | 6 | 19 | 0 |
| X | 79749 | 2 | 4 | 22 | 0 |
| X | 88322 | 2 | 9 | 6 | 0 |
| X | 97518 | 2 | 12 | 4 | 0 |
| X | 142650 | 2 | 16 | 12 | 0 |
| X | 146867 | 2 | 4 | 10 | 0 |
| X | 153547 | 2 | 5 | 15 | 0 |
| X | 156117 | 2 | 12 | 5 | 0 |
| X | 178527 | 2 | 11 | 13 | 0 |
| X | 179450 | 2 | 9 | 16 | 0 |
| X | 202395 | 2 | 14 | 21 | 0 |
| X | 203434 | 2 | 14 | 4 | 0 |
| X | 206104 | 2 | 5 | 18 | 0 |
| X | 332653 | 2 | 13 | 13 | 0 |
| X | 344142 | 2 | 5 | 6 | 0 |
| X | 367036 | 2 | 8 | 16 | 0 |
| X | 385294 | 2 | 12 | 13 | 0 |
| X | 391919 | 2 | 12 | 6 | 0 |
| X | 483177 | 2 | 6 | 6 | 0 |
| X | 487662 | 2 | 6 | 24 | 0 |
| X | 497489 | 2 | 9 | 8 | 0 |
| X | 498560 | 2 | 13 | 7 | 0 |
| X | 501330 | 2 | 24 | 13 | 0 |
| X | 502553 | 2 | 4 | 12 | 0 |
| X | 503864 | 2 | 24 | 4 | 0 |
| X | 504181 | 2 | 10 | 6 | 0 |
| X | 507079 | 2 | 21 | 10 | 0 |
| X | 511485 | 2 | 31 | 13 | 0 |
| X | 511617 | 2 | 10 | 15 | 0 |
| X | 514945 | 2 | 21 | 5 | 0 |
| X | 515344 | 2 | 7 | 6 | 0 |
| X | 518765 | 2 | 25 | 8 | 0 |
| X | 518884 | 2 | 22 | 13 | 0 |
| X | 521738 | 2 | 10 | 4 | 0 |
| X | 521808 | 2 | 25 | 16 | 0 |
| X | 522066 | 2 | 11 | 11 | 0 |
| X | 522240 | 2 | 10 | 5 | 0 |
| X | 524614 | 2 | 12 | 11 | 0 |
| X | 524854 | 2 | 37 | 14 | 0 |
| X | 525044 | 2 | 21 | 11 | 0 |
| X | 525143 | 2 | 26 | 22 | 0 |
| X | 527808 | 2 | 12 | 7 | 0 |
| X | 528142 | 2 | 21 | 16 | 0 |
| X | 528657 | 2 | 18 | 7 | 0 |
| X | 534697 | 2 | 15 | 4 | 0 |
| X | 537980 | 2 | 4 | 14 | 0 |
| X | 556614 | 2 | 18 | 14 | 0 |
| X | 558781 | 2 | 9 | 25 | 0 |
| X | 572750 | 2 | 19 | 11 | 0 |
| X | 576737 | 2 | 4 | 11 | 0 |
| X | 578546 | 2 | 20 | 9 | 0 |
| X | 590030 | 2 | 5 | 18 | 0 |
| X | 590476 | 2 | 9 | 14 | 0 |
| X | 601434 | 2 | 6 | 10 | 0 |
| X | 601944 | 2 | 11 | 10 | 0 |
| X | 602144 | 2 | 23 | 4 | 0 |
| X | 602341 | 2 | 25 | 6 | 0 |
| X | 602787 | 2 | 18 | 15 | 0 |
| X | 610972 | 2 | 21 | 10 | 0 |
| X | 614767 | 2 | 15 | 8 | 0 |
| X | 615697 | 2 | 11 | 7 | 0 |
| X | 617264 | 2 | 15 | 11 | 0 |
| X | 619862 | 2 | 6 | 9 | 0 |
| X | 628150 | 2 | 19 | 23 | 0 |
| X | 633458 | 2 | 18 | 17 | 0 |
| X | 638205 | 2 | 24 | 29 | 0 |
| X | 642755 | 2 | 32 | 20 | 0 |
| X | 660427 | 2 | 11 | 7 | 0 |
| X | 662012 | 2 | 25 | 9 | 0 |
| X | 665209 | 2 | 21 | 13 | 0 |
| X | 668614 | 2 | 22 | 14 | 0 |
| X | 668855 | 2 | 10 | 5 | 0 |
| X | 669238 | 2 | 21 | 8 | 0 |
| X | 670097 | 2 | 20 | 6 | 0 |
| X | 670263 | 2 | 17 | 4 | 0 |
| X | 674368 | 2 | 15 | 7 | 0 |
| X | 679535 | 2 | 27 | 13 | 0 |
| X | 686968 | 2 | 22 | 19 | 0 |
| X | 687475 | 2 | 4 | 22 | 0 |
| X | 689156 | 2 | 18 | 4 | 0 |
| X | 691879 | 2 | 17 | 19 | 0 |
| X | 692256 | 2 | 4 | 22 | 0 |
| X | 695073 | 2 | 13 | 6 | 0 |
| X | 701837 | 2 | 9 | 24 | 0 |
| X | 715838 | 2 | 23 | 15 | 0 |
| X | 717437 | 2 | 23 | 9 | 0 |
| X | 718532 | 2 | 14 | 15 | 0 |
| X | 718644 | 2 | 8 | 17 | 0 |
| X | 731561 | 2 | 24 | 15 | 0 |
| X | 731621 | 2 | 24 | 24 | 0 |
| X | 731735 | 2 | 29 | 15 | 0 |
| X | 736751 | 2 | 14 | 11 | 0 |
| X | 749036 | 2 | 11 | 4 | 0 |
| X | 750996 | 2 | 18 | 5 | 0 |
| X | 751068 | 2 | 27 | 6 | 0 |
| X | 753939 | 2 | 19 | 11 | 0 |
| X | 754798 | 2 | 14 | 10 | 0 |
| X | 756219 | 2 | 12 | 12 | 0 |
| X | 756716 | 2 | 9 | 13 | 0 |
| X | 757844 | 2 | 8 | 8 | 0 |
| X | 765233 | 2 | 22 | 7 | 0 |
| X | 767277 | 2 | 22 | 5 | 0 |
| X | 767894 | 2 | 13 | 10 | 0 |
| X | 767937 | 2 | 27 | 5 | 0 |
| X | 767986 | 2 | 10 | 5 | 0 |
| X | 768222 | 2 | 5 | 22 | 0 |
| X | 769576 | 2 | 18 | 5 | 0 |
| X | 770229 | 2 | 23 | 9 | 0 |
| X | 770445 | 2 | 7 | 13 | 0 |
| X | 770532 | 2 | 27 | 23 | 0 |
| X | 771947 | 2 | 15 | 12 | 0 |
| X | 779799 | 2 | 13 | 9 | 0 |
| X | 781273 | 2 | 23 | 7 | 0 |
| X | 785807 | 2 | 12 | 23 | 0 |
| X | 786005 | 2 | 8 | 10 | 0 |
| X | 786522 | 2 | 17 | 20 | 0 |
| X | 789962 | 2 | 13 | 5 | 0 |
| X | 790401 | 2 | 27 | 20 | 0 |
| X | 791219 | 2 | 11 | 13 | 0 |
| X | 794372 | 2 | 10 | 8 | 0 |
| X | 796305 | 2 | 11 | 5 | 0 |
| X | 796633 | 2 | 28 | 12 | 0 |
| X | 803810 | 2 | 15 | 14 | 0 |
| X | 806724 | 2 | 25 | 7 | 0 |
| X | 815515 | 2 | 8 | 6 | 0 |
| X | 817883 | 2 | 10 | 6 | 0 |
| X | 821204 | 2 | 4 | 9 | 0 |
| X | 821989 | 2 | 14 | 6 | 0 |
| X | 822011 | 2 | 17 | 10 | 0 |
| X | 822064 | 2 | 12 | 6 | 0 |
| X | 823788 | 2 | 8 | 17 | 0 |
| X | 824785 | 2 | 13 | 6 | 0 |
| X | 826913 | 2 | 27 | 20 | 0 |
| X | 831857 | 2 | 9 | 18 | 0 |
| X | 843377 | 2 | 12 | 19 | 0 |
| X | 849369 | 2 | 14 | 19 | 0 |
| X | 857897 | 2 | 4 | 16 | 0 |
| X | 859623 | 2 | 10 | 21 | 0 |
| X | 860468 | 2 | 7 | 14 | 0 |
| X | 868078 | 2 | 6 | 24 | 0 |
| X | 874281 | 2 | 14 | 8 | 0 |
| X | 876541 | 2 | 6 | 8 | 0 |
| X | 876645 | 2 | 4 | 19 | 0 |
| X | 876785 | 2 | 4 | 12 | 0 |
| X | 877238 | 2 | 4 | 12 | 0 |
| X | 883644 | 2 | 6 | 8 | 0 |
| X | 889209 | 2 | 5 | 15 | 0 |
| X | 890845 | 2 | 9 | 7 | 0 |
| X | 892612 | 2 | 12 | 5 | 0 |
| X | 893806 | 2 | 16 | 5 | 0 |
| X | 895951 | 2 | 28 | 4 | 0 |
| X | 897788 | 2 | 5 | 17 | 0 |
| X | 902523 | 2 | 6 | 12 | 0 |
| X | 902792 | 2 | 4 | 17 | 0 |
| X | 902913 | 2 | 5 | 16 | 0 |
| X | 903038 | 2 | 4 | 16 | 0 |
| X | 904081 | 2 | 11 | 15 | 0 |
| X | 906824 | 2 | 10 | 21 | 0 |
| X | 907331 | 2 | 6 | 13 | 0 |
| X | 908189 | 2 | 7 | 13 | 0 |
| X | 909249 | 2 | 10 | 16 | 0 |
| X | 914893 | 2 | 3 | 34 | 0 |
| X | 916150 | 2 | 4 | 13 | 0 |
| X | 917886 | 2 | 10 | 13 | 0 |
| X | 919811 | 2 | 9 | 20 | 0 |
| X | 922020 | 2 | 7 | 19 | 0 |
| X | 924391 | 2 | 10 | 19 | 0 |
| X | 940904 | 2 | 7 | 21 | 0 |
| X | 941409 | 2 | 9 | 13 | 0 |
| X | 946142 | 2 | 6 | 12 | 0 |
| X | 947207 | 2 | 20 | 13 | 0 |
| X | 952443 | 2 | 5 | 14 | 0 |
| X | 954751 | 2 | 10 | 11 | 0 |
| X | 968950 | 2 | 12 | 19 | 0 |
| X | 974492 | 2 | 6 | 28 | 0 |
| X | 980364 | 2 | 16 | 22 | 0 |
| X | 990109 | 2 | 12 | 18 | 0 |
| X | 992260 | 2 | 17 | 19 | 0 |
| X | 994785 | 2 | 7 | 10 | 0 |
| X | 995094 | 2 | 5 | 14 | 0 |
| X | 996027 | 2 | 9 | 14 | 0 |
| X | 996289 | 2 | 9 | 19 | 0 |
| X | 996535 | 2 | 7 | 11 | 0 |
| X | 996945 | 2 | 10 | 11 | 0 |
| X | 997222 | 2 | 13 | 24 | 0 |
| X | 997664 | 2 | 11 | 23 | 0 |
| X | 997741 | 2 | 13 | 22 | 0 |
| X | 997840 | 2 | 15 | 23 | 0 |
| X | 998802 | 2 | 9 | 19 | 0 |
| X | 1003445 | 2 | 6 | 8 | 0 |
| X | 1003539 | 2 | 18 | 12 | 0 |
| X | 1007067 | 2 | 6 | 18 | 0 |
| X | 1021129 | 2 | 14 | 4 | 0 |
| X | 1021393 | 2 | 9 | 10 | 0 |
| X | 1021437 | 2 | 6 | 14 | 0 |
| X | 1035573 | 2 | 14 | 10 | 0 |
| X | 1045669 | 2 | 4 | 10 | 0 |
| X | 1055445 | 2 | 24 | 22 | 0 |
| X | 1062336 | 2 | 12 | 26 | 0 |
| X | 1088117 | 2 | 8 | 17 | 0 |
| X | 1094788 | 2 | 6 | 9 | 0 |
| X | 1096849 | 2 | 5 | 13 | 0 |
| X | 1097844 | 2 | 11 | 18 | 0 |
| X | 1099851 | 2 | 4 | 13 | 0 |
| X | 1100250 | 2 | 8 | 26 | 0 |
| X | 1104881 | 2 | 5 | 5 | 0 |
| X | 1116694 | 2 | 8 | 19 | 0 |
| X | 1118921 | 2 | 4 | 10 | 0 |
| X | 1120421 | 2 | 9 | 17 | 0 |
| X | 1123744 | 2 | 8 | 17 | 0 |
| X | 1140734 | 2 | 12 | 12 | 0 |
| X | 1144193 | 2 | 7 | 18 | 0 |
| X | 1144350 | 2 | 14 | 9 | 0 |
| X | 1146443 | 2 | 7 | 8 | 0 |
| X | 1147150 | 2 | 5 | 21 | 0 |
| X | 1149009 | 2 | 8 | 10 | 0 |
| X | 1149057 | 2 | 14 | 23 | 0 |
| X | 1150139 | 2 | 7 | 10 | 0 |
| X | 1150587 | 2 | 11 | 21 | 0 |
| X | 1150974 | 2 | 15 | 21 | 0 |
| X | 1155922 | 2 | 11 | 24 | 0 |
| X | 1157570 | 2 | 4 | 22 | 0 |
| X | 1157661 | 2 | 5 | 16 | 0 |
| X | 1157931 | 2 | 7 | 9 | 0 |
| X | 1159617 | 2 | 27 | 25 | 0 |
| X | 1160748 | 2 | 5 | 10 | 0 |
| X | 1160943 | 2 | 15 | 22 | 0 |
| X | 1161079 | 2 | 11 | 11 | 0 |
| X | 1161739 | 2 | 9 | 16 | 0 |
| X | 1164140 | 2 | 10 | 8 | 0 |
| X | 1168902 | 2 | 8 | 12 | 0 |
| X | 1170233 | 2 | 8 | 21 | 0 |
| X | 1174278 | 2 | 8 | 10 | 0 |
| X | 1177488 | 2 | 13 | 6 | 0 |
| X | 1183534 | 2 | 9 | 15 | 0 |
| X | 1189782 | 2 | 17 | 17 | 0 |
| X | 1189916 | 2 | 11 | 41 | 0 |
| X | 1189931 | 2 | 5 | 40 | 0 |
| X | 1192950 | 2 | 5 | 10 | 0 |
| X | 1199582 | 2 | 5 | 8 | 0 |
| X | 1207126 | 2 | 5 | 11 | 0 |
| X | 1213405 | 2 | 7 | 15 | 0 |
| X | 1218809 | 2 | 16 | 10 | 0 |
| X | 1221603 | 2 | 18 | 25 | 0 |
| X | 1230196 | 2 | 5 | 21 | 0 |
| X | 1237093 | 2 | 11 | 7 | 0 |
| X | 1244056 | 2 | 14 | 22 | 0 |
| X | 1272269 | 2 | 13 | 14 | 0 |
| X | 1286120 | 2 | 14 | 18 | 0 |
| X | 1293664 | 2 | 14 | 32 | 0 |
| X | 1304064 | 2 | 22 | 21 | 0 |
| X | 1307427 | 2 | 4 | 16 | 0 |
| X | 1309548 | 2 | 6 | 24 | 0 |
| X | 1309878 | 2 | 7 | 12 | 0 |
| X | 1311119 | 2 | 7 | 20 | 0 |
| X | 1312438 | 2 | 13 | 20 | 0 |
| X | 1312645 | 2 | 11 | 34 | 0 |
| X | 1315496 | 2 | 14 | 13 | 0 |
| X | 1319760 | 2 | 4 | 15 | 0 |
| X | 1322591 | 2 | 7 | 15 | 0 |
| X | 1324272 | 2 | 9 | 13 | 0 |
| X | 1326715 | 2 | 15 | 28 | 0 |
| X | 1327240 | 2 | 8 | 13 | 0 |
| X | 1328062 | 2 | 5 | 20 | 0 |
| X | 1328659 | 2 | 5 | 11 | 0 |
| X | 1328825 | 2 | 4 | 12 | 0 |
| X | 1333940 | 2 | 8 | 27 | 0 |
| X | 1335090 | 2 | 11 | 19 | 0 |
| X | 1335212 | 2 | 8 | 19 | 0 |
| X | 1335668 | 2 | 19 | 26 | 0 |
| X | 1336024 | 2 | 8 | 24 | 0 |
| X | 1336774 | 2 | 5 | 23 | 0 |
| X | 1336906 | 2 | 10 | 15 | 0 |
| X | 1338640 | 2 | 14 | 19 | 0 |
| X | 1338741 | 2 | 16 | 22 | 0 |
| X | 1339495 | 2 | 8 | 16 | 0 |
| X | 1339745 | 2 | 7 | 22 | 0 |
| X | 1339842 | 2 | 6 | 31 | 0 |
| X | 1340293 | 2 | 7 | 26 | 0 |
| X | 1340403 | 2 | 17 | 25 | 0 |
| X | 1340492 | 2 | 4 | 10 | 0 |
| X | 1340961 | 2 | 8 | 26 | 0 |
| X | 1340982 | 2 | 4 | 22 | 0 |
| X | 1341184 | 2 | 6 | 12 | 0 |
| X | 1341280 | 2 | 7 | 24 | 0 |
| X | 1341365 | 2 | 8 | 27 | 0 |
| X | 1341408 | 2 | 8 | 29 | 0 |
| X | 1341783 | 2 | 7 | 9 | 0 |
| X | 1341924 | 2 | 6 | 29 | 0 |
| X | 1342110 | 2 | 7 | 19 | 0 |
| X | 1342242 | 2 | 6 | 5 | 0 |
| X | 1342401 | 2 | 21 | 31 | 0 |
| X | 1342680 | 2 | 7 | 12 | 0 |
| X | 1342998 | 2 | 5 | 14 | 0 |
| X | 1343094 | 2 | 19 | 32 | 0 |
| X | 1343129 | 2 | 5 | 15 | 0 |
| X | 1343964 | 2 | 8 | 13 | 0 |
| X | 1344159 | 2 | 9 | 19 | 0 |
| X | 1344700 | 2 | 22 | 23 | 0 |
| X | 1344930 | 2 | 5 | 8 | 0 |
| X | 1345180 | 2 | 6 | 17 | 0 |
| X | 1346321 | 2 | 6 | 25 | 0 |
| X | 1346413 | 2 | 12 | 27 | 0 |
| X | 1346456 | 2 | 4 | 33 | 0 |
| X | 1346885 | 2 | 5 | 19 | 0 |
| X | 1347421 | 2 | 20 | 30 | 0 |
| X | 1347818 | 2 | 18 | 29 | 0 |
| X | 1349212 | 2 | 8 | 28 | 0 |
| X | 1349744 | 2 | 8 | 15 | 0 |
| X | 1349908 | 2 | 6 | 14 | 0 |
| X | 1350182 | 2 | 18 | 15 | 0 |
| X | 1351392 | 2 | 8 | 9 | 0 |
| X | 1351840 | 2 | 15 | 19 | 0 |
| X | 1353689 | 2 | 5 | 9 | 0 |
| X | 1355476 | 2 | 6 | 11 | 0 |
| X | 1356060 | 2 | 10 | 19 | 0 |
| X | 1360219 | 2 | 7 | 7 | 0 |
| X | 1360531 | 2 | 4 | 19 | 0 |
| X | 1364024 | 2 | 6 | 13 | 0 |
| X | 1364400 | 2 | 13 | 17 | 0 |
| X | 1368789 | 2 | 13 | 33 | 0 |
| X | 1370565 | 2 | 5 | 20 | 0 |
| X | 1370587 | 2 | 10 | 11 | 0 |
| X | 1370961 | 2 | 25 | 30 | 0 |
| X | 1371928 | 2 | 7 | 12 | 0 |
| X | 1372438 | 2 | 9 | 14 | 0 |
| X | 1372561 | 2 | 6 | 8 | 0 |
| X | 1372786 | 2 | 4 | 19 | 0 |
| X | 1373115 | 2 | 16 | 13 | 0 |
| X | 1374353 | 2 | 6 | 27 | 0 |
| X | 1374619 | 2 | 8 | 20 | 0 |
| X | 1375041 | 2 | 8 | 26 | 0 |
| X | 1375736 | 2 | 10 | 19 | 0 |
| X | 1389603 | 2 | 9 | 20 | 0 |
| X | 1390094 | 2 | 19 | 11 | 0 |
| X | 1392458 | 2 | 22 | 13 | 0 |
| X | 1393436 | 2 | 7 | 15 | 0 |
| X | 1400766 | 2 | 11 | 10 | 0 |
| X | 1401555 | 2 | 15 | 29 | 0 |
| X | 1406136 | 2 | 14 | 11 | 0 |
| X | 1408739 | 2 | 10 | 9 | 0 |
| X | 1424185 | 2 | 11 | 5 | 0 |
| X | 1425307 | 2 | 6 | 9 | 0 |
| X | 1425587 | 2 | 4 | 10 | 0 |
| X | 1425753 | 2 | 6 | 12 | 0 |
| X | 1426143 | 2 | 7 | 10 | 0 |
| X | 1426512 | 2 | 8 | 21 | 0 |
| X | 1426584 | 2 | 15 | 13 | 0 |
| X | 1431105 | 2 | 5 | 11 | 0 |
| X | 1431149 | 2 | 14 | 14 | 0 |
| X | 1431372 | 2 | 13 | 15 | 0 |
| X | 1431606 | 2 | 4 | 11 | 0 |
| X | 1431814 | 2 | 7 | 13 | 0 |
| X | 1431872 | 2 | 8 | 18 | 0 |
| X | 1431949 | 2 | 6 | 12 | 0 |
| X | 1432079 | 2 | 8 | 27 | 0 |
| X | 1444345 | 2 | 5 | 23 | 0 |
| X | 1455956 | 2 | 10 | 18 | 0 |
| X | 1458016 | 2 | 6 | 8 | 0 |
| X | 1479688 | 2 | 5 | 8 | 0 |
| X | 1486421 | 2 | 11 | 22 | 0 |
| X | 1488896 | 2 | 5 | 8 | 0 |
| X | 1499799 | 2 | 5 | 15 | 0 |
| X | 1504223 | 2 | 8 | 17 | 0 |
| X | 1508260 | 2 | 12 | 21 | 0 |
| X | 1511351 | 2 | 17 | 8 | 0 |
| X | 1521136 | 2 | 14 | 12 | 0 |
| X | 1521549 | 2 | 5 | 20 | 0 |
| X | 1529743 | 2 | 17 | 8 | 0 |
| X | 1529853 | 2 | 18 | 15 | 0 |
| X | 1531066 | 2 | 4 | 14 | 0 |
| X | 1533000 | 2 | 5 | 23 | 0 |
| X | 1533930 | 2 | 15 | 21 | 0 |
| X | 1537854 | 2 | 18 | 8 | 0 |
| X | 1543187 | 2 | 16 | 19 | 0 |
| X | 1544072 | 2 | 8 | 19 | 0 |
| X | 1545092 | 2 | 4 | 16 | 0 |
| X | 1546771 | 2 | 9 | 24 | 0 |
| X | 1547544 | 2 | 6 | 16 | 0 |
| X | 1547689 | 2 | 9 | 19 | 0 |
| X | 1548758 | 2 | 13 | 16 | 0 |
| X | 1549194 | 2 | 4 | 22 | 0 |
| X | 1549829 | 2 | 15 | 40 | 0 |
| X | 1549978 | 2 | 17 | 29 | 0 |
| X | 1550485 | 2 | 16 | 24 | 0 |
| X | 1550957 | 2 | 11 | 31 | 0 |
| X | 1551083 | 2 | 19 | 16 | 0 |
| X | 1551806 | 2 | 4 | 15 | 0 |
| X | 1552208 | 2 | 6 | 9 | 0 |
| X | 1552327 | 2 | 4 | 25 | 0 |
| X | 1554424 | 2 | 7 | 12 | 0 |
| X | 1558589 | 2 | 5 | 14 | 0 |
| X | 1559296 | 2 | 11 | 16 | 0 |
| X | 1559637 | 2 | 14 | 20 | 0 |
| X | 1559778 | 2 | 6 | 14 | 0 |
| X | 1560105 | 2 | 4 | 20 | 0 |
| X | 1560144 | 2 | 9 | 19 | 0 |
| X | 1561515 | 2 | 15 | 17 | 0 |
| X | 1561689 | 2 | 8 | 28 | 0 |
| X | 1563404 | 2 | 4 | 13 | 0 |
| X | 1563732 | 2 | 4 | 21 | 0 |
| X | 1563960 | 2 | 11 | 11 | 0 |
| X | 1564139 | 2 | 12 | 34 | 0 |
| X | 1568778 | 2 | 4 | 17 | 0 |
| X | 1569108 | 2 | 10 | 22 | 0 |
| X | 1571189 | 2 | 15 | 22 | 0 |
| X | 1571819 | 2 | 5 | 29 | 0 |
| X | 1572034 | 2 | 23 | 30 | 0 |
| X | 1572263 | 2 | 30 | 15 | 0 |
| X | 1572334 | 2 | 18 | 35 | 0 |
| X | 1572511 | 2 | 10 | 37 | 0 |
| X | 1572729 | 2 | 16 | 13 | 0 |
| X | 1576976 | 2 | 22 | 12 | 0 |
| X | 1580147 | 2 | 4 | 10 | 0 |
| X | 1581103 | 2 | 18 | 13 | 0 |
| X | 1585377 | 2 | 4 | 14 | 0 |
| X | 1586666 | 2 | 12 | 9 | 0 |
| X | 1592905 | 2 | 12 | 24 | 0 |
| X | 1593480 | 2 | 16 | 4 | 0 |
| X | 1598947 | 2 | 4 | 19 | 0 |
| X | 1599106 | 2 | 10 | 19 | 0 |
| X | 1599204 | 2 | 13 | 18 | 0 |
| X | 1601834 | 2 | 22 | 15 | 0 |
| X | 1601922 | 2 | 17 | 5 | 0 |
| X | 1602061 | 2 | 30 | 18 | 0 |
| X | 1602179 | 2 | 33 | 3 | 0 |
| X | 1602431 | 2 | 36 | 11 | 0 |
| X | 1602692 | 2 | 9 | 6 | 0 |
| X | 1602722 | 2 | 8 | 14 | 0 |
| X | 1602925 | 2 | 21 | 10 | 0 |
| X | 1604038 | 2 | 19 | 4 | 0 |
| X | 1604389 | 2 | 43 | 14 | 0 |
| X | 1604596 | 2 | 18 | 10 | 0 |
| X | 1604763 | 2 | 23 | 11 | 0 |
| X | 1604799 | 2 | 20 | 4 | 0 |
| X | 1605172 | 2 | 4 | 9 | 0 |
| X | 1606782 | 2 | 21 | 5 | 0 |
| X | 1607026 | 2 | 11 | 4 | 0 |
| X | 1607420 | 2 | 7 | 15 | 0 |
| X | 1607968 | 2 | 15 | 12 | 0 |
| X | 1609336 | 2 | 29 | 5 | 0 |
| X | 1612105 | 2 | 18 | 9 | 0 |
| X | 1616575 | 2 | 4 | 25 | 0 |
| X | 1616629 | 2 | 9 | 15 | 0 |
| X | 1617096 | 2 | 7 | 23 | 0 |
| X | 1618906 | 2 | 21 | 7 | 0 |
| X | 1619121 | 2 | 20 | 10 | 0 |
| X | 1629611 | 2 | 11 | 18 | 0 |
| X | 1630293 | 2 | 5 | 15 | 0 |
| X | 1630737 | 2 | 11 | 20 | 0 |
| X | 1640508 | 2 | 17 | 18 | 0 |
| X | 1640679 | 2 | 4 | 29 | 0 |
| X | 1642238 | 2 | 25 | 7 | 0 |
| X | 1643588 | 2 | 8 | 8 | 0 |
| X | 1644438 | 2 | 18 | 6 | 0 |
| X | 1644751 | 2 | 28 | 19 | 0 |
| X | 1646144 | 2 | 12 | 5 | 0 |
| X | 1646614 | 2 | 6 | 16 | 0 |
| X | 1647021 | 2 | 14 | 21 | 0 |
| X | 1647080 | 2 | 5 | 19 | 0 |
| X | 1647169 | 2 | 10 | 19 | 0 |
| X | 1647692 | 2 | 3 | 36 | 0 |
| X | 1647759 | 2 | 3 | 31 | 0 |
| X | 1647793 | 2 | 13 | 20 | 0 |
| X | 1647908 | 2 | 15 | 21 | 0 |
| X | 1647996 | 2 | 16 | 22 | 0 |
| X | 1648495 | 2 | 15 | 13 | 0 |
| X | 1648775 | 2 | 4 | 13 | 0 |
| X | 1649183 | 2 | 7 | 18 | 0 |
| X | 1649358 | 2 | 7 | 17 | 0 |
| X | 1649822 | 2 | 4 | 19 | 0 |
| X | 1649942 | 2 | 8 | 21 | 0 |
| X | 1650148 | 2 | 17 | 19 | 0 |
| X | 1650966 | 2 | 7 | 12 | 0 |
| X | 1651135 | 2 | 8 | 7 | 0 |
| X | 1652110 | 2 | 7 | 5 | 0 |
| X | 1652158 | 2 | 7 | 13 | 0 |
| X | 1654280 | 2 | 6 | 15 | 0 |
| X | 1655087 | 2 | 4 | 20 | 0 |
| X | 1656229 | 2 | 4 | 9 | 0 |
| X | 1656519 | 2 | 9 | 19 | 0 |
| X | 1656567 | 2 | 8 | 13 | 0 |
| X | 1664344 | 2 | 8 | 11 | 0 |
| X | 1673049 | 2 | 11 | 25 | 0 |
| X | 1701242 | 2 | 14 | 13 | 0 |
| X | 1711898 | 2 | 18 | 11 | 0 |
| X | 1714032 | 2 | 7 | 24 | 0 |
| X | 1714210 | 2 | 19 | 36 | 0 |
| X | 1714271 | 2 | 4 | 16 | 0 |
| X | 1714619 | 2 | 14 | 21 | 0 |
| X | 1715478 | 2 | 12 | 29 | 0 |
| X | 1715591 | 2 | 7 | 22 | 0 |
| X | 1715731 | 2 | 5 | 11 | 0 |
| X | 1716286 | 2 | 10 | 21 | 0 |
| X | 1716603 | 2 | 18 | 32 | 0 |
| X | 1716696 | 2 | 15 | 25 | 0 |
| X | 1716876 | 2 | 5 | 16 | 0 |
| X | 1717680 | 2 | 4 | 18 | 0 |
| X | 1717718 | 2 | 6 | 16 | 0 |
| X | 1718697 | 2 | 10 | 16 | 0 |
| X | 1718741 | 2 | 9 | 30 | 0 |
| X | 1721209 | 2 | 27 | 15 | 0 |
| X | 1724122 | 2 | 15 | 21 | 0 |
| X | 1724811 | 2 | 9 | 4 | 0 |
| X | 1724853 | 2 | 10 | 8 | 0 |
| X | 1727275 | 2 | 14 | 14 | 0 |
| X | 1728302 | 2 | 12 | 4 | 0 |
| X | 1734636 | 2 | 11 | 10 | 0 |
| X | 1737116 | 2 | 13 | 7 | 0 |
| X | 1740990 | 2 | 5 | 16 | 0 |
| X | 1744963 | 2 | 14 | 22 | 0 |
| X | 1748167 | 2 | 8 | 6 | 0 |
| X | 1751723 | 2 | 10 | 21 | 0 |
| X | 1766568 | 2 | 5 | 24 | 0 |
| X | 1766605 | 2 | 5 | 17 | 0 |
| X | 1766832 | 2 | 14 | 20 | 0 |
| X | 1767952 | 2 | 5 | 11 | 0 |
| X | 1768086 | 2 | 7 | 20 | 0 |
| X | 1768897 | 2 | 10 | 21 | 0 |
| X | 1769515 | 2 | 8 | 7 | 0 |
| X | 1769936 | 2 | 4 | 15 | 0 |
| X | 1770935 | 2 | 11 | 9 | 0 |
| X | 1773140 | 2 | 7 | 7 | 0 |
| X | 1773387 | 2 | 22 | 24 | 0 |
| X | 1773925 | 2 | 9 | 16 | 0 |
| X | 1779603 | 2 | 9 | 35 | 0 |
| X | 1783154 | 2 | 5 | 18 | 0 |
| X | 1785563 | 2 | 5 | 8 | 0 |
| X | 1789339 | 2 | 13 | 16 | 0 |
| X | 1792778 | 2 | 17 | 22 | 0 |
| X | 1794940 | 2 | 16 | 23 | 0 |
| X | 1795189 | 2 | 4 | 29 | 0 |
| X | 1795872 | 2 | 4 | 20 | 0 |
| X | 1796940 | 2 | 16 | 22 | 0 |
| X | 1802196 | 2 | 11 | 12 | 0 |
| X | 1813236 | 2 | 4 | 16 | 0 |
| X | 1814777 | 2 | 8 | 16 | 0 |
| X | 1817388 | 2 | 13 | 24 | 0 |
| X | 1817624 | 2 | 5 | 37 | 0 |
| X | 1818569 | 2 | 24 | 10 | 0 |
| X | 1820377 | 2 | 22 | 10 | 0 |
| X | 1823012 | 2 | 6 | 21 | 0 |
| X | 1838634 | 2 | 17 | 6 | 0 |
| X | 1842099 | 2 | 22 | 14 | 0 |
| X | 1845445 | 2 | 8 | 12 | 0 |
| X | 1847106 | 2 | 14 | 7 | 0 |
| X | 1859293 | 2 | 7 | 14 | 0 |
| X | 1861329 | 2 | 6 | 16 | 0 |
| X | 1864241 | 2 | 14 | 6 | 0 |
| X | 1866400 | 2 | 6 | 7 | 0 |
| X | 1868231 | 2 | 8 | 26 | 0 |
| X | 1869484 | 2 | 15 | 5 | 0 |
| X | 1874130 | 2 | 13 | 10 | 0 |
| X | 1874204 | 2 | 16 | 11 | 0 |
| X | 1878090 | 2 | 24 | 12 | 0 |
| X | 1879594 | 2 | 11 | 6 | 0 |
| X | 1879897 | 2 | 17 | 7 | 0 |
| X | 1881439 | 2 | 6 | 6 | 0 |
| X | 1890050 | 2 | 16 | 7 | 0 |
| X | 1891119 | 2 | 13 | 8 | 0 |
| X | 1892423 | 2 | 13 | 4 | 0 |
| X | 1892668 | 2 | 13 | 23 | 0 |
| X | 1893140 | 2 | 9 | 5 | 0 |
| X | 1893590 | 2 | 10 | 4 | 0 |
| X | 1897606 | 2 | 30 | 18 | 0 |
| X | 1898524 | 2 | 4 | 22 | 0 |
| X | 1899304 | 2 | 19 | 14 | 0 |
| X | 1901006 | 2 | 26 | 9 | 0 |
| X | 1902204 | 2 | 19 | 15 | 0 |
| X | 1904151 | 2 | 8 | 29 | 0 |
| X | 1904337 | 2 | 5 | 9 | 0 |
| X | 1906087 | 2 | 6 | 22 | 0 |
| X | 1907556 | 2 | 6 | 25 | 0 |
| X | 1907617 | 2 | 7 | 6 | 0 |
| X | 1908658 | 2 | 14 | 12 | 0 |
| X | 1912567 | 2 | 21 | 30 | 0 |
| X | 1913190 | 2 | 11 | 20 | 0 |
| X | 1914463 | 2 | 24 | 29 | 0 |
| X | 1914517 | 2 | 18 | 20 | 0 |
| X | 1925704 | 2 | 11 | 27 | 0 |
| X | 1928000 | 2 | 7 | 14 | 0 |
| X | 1929952 | 2 | 17 | 7 | 0 |
| X | 1935172 | 2 | 24 | 10 | 0 |
| X | 1935561 | 2 | 22 | 14 | 0 |
| X | 1936455 | 2 | 8 | 5 | 0 |
| X | 1936583 | 2 | 5 | 10 | 0 |
| X | 1944947 | 2 | 14 | 5 | 0 |
| X | 1946221 | 2 | 11 | 10 | 0 |
| X | 1950139 | 2 | 12 | 23 | 0 |
| X | 1950147 | 2 | 11 | 24 | 0 |
| X | 1950522 | 2 | 14 | 24 | 0 |
| X | 1954261 | 2 | 18 | 8 | 0 |
| X | 1954901 | 2 | 5 | 10 | 0 |
| X | 1955038 | 2 | 5 | 9 | 0 |
| X | 1955500 | 2 | 27 | 15 | 0 |
| X | 1958445 | 2 | 30 | 14 | 0 |
| X | 1958482 | 2 | 34 | 13 | 0 |
| X | 1963558 | 2 | 19 | 4 | 0 |
| X | 1965071 | 2 | 22 | 7 | 0 |
| X | 1965415 | 2 | 20 | 11 | 0 |
| X | 1965894 | 2 | 29 | 14 | 0 |
| X | 1966057 | 2 | 17 | 8 | 0 |
| X | 1966212 | 2 | 20 | 26 | 0 |
| X | 1966345 | 2 | 12 | 6 | 0 |
| X | 1966439 | 2 | 22 | 7 | 0 |
| X | 1967335 | 2 | 15 | 6 | 0 |
| X | 1967685 | 2 | 23 | 13 | 0 |
| X | 1968192 | 2 | 19 | 6 | 0 |
| X | 1968446 | 2 | 16 | 4 | 0 |
| X | 1969139 | 2 | 19 | 6 | 0 |
| X | 1969152 | 2 | 28 | 9 | 0 |
| X | 1970678 | 2 | 21 | 12 | 0 |
| X | 1970788 | 2 | 22 | 13 | 0 |
| X | 1971109 | 2 | 23 | 7 | 0 |
| X | 1971436 | 2 | 15 | 12 | 0 |
| X | 1972008 | 2 | 9 | 14 | 0 |
| X | 1972323 | 2 | 24 | 18 | 0 |
| X | 1974864 | 2 | 14 | 5 | 0 |
| X | 1975347 | 2 | 10 | 23 | 0 |
| X | 1975933 | 2 | 16 | 23 | 0 |
| X | 1978074 | 2 | 33 | 12 | 0 |
| X | 1978207 | 2 | 29 | 10 | 0 |
| X | 1982463 | 2 | 15 | 6 | 0 |
| X | 1982477 | 2 | 13 | 16 | 0 |
| X | 1987412 | 2 | 24 | 17 | 0 |
| X | 1987469 | 2 | 8 | 5 | 0 |
| X | 1994860 | 2 | 7 | 13 | 0 |
| X | 1995070 | 2 | 12 | 15 | 0 |
| X | 1998412 | 2 | 11 | 5 | 0 |
| X | 1999073 | 2 | 14 | 8 | 0 |
| X | 2000142 | 2 | 30 | 15 | 0 |
| X | 2000498 | 2 | 11 | 6 | 0 |
| X | 2003004 | 2 | 9 | 14 | 0 |
| X | 2003928 | 2 | 4 | 9 | 0 |
| X | 2007720 | 2 | 7 | 13 | 0 |
| X | 2019473 | 2 | 10 | 4 | 0 |
| X | 2027919 | 2 | 6 | 17 | 0 |
| X | 2036921 | 2 | 10 | 7 | 0 |
| X | 2042435 | 2 | 11 | 15 | 0 |
| X | 2044250 | 2 | 4 | 10 | 0 |
| X | 2058020 | 2 | 5 | 7 | 0 |
| X | 2063950 | 2 | 8 | 21 | 0 |
| X | 2064580 | 2 | 11 | 5 | 0 |
| X | 2066230 | 2 | 27 | 6 | 0 |
| X | 2067413 | 2 | 14 | 21 | 0 |
| X | 2069598 | 2 | 22 | 10 | 0 |
| X | 2069996 | 2 | 17 | 19 | 0 |
| X | 2070166 | 2 | 14 | 4 | 0 |
| X | 2070200 | 2 | 15 | 7 | 0 |
| X | 2070473 | 2 | 32 | 14 | 0 |
| X | 2070548 | 2 | 11 | 4 | 0 |
| X | 2070643 | 2 | 15 | 5 | 0 |
| X | 2070807 | 2 | 8 | 19 | 0 |
| X | 2071035 | 2 | 4 | 15 | 0 |
| X | 2072377 | 2 | 13 | 30 | 0 |
| X | 2072467 | 2 | 6 | 24 | 0 |
| X | 2073207 | 2 | 12 | 22 | 0 |
| X | 2073303 | 2 | 16 | 32 | 0 |
| X | 2073408 | 2 | 26 | 24 | 0 |
| X | 2073570 | 2 | 14 | 13 | 0 |
| X | 2074581 | 2 | 16 | 28 | 0 |
| X | 2076057 | 2 | 7 | 21 | 0 |
| X | 2076641 | 2 | 7 | 19 | 0 |
| X | 2077402 | 2 | 7 | 25 | 0 |
| X | 2078974 | 2 | 6 | 10 | 0 |
| X | 2078985 | 2 | 19 | 10 | 0 |
| X | 2079702 | 2 | 5 | 19 | 0 |
| X | 2080051 | 2 | 7 | 14 | 0 |
| X | 2080632 | 2 | 6 | 15 | 0 |
| X | 2080940 | 2 | 21 | 17 | 0 |
| X | 2080953 | 2 | 26 | 12 | 0 |
| X | 2081515 | 2 | 5 | 12 | 0 |
| X | 2082834 | 2 | 24 | 9 | 0 |
| X | 2086653 | 2 | 4 | 10 | 0 |
| X | 2086871 | 2 | 9 | 19 | 0 |
| X | 2086921 | 2 | 7 | 24 | 0 |
| X | 2088403 | 2 | 30 | 13 | 0 |
| X | 2088787 | 2 | 16 | 4 | 0 |
| X | 2088912 | 2 | 4 | 19 | 0 |
| X | 2089017 | 2 | 17 | 26 | 0 |
| X | 2089295 | 2 | 13 | 4 | 0 |
| X | 2089990 | 2 | 15 | 8 | 0 |
| X | 2090045 | 2 | 29 | 21 | 0 |
| X | 2090363 | 2 | 9 | 8 | 0 |
| X | 2090408 | 2 | 18 | 12 | 0 |
| X | 2090818 | 2 | 17 | 11 | 0 |
| X | 2092012 | 2 | 21 | 4 | 0 |
| X | 2092097 | 2 | 11 | 4 | 0 |
| X | 2092485 | 2 | 34 | 6 | 0 |
| X | 2093653 | 2 | 17 | 5 | 0 |
| X | 2094216 | 2 | 15 | 9 | 0 |
| X | 2094522 | 2 | 13 | 6 | 0 |
| X | 2095202 | 2 | 19 | 29 | 0 |
| X | 2095710 | 2 | 29 | 28 | 0 |
| X | 2096260 | 2 | 32 | 33 | 0 |
| X | 2098790 | 2 | 21 | 34 | 0 |
| X | 2099732 | 2 | 13 | 12 | 0 |
| X | 2107584 | 2 | 33 | 16 | 0 |
| X | 2110740 | 2 | 12 | 13 | 0 |
| X | 2115468 | 2 | 15 | 27 | 0 |
| X | 2122748 | 2 | 12 | 19 | 0 |
| X | 2125288 | 2 | 16 | 9 | 0 |
| X | 2132247 | 2 | 23 | 14 | 0 |
| X | 2133287 | 2 | 15 | 5 | 0 |
| X | 2133362 | 2 | 17 | 12 | 0 |
| X | 2133395 | 2 | 14 | 7 | 0 |
| X | 2137013 | 2 | 13 | 4 | 0 |
| X | 2138643 | 2 | 24 | 20 | 0 |
| X | 2139659 | 2 | 6 | 13 | 0 |
| X | 2145023 | 2 | 7 | 26 | 0 |
| X | 2145127 | 2 | 11 | 13 | 0 |
| X | 2146008 | 2 | 9 | 17 | 0 |
| X | 2147401 | 2 | 15 | 28 | 0 |
| X | 2147463 | 2 | 21 | 16 | 0 |
| X | 2147663 | 2 | 6 | 32 | 0 |
| X | 2148093 | 2 | 4 | 20 | 0 |
| X | 2148131 | 2 | 11 | 19 | 0 |
| X | 2148848 | 2 | 12 | 26 | 0 |
| X | 2149239 | 2 | 8 | 26 | 0 |
| X | 2149503 | 2 | 8 | 6 | 0 |
| X | 2149561 | 2 | 22 | 15 | 0 |
| X | 2149613 | 2 | 13 | 4 | 0 |
| X | 2150857 | 2 | 14 | 40 | 0 |
| X | 2150964 | 2 | 6 | 12 | 0 |
| X | 2151052 | 2 | 12 | 13 | 0 |
| X | 2151492 | 2 | 7 | 10 | 0 |
| X | 2151574 | 2 | 7 | 26 | 0 |
| X | 2151827 | 2 | 9 | 16 | 0 |
| X | 2151878 | 2 | 4 | 25 | 0 |
| X | 2152227 | 2 | 5 | 15 | 0 |
| X | 2152533 | 2 | 9 | 19 | 0 |
| X | 2152633 | 2 | 4 | 16 | 0 |
| X | 2152715 | 2 | 4 | 13 | 0 |
| X | 2152809 | 2 | 11 | 17 | 0 |
| X | 2152959 | 2 | 24 | 9 | 0 |
| X | 2153008 | 2 | 21 | 6 | 0 |
| X | 2153049 | 2 | 10 | 10 | 0 |
| X | 2153278 | 2 | 11 | 6 | 0 |
| X | 2153339 | 2 | 20 | 10 | 0 |
| X | 2154312 | 2 | 13 | 12 | 0 |
| X | 2154422 | 2 | 12 | 18 | 0 |
| X | 2155003 | 2 | 21 | 4 | 0 |
| X | 2155219 | 2 | 20 | 11 | 0 |
| X | 2155372 | 2 | 11 | 7 | 0 |
| X | 2155901 | 2 | 19 | 5 | 0 |
| X | 2156287 | 2 | 5 | 13 | 0 |
| X | 2156552 | 2 | 5 | 15 | 0 |
| X | 2157402 | 2 | 5 | 7 | 0 |
| X | 2157773 | 2 | 13 | 7 | 0 |
| X | 2157806 | 2 | 12 | 9 | 0 |
| X | 2157896 | 2 | 23 | 15 | 0 |
| X | 2159808 | 2 | 9 | 21 | 0 |
| X | 2159896 | 2 | 15 | 27 | 0 |
| X | 2160111 | 2 | 12 | 17 | 0 |
| X | 2160246 | 2 | 14 | 34 | 0 |
| X | 2160272 | 2 | 10 | 32 | 0 |
| X | 2160758 | 2 | 6 | 16 | 0 |
| X | 2160809 | 2 | 9 | 19 | 0 |
| X | 2161376 | 2 | 9 | 9 | 0 |
| X | 2161428 | 2 | 6 | 23 | 0 |
| X | 2161527 | 2 | 6 | 8 | 0 |
| X | 2161615 | 2 | 19 | 8 | 0 |
| X | 2161688 | 2 | 5 | 17 | 0 |
| X | 2162599 | 2 | 17 | 27 | 0 |
| X | 2163221 | 2 | 6 | 9 | 0 |
| X | 2164940 | 2 | 14 | 5 | 0 |
| X | 2168253 | 2 | 11 | 21 | 0 |
| X | 2168428 | 2 | 10 | 17 | 0 |
| X | 2168524 | 2 | 5 | 15 | 0 |
| X | 2168621 | 2 | 4 | 24 | 0 |
| X | 2171932 | 2 | 10 | 31 | 0 |
| X | 2181270 | 2 | 9 | 25 | 0 |
| X | 2183639 | 2 | 13 | 12 | 0 |
| X | 2185817 | 2 | 10 | 4 | 0 |
| X | 2185896 | 2 | 7 | 11 | 0 |
| X | 2188448 | 2 | 24 | 8 | 0 |
| X | 2191805 | 2 | 15 | 15 | 0 |
| X | 2191939 | 2 | 10 | 11 | 0 |
| X | 2192368 | 2 | 22 | 28 | 0 |
| X | 2193228 | 2 | 7 | 29 | 0 |
| X | 2194729 | 2 | 10 | 25 | 0 |
| X | 2200750 | 2 | 13 | 16 | 0 |
| X | 2201159 | 2 | 13 | 7 | 0 |
| X | 2201471 | 2 | 15 | 4 | 0 |
| X | 2201710 | 2 | 38 | 13 | 0 |
| X | 2201843 | 2 | 14 | 7 | 0 |
| X | 2202279 | 2 | 16 | 4 | 0 |
| X | 2202327 | 2 | 5 | 6 | 0 |
| X | 2202381 | 2 | 27 | 11 | 0 |
| X | 2202712 | 2 | 8 | 5 | 0 |
| X | 2211436 | 2 | 13 | 4 | 0 |
| X | 2211532 | 2 | 8 | 10 | 0 |
| X | 2212152 | 2 | 22 | 21 | 0 |
| X | 2212348 | 2 | 26 | 7 | 0 |
| X | 2212468 | 2 | 5 | 34 | 0 |
| X | 2212644 | 2 | 4 | 23 | 0 |
| X | 2212719 | 2 | 22 | 17 | 0 |
| X | 2214728 | 2 | 15 | 4 | 0 |
| X | 2215797 | 2 | 21 | 4 | 0 |
| X | 2216855 | 2 | 7 | 20 | 0 |
| X | 2216916 | 2 | 6 | 21 | 0 |
| X | 2218075 | 2 | 14 | 20 | 0 |
| X | 2219071 | 2 | 20 | 4 | 0 |
| X | 2219760 | 2 | 20 | 4 | 0 |
| X | 2220068 | 2 | 10 | 9 | 0 |
| X | 2220174 | 2 | 9 | 7 | 0 |
| X | 2220398 | 2 | 8 | 11 | 0 |
| X | 2221732 | 2 | 20 | 16 | 0 |
| X | 2221749 | 2 | 14 | 18 | 0 |
| X | 2222244 | 2 | 4 | 24 | 0 |
| X | 2222793 | 2 | 15 | 23 | 0 |
| X | 2223170 | 2 | 19 | 8 | 0 |
| X | 2224361 | 2 | 10 | 32 | 0 |
| X | 2224407 | 2 | 7 | 24 | 0 |
| X | 2225147 | 2 | 19 | 10 | 0 |
| X | 2226260 | 2 | 5 | 10 | 0 |
| X | 2370401 | 2 | 5 | 5 | 0 |
| X | 2383061 | 2 | 16 | 4 | 0 |
| X | 2390295 | 2 | 6 | 8 | 0 |
| X | 2396358 | 2 | 7 | 8 | 0 |
| X | 2412991 | 2 | 5 | 5 | 0 |
| X | 2423972 | 2 | 11 | 4 | 0 |
| X | 2540533 | 2 | 10 | 4 | 0 |
| X | 2577318 | 2 | 5 | 6 | 0 |
| X | 2653150 | 2 | 9 | 4 | 0 |
| X | 2806380 | 2 | 5 | 5 | 0 |
| X | 2948513 | 2 | 6 | 9 | 0 |
| X | 3085776 | 2 | 6 | 6 | 0 |
| X | 3096748 | 2 | 6 | 5 | 0 |
| X | 3361491 | 2 | 6 | 6 | 0 |
| X | 3440536 | 2 | 4 | 12 | 0 |
| X | 3447473 | 2 | 5 | 5 | 0 |
| X | 3461475 | 2 | 7 | 5 | 0 |
| X | 3821979 | 2 | 4 | 18 | 0 |
| X | 3822585 | 2 | 10 | 10 | 0 |
| X | 3822862 | 2 | 8 | 6 | 0 |
| X | 3823831 | 2 | 4 | 11 | 0 |
| X | 3833327 | 2 | 10 | 5 | 0 |
| X | 3845828 | 2 | 16 | 8 | 0 |
| X | 3850497 | 2 | 9 | 6 | 0 |
| X | 3851006 | 2 | 17 | 7 | 0 |
| X | 3856331 | 2 | 23 | 6 | 0 |
| X | 3856379 | 2 | 22 | 8 | 0 |
| X | 3856598 | 2 | 26 | 4 | 0 |
| X | 3856673 | 2 | 21 | 5 | 0 |
| X | 3856811 | 2 | 7 | 7 | 0 |
| X | 3857112 | 2 | 14 | 7 | 0 |
| X | 3857563 | 2 | 21 | 9 | 0 |
| X | 3857621 | 2 | 25 | 20 | 0 |
| X | 3857876 | 2 | 17 | 9 | 0 |
| X | 3858259 | 2 | 17 | 10 | 0 |
| X | 3858500 | 2 | 13 | 8 | 0 |
| X | 3860924 | 2 | 17 | 7 | 0 |
| X | 3861002 | 2 | 14 | 26 | 0 |
| X | 3861865 | 2 | 14 | 5 | 0 |
| X | 3862021 | 2 | 4 | 20 | 0 |
| X | 3862049 | 2 | 12 | 19 | 0 |
| X | 3863295 | 2 | 17 | 8 | 0 |
| X | 3863469 | 2 | 19 | 5 | 0 |
| X | 3863682 | 2 | 14 | 22 | 0 |
| X | 3863748 | 2 | 17 | 5 | 0 |
| X | 3864291 | 2 | 10 | 4 | 0 |
| X | 3865445 | 2 | 27 | 10 | 0 |
| X | 3865618 | 2 | 7 | 12 | 0 |
| X | 3866607 | 2 | 6 | 15 | 0 |
| X | 3867097 | 2 | 5 | 15 | 0 |
| X | 3867903 | 2 | 4 | 17 | 0 |
| X | 3873005 | 2 | 10 | 17 | 0 |
| X | 3874006 | 2 | 5 | 18 | 0 |
| X | 3874089 | 2 | 15 | 23 | 0 |
| X | 3874470 | 2 | 10 | 14 | 0 |
| X | 3874669 | 2 | 4 | 22 | 0 |
| X | 3874719 | 2 | 6 | 17 | 0 |
| X | 3876251 | 2 | 18 | 4 | 0 |
| X | 3877988 | 2 | 11 | 9 | 0 |
| X | 3880718 | 2 | 20 | 30 | 0 |
| X | 3880871 | 2 | 7 | 22 | 0 |
| X | 3890637 | 2 | 8 | 13 | 0 |
| X | 3890755 | 2 | 8 | 10 | 0 |
| X | 3891142 | 2 | 9 | 22 | 0 |
| X | 3891593 | 2 | 6 | 7 | 0 |
| X | 3891955 | 2 | 27 | 8 | 0 |
| X | 3894816 | 2 | 10 | 4 | 0 |
| X | 3894877 | 2 | 18 | 4 | 0 |
| X | 3894992 | 2 | 13 | 8 | 0 |
| X | 3895591 | 2 | 19 | 11 | 0 |
| X | 3895838 | 2 | 7 | 10 | 0 |
| X | 3896310 | 2 | 12 | 6 | 0 |
| X | 3896933 | 2 | 10 | 35 | 0 |
| X | 3897679 | 2 | 8 | 5 | 0 |
| X | 3897812 | 2 | 21 | 20 | 0 |
| X | 3898476 | 2 | 17 | 4 | 0 |
| X | 3904584 | 2 | 4 | 19 | 0 |
| X | 3907049 | 2 | 7 | 21 | 0 |
| X | 3909387 | 2 | 26 | 7 | 0 |
| X | 3913619 | 2 | 6 | 24 | 0 |
| X | 3915165 | 2 | 9 | 4 | 0 |
| X | 3915561 | 2 | 17 | 4 | 0 |
| X | 3917444 | 2 | 7 | 23 | 0 |
| X | 3922259 | 2 | 7 | 20 | 0 |
| X | 3922668 | 2 | 12 | 22 | 0 |
| X | 3922847 | 2 | 7 | 13 | 0 |
| X | 3926664 | 2 | 26 | 14 | 0 |
| X | 3928163 | 2 | 16 | 10 | 0 |
| X | 3929193 | 2 | 18 | 7 | 0 |
| X | 3929296 | 2 | 4 | 10 | 0 |
| X | 3930491 | 2 | 13 | 14 | 0 |
| X | 3930823 | 2 | 10 | 15 | 0 |
| X | 3934182 | 2 | 24 | 12 | 0 |
| X | 3936895 | 2 | 12 | 9 | 0 |
| X | 3937507 | 2 | 13 | 6 | 0 |
| X | 3937863 | 2 | 4 | 9 | 0 |
| X | 3939230 | 2 | 13 | 22 | 0 |
| X | 3940134 | 2 | 10 | 22 | 0 |
| X | 3943048 | 2 | 18 | 4 | 0 |
| X | 3945087 | 2 | 10 | 20 | 0 |
| X | 3947415 | 2 | 6 | 19 | 0 |
| X | 3948585 | 2 | 12 | 4 | 0 |
| X | 3949334 | 2 | 23 | 14 | 0 |
| X | 3950574 | 2 | 23 | 9 | 0 |
| X | 3950633 | 2 | 15 | 13 | 0 |
| X | 3950648 | 2 | 6 | 12 | 0 |
| X | 3953588 | 2 | 16 | 10 | 0 |
| X | 3954005 | 2 | 16 | 7 | 0 |
| X | 3955308 | 2 | 13 | 6 | 0 |
| X | 3955632 | 2 | 4 | 15 | 0 |
| X | 3956168 | 2 | 12 | 7 | 0 |
| X | 4142867 | 2 | 10 | 5 | 0 |
| X | 4176740 | 2 | 5 | 5 | 0 |
| X | 4273318 | 2 | 10 | 7 | 0 |
| X | 4290049 | 2 | 9 | 5 | 0 |
| X | 4295041 | 2 | 5 | 5 | 0 |
| X | 4315317 | 2 | 8 | 5 | 0 |
| X | 4324268 | 2 | 11 | 6 | 0 |
| X | 4343964 | 2 | 13 | 6 | 0 |
| X | 4344267 | 2 | 16 | 6 | 0 |
| X | 4350318 | 2 | 12 | 4 | 0 |
| X | 4352314 | 2 | 8 | 15 | 0 |
| X | 4358444 | 2 | 14 | 18 | 0 |
| X | 4359676 | 2 | 15 | 9 | 0 |
| X | 4362182 | 2 | 28 | 13 | 0 |
| X | 4362631 | 2 | 11 | 4 | 0 |
| X | 4362683 | 2 | 11 | 5 | 0 |
| X | 4364063 | 2 | 10 | 17 | 0 |
| X | 4364090 | 2 | 5 | 12 | 0 |
| X | 4366096 | 2 | 20 | 21 | 0 |
| X | 4366334 | 2 | 19 | 14 | 0 |
| X | 4366458 | 2 | 23 | 13 | 0 |
| X | 4372537 | 2 | 8 | 18 | 0 |
| X | 4376227 | 2 | 5 | 8 | 0 |
| X | 4379875 | 2 | 6 | 13 | 0 |
| X | 4380980 | 2 | 6 | 18 | 0 |
| X | 4387452 | 2 | 9 | 6 | 0 |
| X | 4387759 | 2 | 4 | 11 | 0 |
| X | 4391602 | 2 | 6 | 7 | 0 |
| X | 4391858 | 2 | 12 | 7 | 0 |
| X | 4393058 | 2 | 13 | 9 | 0 |
| X | 4394067 | 2 | 19 | 5 | 0 |
| X | 4397785 | 2 | 21 | 10 | 0 |
| X | 4398192 | 2 | 4 | 23 | 0 |
| X | 4399125 | 2 | 14 | 27 | 0 |
| X | 4401421 | 2 | 6 | 12 | 0 |
| X | 4411912 | 2 | 10 | 21 | 0 |
| X | 4414228 | 2 | 10 | 24 | 0 |
| X | 4416638 | 2 | 19 | 6 | 0 |
| X | 4418108 | 2 | 7 | 10 | 0 |
| X | 4418557 | 2 | 6 | 10 | 0 |
| X | 4419765 | 2 | 13 | 9 | 0 |
| X | 4421330 | 2 | 7 | 8 | 0 |
| X | 4423626 | 2 | 15 | 6 | 0 |
| X | 4424162 | 2 | 8 | 9 | 0 |
| X | 4424560 | 2 | 7 | 6 | 0 |
| X | 4424762 | 2 | 7 | 9 | 0 |
| X | 4425357 | 2 | 15 | 15 | 0 |
| X | 4425385 | 2 | 24 | 15 | 0 |
| X | 4425553 | 2 | 8 | 13 | 0 |
| X | 4426029 | 2 | 5 | 7 | 0 |
| X | 4426053 | 2 | 4 | 10 | 0 |
| X | 4427040 | 2 | 7 | 22 | 0 |
| X | 4429167 | 2 | 16 | 5 | 0 |
| X | 4430198 | 2 | 12 | 12 | 0 |
| X | 4430513 | 2 | 4 | 12 | 0 |
| X | 4431334 | 2 | 27 | 9 | 0 |
| X | 4431388 | 2 | 11 | 6 | 0 |
| X | 4432108 | 2 | 5 | 10 | 0 |
| X | 4433003 | 2 | 15 | 9 | 0 |
| X | 4433253 | 2 | 10 | 4 | 0 |
| X | 4433403 | 2 | 25 | 23 | 0 |
| X | 4434162 | 2 | 4 | 10 | 0 |
| X | 4434354 | 2 | 14 | 11 | 0 |
| X | 4434502 | 2 | 26 | 22 | 0 |
| X | 4435296 | 2 | 18 | 29 | 0 |
| X | 4441463 | 2 | 7 | 14 | 0 |
| X | 4443812 | 2 | 5 | 18 | 0 |
| X | 4447851 | 2 | 14 | 7 | 0 |
| X | 4455420 | 2 | 20 | 18 | 0 |
| X | 4461691 | 2 | 20 | 5 | 0 |
| X | 4461711 | 2 | 11 | 4 | 0 |
| X | 4462433 | 2 | 19 | 9 | 0 |
| X | 4462958 | 2 | 20 | 7 | 0 |
| X | 4463117 | 2 | 15 | 4 | 0 |
| X | 4465678 | 2 | 12 | 4 | 0 |
| X | 4467499 | 2 | 6 | 16 | 0 |
| X | 4469211 | 2 | 17 | 29 | 0 |
| X | 4469275 | 2 | 10 | 18 | 0 |
| X | 4469394 | 2 | 4 | 20 | 0 |
| X | 4469835 | 2 | 7 | 37 | 0 |
| X | 4472481 | 2 | 5 | 17 | 0 |
| X | 4476158 | 2 | 18 | 16 | 0 |
| X | 4481046 | 2 | 19 | 7 | 0 |
| X | 4481490 | 2 | 11 | 11 | 0 |
| X | 4485072 | 2 | 15 | 28 | 0 |
| X | 4486618 | 2 | 10 | 24 | 0 |
| X | 4487149 | 2 | 6 | 18 | 0 |
| X | 4487355 | 2 | 7 | 12 | 0 |
| X | 4488148 | 2 | 14 | 11 | 0 |
| X | 4488609 | 2 | 5 | 21 | 0 |
| X | 4490045 | 2 | 9 | 16 | 0 |
| X | 4490520 | 2 | 5 | 26 | 0 |
| X | 4491492 | 2 | 6 | 25 | 0 |
| X | 4491979 | 2 | 9 | 13 | 0 |
| X | 4492974 | 2 | 6 | 21 | 0 |
| X | 4498309 | 2 | 4 | 14 | 0 |
| X | 4498725 | 2 | 18 | 7 | 0 |
| X | 4500109 | 2 | 12 | 24 | 0 |
| X | 4510393 | 2 | 9 | 14 | 0 |
| X | 4510459 | 2 | 4 | 13 | 0 |
| X | 4514854 | 2 | 9 | 4 | 0 |
| X | 4516138 | 2 | 16 | 30 | 0 |
| X | 4519489 | 2 | 13 | 6 | 0 |
| X | 4520016 | 2 | 19 | 29 | 0 |
| X | 4520119 | 2 | 9 | 4 | 0 |
| X | 4520820 | 2 | 4 | 18 | 0 |
| X | 4522861 | 2 | 8 | 19 | 0 |
| X | 4526638 | 2 | 8 | 31 | 0 |
| X | 4530572 | 2 | 5 | 14 | 0 |
| X | 4532126 | 2 | 5 | 26 | 0 |
| X | 4532149 | 2 | 7 | 21 | 0 |
| X | 4532835 | 2 | 8 | 16 | 0 |
| X | 4533152 | 2 | 16 | 12 | 0 |
| X | 4536891 | 2 | 10 | 6 | 0 |
| X | 4537361 | 2 | 7 | 7 | 0 |
| X | 4538953 | 2 | 7 | 8 | 0 |
| X | 4539927 | 2 | 9 | 13 | 0 |
| X | 4541941 | 2 | 15 | 4 | 0 |
| X | 4543569 | 2 | 12 | 4 | 0 |
| X | 4547002 | 2 | 33 | 17 | 0 |
| X | 4548151 | 2 | 10 | 18 | 0 |
| X | 4550761 | 2 | 7 | 26 | 0 |
| X | 4551338 | 2 | 6 | 21 | 0 |
| X | 4553200 | 2 | 14 | 16 | 0 |
| X | 4554273 | 2 | 7 | 5 | 0 |
| X | 4555535 | 2 | 15 | 5 | 0 |
| X | 4555598 | 2 | 23 | 11 | 0 |
| X | 4556249 | 2 | 11 | 24 | 0 |
| X | 4557376 | 2 | 8 | 12 | 0 |
| X | 4562164 | 2 | 10 | 25 | 0 |
| X | 4563212 | 2 | 6 | 28 | 0 |
| X | 4563392 | 2 | 8 | 5 | 0 |
| X | 4564518 | 2 | 7 | 13 | 0 |
| X | 4566057 | 2 | 13 | 13 | 0 |
| X | 4566123 | 2 | 9 | 27 | 0 |
| X | 4568625 | 2 | 8 | 13 | 0 |
| X | 4570370 | 2 | 5 | 25 | 0 |
| X | 4571328 | 2 | 6 | 18 | 0 |
| X | 4571596 | 2 | 13 | 8 | 0 |
| X | 4573268 | 2 | 5 | 22 | 0 |
| X | 4573444 | 2 | 10 | 4 | 0 |
| X | 4573617 | 2 | 26 | 21 | 0 |
| X | 4574151 | 2 | 15 | 8 | 0 |
| X | 4574815 | 2 | 16 | 14 | 0 |
| X | 4575007 | 2 | 8 | 8 | 0 |
| X | 4575447 | 2 | 10 | 14 | 0 |
| X | 4575871 | 2 | 4 | 22 | 0 |
| X | 4576081 | 2 | 10 | 22 | 0 |
| X | 4586018 | 2 | 22 | 4 | 0 |
| X | 4586351 | 2 | 15 | 4 | 0 |
| X | 4590617 | 2 | 8 | 20 | 0 |
| X | 4593287 | 2 | 13 | 19 | 0 |
| X | 4598184 | 2 | 11 | 21 | 0 |
| X | 4598343 | 2 | 16 | 13 | 0 |
| X | 4599333 | 2 | 9 | 11 | 0 |
| X | 4599916 | 2 | 12 | 4 | 0 |
| X | 4603851 | 2 | 6 | 21 | 0 |
| X | 4606877 | 2 | 5 | 22 | 0 |
| X | 4609181 | 2 | 7 | 13 | 0 |
| X | 4611412 | 2 | 17 | 6 | 0 |
| X | 4611460 | 2 | 8 | 13 | 0 |
| X | 4611954 | 2 | 12 | 9 | 0 |
| X | 4612596 | 2 | 12 | 16 | 0 |
| X | 4613016 | 2 | 22 | 17 | 0 |
| X | 4613502 | 2 | 9 | 8 | 0 |
| X | 4613841 | 2 | 18 | 15 | 0 |
| X | 4613929 | 2 | 8 | 6 | 0 |
| X | 4615172 | 2 | 9 | 19 | 0 |
| X | 4616306 | 2 | 8 | 11 | 0 |
| X | 4616477 | 2 | 14 | 10 | 0 |
| X | 4618436 | 2 | 13 | 31 | 0 |
| X | 4618585 | 2 | 10 | 11 | 0 |
| X | 4618689 | 2 | 8 | 15 | 0 |
| X | 4621475 | 2 | 11 | 26 | 0 |
| X | 4622495 | 2 | 4 | 13 | 0 |
| X | 4622570 | 2 | 5 | 19 | 0 |
| X | 4624678 | 2 | 8 | 20 | 0 |
| X | 4632249 | 2 | 13 | 9 | 0 |
| X | 4635055 | 2 | 21 | 6 | 0 |
| X | 4635998 | 2 | 10 | 7 | 0 |
| X | 4637527 | 2 | 4 | 25 | 0 |
| X | 4638814 | 2 | 17 | 5 | 0 |
| X | 4640437 | 2 | 9 | 6 | 0 |
| X | 4641839 | 2 | 11 | 5 | 0 |
| X | 4641893 | 2 | 11 | 7 | 0 |
| X | 4643362 | 2 | 12 | 5 | 0 |
| X | 4643477 | 2 | 7 | 13 | 0 |
| X | 4644143 | 2 | 16 | 6 | 0 |
| X | 4644763 | 2 | 5 | 18 | 0 |
| X | 4645038 | 2 | 9 | 7 | 0 |
| X | 4645841 | 2 | 7 | 6 | 0 |
| X | 4646008 | 2 | 14 | 5 | 0 |
| X | 4649133 | 2 | 9 | 10 | 0 |
| X | 4650171 | 2 | 11 | 5 | 0 |
| X | 4651162 | 2 | 5 | 12 | 0 |
| X | 4652483 | 2 | 4 | 19 | 0 |
| X | 4654100 | 2 | 8 | 19 | 0 |
| X | 4654186 | 2 | 7 | 23 | 0 |
| X | 4658543 | 2 | 5 | 7 | 0 |
| X | 4658773 | 2 | 12 | 22 | 0 |
| X | 4665627 | 2 | 24 | 7 | 0 |
| X | 4665811 | 2 | 7 | 5 | 0 |
| X | 4665941 | 2 | 16 | 7 | 0 |
| X | 4669454 | 2 | 22 | 12 | 0 |
| X | 4672857 | 2 | 6 | 7 | 0 |
| X | 4677398 | 2 | 5 | 28 | 0 |
| X | 4677508 | 2 | 9 | 16 | 0 |
| X | 4678944 | 2 | 15 | 5 | 0 |
| X | 4680262 | 2 | 12 | 13 | 0 |
| X | 4687758 | 2 | 4 | 13 | 0 |
| X | 4687946 | 2 | 7 | 15 | 0 |
| X | 4687998 | 2 | 5 | 19 | 0 |
| X | 4688590 | 2 | 4 | 10 | 0 |
| X | 4698362 | 2 | 9 | 12 | 0 |
| X | 4698619 | 2 | 7 | 25 | 0 |
| X | 4699045 | 2 | 6 | 12 | 0 |
| X | 4703222 | 2 | 4 | 16 | 0 |
| X | 4705149 | 2 | 9 | 14 | 0 |
| X | 4711658 | 2 | 4 | 24 | 0 |
| X | 4711712 | 2 | 7 | 24 | 0 |
| X | 4712198 | 2 | 7 | 18 | 0 |
| X | 4713742 | 2 | 11 | 25 | 0 |
| X | 4714875 | 2 | 6 | 5 | 0 |
| X | 4719263 | 2 | 19 | 14 | 0 |
| X | 4721481 | 2 | 14 | 4 | 0 |
| X | 4721568 | 2 | 13 | 8 | 0 |
| X | 4729529 | 2 | 5 | 8 | 0 |
| X | 4730181 | 2 | 10 | 5 | 0 |
| X | 4730651 | 2 | 8 | 18 | 0 |
| X | 4732319 | 2 | 8 | 31 | 0 |
| X | 4740837 | 2 | 7 | 14 | 0 |
| X | 4741598 | 2 | 14 | 6 | 0 |
| X | 4742660 | 2 | 8 | 18 | 0 |
| X | 4744351 | 2 | 12 | 6 | 0 |
| X | 4744695 | 2 | 12 | 11 | 0 |
| X | 4744932 | 2 | 14 | 8 | 0 |
| X | 4748022 | 2 | 4 | 18 | 0 |
| X | 4750164 | 2 | 5 | 10 | 0 |
| X | 4752996 | 2 | 5 | 12 | 0 |
| X | 4754572 | 2 | 6 | 10 | 0 |
| X | 4757786 | 2 | 27 | 7 | 0 |
| X | 4760054 | 2 | 22 | 10 | 0 |
| X | 4760074 | 2 | 25 | 10 | 0 |
| X | 4760800 | 2 | 26 | 4 | 0 |
| X | 4762956 | 2 | 8 | 5 | 0 |
| X | 4766514 | 2 | 9 | 14 | 0 |
| X | 4767956 | 2 | 12 | 4 | 0 |
| X | 4768224 | 2 | 16 | 7 | 0 |
| X | 4769583 | 2 | 18 | 11 | 0 |
| X | 4772229 | 2 | 9 | 22 | 0 |
| X | 4781058 | 2 | 7 | 19 | 0 |
| X | 4784013 | 2 | 14 | 8 | 0 |
| X | 4784765 | 2 | 16 | 6 | 0 |
| X | 4786220 | 2 | 10 | 9 | 0 |
| X | 4791152 | 2 | 10 | 6 | 0 |
| X | 4792114 | 2 | 12 | 8 | 0 |
| X | 4794239 | 2 | 12 | 8 | 0 |
| X | 4795684 | 2 | 15 | 7 | 0 |
| X | 4795976 | 2 | 12 | 9 | 0 |
| X | 4796268 | 2 | 24 | 6 | 0 |
| X | 4797441 | 2 | 10 | 4 | 0 |
| X | 4797958 | 2 | 5 | 21 | 0 |
| X | 4801984 | 2 | 12 | 7 | 0 |
| X | 4802085 | 2 | 16 | 13 | 0 |
| X | 4805082 | 2 | 8 | 6 | 0 |
| X | 4806727 | 2 | 32 | 9 | 0 |
| X | 4807039 | 2 | 5 | 18 | 0 |
| X | 4807560 | 2 | 8 | 14 | 0 |
| X | 4808602 | 2 | 16 | 5 | 0 |
| X | 4809568 | 2 | 4 | 23 | 0 |
| X | 4810340 | 2 | 10 | 32 | 0 |
| X | 4811711 | 2 | 10 | 20 | 0 |
| X | 4828807 | 2 | 11 | 4 | 0 |
| X | 4832397 | 2 | 6 | 11 | 0 |
| X | 4835487 | 2 | 4 | 20 | 0 |
| X | 4836041 | 2 | 10 | 15 | 0 |
| X | 4839261 | 2 | 14 | 8 | 0 |
| X | 4840785 | 2 | 15 | 26 | 0 |
| X | 4841798 | 2 | 25 | 8 | 0 |
| X | 4842343 | 2 | 17 | 33 | 0 |
| X | 4843304 | 2 | 9 | 6 | 0 |
| X | 4847002 | 2 | 10 | 24 | 0 |
| X | 4847227 | 2 | 6 | 24 | 0 |
| X | 4847368 | 2 | 5 | 6 | 0 |
| X | 4847701 | 2 | 16 | 19 | 0 |
| X | 4847901 | 2 | 20 | 7 | 0 |
| X | 4848057 | 2 | 6 | 28 | 0 |
| X | 4857797 | 2 | 9 | 18 | 0 |
| X | 4860345 | 2 | 5 | 10 | 0 |
| X | 4862763 | 2 | 4 | 14 | 0 |
| X | 4863828 | 2 | 8 | 25 | 0 |
| X | 4864752 | 2 | 9 | 24 | 0 |
| X | 4869234 | 2 | 4 | 9 | 0 |
| X | 4873192 | 2 | 6 | 23 | 0 |
| X | 4875012 | 2 | 8 | 15 | 0 |
| X | 4875332 | 2 | 15 | 15 | 0 |
| X | 4875657 | 2 | 8 | 15 | 0 |
| X | 4877751 | 2 | 10 | 10 | 0 |
| X | 4878600 | 2 | 6 | 14 | 0 |
| X | 4878944 | 2 | 4 | 28 | 0 |
| X | 4882113 | 2 | 9 | 22 | 0 |
| X | 4882668 | 2 | 5 | 28 | 0 |
| X | 4882727 | 2 | 6 | 16 | 0 |
| X | 4892492 | 2 | 11 | 22 | 0 |
| X | 4895958 | 2 | 6 | 17 | 0 |
| X | 4897193 | 2 | 7 | 23 | 0 |
| X | 4897947 | 2 | 8 | 23 | 0 |
| X | 4898999 | 2 | 5 | 8 | 0 |
| X | 4901521 | 2 | 11 | 11 | 0 |
| X | 4906939 | 2 | 19 | 10 | 0 |
| X | 4912620 | 2 | 10 | 16 | 0 |
| X | 4918762 | 2 | 6 | 5 | 0 |
| X | 4921109 | 2 | 8 | 25 | 0 |
| X | 4927829 | 2 | 5 | 5 | 0 |
| X | 4928010 | 2 | 16 | 10 | 0 |
| X | 4932058 | 2 | 4 | 25 | 0 |
| X | 4933062 | 2 | 13 | 10 | 0 |
| X | 4933132 | 2 | 11 | 6 | 0 |
| X | 4934609 | 2 | 10 | 12 | 0 |
| X | 4935896 | 2 | 8 | 5 | 0 |
| X | 4936626 | 2 | 15 | 8 | 0 |
| X | 4938395 | 2 | 7 | 15 | 0 |
| X | 4938420 | 2 | 5 | 12 | 0 |
| X | 4942575 | 2 | 6 | 5 | 0 |
| X | 4946931 | 2 | 6 | 30 | 0 |
| X | 4951027 | 2 | 18 | 7 | 0 |
| X | 4965559 | 2 | 11 | 9 | 0 |
| X | 4976067 | 2 | 12 | 8 | 0 |
| X | 4976381 | 2 | 5 | 6 | 0 |
| X | 4982668 | 2 | 4 | 14 | 0 |
| X | 4983506 | 2 | 22 | 4 | 0 |
| X | 4983933 | 2 | 25 | 7 | 0 |
| X | 4989352 | 2 | 6 | 21 | 0 |
| X | 4989579 | 2 | 9 | 16 | 0 |
| X | 5001524 | 2 | 12 | 16 | 0 |
| X | 5003599 | 2 | 18 | 5 | 0 |
| X | 5005903 | 2 | 4 | 9 | 0 |
| X | 5007134 | 2 | 9 | 7 | 0 |
| X | 5010924 | 2 | 21 | 5 | 0 |
| X | 5011070 | 2 | 17 | 4 | 0 |
| X | 5011192 | 2 | 25 | 7 | 0 |
| X | 5011932 | 2 | 11 | 9 | 0 |
| X | 5012444 | 2 | 13 | 19 | 0 |
| X | 5012495 | 2 | 8 | 10 | 0 |
| X | 5016273 | 2 | 29 | 22 | 0 |
| X | 5018773 | 2 | 4 | 10 | 0 |
| X | 5019193 | 2 | 8 | 9 | 0 |
| X | 5020319 | 2 | 10 | 25 | 0 |
| X | 5020472 | 2 | 6 | 16 | 0 |
| X | 5020991 | 2 | 5 | 9 | 0 |
| X | 5025901 | 2 | 4 | 17 | 0 |
| X | 5026191 | 2 | 4 | 13 | 0 |
| X | 5028526 | 2 | 17 | 9 | 0 |
| X | 5029140 | 2 | 9 | 19 | 0 |
| X | 5031020 | 2 | 18 | 27 | 0 |
| X | 5033133 | 2 | 6 | 11 | 0 |
| X | 5034060 | 2 | 5 | 18 | 0 |
| X | 5034205 | 2 | 7 | 12 | 0 |
| X | 5034434 | 2 | 4 | 14 | 0 |
| X | 5035722 | 2 | 11 | 11 | 0 |
| X | 5038603 | 2 | 15 | 6 | 0 |
| X | 5038860 | 2 | 6 | 13 | 0 |
| X | 5041026 | 2 | 4 | 12 | 0 |
| X | 5045882 | 2 | 13 | 7 | 0 |
| X | 5046692 | 2 | 16 | 7 | 0 |
| X | 5047749 | 2 | 7 | 18 | 0 |
| X | 5048639 | 2 | 7 | 20 | 0 |
| X | 5049093 | 2 | 5 | 11 | 0 |
| X | 5050075 | 2 | 5 | 6 | 0 |
| X | 5052538 | 2 | 24 | 4 | 0 |
| X | 5055557 | 2 | 7 | 7 | 0 |
| X | 5055592 | 2 | 5 | 15 | 0 |
| X | 5055648 | 2 | 4 | 20 | 0 |
| X | 5058924 | 2 | 13 | 5 | 0 |
| X | 5061367 | 2 | 7 | 25 | 0 |
| X | 5070295 | 2 | 5 | 25 | 0 |
| X | 5071111 | 2 | 7 | 14 | 0 |
| X | 5072123 | 2 | 12 | 17 | 0 |
| X | 5072191 | 2 | 9 | 29 | 0 |
| X | 5073593 | 2 | 8 | 12 | 0 |
| X | 5075686 | 2 | 15 | 5 | 0 |
| X | 5076275 | 2 | 11 | 6 | 0 |
| X | 5077138 | 2 | 5 | 13 | 0 |
| X | 5078718 | 2 | 5 | 23 | 0 |
| X | 5078823 | 2 | 7 | 17 | 0 |
| X | 5078933 | 2 | 11 | 21 | 0 |
| X | 5079091 | 2 | 15 | 10 | 0 |
| X | 5085992 | 2 | 27 | 6 | 0 |
| X | 5088298 | 2 | 14 | 15 | 0 |
| X | 5091924 | 2 | 14 | 9 | 0 |
| X | 5092301 | 2 | 8 | 24 | 0 |
| X | 5092610 | 2 | 7 | 14 | 0 |
| X | 5093625 | 2 | 6 | 5 | 0 |
| X | 5093904 | 2 | 12 | 16 | 0 |
| X | 5095364 | 2 | 23 | 23 | 0 |
| X | 5096566 | 2 | 11 | 6 | 0 |
| X | 5096943 | 2 | 18 | 8 | 0 |
| X | 5098901 | 2 | 7 | 26 | 0 |
| X | 5099856 | 2 | 9 | 7 | 0 |
| X | 5100067 | 2 | 5 | 6 | 0 |
| X | 5100234 | 2 | 6 | 7 | 0 |
| X | 5101304 | 2 | 18 | 8 | 0 |
| X | 5103133 | 2 | 15 | 15 | 0 |
| X | 5109228 | 2 | 4 | 9 | 0 |
| X | 5110329 | 2 | 16 | 6 | 0 |
| X | 5110343 | 2 | 17 | 6 | 0 |
| X | 5110911 | 2 | 11 | 16 | 0 |
| X | 5110920 | 2 | 8 | 17 | 0 |
| X | 5111192 | 2 | 21 | 11 | 0 |
| X | 5111376 | 2 | 25 | 15 | 0 |
| X | 5114751 | 2 | 16 | 15 | 0 |
| X | 5116099 | 2 | 5 | 6 | 0 |
| X | 5116910 | 2 | 10 | 7 | 0 |
| X | 5117227 | 2 | 10 | 19 | 0 |
| X | 5117274 | 2 | 9 | 16 | 0 |
| X | 5119409 | 2 | 11 | 14 | 0 |
| X | 5123520 | 2 | 5 | 14 | 0 |
| X | 5130418 | 2 | 21 | 5 | 0 |
| X | 5130455 | 2 | 20 | 4 | 0 |
| X | 5139452 | 2 | 13 | 9 | 0 |
| X | 5140268 | 2 | 22 | 8 | 0 |
| X | 5145254 | 2 | 5 | 26 | 0 |
| X | 5148120 | 2 | 4 | 21 | 0 |
| X | 5151720 | 2 | 15 | 29 | 0 |
| X | 5153106 | 2 | 8 | 11 | 0 |
| X | 5153873 | 2 | 8 | 7 | 0 |
| X | 5154001 | 2 | 8 | 22 | 0 |
| X | 5154676 | 2 | 21 | 12 | 0 |
| X | 5155639 | 2 | 8 | 20 | 0 |
| X | 5157498 | 2 | 5 | 17 | 0 |
| X | 5157536 | 2 | 17 | 17 | 0 |
| X | 5166308 | 2 | 9 | 11 | 0 |
| X | 5171152 | 2 | 11 | 8 | 0 |
| X | 5173349 | 2 | 7 | 5 | 0 |
| X | 5175694 | 2 | 11 | 8 | 0 |
| X | 5177507 | 2 | 9 | 19 | 0 |
| X | 5180424 | 2 | 22 | 9 | 0 |
| X | 5184486 | 2 | 12 | 5 | 0 |
| X | 5187403 | 2 | 5 | 5 | 0 |
| X | 5191868 | 2 | 23 | 14 | 0 |
| X | 5192851 | 2 | 12 | 13 | 0 |
| X | 5193197 | 2 | 19 | 13 | 0 |
| X | 5193967 | 2 | 10 | 4 | 0 |
| X | 5194753 | 2 | 11 | 6 | 0 |
| X | 5195309 | 2 | 14 | 9 | 0 |
| X | 5195372 | 2 | 18 | 15 | 0 |
| X | 5196070 | 2 | 10 | 7 | 0 |
| X | 5196711 | 2 | 19 | 15 | 0 |
| X | 5198254 | 2 | 18 | 13 | 0 |
| X | 5199434 | 2 | 8 | 7 | 0 |
| X | 5199533 | 2 | 6 | 20 | 0 |
| X | 5202611 | 2 | 20 | 7 | 0 |
| X | 5202933 | 2 | 21 | 12 | 0 |
| X | 5203849 | 2 | 12 | 4 | 0 |
| X | 5204222 | 2 | 15 | 19 | 0 |
| X | 5208831 | 2 | 19 | 7 | 0 |
| X | 5213595 | 2 | 16 | 8 | 0 |
| X | 5213689 | 2 | 19 | 6 | 0 |
| X | 5213783 | 2 | 26 | 5 | 0 |
| X | 5214770 | 2 | 21 | 5 | 0 |
| X | 5214856 | 2 | 14 | 4 | 0 |
| X | 5215067 | 2 | 10 | 19 | 0 |
| X | 5219154 | 2 | 13 | 4 | 0 |
| X | 5219762 | 2 | 4 | 12 | 0 |
| X | 5219978 | 2 | 10 | 5 | 0 |
| X | 5220566 | 2 | 13 | 8 | 0 |
| X | 5220947 | 2 | 25 | 24 | 0 |
| X | 5221846 | 2 | 13 | 11 | 0 |
| X | 5222015 | 2 | 17 | 4 | 0 |
| X | 5222349 | 2 | 5 | 7 | 0 |
| X | 5222730 | 2 | 4 | 10 | 0 |
| X | 5224420 | 2 | 15 | 4 | 0 |
| X | 5225035 | 2 | 16 | 7 | 0 |
| X | 5225274 | 2 | 15 | 11 | 0 |
| X | 5225516 | 2 | 24 | 36 | 0 |
| X | 5226990 | 2 | 13 | 14 | 0 |
| X | 5227427 | 2 | 32 | 21 | 0 |
| X | 5228054 | 2 | 27 | 14 | 0 |
| X | 5228646 | 2 | 4 | 12 | 0 |
| X | 5229008 | 2 | 19 | 8 | 0 |
| X | 5230324 | 2 | 14 | 4 | 0 |
| X | 5234717 | 2 | 5 | 26 | 0 |
| X | 5235843 | 2 | 9 | 4 | 0 |
| X | 5236156 | 2 | 4 | 22 | 0 |
| X | 5238185 | 2 | 16 | 6 | 0 |
| X | 5238335 | 2 | 11 | 15 | 0 |
| X | 5239695 | 2 | 18 | 4 | 0 |
| X | 5239739 | 2 | 14 | 6 | 0 |
| X | 5240211 | 2 | 14 | 10 | 0 |
| X | 5240564 | 2 | 19 | 7 | 0 |
| X | 5242562 | 2 | 9 | 6 | 0 |
| X | 5243178 | 2 | 8 | 26 | 0 |
| X | 5245792 | 2 | 12 | 7 | 0 |
| X | 5246236 | 2 | 15 | 4 | 0 |
| X | 5246266 | 2 | 14 | 15 | 0 |
| X | 5246515 | 2 | 10 | 4 | 0 |
| X | 5249309 | 2 | 17 | 23 | 0 |
| X | 5253613 | 2 | 14 | 8 | 0 |
| X | 5254326 | 2 | 8 | 19 | 0 |
| X | 5254462 | 2 | 6 | 12 | 0 |
| X | 5257559 | 2 | 18 | 7 | 0 |
| X | 5257871 | 2 | 19 | 16 | 0 |
| X | 5258192 | 2 | 11 | 9 | 0 |
| X | 5258697 | 2 | 4 | 20 | 0 |
| X | 5258976 | 2 | 13 | 8 | 0 |
| X | 5260325 | 2 | 13 | 10 | 0 |
| X | 5260871 | 2 | 12 | 6 | 0 |
| X | 5261430 | 2 | 18 | 10 | 0 |
| X | 5263427 | 2 | 20 | 4 | 0 |
| X | 5278713 | 2 | 11 | 5 | 0 |
| X | 5279442 | 2 | 8 | 9 | 0 |
| X | 5288403 | 2 | 18 | 4 | 0 |
| X | 5290321 | 2 | 21 | 4 | 0 |
| X | 5290360 | 2 | 10 | 7 | 0 |
| X | 5297073 | 2 | 14 | 11 | 0 |
| X | 5299887 | 2 | 4 | 20 | 0 |
| X | 5300887 | 2 | 11 | 30 | 0 |
| X | 5302265 | 2 | 5 | 8 | 0 |
| X | 5304066 | 2 | 29 | 17 | 0 |
| X | 5304207 | 2 | 5 | 16 | 0 |
| X | 5305382 | 2 | 6 | 18 | 0 |
| X | 5305577 | 2 | 5 | 8 | 0 |
| X | 5305928 | 2 | 22 | 14 | 0 |
| X | 5317415 | 2 | 5 | 16 | 0 |
| X | 5321102 | 2 | 11 | 19 | 0 |
| X | 5321430 | 2 | 20 | 13 | 0 |
| X | 5321473 | 2 | 15 | 14 | 0 |
| X | 5321656 | 2 | 6 | 11 | 0 |
| X | 5323801 | 2 | 5 | 5 | 0 |
| X | 5325840 | 2 | 9 | 22 | 0 |
| X | 5328195 | 2 | 7 | 7 | 0 |
| X | 5329901 | 2 | 12 | 5 | 0 |
| X | 5331343 | 2 | 6 | 14 | 0 |
| X | 5331407 | 2 | 12 | 4 | 0 |
| X | 5332057 | 2 | 25 | 9 | 0 |
| X | 5333270 | 2 | 4 | 15 | 0 |
| X | 5333391 | 2 | 11 | 6 | 0 |
| X | 5334998 | 2 | 17 | 7 | 0 |
| X | 5335946 | 2 | 6 | 16 | 0 |
| X | 5340248 | 2 | 17 | 13 | 0 |
| X | 5344331 | 2 | 9 | 4 | 0 |
| X | 5346597 | 2 | 3 | 31 | 0 |
| X | 5347485 | 2 | 7 | 17 | 0 |
| X | 5347876 | 2 | 17 | 11 | 0 |
| X | 5349881 | 2 | 5 | 13 | 0 |
| X | 5350781 | 2 | 9 | 9 | 0 |
| X | 5351083 | 2 | 11 | 25 | 0 |
| X | 5351095 | 2 | 9 | 25 | 0 |
| X | 5351939 | 2 | 7 | 8 | 0 |
| X | 5353315 | 2 | 5 | 6 | 0 |
| X | 5353401 | 2 | 9 | 7 | 0 |
| X | 5355904 | 2 | 4 | 9 | 0 |
| X | 5356318 | 2 | 5 | 14 | 0 |
| X | 5358532 | 2 | 8 | 12 | 0 |
| X | 5358637 | 2 | 15 | 8 | 0 |
| X | 5359010 | 2 | 25 | 11 | 0 |
| X | 5359336 | 2 | 10 | 14 | 0 |
| X | 5360651 | 2 | 21 | 16 | 0 |
| X | 5361250 | 2 | 4 | 17 | 0 |
| X | 5361726 | 2 | 6 | 6 | 0 |
| X | 5362172 | 2 | 9 | 10 | 0 |
| X | 5362623 | 2 | 6 | 12 | 0 |
| X | 5364227 | 2 | 23 | 13 | 0 |
| X | 5364874 | 2 | 15 | 8 | 0 |
| X | 5364958 | 2 | 11 | 7 | 0 |
| X | 5366385 | 2 | 15 | 4 | 0 |
| X | 5367764 | 2 | 24 | 8 | 0 |
| X | 5367789 | 2 | 26 | 7 | 0 |
| X | 5368063 | 2 | 10 | 4 | 0 |
| X | 5369623 | 2 | 30 | 12 | 0 |
| X | 5369928 | 2 | 7 | 8 | 0 |
| X | 5370622 | 2 | 14 | 6 | 0 |
| X | 5373637 | 2 | 11 | 6 | 0 |
| X | 5374453 | 2 | 7 | 7 | 0 |
| X | 5374616 | 2 | 9 | 32 | 0 |
| X | 5374750 | 2 | 12 | 5 | 0 |
| X | 5379645 | 2 | 4 | 12 | 0 |
| X | 5384027 | 2 | 4 | 23 | 0 |
| X | 5387824 | 2 | 11 | 28 | 0 |
| X | 5387923 | 2 | 7 | 29 | 0 |
| X | 5388881 | 2 | 10 | 9 | 0 |
| X | 5388885 | 2 | 9 | 12 | 0 |
| X | 5389160 | 2 | 12 | 26 | 0 |
| X | 5389688 | 2 | 15 | 9 | 0 |
| X | 5389932 | 2 | 12 | 8 | 0 |
| X | 5393917 | 2 | 28 | 14 | 0 |
| X | 5393966 | 2 | 19 | 18 | 0 |
| X | 5393977 | 2 | 14 | 21 | 0 |
| X | 5396401 | 2 | 14 | 5 | 0 |
| X | 5399011 | 2 | 11 | 11 | 0 |
| X | 5403237 | 2 | 13 | 15 | 0 |
| X | 5404242 | 2 | 10 | 7 | 0 |
| X | 5405229 | 2 | 13 | 5 | 0 |
| X | 5405601 | 2 | 10 | 5 | 0 |
| X | 5407045 | 2 | 18 | 9 | 0 |
| X | 5407202 | 2 | 16 | 5 | 0 |
| X | 5407263 | 2 | 6 | 8 | 0 |
| X | 5407285 | 2 | 17 | 9 | 0 |
| X | 5408214 | 2 | 14 | 22 | 0 |
| X | 5424670 | 2 | 20 | 16 | 0 |
| X | 5425087 | 2 | 7 | 16 | 0 |
| X | 5425231 | 2 | 4 | 14 | 0 |
| X | 5426393 | 2 | 8 | 9 | 0 |
| X | 5427463 | 2 | 28 | 5 | 0 |
| X | 5429721 | 2 | 11 | 4 | 0 |
| X | 5430082 | 2 | 13 | 19 | 0 |
| X | 5432239 | 2 | 10 | 9 | 0 |
| X | 5433293 | 2 | 6 | 17 | 0 |
| X | 5436033 | 2 | 6 | 11 | 0 |
| X | 5436562 | 2 | 17 | 11 | 0 |
| X | 5437356 | 2 | 7 | 20 | 0 |
| X | 5437475 | 2 | 17 | 19 | 0 |
| X | 5440743 | 2 | 6 | 13 | 0 |
| X | 5441216 | 2 | 7 | 6 | 0 |
| X | 5442777 | 2 | 8 | 12 | 0 |
| X | 5443393 | 2 | 6 | 10 | 0 |
| X | 5443743 | 2 | 19 | 20 | 0 |
| X | 5443782 | 2 | 4 | 24 | 0 |
| X | 5444182 | 2 | 12 | 12 | 0 |
| X | 5446282 | 2 | 25 | 17 | 0 |
| X | 5446886 | 2 | 8 | 8 | 0 |
| X | 5448058 | 2 | 18 | 10 | 0 |
| X | 5448291 | 2 | 5 | 15 | 0 |
| X | 5449042 | 2 | 15 | 9 | 0 |
| X | 5449276 | 2 | 12 | 18 | 0 |
| X | 5451550 | 2 | 21 | 13 | 0 |
| X | 5453092 | 2 | 12 | 16 | 0 |
| X | 5454152 | 2 | 13 | 4 | 0 |
| X | 5454820 | 2 | 11 | 25 | 0 |
| X | 5455449 | 2 | 12 | 4 | 0 |
| X | 5455705 | 2 | 14 | 18 | 0 |
| X | 5458931 | 2 | 13 | 5 | 0 |
| X | 5460308 | 2 | 4 | 11 | 0 |
| X | 5460916 | 2 | 14 | 4 | 0 |
| X | 5464309 | 2 | 10 | 5 | 0 |
| X | 5464386 | 2 | 7 | 9 | 0 |
| X | 5467731 | 2 | 4 | 18 | 0 |
| X | 5467811 | 2 | 5 | 19 | 0 |
| X | 5467912 | 2 | 4 | 9 | 0 |
| X | 5469879 | 2 | 8 | 10 | 0 |
| X | 5470476 | 2 | 12 | 6 | 0 |
| X | 5470669 | 2 | 8 | 8 | 0 |
| X | 5470909 | 2 | 8 | 5 | 0 |
| X | 5472752 | 2 | 30 | 16 | 0 |
| X | 5473552 | 2 | 16 | 23 | 0 |
| X | 5473714 | 2 | 12 | 11 | 0 |
| X | 5478022 | 2 | 8 | 7 | 0 |
| X | 5478266 | 2 | 6 | 18 | 0 |
| X | 5482809 | 2 | 5 | 17 | 0 |
| X | 5483839 | 2 | 12 | 4 | 0 |
| X | 5484309 | 2 | 11 | 5 | 0 |
| X | 5484391 | 2 | 17 | 7 | 0 |
| X | 5486381 | 2 | 5 | 6 | 0 |
| X | 5487232 | 2 | 8 | 13 | 0 |
| X | 5487286 | 2 | 6 | 15 | 0 |
| X | 5488175 | 2 | 8 | 5 | 0 |
| X | 5489730 | 2 | 7 | 15 | 0 |
| X | 5490353 | 2 | 5 | 7 | 0 |
| X | 5490456 | 2 | 13 | 13 | 0 |
| X | 5490556 | 2 | 20 | 16 | 0 |
| X | 5493329 | 2 | 27 | 10 | 0 |
| X | 5496494 | 2 | 14 | 4 | 0 |
| X | 5508297 | 2 | 17 | 4 | 0 |
| X | 5508669 | 2 | 5 | 6 | 0 |
| X | 5511053 | 2 | 9 | 6 | 0 |
| X | 5512137 | 2 | 25 | 10 | 0 |
| X | 5516061 | 2 | 26 | 15 | 0 |
| X | 5518663 | 2 | 10 | 35 | 0 |
| X | 5518762 | 2 | 12 | 9 | 0 |
| X | 5518864 | 2 | 5 | 11 | 0 |
| X | 5521044 | 2 | 12 | 37 | 0 |
| X | 5521545 | 2 | 6 | 20 | 0 |
| X | 5522538 | 2 | 17 | 5 | 0 |
| X | 5522954 | 2 | 5 | 10 | 0 |
| X | 5523343 | 2 | 19 | 34 | 0 |
| X | 5523941 | 2 | 5 | 28 | 0 |
| X | 5523999 | 2 | 5 | 17 | 0 |
| X | 5524632 | 2 | 13 | 18 | 0 |
| X | 5526703 | 2 | 12 | 4 | 0 |
| X | 5527662 | 2 | 5 | 8 | 0 |
| X | 5527896 | 2 | 6 | 22 | 0 |
| X | 5528728 | 2 | 17 | 5 | 0 |
| X | 5528769 | 2 | 21 | 4 | 0 |
| X | 5530181 | 2 | 14 | 22 | 0 |
| X | 5531402 | 2 | 11 | 27 | 0 |
| X | 5532992 | 2 | 5 | 16 | 0 |
| X | 5534414 | 2 | 16 | 9 | 0 |
| X | 5535222 | 2 | 4 | 20 | 0 |
| X | 5535453 | 2 | 11 | 24 | 0 |
| X | 5535512 | 2 | 12 | 23 | 0 |
| X | 5535660 | 2 | 10 | 30 | 0 |
| X | 5535738 | 2 | 10 | 13 | 0 |
| X | 5536506 | 2 | 5 | 19 | 0 |
| X | 5536823 | 2 | 15 | 10 | 0 |
| X | 5537115 | 2 | 5 | 14 | 0 |
| X | 5539174 | 2 | 8 | 8 | 0 |
| X | 5539253 | 2 | 8 | 22 | 0 |
| X | 5539390 | 2 | 5 | 15 | 0 |
| X | 5539428 | 2 | 9 | 25 | 0 |
| X | 5546524 | 2 | 15 | 18 | 0 |
| X | 5546944 | 2 | 12 | 4 | 0 |
| X | 5548937 | 2 | 18 | 18 | 0 |
| X | 5549060 | 2 | 15 | 8 | 0 |
| X | 5549562 | 2 | 28 | 17 | 0 |
| X | 5549650 | 2 | 16 | 15 | 0 |
| X | 5550130 | 2 | 13 | 5 | 0 |
| X | 5550241 | 2 | 12 | 6 | 0 |
| X | 5552624 | 2 | 34 | 3 | 0 |
| X | 5552734 | 2 | 26 | 14 | 0 |
| X | 5553016 | 2 | 14 | 9 | 0 |
| X | 5553373 | 2 | 7 | 13 | 0 |
| X | 5553815 | 2 | 7 | 13 | 0 |
| X | 5554973 | 2 | 18 | 14 | 0 |
| X | 5556124 | 2 | 8 | 9 | 0 |
| X | 5557195 | 2 | 12 | 23 | 0 |
| X | 5557350 | 2 | 6 | 15 | 0 |
| X | 5558237 | 2 | 5 | 14 | 0 |
| X | 5558588 | 2 | 4 | 16 | 0 |
| X | 5559011 | 2 | 10 | 10 | 0 |
| X | 5563355 | 2 | 13 | 15 | 0 |
| X | 5565162 | 2 | 28 | 17 | 0 |
| X | 5565195 | 2 | 20 | 26 | 0 |
| X | 5565571 | 2 | 7 | 27 | 0 |
| X | 5565906 | 2 | 6 | 12 | 0 |
| X | 5570153 | 2 | 13 | 4 | 0 |
| X | 5570457 | 2 | 5 | 10 | 0 |
| X | 5570517 | 2 | 13 | 13 | 0 |
| X | 5570557 | 2 | 9 | 20 | 0 |
| X | 5570569 | 2 | 12 | 21 | 0 |
| X | 5570659 | 2 | 5 | 6 | 0 |
| X | 5570863 | 2 | 32 | 9 | 0 |
| X | 5570935 | 2 | 23 | 8 | 0 |
| X | 5570984 | 2 | 7 | 6 | 0 |
| X | 5571097 | 2 | 11 | 5 | 0 |
| X | 5571299 | 2 | 17 | 12 | 0 |
| X | 5571880 | 2 | 13 | 28 | 0 |
| X | 5572426 | 2 | 12 | 4 | 0 |
| X | 5574799 | 2 | 20 | 7 | 0 |
| X | 5575696 | 2 | 35 | 11 | 0 |
| X | 5575986 | 2 | 16 | 17 | 0 |
| X | 5576069 | 2 | 16 | 20 | 0 |
| X | 5576816 | 2 | 30 | 21 | 0 |
| X | 5578049 | 2 | 13 | 6 | 0 |
| X | 5578177 | 2 | 23 | 21 | 0 |
| X | 5579407 | 2 | 11 | 13 | 0 |
| X | 5579952 | 2 | 4 | 22 | 0 |
| X | 5580182 | 2 | 5 | 9 | 0 |
| X | 5580790 | 2 | 19 | 6 | 0 |
| X | 5581555 | 2 | 10 | 18 | 0 |
| X | 5583221 | 2 | 13 | 13 | 0 |
| X | 5585576 | 2 | 5 | 18 | 0 |
| X | 5586205 | 2 | 17 | 35 | 0 |
| X | 5586349 | 2 | 16 | 13 | 0 |
| X | 5588120 | 2 | 14 | 6 | 0 |
| X | 5588192 | 2 | 24 | 12 | 0 |
| X | 5588504 | 2 | 14 | 6 | 0 |
| X | 5588570 | 2 | 21 | 22 | 0 |
| X | 5588757 | 2 | 13 | 7 | 0 |
| X | 5593044 | 2 | 5 | 8 | 0 |
| X | 5594000 | 2 | 8 | 5 | 0 |
| X | 5596754 | 2 | 10 | 21 | 0 |
| X | 5597490 | 2 | 7 | 10 | 0 |
| X | 5597665 | 2 | 16 | 22 | 0 |
| X | 5599063 | 2 | 9 | 17 | 0 |
| X | 5601995 | 2 | 12 | 18 | 0 |
| X | 5603594 | 2 | 11 | 15 | 0 |
| X | 5603789 | 2 | 8 | 7 | 0 |
| X | 5607727 | 2 | 17 | 7 | 0 |
| X | 5608795 | 2 | 6 | 18 | 0 |
| X | 5609107 | 2 | 12 | 5 | 0 |
| X | 5609376 | 2 | 16 | 8 | 0 |
| X | 5613557 | 2 | 15 | 9 | 0 |
| X | 5614337 | 2 | 11 | 9 | 0 |
| X | 5614342 | 2 | 12 | 11 | 0 |
| X | 5614549 | 2 | 18 | 4 | 0 |
| X | 5615630 | 2 | 26 | 10 | 0 |
| X | 5616347 | 2 | 10 | 10 | 0 |
| X | 5616434 | 2 | 12 | 4 | 0 |
| X | 5618524 | 2 | 15 | 12 | 0 |
| X | 5618629 | 2 | 10 | 7 | 0 |
| X | 5619924 | 2 | 11 | 14 | 0 |
| X | 5620845 | 2 | 13 | 6 | 0 |
| X | 5627032 | 2 | 11 | 9 | 0 |
| X | 5627107 | 2 | 16 | 5 | 0 |
| X | 5627239 | 2 | 18 | 8 | 0 |
| X | 5628473 | 2 | 5 | 11 | 0 |
| X | 5629429 | 2 | 16 | 9 | 0 |
| X | 5629580 | 2 | 25 | 4 | 0 |
| X | 5632252 | 2 | 12 | 5 | 0 |
| X | 5633055 | 2 | 13 | 6 | 0 |
| X | 5646317 | 2 | 18 | 10 | 0 |
| X | 5646442 | 2 | 25 | 4 | 0 |
| X | 5646693 | 2 | 11 | 4 | 0 |
| X | 5650919 | 2 | 13 | 6 | 0 |
| X | 5653155 | 2 | 18 | 17 | 0 |
| X | 5653547 | 2 | 16 | 11 | 0 |
| X | 5653894 | 2 | 7 | 16 | 0 |
| X | 5654343 | 2 | 16 | 15 | 0 |
| X | 5655751 | 2 | 17 | 4 | 0 |
| X | 5656033 | 2 | 13 | 6 | 0 |
| X | 5661340 | 2 | 32 | 21 | 0 |
| X | 5661524 | 2 | 23 | 15 | 0 |
| X | 5665905 | 2 | 8 | 15 | 0 |
| X | 5666875 | 2 | 30 | 13 | 0 |
| X | 5669091 | 2 | 6 | 11 | 0 |
| X | 5670273 | 2 | 12 | 13 | 0 |
| X | 5670490 | 2 | 13 | 13 | 0 |
| X | 5671757 | 2 | 5 | 12 | 0 |
| X | 5673356 | 2 | 9 | 9 | 0 |
| X | 5679990 | 2 | 11 | 8 | 0 |
| X | 5688207 | 2 | 42 | 10 | 0 |
| X | 5690823 | 2 | 9 | 11 | 0 |
| X | 5693170 | 2 | 19 | 8 | 0 |
| X | 5693481 | 2 | 21 | 10 | 0 |
| X | 5694650 | 2 | 29 | 5 | 0 |
| X | 5696439 | 2 | 17 | 8 | 0 |
| X | 5697955 | 2 | 13 | 6 | 0 |
| X | 5700584 | 2 | 13 | 22 | 0 |
| X | 5700653 | 2 | 9 | 13 | 0 |
| X | 5702342 | 2 | 27 | 13 | 0 |
| X | 5702660 | 2 | 10 | 19 | 0 |
| X | 5702778 | 2 | 7 | 10 | 0 |
| X | 5703221 | 2 | 6 | 23 | 0 |
| X | 5703248 | 2 | 11 | 31 | 0 |
| X | 5703508 | 2 | 16 | 19 | 0 |
| X | 5709316 | 2 | 6 | 13 | 0 |
| X | 5709736 | 2 | 7 | 21 | 0 |
| X | 5709829 | 2 | 6 | 5 | 0 |
| X | 5709889 | 2 | 13 | 29 | 0 |
| X | 5710066 | 2 | 5 | 19 | 0 |
| X | 5710567 | 2 | 13 | 5 | 0 |
| X | 5710688 | 2 | 6 | 8 | 0 |
| X | 5710783 | 2 | 19 | 10 | 0 |
| X | 5713700 | 2 | 14 | 7 | 0 |
| X | 5714393 | 2 | 10 | 10 | 0 |
| X | 5714993 | 2 | 5 | 18 | 0 |
| X | 5715556 | 2 | 4 | 11 | 0 |
| X | 5716171 | 2 | 12 | 6 | 0 |
| X | 5716480 | 2 | 5 | 15 | 0 |
| X | 5716933 | 2 | 10 | 13 | 0 |
| X | 5717154 | 2 | 8 | 24 | 0 |
| X | 5718460 | 2 | 6 | 9 | 0 |
| X | 5719552 | 2 | 17 | 5 | 0 |
| X | 5720883 | 2 | 12 | 9 | 0 |
| X | 5721290 | 2 | 6 | 26 | 0 |
| X | 5722257 | 2 | 5 | 10 | 0 |
| X | 5722826 | 2 | 20 | 33 | 0 |
| X | 5724450 | 2 | 8 | 19 | 0 |
| X | 5724634 | 2 | 9 | 26 | 0 |
| X | 5726906 | 2 | 4 | 14 | 0 |
| X | 5727198 | 2 | 4 | 27 | 0 |
| X | 5729317 | 2 | 12 | 23 | 0 |
| X | 5729979 | 2 | 14 | 14 | 0 |
| X | 5730229 | 2 | 30 | 16 | 0 |
| X | 5730439 | 2 | 20 | 12 | 0 |
| X | 5731301 | 2 | 4 | 12 | 0 |
| X | 5731634 | 2 | 12 | 21 | 0 |
| X | 5731697 | 2 | 8 | 18 | 0 |
| X | 5731877 | 2 | 9 | 8 | 0 |
| X | 5731951 | 2 | 24 | 10 | 0 |
| X | 5732012 | 2 | 7 | 10 | 0 |
| X | 5732628 | 2 | 27 | 10 | 0 |
| X | 5732650 | 2 | 20 | 6 | 0 |
| X | 5732695 | 2 | 19 | 12 | 0 |
| X | 5733972 | 2 | 30 | 10 | 0 |
| X | 5735565 | 2 | 5 | 28 | 0 |
| X | 5737213 | 2 | 13 | 8 | 0 |
| X | 5738002 | 2 | 4 | 13 | 0 |
| X | 5738101 | 2 | 7 | 17 | 0 |
| X | 5739146 | 2 | 11 | 4 | 0 |
| X | 5739627 | 2 | 10 | 14 | 0 |
| X | 5740098 | 2 | 13 | 4 | 0 |
| X | 5741703 | 2 | 4 | 11 | 0 |
| X | 5741752 | 2 | 8 | 14 | 0 |
| X | 5744161 | 2 | 5 | 26 | 0 |
| X | 5744418 | 2 | 24 | 11 | 0 |
| X | 5745235 | 2 | 15 | 12 | 0 |
| X | 5746539 | 2 | 5 | 20 | 0 |
| X | 5747770 | 2 | 4 | 15 | 0 |
| X | 5748092 | 2 | 20 | 14 | 0 |
| X | 5749025 | 2 | 18 | 8 | 0 |
| X | 5749742 | 2 | 13 | 7 | 0 |
| X | 5749940 | 2 | 15 | 5 | 0 |
| X | 5750650 | 2 | 6 | 8 | 0 |
| X | 5751204 | 2 | 13 | 5 | 0 |
| X | 5751342 | 2 | 6 | 9 | 0 |
| X | 5751782 | 2 | 15 | 17 | 0 |
| X | 5752526 | 2 | 4 | 13 | 0 |
| X | 5753553 | 2 | 12 | 13 | 0 |
| X | 5753659 | 2 | 20 | 4 | 0 |
| X | 5754375 | 2 | 18 | 24 | 0 |
| X | 5754418 | 2 | 11 | 30 | 0 |
| X | 5754457 | 2 | 15 | 32 | 0 |
| X | 5754870 | 2 | 14 | 9 | 0 |
| X | 5756222 | 2 | 4 | 12 | 0 |
| X | 5756891 | 2 | 5 | 9 | 0 |
| X | 5757506 | 2 | 12 | 10 | 0 |
| X | 5757697 | 2 | 12 | 14 | 0 |
| X | 5757815 | 2 | 6 | 10 | 0 |
| X | 5757985 | 2 | 7 | 12 | 0 |
| X | 5760072 | 2 | 7 | 20 | 0 |
| X | 5761705 | 2 | 4 | 24 | 0 |
| X | 5762134 | 2 | 5 | 10 | 0 |
| X | 5765079 | 2 | 19 | 21 | 0 |
| X | 5765670 | 2 | 5 | 17 | 0 |
| X | 5765718 | 2 | 5 | 14 | 0 |
| X | 5765964 | 2 | 7 | 17 | 0 |
| X | 5766018 | 2 | 5 | 30 | 0 |
| X | 5766338 | 2 | 7 | 10 | 0 |
| X | 5767221 | 2 | 6 | 12 | 0 |
| X | 5776750 | 2 | 5 | 8 | 0 |
| X | 5780749 | 2 | 10 | 31 | 0 |
| X | 5784237 | 2 | 17 | 11 | 0 |
| X | 5789323 | 2 | 13 | 13 | 0 |
| X | 5790482 | 2 | 20 | 20 | 0 |
| X | 5792100 | 2 | 18 | 19 | 0 |
| X | 5793779 | 2 | 16 | 16 | 0 |
| X | 5797696 | 2 | 4 | 13 | 0 |
| X | 5797892 | 2 | 19 | 18 | 0 |
| X | 5799188 | 2 | 7 | 11 | 0 |
| X | 5799461 | 2 | 7 | 19 | 0 |
| X | 5799528 | 2 | 4 | 17 | 0 |
| X | 5799751 | 2 | 17 | 29 | 0 |
| X | 5799860 | 2 | 5 | 8 | 0 |
| X | 5800501 | 2 | 4 | 21 | 0 |
| X | 5800928 | 2 | 11 | 14 | 0 |
| X | 5801691 | 2 | 5 | 9 | 0 |
| X | 5802819 | 2 | 16 | 6 | 0 |
| X | 5802978 | 2 | 21 | 4 | 0 |
| X | 5803500 | 2 | 22 | 18 | 0 |
| X | 5804644 | 2 | 25 | 14 | 0 |
| X | 5804692 | 2 | 9 | 7 | 0 |
| X | 5804993 | 2 | 10 | 25 | 0 |
| X | 5805155 | 2 | 7 | 24 | 0 |
| X | 5808799 | 2 | 5 | 9 | 0 |
| X | 5809171 | 2 | 10 | 19 | 0 |
| X | 5810079 | 2 | 11 | 6 | 0 |
| X | 5810495 | 2 | 7 | 26 | 0 |
| X | 5815176 | 2 | 5 | 10 | 0 |
| X | 5818723 | 2 | 12 | 5 | 0 |
| X | 5828319 | 2 | 7 | 8 | 0 |
| X | 5829895 | 2 | 6 | 21 | 0 |
| X | 5830441 | 2 | 17 | 9 | 0 |
| X | 5830530 | 2 | 6 | 26 | 0 |
| X | 5830940 | 2 | 7 | 5 | 0 |
| X | 5833987 | 2 | 6 | 11 | 0 |
| X | 5834108 | 2 | 4 | 17 | 0 |
| X | 5842364 | 2 | 17 | 5 | 0 |
| X | 5843671 | 2 | 9 | 17 | 0 |
| X | 5844031 | 2 | 31 | 10 | 0 |
| X | 5846598 | 2 | 19 | 16 | 0 |
| X | 5847187 | 2 | 10 | 11 | 0 |
| X | 5848034 | 2 | 7 | 15 | 0 |
| X | 5848969 | 2 | 8 | 29 | 0 |
| X | 5849067 | 2 | 19 | 16 | 0 |
| X | 5851452 | 2 | 21 | 18 | 0 |
| X | 5851585 | 2 | 28 | 18 | 0 |
| X | 5851702 | 2 | 30 | 20 | 0 |
| X | 5852065 | 2 | 29 | 11 | 0 |
| X | 5852139 | 2 | 9 | 4 | 0 |
| X | 5852398 | 2 | 30 | 18 | 0 |
| X | 5852482 | 2 | 14 | 7 | 0 |
| X | 5852573 | 2 | 26 | 4 | 0 |
| X | 5852671 | 2 | 10 | 16 | 0 |
| X | 5852866 | 2 | 6 | 17 | 0 |
| X | 5852893 | 2 | 9 | 13 | 0 |
| X | 5852956 | 2 | 10 | 9 | 0 |
| X | 5853115 | 2 | 17 | 8 | 0 |
| X | 5853718 | 2 | 21 | 37 | 0 |
| X | 5853955 | 2 | 6 | 11 | 0 |
| X | 5853997 | 2 | 14 | 16 | 0 |
| X | 5854417 | 2 | 4 | 24 | 0 |
| X | 5854445 | 2 | 4 | 29 | 0 |
| X | 5854809 | 2 | 8 | 26 | 0 |
| X | 5854842 | 2 | 14 | 25 | 0 |
| X | 5854980 | 2 | 13 | 13 | 0 |
| X | 5855032 | 2 | 19 | 18 | 0 |
| X | 5855248 | 2 | 8 | 33 | 0 |
| X | 5855323 | 2 | 5 | 13 | 0 |
| X | 5855398 | 2 | 7 | 22 | 0 |
| X | 5855488 | 2 | 11 | 22 | 0 |
| X | 5855639 | 2 | 6 | 9 | 0 |
| X | 5855699 | 2 | 4 | 9 | 0 |
| X | 5856938 | 2 | 5 | 23 | 0 |
| X | 5856975 | 2 | 11 | 25 | 0 |
| X | 5858071 | 2 | 14 | 5 | 0 |
| X | 5858267 | 2 | 11 | 5 | 0 |
| X | 5859380 | 2 | 19 | 21 | 0 |
| X | 5862194 | 2 | 4 | 9 | 0 |
| X | 5862903 | 2 | 6 | 8 | 0 |
| X | 5864086 | 2 | 6 | 20 | 0 |
| X | 5866486 | 2 | 13 | 8 | 0 |
| X | 5867190 | 2 | 34 | 11 | 0 |
| X | 5867527 | 2 | 16 | 8 | 0 |
| X | 5867578 | 2 | 10 | 4 | 0 |
| X | 5869060 | 2 | 18 | 4 | 0 |
| X | 5869442 | 2 | 21 | 6 | 0 |
| X | 5869791 | 2 | 16 | 4 | 0 |
| X | 5869989 | 2 | 16 | 4 | 0 |
| X | 5870258 | 2 | 27 | 4 | 0 |
| X | 5872711 | 2 | 4 | 12 | 0 |
| X | 5876778 | 2 | 13 | 4 | 0 |
| X | 5878203 | 2 | 22 | 13 | 0 |
| X | 5878369 | 2 | 13 | 5 | 0 |
| X | 5878622 | 2 | 14 | 7 | 0 |
| X | 5878694 | 2 | 16 | 7 | 0 |
| X | 5878721 | 2 | 12 | 5 | 0 |
| X | 5879133 | 2 | 11 | 8 | 0 |
| X | 5879270 | 2 | 16 | 5 | 0 |
| X | 5879354 | 2 | 16 | 11 | 0 |
| X | 5880044 | 2 | 28 | 6 | 0 |
| X | 5880212 | 2 | 14 | 6 | 0 |
| X | 5880239 | 2 | 10 | 12 | 0 |
| X | 5880577 | 2 | 10 | 19 | 0 |
| X | 5880594 | 2 | 6 | 11 | 0 |
| X | 5880610 | 2 | 5 | 15 | 0 |
| X | 5882438 | 2 | 20 | 32 | 0 |
| X | 5882845 | 2 | 16 | 16 | 0 |
| X | 5885715 | 2 | 29 | 9 | 0 |
| X | 5886052 | 2 | 27 | 4 | 0 |
| X | 5886133 | 2 | 10 | 4 | 0 |
| X | 5888363 | 2 | 5 | 17 | 0 |
| X | 5888484 | 2 | 5 | 16 | 0 |
| X | 5889724 | 2 | 13 | 12 | 0 |
| X | 5891109 | 2 | 10 | 19 | 0 |
| X | 5894257 | 2 | 10 | 14 | 0 |
| X | 5895081 | 2 | 4 | 19 | 0 |
| X | 5895244 | 2 | 7 | 10 | 0 |
| X | 5897562 | 2 | 9 | 18 | 0 |
| X | 6119956 | 2 | 4 | 19 | 0 |
| X | 6168843 | 2 | 10 | 17 | 0 |
| X | 6465592 | 2 | 18 | 34 | 0 |
| X | 6571844 | 2 | 6 | 14 | 0 |
| X | 6572395 | 2 | 26 | 11 | 0 |
| X | 6575773 | 2 | 17 | 17 | 0 |
| X | 6575800 | 2 | 25 | 9 | 0 |
| X | 6582487 | 2 | 10 | 21 | 0 |
| X | 6583757 | 2 | 13 | 21 | 0 |
| X | 6585289 | 2 | 10 | 4 | 0 |
| X | 6593878 | 2 | 6 | 8 | 0 |
| X | 6597739 | 2 | 13 | 8 | 0 |
| X | 6598318 | 2 | 16 | 10 | 0 |
| X | 6600512 | 2 | 8 | 19 | 0 |
| X | 6601533 | 2 | 8 | 10 | 0 |
| X | 6601767 | 2 | 6 | 19 | 0 |
| X | 6603468 | 2 | 12 | 13 | 0 |
| X | 6604261 | 2 | 13 | 18 | 0 |
| X | 6604946 | 2 | 16 | 5 | 0 |
| X | 6606180 | 2 | 6 | 13 | 0 |
| X | 6606813 | 2 | 13 | 18 | 0 |
| X | 6608733 | 2 | 6 | 26 | 0 |
| X | 6609190 | 2 | 5 | 9 | 0 |
| X | 6610012 | 2 | 8 | 19 | 0 |
| X | 6610574 | 2 | 11 | 9 | 0 |
| X | 6611198 | 2 | 17 | 12 | 0 |
| X | 6611424 | 2 | 9 | 7 | 0 |
| X | 6611973 | 2 | 26 | 12 | 0 |
| X | 6612164 | 2 | 6 | 18 | 0 |
| X | 6612183 | 2 | 5 | 7 | 0 |
| X | 6612418 | 2 | 13 | 6 | 0 |
| X | 6613724 | 2 | 10 | 5 | 0 |
| X | 6613791 | 2 | 8 | 6 | 0 |
| X | 6613900 | 2 | 18 | 9 | 0 |
| X | 6615486 | 2 | 16 | 10 | 0 |
| X | 6618395 | 2 | 11 | 4 | 0 |
| X | 6618940 | 2 | 21 | 8 | 0 |
| X | 6619041 | 2 | 28 | 15 | 0 |
| X | 6619085 | 2 | 21 | 11 | 0 |
| X | 6619362 | 2 | 20 | 23 | 0 |
| X | 6619933 | 2 | 12 | 5 | 0 |
| X | 6620581 | 2 | 10 | 5 | 0 |
| X | 6620645 | 2 | 6 | 16 | 0 |
| X | 6620964 | 2 | 9 | 16 | 0 |
| X | 6621894 | 2 | 4 | 16 | 0 |
| X | 6622969 | 2 | 9 | 16 | 0 |
| X | 6625195 | 2 | 5 | 13 | 0 |
| X | 6625387 | 2 | 6 | 17 | 0 |
| X | 6625440 | 2 | 23 | 7 | 0 |
| X | 6626796 | 2 | 16 | 13 | 0 |
| X | 6626821 | 2 | 5 | 11 | 0 |
| X | 6628403 | 2 | 15 | 5 | 0 |
| X | 6628753 | 2 | 10 | 5 | 0 |
| X | 6628776 | 2 | 13 | 4 | 0 |
| X | 6628918 | 2 | 9 | 30 | 0 |
| X | 6629830 | 2 | 10 | 15 | 0 |
| X | 6630375 | 2 | 10 | 5 | 0 |
| X | 6631402 | 2 | 13 | 4 | 0 |
| X | 6632680 | 2 | 21 | 8 | 0 |
| X | 6635566 | 2 | 4 | 16 | 0 |
| X | 6636974 | 2 | 4 | 14 | 0 |
| X | 6638982 | 2 | 14 | 19 | 0 |
| X | 6639441 | 2 | 16 | 8 | 0 |
| X | 6639606 | 2 | 5 | 5 | 0 |
| X | 6640035 | 2 | 33 | 6 | 0 |
| X | 6640052 | 2 | 26 | 4 | 0 |
| X | 6640218 | 2 | 13 | 14 | 0 |
| X | 6640671 | 2 | 6 | 29 | 0 |
| X | 6644439 | 2 | 7 | 5 | 0 |
| X | 6644566 | 2 | 8 | 12 | 0 |
| X | 6646564 | 2 | 11 | 7 | 0 |
| X | 6647788 | 2 | 11 | 4 | 0 |
| X | 6647844 | 2 | 31 | 18 | 0 |
| X | 6647893 | 2 | 8 | 10 | 0 |
| X | 6648750 | 2 | 6 | 16 | 0 |
| X | 6655602 | 2 | 12 | 13 | 0 |
| X | 6657921 | 2 | 5 | 14 | 0 |
| X | 6658333 | 2 | 18 | 17 | 0 |
| X | 6660590 | 2 | 6 | 9 | 0 |
| X | 6660915 | 2 | 6 | 13 | 0 |
| X | 6661164 | 2 | 28 | 20 | 0 |
| X | 6664242 | 2 | 8 | 7 | 0 |
| X | 6665700 | 2 | 6 | 11 | 0 |
| X | 6665972 | 2 | 14 | 6 | 0 |
| X | 6666218 | 2 | 20 | 13 | 0 |
| X | 6666564 | 2 | 17 | 18 | 0 |
| X | 6666776 | 2 | 19 | 13 | 0 |
| X | 6667122 | 2 | 13 | 5 | 0 |
| X | 6667405 | 2 | 12 | 9 | 0 |
| X | 6668441 | 2 | 12 | 22 | 0 |
| X | 6716548 | 2 | 5 | 12 | 0 |
| X | 6716573 | 2 | 21 | 14 | 0 |
| X | 6716854 | 2 | 11 | 5 | 0 |
| X | 6716926 | 2 | 16 | 9 | 0 |
| X | 6718300 | 2 | 25 | 15 | 0 |
| X | 6720115 | 2 | 17 | 13 | 0 |
| X | 6720996 | 2 | 24 | 9 | 0 |
| X | 6722752 | 2 | 6 | 7 | 0 |
| X | 6722847 | 2 | 13 | 11 | 0 |
| X | 6724352 | 2 | 11 | 23 | 0 |
| X | 6724799 | 2 | 7 | 18 | 0 |
| X | 6725497 | 2 | 3 | 35 | 0 |
| X | 6725519 | 2 | 6 | 32 | 0 |
| X | 6726088 | 2 | 23 | 6 | 0 |
| X | 6726206 | 2 | 14 | 4 | 0 |
| X | 6726230 | 2 | 23 | 17 | 0 |
| X | 6726745 | 2 | 5 | 6 | 0 |
| X | 6726764 | 2 | 6 | 5 | 0 |
| X | 6726779 | 2 | 5 | 12 | 0 |
| X | 6726996 | 2 | 13 | 4 | 0 |
| X | 6727165 | 2 | 24 | 19 | 0 |
| X | 6728529 | 2 | 11 | 15 | 0 |
| X | 6728538 | 2 | 13 | 14 | 0 |
| X | 6728583 | 2 | 20 | 6 | 0 |
| X | 6729387 | 2 | 12 | 5 | 0 |
| X | 6731065 | 2 | 25 | 16 | 0 |
| X | 6731246 | 2 | 15 | 9 | 0 |
| X | 6732178 | 2 | 9 | 17 | 0 |
| X | 6732826 | 2 | 17 | 4 | 0 |
| X | 6734234 | 2 | 11 | 6 | 0 |
| X | 6734256 | 2 | 7 | 20 | 0 |
| X | 6734434 | 2 | 11 | 16 | 0 |
| X | 6734541 | 2 | 17 | 12 | 0 |
| X | 6736540 | 2 | 15 | 17 | 0 |
| X | 6739302 | 2 | 25 | 18 | 0 |
| X | 6739895 | 2 | 19 | 16 | 0 |
| X | 6740501 | 2 | 4 | 16 | 0 |
| X | 6741301 | 2 | 6 | 10 | 0 |
| X | 6744436 | 2 | 10 | 6 | 0 |
| X | 6744691 | 2 | 12 | 26 | 0 |
| X | 6746169 | 2 | 20 | 22 | 0 |
| X | 6752011 | 2 | 9 | 13 | 0 |
| X | 6752719 | 2 | 7 | 5 | 0 |
| X | 6753033 | 2 | 13 | 13 | 0 |
| X | 6753439 | 2 | 4 | 10 | 0 |
| X | 6754217 | 2 | 19 | 16 | 0 |
| X | 6754828 | 2 | 13 | 9 | 0 |
| X | 6754883 | 2 | 18 | 4 | 0 |
| X | 6755668 | 2 | 7 | 19 | 0 |
| X | 6757809 | 2 | 17 | 12 | 0 |
| X | 6758102 | 2 | 13 | 5 | 0 |
| X | 6759456 | 2 | 6 | 14 | 0 |
| X | 6759662 | 2 | 7 | 6 | 0 |
| X | 6760014 | 2 | 9 | 22 | 0 |
| X | 6760115 | 2 | 12 | 25 | 0 |
| X | 6766381 | 2 | 7 | 17 | 0 |
| X | 6770041 | 2 | 21 | 9 | 0 |
| X | 6770464 | 2 | 22 | 9 | 0 |
| X | 6771120 | 2 | 11 | 22 | 0 |
| X | 6771542 | 2 | 13 | 4 | 0 |
| X | 6772157 | 2 | 11 | 28 | 0 |
| X | 6772781 | 2 | 11 | 20 | 0 |
| X | 6772818 | 2 | 14 | 25 | 0 |
| X | 6778360 | 2 | 4 | 13 | 0 |
| X | 6778766 | 2 | 6 | 23 | 0 |
| X | 6780180 | 2 | 9 | 17 | 0 |
| X | 6781399 | 2 | 19 | 8 | 0 |
| X | 6781806 | 2 | 15 | 27 | 0 |
| X | 6784537 | 2 | 5 | 21 | 0 |
| X | 6785098 | 2 | 5 | 8 | 0 |
| X | 6785534 | 2 | 10 | 19 | 0 |
| X | 6785927 | 2 | 4 | 20 | 0 |
| X | 6786133 | 2 | 9 | 18 | 0 |
| X | 6788679 | 2 | 9 | 7 | 0 |
| X | 6788875 | 2 | 9 | 7 | 0 |
| X | 6788901 | 2 | 9 | 17 | 0 |
| X | 6789111 | 2 | 6 | 9 | 0 |
| X | 6789135 | 2 | 5 | 7 | 0 |
| X | 6793518 | 2 | 16 | 12 | 0 |
| X | 6793563 | 2 | 14 | 19 | 0 |
| X | 6793572 | 2 | 14 | 21 | 0 |
| X | 6796698 | 2 | 8 | 9 | 0 |
| X | 6797073 | 2 | 12 | 11 | 0 |
| X | 6800581 | 2 | 16 | 5 | 0 |
| X | 6800665 | 2 | 11 | 31 | 0 |
| X | 6801075 | 2 | 20 | 13 | 0 |
| X | 6804122 | 2 | 4 | 16 | 0 |
| X | 6807227 | 2 | 10 | 4 | 0 |
| X | 6815074 | 2 | 16 | 18 | 0 |
| X | 6820593 | 2 | 6 | 10 | 0 |
| X | 6824904 | 2 | 5 | 18 | 0 |
| X | 6826104 | 2 | 5 | 12 | 0 |
| X | 6826173 | 2 | 5 | 10 | 0 |
| X | 6826302 | 2 | 20 | 12 | 0 |
| X | 6826422 | 2 | 13 | 10 | 0 |
| X | 6828122 | 2 | 5 | 14 | 0 |
| X | 6828567 | 2 | 19 | 10 | 0 |
| X | 6829040 | 2 | 24 | 4 | 0 |
| X | 6829215 | 2 | 5 | 16 | 0 |
| X | 6829441 | 2 | 11 | 12 | 0 |
| X | 6830281 | 2 | 4 | 15 | 0 |
| X | 6830509 | 2 | 13 | 26 | 0 |
| X | 6830567 | 2 | 20 | 33 | 0 |
| X | 6832372 | 2 | 8 | 7 | 0 |
| X | 6833066 | 2 | 9 | 25 | 0 |
| X | 6833116 | 2 | 10 | 23 | 0 |
| X | 6833233 | 2 | 11 | 6 | 0 |
| X | 6834427 | 2 | 11 | 6 | 0 |
| X | 6834690 | 2 | 10 | 8 | 0 |
| X | 6837432 | 2 | 11 | 5 | 0 |
| X | 6837890 | 2 | 11 | 13 | 0 |
| X | 6838967 | 2 | 16 | 33 | 0 |
| X | 6840540 | 2 | 20 | 10 | 0 |
| X | 6841072 | 2 | 5 | 33 | 0 |
| X | 6841134 | 2 | 5 | 15 | 0 |
| X | 6841736 | 2 | 7 | 8 | 0 |
| X | 6852488 | 2 | 26 | 17 | 0 |
| X | 6853144 | 2 | 22 | 15 | 0 |
| X | 6853608 | 2 | 6 | 20 | 0 |
| X | 6853728 | 2 | 10 | 12 | 0 |
| X | 6863823 | 2 | 6 | 17 | 0 |
| X | 6863853 | 2 | 7 | 19 | 0 |
| X | 6864398 | 2 | 16 | 18 | 0 |
| X | 6864953 | 2 | 24 | 15 | 0 |
| X | 6865076 | 2 | 15 | 4 | 0 |
| X | 6865418 | 2 | 13 | 7 | 0 |
| X | 6865436 | 2 | 6 | 14 | 0 |
| X | 6865772 | 2 | 5 | 5 | 0 |
| X | 6865776 | 2 | 6 | 5 | 0 |
| X | 6866150 | 2 | 22 | 13 | 0 |
| X | 6866242 | 2 | 10 | 4 | 0 |
| X | 6866705 | 2 | 19 | 5 | 0 |
| X | 6866894 | 2 | 18 | 7 | 0 |
| X | 6867089 | 2 | 11 | 13 | 0 |
| X | 6867224 | 2 | 18 | 9 | 0 |
| X | 6867471 | 2 | 24 | 7 | 0 |
| X | 6867533 | 2 | 25 | 16 | 0 |
| X | 6872304 | 2 | 21 | 9 | 0 |
| X | 6873307 | 2 | 5 | 9 | 0 |
| X | 6873907 | 2 | 13 | 17 | 0 |
| X | 6873973 | 2 | 10 | 21 | 0 |
| X | 6874380 | 2 | 12 | 25 | 0 |
| X | 6874526 | 2 | 9 | 14 | 0 |
| X | 6874760 | 2 | 6 | 18 | 0 |
| X | 6874849 | 2 | 5 | 11 | 0 |
| X | 6874955 | 2 | 12 | 5 | 0 |
| X | 6875016 | 2 | 5 | 24 | 0 |
| X | 6877702 | 2 | 15 | 8 | 0 |
| X | 6883324 | 2 | 5 | 24 | 0 |
| X | 6884373 | 2 | 13 | 8 | 0 |
| X | 6887814 | 2 | 13 | 12 | 0 |
| X | 6888285 | 2 | 9 | 5 | 0 |
| X | 6888483 | 2 | 17 | 8 | 0 |
| X | 6888697 | 2 | 20 | 11 | 0 |
| X | 6890251 | 2 | 8 | 9 | 0 |
| X | 6890387 | 2 | 24 | 15 | 0 |
| X | 6890480 | 2 | 25 | 18 | 0 |
| X | 6890537 | 2 | 5 | 24 | 0 |
| X | 6891027 | 2 | 13 | 5 | 0 |
| X | 6893009 | 2 | 16 | 7 | 0 |
| X | 6893788 | 2 | 25 | 24 | 0 |
| X | 6893854 | 2 | 23 | 16 | 0 |
| X | 6893933 | 2 | 20 | 9 | 0 |
| X | 6894906 | 2 | 13 | 8 | 0 |
| X | 6895007 | 2 | 18 | 13 | 0 |
| X | 6895275 | 2 | 18 | 6 | 0 |
| X | 6895315 | 2 | 15 | 4 | 0 |
| X | 6899450 | 2 | 7 | 9 | 0 |
| X | 6902884 | 2 | 5 | 11 | 0 |
| X | 6903008 | 2 | 21 | 19 | 0 |
| X | 6903336 | 2 | 10 | 23 | 0 |
| X | 6903753 | 2 | 5 | 12 | 0 |
| X | 6903888 | 2 | 8 | 21 | 0 |
| X | 6903975 | 2 | 8 | 29 | 0 |
| X | 6904077 | 2 | 4 | 19 | 0 |
| X | 6904662 | 2 | 5 | 21 | 0 |
| X | 6905227 | 2 | 5 | 7 | 0 |
| X | 6905734 | 2 | 6 | 14 | 0 |
| X | 6905903 | 2 | 14 | 20 | 0 |
| X | 6906025 | 2 | 13 | 33 | 0 |
| X | 6906193 | 2 | 5 | 16 | 0 |
| X | 6910461 | 2 | 11 | 34 | 0 |
| X | 6915771 | 2 | 7 | 18 | 0 |
| X | 6925482 | 2 | 11 | 12 | 0 |
| X | 6927637 | 2 | 19 | 13 | 0 |
| X | 6927719 | 2 | 6 | 19 | 0 |
| X | 6928155 | 2 | 9 | 5 | 0 |
| X | 6929846 | 2 | 4 | 21 | 0 |
| X | 6930071 | 2 | 12 | 8 | 0 |
| X | 6931024 | 2 | 10 | 16 | 0 |
| X | 6931439 | 2 | 11 | 14 | 0 |
| X | 6933249 | 2 | 28 | 5 | 0 |
| X | 6933861 | 2 | 16 | 9 | 0 |
| X | 6934450 | 2 | 23 | 9 | 0 |
| X | 6934656 | 2 | 28 | 14 | 0 |
| X | 6935235 | 2 | 21 | 8 | 0 |
| X | 6935304 | 2 | 8 | 7 | 0 |
| X | 6938470 | 2 | 33 | 10 | 0 |
| X | 6938528 | 2 | 27 | 7 | 0 |
| X | 6938990 | 2 | 9 | 16 | 0 |
| X | 6940471 | 2 | 17 | 6 | 0 |
| X | 6941907 | 2 | 7 | 23 | 0 |
| X | 6942401 | 2 | 11 | 10 | 0 |
| X | 6943008 | 2 | 8 | 23 | 0 |
| X | 6943215 | 2 | 9 | 18 | 0 |
| X | 6944239 | 2 | 10 | 9 | 0 |
| X | 6945966 | 2 | 8 | 24 | 0 |
| X | 6947039 | 2 | 7 | 7 | 0 |
| X | 6947120 | 2 | 18 | 6 | 0 |
| X | 6947162 | 2 | 17 | 6 | 0 |
| X | 6947199 | 2 | 6 | 7 | 0 |
| X | 6947207 | 2 | 6 | 11 | 0 |
| X | 6947615 | 2 | 11 | 33 | 0 |
| X | 6951380 | 2 | 10 | 4 | 0 |
| X | 6952118 | 2 | 4 | 14 | 0 |
| X | 6956161 | 2 | 12 | 12 | 0 |
| X | 6956979 | 2 | 6 | 21 | 0 |
| X | 6962311 | 2 | 14 | 6 | 0 |
| X | 6962367 | 2 | 16 | 9 | 0 |
| X | 6962395 | 2 | 12 | 21 | 0 |
| X | 6963192 | 2 | 7 | 7 | 0 |
| X | 6964189 | 2 | 19 | 4 | 0 |
| X | 6964611 | 2 | 24 | 10 | 0 |
| X | 6964671 | 2 | 6 | 11 | 0 |
| X | 6964698 | 2 | 6 | 7 | 0 |
| X | 6964975 | 2 | 19 | 11 | 0 |
| X | 6965050 | 2 | 6 | 15 | 0 |
| X | 6965368 | 2 | 6 | 6 | 0 |
| X | 6965601 | 2 | 7 | 18 | 0 |
| X | 6967871 | 2 | 15 | 7 | 0 |
| X | 6967880 | 2 | 17 | 6 | 0 |
| X | 6970406 | 2 | 15 | 20 | 0 |
| X | 6970571 | 2 | 21 | 17 | 0 |
| X | 6972670 | 2 | 5 | 9 | 0 |
| X | 6973869 | 2 | 13 | 21 | 0 |
| X | 6981350 | 2 | 7 | 13 | 0 |
| X | 6983310 | 2 | 9 | 4 | 0 |
| X | 6984954 | 2 | 12 | 12 | 0 |
| X | 6996014 | 2 | 23 | 6 | 0 |
| X | 6997713 | 2 | 7 | 16 | 0 |
| X | 7001469 | 2 | 20 | 21 | 0 |
| X | 7001513 | 2 | 9 | 4 | 0 |
| X | 7001714 | 2 | 16 | 9 | 0 |
| X | 7002175 | 2 | 21 | 6 | 0 |
| X | 7002298 | 2 | 16 | 8 | 0 |
| X | 7002352 | 2 | 25 | 14 | 0 |
| X | 7004221 | 2 | 10 | 7 | 0 |
| X | 7004304 | 2 | 29 | 11 | 0 |
| X | 7004932 | 2 | 9 | 18 | 0 |
| X | 7005159 | 2 | 13 | 6 | 0 |
| X | 7005476 | 2 | 17 | 7 | 0 |
| X | 7007598 | 2 | 17 | 24 | 0 |
| X | 7010343 | 2 | 11 | 6 | 0 |
| X | 7018239 | 2 | 18 | 10 | 0 |
| X | 7031999 | 2 | 7 | 18 | 0 |
| X | 7050579 | 2 | 17 | 9 | 0 |
| X | 7052671 | 2 | 4 | 17 | 0 |
| X | 7055336 | 2 | 10 | 25 | 0 |
| X | 7056301 | 2 | 8 | 13 | 0 |
| X | 7059080 | 2 | 4 | 13 | 0 |
| X | 7059908 | 2 | 4 | 9 | 0 |
| X | 7060366 | 2 | 9 | 16 | 0 |
| X | 7063783 | 2 | 29 | 16 | 0 |
| X | 7063945 | 2 | 17 | 11 | 0 |
| X | 7065346 | 2 | 6 | 18 | 0 |
| X | 7065937 | 2 | 8 | 12 | 0 |
| X | 7068278 | 2 | 9 | 10 | 0 |
| X | 7068476 | 2 | 6 | 12 | 0 |
| X | 7068677 | 2 | 7 | 8 | 0 |
| X | 7068853 | 2 | 19 | 15 | 0 |
| X | 7068918 | 2 | 5 | 13 | 0 |
| X | 7069918 | 2 | 15 | 15 | 0 |
| X | 7070106 | 2 | 4 | 18 | 0 |
| X | 7072132 | 2 | 11 | 21 | 0 |
| X | 7073285 | 2 | 9 | 13 | 0 |
| X | 7073587 | 2 | 4 | 10 | 0 |
| X | 7075574 | 2 | 7 | 5 | 0 |
| X | 7077299 | 2 | 4 | 11 | 0 |
| X | 7079551 | 2 | 14 | 4 | 0 |
| X | 7079893 | 2 | 35 | 4 | 0 |
| X | 7083832 | 2 | 15 | 20 | 0 |
| X | 7085999 | 2 | 17 | 4 | 0 |
| X | 7088611 | 2 | 6 | 11 | 0 |
| X | 7090942 | 2 | 12 | 14 | 0 |
| X | 7091672 | 2 | 6 | 15 | 0 |
| X | 7091723 | 2 | 7 | 16 | 0 |
| X | 7400266 | 2 | 16 | 6 | 0 |
| X | 7400407 | 2 | 6 | 6 | 0 |
| X | 7400689 | 2 | 8 | 5 | 0 |
| X | 7401320 | 2 | 22 | 15 | 0 |
| X | 7404404 | 2 | 13 | 7 | 0 |
| X | 7405239 | 2 | 17 | 7 | 0 |
| X | 7408151 | 2 | 7 | 8 | 0 |
| X | 7408850 | 2 | 7 | 6 | 0 |
| X | 7410686 | 2 | 21 | 14 | 0 |
| X | 7415911 | 2 | 6 | 7 | 0 |
| X | 7416373 | 2 | 16 | 10 | 0 |
| X | 7421878 | 2 | 23 | 5 | 0 |
| X | 7422121 | 2 | 8 | 5 | 0 |
| X | 7423338 | 2 | 13 | 4 | 0 |
| X | 7424965 | 2 | 11 | 11 | 0 |
| X | 7427971 | 2 | 4 | 11 | 0 |
| X | 7428331 | 2 | 15 | 9 | 0 |
| X | 7428786 | 2 | 16 | 11 | 0 |
| X | 7429367 | 2 | 14 | 7 | 0 |
| X | 7429459 | 2 | 21 | 16 | 0 |
| X | 7430029 | 2 | 20 | 12 | 0 |
| X | 7432557 | 2 | 21 | 8 | 0 |
| X | 7432761 | 2 | 14 | 24 | 0 |
| X | 7434329 | 2 | 24 | 11 | 0 |
| X | 7436378 | 2 | 9 | 5 | 0 |
| X | 7437091 | 2 | 13 | 13 | 0 |
| X | 7438477 | 2 | 24 | 10 | 0 |
| X | 7438604 | 2 | 9 | 4 | 0 |
| X | 7439290 | 2 | 9 | 7 | 0 |
| X | 7439410 | 2 | 10 | 6 | 0 |
| X | 7439668 | 2 | 35 | 13 | 0 |
| X | 7440155 | 2 | 14 | 16 | 0 |
| X | 7440242 | 2 | 25 | 8 | 0 |
| X | 7440795 | 2 | 12 | 9 | 0 |
| X | 7440964 | 2 | 11 | 4 | 0 |
| X | 7448243 | 2 | 9 | 5 | 0 |
| X | 7449825 | 2 | 23 | 13 | 0 |
| X | 7453023 | 2 | 25 | 9 | 0 |
| X | 7453285 | 2 | 21 | 7 | 0 |
| X | 7453654 | 2 | 22 | 8 | 0 |
| X | 7454608 | 2 | 13 | 17 | 0 |
| X | 7454677 | 2 | 17 | 6 | 0 |
| X | 7454719 | 2 | 12 | 6 | 0 |
| X | 7454792 | 2 | 23 | 4 | 0 |
| X | 7455424 | 2 | 9 | 5 | 0 |
| X | 7455645 | 2 | 17 | 4 | 0 |
| X | 7457369 | 2 | 12 | 15 | 0 |
| X | 7458538 | 2 | 16 | 5 | 0 |
| X | 7459884 | 2 | 14 | 6 | 0 |
| X | 7463678 | 2 | 23 | 12 | 0 |
| X | 7466270 | 2 | 8 | 8 | 0 |
| X | 7467098 | 2 | 16 | 9 | 0 |
| X | 7467150 | 2 | 12 | 16 | 0 |
| X | 7467295 | 2 | 20 | 14 | 0 |
| X | 7473090 | 2 | 13 | 4 | 0 |
| X | 7476695 | 2 | 14 | 5 | 0 |
| X | 7476972 | 2 | 10 | 4 | 0 |
| X | 7478095 | 2 | 8 | 6 | 0 |
| X | 7479771 | 2 | 11 | 7 | 0 |
| X | 7481332 | 2 | 16 | 13 | 0 |
| X | 7482665 | 2 | 11 | 5 | 0 |
| X | 7484032 | 2 | 19 | 8 | 0 |
| X | 7484437 | 2 | 15 | 4 | 0 |
| X | 7484500 | 2 | 18 | 11 | 0 |
| X | 7484874 | 2 | 9 | 4 | 0 |
| X | 7485603 | 2 | 15 | 8 | 0 |
| X | 7487087 | 2 | 20 | 5 | 0 |
| X | 7487546 | 2 | 14 | 6 | 0 |
| X | 7487598 | 2 | 17 | 18 | 0 |
| X | 7488104 | 2 | 20 | 4 | 0 |
| X | 7489161 | 2 | 9 | 4 | 0 |
| X | 7489772 | 2 | 5 | 5 | 0 |
| X | 7490469 | 2 | 9 | 5 | 0 |
| X | 7490544 | 2 | 17 | 14 | 0 |
| X | 7490828 | 2 | 12 | 15 | 0 |
| X | 7494106 | 2 | 11 | 4 | 0 |
| X | 7495129 | 2 | 8 | 6 | 0 |
| X | 7498869 | 2 | 9 | 6 | 0 |
| X | 7500050 | 2 | 14 | 4 | 0 |
| X | 7500562 | 2 | 20 | 15 | 0 |
| X | 7500650 | 2 | 6 | 5 | 0 |
| X | 7501884 | 2 | 6 | 8 | 0 |
| X | 7502720 | 2 | 5 | 6 | 0 |
| X | 7503399 | 2 | 9 | 14 | 0 |
| X | 7503731 | 2 | 16 | 7 | 0 |
| X | 7507623 | 2 | 21 | 17 | 0 |
| X | 7509481 | 2 | 26 | 12 | 0 |
| X | 7514347 | 2 | 21 | 7 | 0 |
| X | 7514597 | 2 | 14 | 11 | 0 |
| X | 7514923 | 2 | 13 | 6 | 0 |
| X | 7515090 | 2 | 15 | 6 | 0 |
| X | 7515270 | 2 | 23 | 25 | 0 |
| X | 7515808 | 2 | 9 | 8 | 0 |
| X | 7515942 | 2 | 12 | 7 | 0 |
| X | 7517677 | 2 | 11 | 4 | 0 |
| X | 7524038 | 2 | 14 | 7 | 0 |
| X | 7524198 | 2 | 19 | 24 | 0 |
| X | 7526986 | 2 | 12 | 12 | 0 |
| X | 7531574 | 2 | 19 | 7 | 0 |
| X | 7531649 | 2 | 9 | 4 | 0 |
| X | 7543383 | 2 | 9 | 6 | 0 |
| X | 7543449 | 2 | 12 | 6 | 0 |
| X | 7545377 | 2 | 15 | 11 | 0 |
| X | 7545557 | 2 | 10 | 15 | 0 |
| X | 7545694 | 2 | 24 | 12 | 0 |
| X | 7547366 | 2 | 10 | 6 | 0 |
| X | 7547917 | 2 | 8 | 12 | 0 |
| X | 7548337 | 2 | 9 | 5 | 0 |
| X | 7550106 | 2 | 17 | 13 | 0 |
| X | 7551401 | 2 | 19 | 10 | 0 |
| X | 7551772 | 2 | 17 | 4 | 0 |
| X | 7557304 | 2 | 10 | 15 | 0 |
| X | 7559689 | 2 | 7 | 8 | 0 |
| X | 7559870 | 2 | 12 | 5 | 0 |
| X | 7560105 | 2 | 11 | 5 | 0 |
| X | 7560194 | 2 | 7 | 5 | 0 |
| X | 7561480 | 2 | 16 | 7 | 0 |
| X | 7574305 | 2 | 26 | 10 | 0 |
| X | 7577325 | 2 | 6 | 6 | 0 |
| X | 7577896 | 2 | 19 | 6 | 0 |
| X | 7579234 | 2 | 14 | 9 | 0 |
| X | 7579413 | 2 | 7 | 6 | 0 |
| X | 7579547 | 2 | 10 | 9 | 0 |
| X | 7583207 | 2 | 16 | 9 | 0 |
| X | 7583612 | 2 | 18 | 10 | 0 |
| X | 7584011 | 2 | 13 | 9 | 0 |
| X | 7584146 | 2 | 19 | 13 | 0 |
| X | 7584274 | 2 | 9 | 16 | 0 |
| X | 7584339 | 2 | 17 | 11 | 0 |
| X | 7584575 | 2 | 28 | 8 | 0 |
| X | 7585144 | 2 | 17 | 5 | 0 |
| X | 7585306 | 2 | 19 | 5 | 0 |
| X | 7585449 | 2 | 28 | 6 | 0 |
| X | 7585873 | 2 | 14 | 10 | 0 |
| X | 7586534 | 2 | 6 | 7 | 0 |
| X | 7587120 | 2 | 10 | 7 | 0 |
| X | 7587379 | 2 | 24 | 11 | 0 |
| X | 7588311 | 2 | 11 | 4 | 0 |
| X | 7588382 | 2 | 12 | 5 | 0 |
| X | 7589800 | 2 | 6 | 10 | 0 |
| X | 7589900 | 2 | 16 | 7 | 0 |
| X | 7589953 | 2 | 13 | 12 | 0 |
| X | 7590194 | 2 | 10 | 6 | 0 |
| X | 7590308 | 2 | 11 | 6 | 0 |
| X | 7590397 | 2 | 8 | 8 | 0 |
| X | 7590869 | 2 | 5 | 5 | 0 |
| X | 7591156 | 2 | 10 | 7 | 0 |
| X | 7591601 | 2 | 13 | 11 | 0 |
| X | 7592075 | 2 | 9 | 8 | 0 |
| X | 7593152 | 2 | 24 | 6 | 0 |
| X | 7593350 | 2 | 8 | 5 | 0 |
| X | 7593592 | 2 | 12 | 4 | 0 |
| X | 7594258 | 2 | 19 | 6 | 0 |
| X | 7594589 | 2 | 21 | 4 | 0 |
| X | 7594641 | 2 | 15 | 11 | 0 |
| X | 7595254 | 2 | 7 | 5 | 0 |
| X | 7595790 | 2 | 10 | 7 | 0 |
| X | 7596017 | 2 | 12 | 4 | 0 |
| X | 7596747 | 2 | 19 | 16 | 0 |
| X | 7599680 | 2 | 10 | 12 | 0 |
| X | 7600115 | 2 | 13 | 8 | 0 |
| X | 7602187 | 2 | 17 | 7 | 0 |
| X | 7602531 | 2 | 12 | 10 | 0 |
| X | 7603002 | 2 | 14 | 10 | 0 |
| X | 7603365 | 2 | 22 | 16 | 0 |
| X | 7603428 | 2 | 16 | 21 | 0 |
| X | 7603605 | 2 | 19 | 4 | 0 |
| X | 7603626 | 2 | 15 | 8 | 0 |
| X | 7604178 | 2 | 9 | 9 | 0 |
| X | 7604980 | 2 | 13 | 5 | 0 |
| X | 7606345 | 2 | 11 | 8 | 0 |
| X | 7607554 | 2 | 18 | 5 | 0 |
| X | 7608541 | 2 | 18 | 8 | 0 |
| X | 7608593 | 2 | 21 | 14 | 0 |
| X | 7608863 | 2 | 18 | 23 | 0 |
| X | 7610845 | 2 | 17 | 12 | 0 |
| X | 7612380 | 2 | 10 | 11 | 0 |
| X | 7612504 | 2 | 8 | 5 | 0 |
| X | 7612745 | 2 | 16 | 7 | 0 |
| X | 7613156 | 2 | 15 | 7 | 0 |
| X | 7613626 | 2 | 16 | 11 | 0 |
| X | 7613947 | 2 | 12 | 11 | 0 |
| X | 7614053 | 2 | 17 | 9 | 0 |
| X | 7614180 | 2 | 14 | 16 | 0 |
| X | 7615724 | 2 | 19 | 10 | 0 |
| X | 7621051 | 2 | 19 | 4 | 0 |
| X | 7621807 | 2 | 29 | 5 | 0 |
| X | 7623480 | 2 | 11 | 7 | 0 |
| X | 7623732 | 2 | 21 | 8 | 0 |
| X | 7624038 | 2 | 17 | 9 | 0 |
| X | 7624591 | 2 | 17 | 5 | 0 |
| X | 7624737 | 2 | 21 | 14 | 0 |
| X | 7624803 | 2 | 24 | 19 | 0 |
| X | 7625061 | 2 | 27 | 15 | 0 |
| X | 7626150 | 2 | 22 | 6 | 0 |
| X | 7626185 | 2 | 16 | 12 | 0 |
| X | 7626248 | 2 | 15 | 7 | 0 |
| X | 7626918 | 2 | 13 | 6 | 0 |
| X | 7626954 | 2 | 14 | 6 | 0 |
| X | 7627061 | 2 | 14 | 5 | 0 |
| X | 7627130 | 2 | 20 | 5 | 0 |
| X | 7634435 | 2 | 21 | 4 | 0 |
| X | 7639627 | 2 | 15 | 14 | 0 |
| X | 7639893 | 2 | 22 | 13 | 0 |
| X | 7640310 | 2 | 6 | 5 | 0 |
| X | 7641034 | 2 | 9 | 5 | 0 |
| X | 7641131 | 2 | 10 | 9 | 0 |
| X | 7641800 | 2 | 14 | 5 | 0 |
| X | 7642443 | 2 | 11 | 4 | 0 |
| X | 7643951 | 2 | 20 | 7 | 0 |
| X | 7646303 | 2 | 19 | 7 | 0 |
| X | 7647023 | 2 | 20 | 9 | 0 |
| X | 7647962 | 2 | 5 | 7 | 0 |
| X | 7648294 | 2 | 18 | 10 | 0 |
| X | 7648404 | 2 | 13 | 4 | 0 |
| X | 7648810 | 2 | 8 | 5 | 0 |
| X | 7649006 | 2 | 18 | 9 | 0 |
| X | 7649296 | 2 | 15 | 4 | 0 |
| X | 7649446 | 2 | 10 | 10 | 0 |
| X | 7649552 | 2 | 7 | 11 | 0 |
| X | 7649617 | 2 | 13 | 5 | 0 |
| X | 7651273 | 2 | 11 | 4 | 0 |
| X | 7651889 | 2 | 29 | 9 | 0 |
| X | 7654116 | 2 | 19 | 6 | 0 |
| X | 7655219 | 2 | 7 | 6 | 0 |
| X | 7655905 | 2 | 19 | 13 | 0 |
| X | 7657223 | 2 | 31 | 9 | 0 |
| X | 7657391 | 2 | 5 | 6 | 0 |
| X | 7657672 | 2 | 22 | 5 | 0 |
| X | 7658888 | 2 | 20 | 8 | 0 |
| X | 7661999 | 2 | 29 | 22 | 0 |
| X | 7665482 | 2 | 7 | 6 | 0 |
| X | 7666112 | 2 | 18 | 5 | 0 |
| X | 7666532 | 2 | 16 | 6 | 0 |
| X | 7667255 | 2 | 18 | 8 | 0 |
| X | 7669159 | 2 | 21 | 6 | 0 |
| X | 7672059 | 2 | 11 | 7 | 0 |
| X | 7673014 | 2 | 16 | 10 | 0 |
| X | 7674868 | 2 | 12 | 5 | 0 |
| X | 7675642 | 2 | 5 | 8 | 0 |
| X | 7675783 | 2 | 27 | 7 | 0 |
| X | 7675927 | 2 | 12 | 8 | 0 |
| X | 7676167 | 2 | 8 | 8 | 0 |
| X | 7680626 | 2 | 8 | 5 | 0 |
| X | 7681007 | 2 | 12 | 8 | 0 |
| X | 7683558 | 2 | 19 | 8 | 0 |
| X | 7684651 | 2 | 13 | 5 | 0 |
| X | 7685263 | 2 | 19 | 7 | 0 |
| X | 7685499 | 2 | 16 | 6 | 0 |
| X | 7685736 | 2 | 22 | 7 | 0 |
| X | 7686184 | 2 | 15 | 7 | 0 |
| X | 7686390 | 2 | 25 | 16 | 0 |
| X | 7686553 | 2 | 16 | 4 | 0 |
| X | 7686685 | 2 | 20 | 4 | 0 |
| X | 7687132 | 2 | 8 | 7 | 0 |
| X | 7687761 | 2 | 14 | 13 | 0 |
| X | 7687821 | 2 | 23 | 7 | 0 |
| X | 7688184 | 2 | 11 | 8 | 0 |
| X | 7688440 | 2 | 7 | 7 | 0 |
| X | 7688706 | 2 | 10 | 11 | 0 |
| X | 7688758 | 2 | 9 | 7 | 0 |
| X | 7688810 | 2 | 8 | 12 | 0 |
| X | 7691159 | 2 | 24 | 9 | 0 |
| X | 7691958 | 2 | 12 | 8 | 0 |
| X | 7692233 | 2 | 17 | 12 | 0 |
| X | 7692723 | 2 | 9 | 6 | 0 |
| X | 7692903 | 2 | 21 | 10 | 0 |
| X | 7692982 | 2 | 11 | 12 | 0 |
| X | 7693623 | 2 | 7 | 8 | 0 |
| X | 7696366 | 2 | 11 | 16 | 0 |
| X | 7696917 | 2 | 10 | 8 | 0 |
| X | 7697112 | 2 | 5 | 5 | 0 |
| X | 7698948 | 2 | 12 | 4 | 0 |
| X | 7700406 | 2 | 7 | 7 | 0 |
| X | 7703483 | 2 | 22 | 7 | 0 |
| X | 7703506 | 2 | 17 | 6 | 0 |
| X | 7703620 | 2 | 5 | 5 | 0 |
| X | 7703887 | 2 | 20 | 5 | 0 |
| X | 7705033 | 2 | 11 | 4 | 0 |
| X | 7705143 | 2 | 9 | 4 | 0 |
| X | 7706579 | 2 | 12 | 5 | 0 |
| X | 7716665 | 2 | 19 | 9 | 0 |
| X | 7719698 | 2 | 12 | 6 | 0 |
| X | 7720616 | 2 | 16 | 6 | 0 |
| X | 7721507 | 2 | 14 | 5 | 0 |
| X | 7722013 | 2 | 9 | 8 | 0 |
| X | 7724105 | 2 | 7 | 12 | 0 |
| X | 7724731 | 2 | 5 | 6 | 0 |
| X | 7724786 | 2 | 14 | 7 | 0 |
| X | 7725085 | 2 | 12 | 6 | 0 |
| X | 7725792 | 2 | 9 | 6 | 0 |
| X | 7727658 | 2 | 10 | 5 | 0 |
| X | 7729430 | 2 | 15 | 9 | 0 |
| X | 7734118 | 2 | 5 | 5 | 0 |
| X | 7734749 | 2 | 4 | 14 | 0 |
| X | 7737477 | 2 | 11 | 21 | 0 |
| X | 7737500 | 2 | 15 | 19 | 0 |
| X | 7737940 | 2 | 19 | 11 | 0 |
| X | 7738556 | 2 | 15 | 5 | 0 |
| X | 7739423 | 2 | 14 | 6 | 0 |
| X | 7739583 | 2 | 20 | 10 | 0 |
| X | 7739732 | 2 | 20 | 17 | 0 |
| X | 7741430 | 2 | 14 | 7 | 0 |
| X | 7743111 | 2 | 4 | 22 | 0 |
| X | 7744074 | 2 | 8 | 10 | 0 |
| X | 7744084 | 2 | 10 | 9 | 0 |
| X | 7745225 | 2 | 20 | 10 | 0 |
| X | 7746426 | 2 | 21 | 15 | 0 |
| X | 7747330 | 2 | 4 | 21 | 0 |
| X | 7747354 | 2 | 5 | 29 | 0 |
| X | 7749702 | 2 | 16 | 10 | 0 |
| X | 7751905 | 2 | 9 | 25 | 0 |
| X | 7752373 | 2 | 11 | 18 | 0 |
| X | 7752487 | 2 | 12 | 15 | 0 |
| X | 7752613 | 2 | 18 | 18 | 0 |
| X | 7753417 | 2 | 6 | 8 | 0 |
| X | 7754653 | 2 | 10 | 18 | 0 |
| X | 7755064 | 2 | 10 | 15 | 0 |
| X | 7755101 | 2 | 7 | 27 | 0 |
| X | 7755773 | 2 | 4 | 20 | 0 |
| X | 7756386 | 2 | 4 | 13 | 0 |
| X | 7756542 | 2 | 5 | 10 | 0 |
| X | 7759446 | 2 | 27 | 30 | 0 |
| X | 7762434 | 2 | 5 | 18 | 0 |
| X | 7765932 | 2 | 17 | 21 | 0 |
| X | 7766253 | 2 | 14 | 4 | 0 |
| X | 7766340 | 2 | 27 | 5 | 0 |
| X | 7769807 | 2 | 17 | 5 | 0 |
| X | 7769889 | 2 | 23 | 25 | 0 |
| X | 7769980 | 2 | 7 | 13 | 0 |
| X | 7770020 | 2 | 20 | 4 | 0 |
| X | 7770070 | 2 | 13 | 6 | 0 |
| X | 7770124 | 2 | 21 | 14 | 0 |
| X | 7771042 | 2 | 39 | 5 | 0 |
| X | 7771187 | 2 | 20 | 13 | 0 |
| X | 7771678 | 2 | 6 | 15 | 0 |
| X | 7772442 | 2 | 12 | 5 | 0 |
| X | 7774888 | 2 | 9 | 7 | 0 |
| X | 7774999 | 2 | 10 | 21 | 0 |
| X | 7775074 | 2 | 7 | 33 | 0 |
| X | 7775247 | 2 | 4 | 23 | 0 |
| X | 7775284 | 2 | 3 | 35 | 0 |
| X | 7775320 | 2 | 8 | 28 | 0 |
| X | 7775781 | 2 | 15 | 24 | 0 |
| X | 7777215 | 2 | 6 | 15 | 0 |
| X | 7777404 | 2 | 9 | 17 | 0 |
| X | 7777569 | 2 | 5 | 37 | 0 |
| X | 7777623 | 2 | 15 | 24 | 0 |
| X | 7777767 | 2 | 6 | 13 | 0 |
| X | 7778210 | 2 | 4 | 13 | 0 |
| X | 7778366 | 2 | 12 | 4 | 0 |
| X | 7778407 | 2 | 21 | 12 | 0 |
| X | 7778650 | 2 | 5 | 9 | 0 |
| X | 7780412 | 2 | 13 | 21 | 0 |
| X | 7786718 | 2 | 6 | 12 | 0 |
| X | 7787359 | 2 | 11 | 24 | 0 |
| X | 7800341 | 2 | 19 | 13 | 0 |
| X | 7804678 | 2 | 12 | 4 | 0 |
| X | 7804852 | 2 | 26 | 6 | 0 |
| X | 7805494 | 2 | 20 | 5 | 0 |
| X | 7805622 | 2 | 20 | 7 | 0 |
| X | 7805664 | 2 | 14 | 5 | 0 |
| X | 7807628 | 2 | 4 | 14 | 0 |
| X | 7807774 | 2 | 4 | 14 | 0 |
| X | 7808026 | 2 | 13 | 9 | 0 |
| X | 7812158 | 2 | 18 | 11 | 0 |
| X | 7812374 | 2 | 17 | 19 | 0 |
| X | 7812726 | 2 | 10 | 8 | 0 |
| X | 7812923 | 2 | 12 | 22 | 0 |
| X | 7812964 | 2 | 11 | 25 | 0 |
| X | 7813471 | 2 | 9 | 25 | 0 |
| X | 7815292 | 2 | 20 | 10 | 0 |
| X | 7815898 | 2 | 11 | 11 | 0 |
| X | 7815973 | 2 | 6 | 21 | 0 |
| X | 7816391 | 2 | 13 | 5 | 0 |
| X | 7818730 | 2 | 15 | 11 | 0 |
| X | 7819371 | 2 | 11 | 8 | 0 |
| X | 7819430 | 2 | 12 | 14 | 0 |
| X | 7820811 | 2 | 19 | 15 | 0 |
| X | 7830565 | 2 | 9 | 6 | 0 |
| X | 7852355 | 2 | 8 | 6 | 0 |
| X | 7855784 | 2 | 10 | 8 | 0 |
| X | 7857438 | 2 | 19 | 30 | 0 |
| X | 7857597 | 2 | 8 | 17 | 0 |
| X | 7859504 | 2 | 4 | 32 | 0 |
| X | 7859566 | 2 | 6 | 18 | 0 |
| X | 7859843 | 2 | 6 | 7 | 0 |
| X | 7859871 | 2 | 15 | 14 | 0 |
| X | 7863423 | 2 | 13 | 6 | 0 |
| X | 7867378 | 2 | 32 | 15 | 0 |
| X | 7867418 | 2 | 15 | 13 | 0 |
| X | 7870516 | 2 | 9 | 9 | 0 |
| X | 7872270 | 2 | 14 | 11 | 0 |
| X | 7873112 | 2 | 4 | 23 | 0 |
| X | 7873291 | 2 | 18 | 19 | 0 |
| X | 7874197 | 2 | 5 | 19 | 0 |
| X | 7874278 | 2 | 7 | 9 | 0 |
| X | 7875816 | 2 | 12 | 21 | 0 |
| X | 7876090 | 2 | 12 | 11 | 0 |
| X | 7876387 | 2 | 21 | 7 | 0 |
| X | 7876420 | 2 | 18 | 4 | 0 |
| X | 7876695 | 2 | 17 | 5 | 0 |
| X | 7876713 | 2 | 21 | 8 | 0 |
| X | 7876756 | 2 | 20 | 4 | 0 |
| X | 7876840 | 2 | 15 | 4 | 0 |
| X | 7878450 | 2 | 6 | 26 | 0 |
| X | 7878560 | 2 | 4 | 11 | 0 |
| X | 7878701 | 2 | 6 | 12 | 0 |
| X | 7879067 | 2 | 11 | 26 | 0 |
| X | 7879195 | 2 | 8 | 21 | 0 |
| X | 7881150 | 2 | 6 | 18 | 0 |
| X | 7881237 | 2 | 11 | 13 | 0 |
| X | 7881418 | 2 | 4 | 11 | 0 |
| X | 7885851 | 2 | 4 | 18 | 0 |
| X | 7886888 | 2 | 10 | 15 | 0 |
| X | 7887365 | 2 | 11 | 10 | 0 |
| X | 7887566 | 2 | 10 | 6 | 0 |
| X | 7888001 | 2 | 12 | 5 | 0 |
| X | 7888856 | 2 | 11 | 8 | 0 |
| X | 7890524 | 2 | 5 | 13 | 0 |
| X | 7890617 | 2 | 7 | 16 | 0 |
| X | 7891451 | 2 | 4 | 17 | 0 |
| X | 7892050 | 2 | 21 | 6 | 0 |
| X | 7894371 | 2 | 11 | 11 | 0 |
| X | 7894547 | 2 | 27 | 18 | 0 |
| X | 7895767 | 2 | 18 | 5 | 0 |
| X | 7895908 | 2 | 27 | 15 | 0 |
| X | 7896815 | 2 | 15 | 7 | 0 |
| X | 7897936 | 2 | 7 | 19 | 0 |
| X | 7898383 | 2 | 5 | 7 | 0 |
| X | 7901174 | 2 | 12 | 16 | 0 |
| X | 7904216 | 2 | 12 | 6 | 0 |
| X | 7905397 | 2 | 14 | 5 | 0 |
| X | 7908945 | 2 | 5 | 19 | 0 |
| X | 7909216 | 2 | 16 | 12 | 0 |
| X | 7910634 | 2 | 5 | 11 | 0 |
| X | 7911754 | 2 | 5 | 6 | 0 |
| X | 7912804 | 2 | 23 | 10 | 0 |
| X | 7913774 | 2 | 18 | 6 | 0 |
| X | 7914953 | 2 | 15 | 5 | 0 |
| X | 7915220 | 2 | 26 | 7 | 0 |
| X | 7915475 | 2 | 14 | 14 | 0 |
| X | 7917050 | 2 | 5 | 7 | 0 |
| X | 7920598 | 2 | 13 | 15 | 0 |
| X | 7922327 | 2 | 15 | 7 | 0 |
| X | 7922713 | 2 | 11 | 10 | 0 |
| X | 7924130 | 2 | 10 | 8 | 0 |
| X | 7924159 | 2 | 8 | 5 | 0 |
| X | 7924605 | 2 | 13 | 4 | 0 |
| X | 7925240 | 2 | 19 | 4 | 0 |
| X | 7925297 | 2 | 22 | 7 | 0 |
| X | 7925415 | 2 | 5 | 12 | 0 |
| X | 7925461 | 2 | 26 | 17 | 0 |
| X | 7929959 | 2 | 23 | 5 | 0 |
| X | 7929981 | 2 | 31 | 8 | 0 |
| X | 7930415 | 2 | 30 | 6 | 0 |
| X | 7930582 | 2 | 36 | 5 | 0 |
| X | 7931406 | 2 | 23 | 5 | 0 |
| X | 7932524 | 2 | 35 | 6 | 0 |
| X | 7933157 | 2 | 11 | 17 | 0 |
| X | 7935090 | 2 | 8 | 28 | 0 |
| X | 7936574 | 2 | 17 | 15 | 0 |
| X | 7936931 | 2 | 20 | 7 | 0 |
| X | 7937243 | 2 | 6 | 5 | 0 |
| X | 7937261 | 2 | 5 | 13 | 0 |
| X | 7937787 | 2 | 10 | 30 | 0 |
| X | 7938088 | 2 | 12 | 23 | 0 |
| X | 7938159 | 2 | 7 | 6 | 0 |
| X | 7938320 | 2 | 4 | 20 | 0 |
| X | 7938544 | 2 | 13 | 9 | 0 |
| X | 7940801 | 2 | 8 | 9 | 0 |
| X | 7941348 | 2 | 10 | 13 | 0 |
| X | 7941967 | 2 | 23 | 18 | 0 |
| X | 7943668 | 2 | 9 | 23 | 0 |
| X | 7944331 | 2 | 6 | 20 | 0 |
| X | 7947508 | 2 | 5 | 5 | 0 |
| X | 7948801 | 2 | 6 | 7 | 0 |
| X | 7950498 | 2 | 12 | 28 | 0 |
| X | 7951007 | 2 | 5 | 15 | 0 |
| X | 7951804 | 2 | 17 | 19 | 0 |
| X | 7952006 | 2 | 10 | 17 | 0 |
| X | 7952110 | 2 | 13 | 23 | 0 |
| X | 7952663 | 2 | 5 | 7 | 0 |
| X | 7952946 | 2 | 7 | 23 | 0 |
| X | 7953814 | 2 | 6 | 13 | 0 |
| X | 7955794 | 2 | 4 | 19 | 0 |
| X | 7956868 | 2 | 7 | 13 | 0 |
| X | 7957009 | 2 | 11 | 15 | 0 |
| X | 7957446 | 2 | 4 | 12 | 0 |
| X | 7957907 | 2 | 11 | 28 | 0 |
| X | 7958107 | 2 | 6 | 10 | 0 |
| X | 7958544 | 2 | 14 | 27 | 0 |
| X | 7958651 | 2 | 18 | 28 | 0 |
| X | 7960570 | 2 | 5 | 12 | 0 |
| X | 7960909 | 2 | 8 | 30 | 0 |
| X | 7961358 | 2 | 4 | 27 | 0 |
| X | 7962654 | 2 | 11 | 8 | 0 |
| X | 7963154 | 2 | 14 | 29 | 0 |
| X | 7964852 | 2 | 24 | 10 | 0 |
| X | 7965345 | 2 | 11 | 8 | 0 |
| X | 7966162 | 2 | 24 | 16 | 0 |
| X | 7966500 | 2 | 19 | 11 | 0 |
| X | 7966972 | 2 | 16 | 9 | 0 |
| X | 7967170 | 2 | 10 | 19 | 0 |
| X | 7967428 | 2 | 11 | 12 | 0 |
| X | 7967713 | 2 | 22 | 11 | 0 |
| X | 7967740 | 2 | 11 | 17 | 0 |
| X | 7968034 | 2 | 13 | 8 | 0 |
| X | 7968154 | 2 | 30 | 15 | 0 |
| X | 7969323 | 2 | 5 | 23 | 0 |
| X | 7969756 | 2 | 5 | 9 | 0 |
| X | 7970291 | 2 | 10 | 11 | 0 |
| X | 7970367 | 2 | 16 | 8 | 0 |
| X | 7970522 | 2 | 17 | 35 | 0 |
| X | 7970888 | 2 | 15 | 7 | 0 |
| X | 7971686 | 2 | 12 | 12 | 0 |
| X | 7971722 | 2 | 10 | 17 | 0 |
| X | 7971839 | 2 | 18 | 12 | 0 |
| X | 7971845 | 2 | 18 | 11 | 0 |
| X | 7972058 | 2 | 16 | 4 | 0 |
| X | 7972250 | 2 | 9 | 12 | 0 |
| X | 7972854 | 2 | 12 | 21 | 0 |
| X | 7972973 | 2 | 20 | 7 | 0 |
| X | 7973443 | 2 | 4 | 30 | 0 |
| X | 7973673 | 2 | 15 | 23 | 0 |
| X | 7973904 | 2 | 10 | 13 | 0 |
| X | 7974232 | 2 | 6 | 12 | 0 |
| X | 7974488 | 2 | 10 | 14 | 0 |
| X | 7974749 | 2 | 7 | 24 | 0 |
| X | 7975209 | 2 | 4 | 18 | 0 |
| X | 7977084 | 2 | 6 | 13 | 0 |
| X | 7977647 | 2 | 16 | 18 | 0 |
| X | 7977960 | 2 | 13 | 19 | 0 |
| X | 7978612 | 2 | 13 | 4 | 0 |
| X | 7978768 | 2 | 9 | 9 | 0 |
| X | 7978956 | 2 | 10 | 27 | 0 |
| X | 7979346 | 2 | 5 | 12 | 0 |
| X | 7979862 | 2 | 22 | 6 | 0 |
| X | 7979887 | 2 | 24 | 5 | 0 |
| X | 7980126 | 2 | 6 | 6 | 0 |
| X | 7980308 | 2 | 5 | 12 | 0 |
| X | 7981871 | 2 | 21 | 15 | 0 |
| X | 7982289 | 2 | 18 | 7 | 0 |
| X | 7982688 | 2 | 21 | 8 | 0 |
| X | 7984886 | 2 | 9 | 11 | 0 |
| X | 7985380 | 2 | 5 | 17 | 0 |
| X | 7985577 | 2 | 14 | 25 | 0 |
| X | 7986298 | 2 | 15 | 6 | 0 |
| X | 7987515 | 2 | 7 | 11 | 0 |
| X | 7988361 | 2 | 14 | 4 | 0 |
| X | 7988960 | 2 | 12 | 9 | 0 |
| X | 7990561 | 2 | 7 | 7 | 0 |
| X | 8036209 | 2 | 8 | 17 | 0 |
| X | 8487714 | 2 | 12 | 6 | 0 |
| X | 8491505 | 2 | 22 | 7 | 0 |
| X | 8493370 | 2 | 4 | 11 | 0 |
| X | 8499315 | 2 | 7 | 14 | 0 |
| X | 8501683 | 2 | 9 | 13 | 0 |
| X | 8504648 | 2 | 22 | 5 | 0 |
| X | 8505465 | 2 | 8 | 10 | 0 |
| X | 8510111 | 2 | 5 | 19 | 0 |
| X | 8510529 | 2 | 7 | 14 | 0 |
| X | 8510686 | 2 | 5 | 23 | 0 |
| X | 8511170 | 2 | 9 | 18 | 0 |
| X | 8511791 | 2 | 18 | 7 | 0 |
| X | 8511919 | 2 | 24 | 7 | 0 |
| X | 8513934 | 2 | 11 | 6 | 0 |
| X | 8514402 | 2 | 14 | 13 | 0 |
| X | 8515103 | 2 | 8 | 6 | 0 |
| X | 8521299 | 2 | 6 | 15 | 0 |
| X | 8524577 | 2 | 9 | 20 | 0 |
| X | 8527087 | 2 | 4 | 17 | 0 |
| X | 8527795 | 2 | 28 | 7 | 0 |
| X | 8529029 | 2 | 12 | 17 | 0 |
| X | 8529167 | 2 | 4 | 15 | 0 |
| X | 8531308 | 2 | 8 | 10 | 0 |
| X | 8531623 | 2 | 13 | 5 | 0 |
| X | 8532011 | 2 | 4 | 23 | 0 |
| X | 8533874 | 2 | 11 | 6 | 0 |
| X | 8538314 | 2 | 8 | 22 | 0 |
| X | 8539442 | 2 | 8 | 19 | 0 |
| X | 8539921 | 2 | 5 | 10 | 0 |
| X | 8545712 | 2 | 16 | 8 | 0 |
| X | 8549529 | 2 | 6 | 17 | 0 |
| X | 8552725 | 2 | 5 | 5 | 0 |
| X | 8553670 | 2 | 6 | 25 | 0 |
| X | 8555143 | 2 | 13 | 5 | 0 |
| X | 8560455 | 2 | 15 | 4 | 0 |
| X | 8564106 | 2 | 11 | 13 | 0 |
| X | 8565533 | 2 | 4 | 20 | 0 |
| X | 8565818 | 2 | 8 | 6 | 0 |
| X | 8566661 | 2 | 20 | 5 | 0 |
| X | 8567053 | 2 | 21 | 11 | 0 |
| X | 8567668 | 2 | 25 | 11 | 0 |
| X | 8568935 | 2 | 16 | 6 | 0 |
| X | 8569872 | 2 | 9 | 7 | 0 |
| X | 8570539 | 2 | 4 | 11 | 0 |
| X | 8571749 | 2 | 19 | 12 | 0 |
| X | 8572688 | 2 | 11 | 5 | 0 |
| X | 8575714 | 2 | 7 | 6 | 0 |
| X | 8575808 | 2 | 11 | 5 | 0 |
| X | 8580597 | 2 | 15 | 4 | 0 |
| X | 8606163 | 2 | 18 | 10 | 0 |
| X | 8606664 | 2 | 11 | 5 | 0 |
| X | 8612844 | 2 | 14 | 8 | 0 |
| X | 8614689 | 2 | 21 | 8 | 0 |
| X | 8614737 | 2 | 19 | 17 | 0 |
| X | 8615344 | 2 | 10 | 23 | 0 |
| X | 8618271 | 2 | 14 | 4 | 0 |
| X | 8624461 | 2 | 20 | 5 | 0 |
| X | 8624749 | 2 | 5 | 14 | 0 |
| X | 8631806 | 2 | 9 | 13 | 0 |
| X | 8632307 | 2 | 9 | 5 | 0 |
| X | 8635952 | 2 | 11 | 5 | 0 |
| X | 8638401 | 2 | 12 | 9 | 0 |
| X | 8640120 | 2 | 7 | 18 | 0 |
| X | 8641218 | 2 | 20 | 11 | 0 |
| X | 8642148 | 2 | 4 | 14 | 0 |
| X | 8650117 | 2 | 15 | 5 | 0 |
| X | 8650505 | 2 | 26 | 11 | 0 |
| X | 8653347 | 2 | 10 | 17 | 0 |
| X | 8661725 | 2 | 13 | 5 | 0 |
| X | 8662505 | 2 | 24 | 6 | 0 |
| X | 8666619 | 2 | 14 | 8 | 0 |
| X | 8678733 | 2 | 7 | 7 | 0 |
| X | 8684896 | 2 | 5 | 6 | 0 |
| X | 8692095 | 2 | 11 | 6 | 0 |
| X | 8695693 | 2 | 15 | 5 | 0 |
| X | 8697140 | 2 | 19 | 5 | 0 |
| X | 8699581 | 2 | 8 | 19 | 0 |
| X | 8704924 | 2 | 11 | 6 | 0 |
| X | 8706035 | 2 | 6 | 19 | 0 |
| X | 8708973 | 2 | 13 | 4 | 0 |
| X | 8714985 | 2 | 8 | 5 | 0 |
| X | 8717546 | 2 | 10 | 9 | 0 |
| X | 8723912 | 2 | 17 | 6 | 0 |
| X | 8726814 | 2 | 6 | 5 | 0 |
| X | 8727107 | 2 | 6 | 13 | 0 |
| X | 8728169 | 2 | 27 | 11 | 0 |
| X | 8728753 | 2 | 15 | 4 | 0 |
| X | 8733503 | 2 | 11 | 14 | 0 |
| X | 8733831 | 2 | 17 | 6 | 0 |
| X | 8740352 | 2 | 7 | 9 | 0 |
| X | 8741637 | 2 | 12 | 4 | 0 |
| X | 8745173 | 2 | 9 | 4 | 0 |
| X | 8747149 | 2 | 6 | 15 | 0 |
| X | 8751030 | 2 | 7 | 21 | 0 |
| X | 8751119 | 2 | 5 | 12 | 0 |
| X | 8758880 | 2 | 11 | 9 | 0 |
| X | 8765229 | 2 | 7 | 8 | 0 |
| X | 8768842 | 2 | 9 | 13 | 0 |
| X | 8768881 | 2 | 14 | 5 | 0 |
| X | 8772885 | 2 | 9 | 11 | 0 |
| X | 8773085 | 2 | 15 | 13 | 0 |
| X | 8775122 | 2 | 12 | 4 | 0 |
| X | 8775262 | 2 | 5 | 8 | 0 |
| X | 8777741 | 2 | 23 | 9 | 0 |
| X | 8780278 | 2 | 23 | 12 | 0 |
| X | 8780314 | 2 | 22 | 12 | 0 |
| X | 8791522 | 2 | 10 | 4 | 0 |
| X | 8796439 | 2 | 12 | 5 | 0 |
| X | 8798865 | 2 | 15 | 10 | 0 |
| X | 8799112 | 2 | 6 | 16 | 0 |
| X | 8802828 | 2 | 4 | 12 | 0 |
| X | 8806039 | 2 | 8 | 5 | 0 |
| X | 9597793 | 2 | 21 | 11 | 0 |
| X | 9601645 | 2 | 19 | 5 | 0 |
| X | 9602106 | 2 | 30 | 4 | 0 |
| X | 9602734 | 2 | 20 | 6 | 0 |
| X | 9603557 | 2 | 20 | 10 | 0 |
| X | 9611528 | 2 | 11 | 7 | 0 |
| X | 9615995 | 2 | 5 | 9 | 0 |
| X | 9618231 | 2 | 16 | 7 | 0 |
| X | 9625657 | 2 | 22 | 14 | 0 |
| X | 9627309 | 2 | 10 | 12 | 0 |
| X | 9627839 | 2 | 10 | 11 | 0 |
| X | 9628907 | 2 | 13 | 12 | 0 |
| X | 9629335 | 2 | 23 | 6 | 0 |
| X | 9629956 | 2 | 6 | 12 | 0 |
| X | 9630867 | 2 | 14 | 10 | 0 |
| X | 9630948 | 2 | 6 | 5 | 0 |
| X | 9631597 | 2 | 19 | 13 | 0 |
| X | 9634021 | 2 | 10 | 4 | 0 |
| X | 9636020 | 2 | 4 | 30 | 0 |
| X | 9641019 | 2 | 4 | 13 | 0 |
| X | 9641705 | 2 | 21 | 10 | 0 |
| X | 9651206 | 2 | 7 | 5 | 0 |
| X | 9651269 | 2 | 10 | 6 | 0 |
| X | 9652605 | 2 | 23 | 9 | 0 |
| X | 9654455 | 2 | 19 | 15 | 0 |
| X | 9656532 | 2 | 4 | 13 | 0 |
| X | 9658615 | 2 | 5 | 7 | 0 |
| X | 9662485 | 2 | 11 | 9 | 0 |
| X | 9667968 | 2 | 4 | 23 | 0 |
| X | 9670601 | 2 | 6 | 17 | 0 |
| X | 9675170 | 2 | 10 | 9 | 0 |
| X | 9675467 | 2 | 9 | 17 | 0 |
| X | 9675515 | 2 | 7 | 14 | 0 |
| X | 9676028 | 2 | 22 | 6 | 0 |
| X | 9677176 | 2 | 15 | 6 | 0 |
| X | 9677446 | 2 | 18 | 5 | 0 |
| X | 9677562 | 2 | 11 | 4 | 0 |
| X | 9679038 | 2 | 4 | 11 | 0 |
| X | 9682766 | 2 | 13 | 10 | 0 |
| X | 9683334 | 2 | 13 | 24 | 0 |
| X | 9683395 | 2 | 12 | 5 | 0 |
| X | 9688125 | 2 | 13 | 15 | 0 |
| X | 9688497 | 2 | 7 | 5 | 0 |
| X | 9689106 | 2 | 8 | 8 | 0 |
| X | 9692076 | 2 | 11 | 10 | 0 |
| X | 9700477 | 2 | 18 | 8 | 0 |
| X | 9702571 | 2 | 12 | 4 | 0 |
| X | 9702811 | 2 | 11 | 4 | 0 |
| X | 9714680 | 2 | 11 | 14 | 0 |
| X | 9717786 | 2 | 4 | 17 | 0 |
| X | 9721863 | 2 | 10 | 19 | 0 |
| X | 9727544 | 2 | 4 | 12 | 0 |
| X | 9728233 | 2 | 10 | 7 | 0 |
| X | 9728514 | 2 | 10 | 7 | 0 |
| X | 9729305 | 2 | 4 | 15 | 0 |
| X | 9729526 | 2 | 24 | 4 | 0 |
| X | 9732097 | 2 | 43 | 5 | 0 |
| X | 9734277 | 2 | 9 | 14 | 0 |
| X | 9736527 | 2 | 13 | 13 | 0 |
| X | 9737703 | 2 | 5 | 5 | 0 |
| X | 9740448 | 2 | 5 | 26 | 0 |
| X | 9741161 | 2 | 12 | 18 | 0 |
| X | 9743163 | 2 | 20 | 14 | 0 |
| X | 9744144 | 2 | 4 | 12 | 0 |
| X | 9746625 | 2 | 13 | 7 | 0 |
| X | 9746820 | 2 | 16 | 7 | 0 |
| X | 9748309 | 2 | 12 | 8 | 0 |
| X | 9751110 | 2 | 10 | 14 | 0 |
| X | 9752281 | 2 | 7 | 7 | 0 |
| X | 9754162 | 2 | 13 | 12 | 0 |
| X | 9756765 | 2 | 10 | 19 | 0 |
| X | 9757853 | 2 | 8 | 5 | 0 |
| X | 9762803 | 2 | 6 | 10 | 0 |
| X | 9766180 | 2 | 10 | 22 | 0 |
| X | 9767878 | 2 | 17 | 7 | 0 |
| X | 9768405 | 2 | 15 | 11 | 0 |
| X | 9768637 | 2 | 23 | 4 | 0 |
| X | 9770452 | 2 | 5 | 17 | 0 |
| X | 9771205 | 2 | 22 | 5 | 0 |
| X | 9775823 | 2 | 5 | 6 | 0 |
| X | 9777924 | 2 | 18 | 15 | 0 |
| X | 9780540 | 2 | 5 | 15 | 0 |
| X | 9784672 | 2 | 10 | 10 | 0 |
| X | 9797926 | 2 | 8 | 8 | 0 |
| X | 9806386 | 2 | 20 | 8 | 0 |
| X | 9813110 | 2 | 8 | 22 | 0 |
| X | 9816471 | 2 | 4 | 10 | 0 |
| X | 9820158 | 2 | 10 | 18 | 0 |
| X | 9821277 | 2 | 10 | 12 | 0 |
| X | 9832090 | 2 | 16 | 7 | 0 |
| X | 9835115 | 2 | 5 | 18 | 0 |
| X | 9835381 | 2 | 16 | 9 | 0 |
| X | 9836055 | 2 | 7 | 11 | 0 |
| X | 9836893 | 2 | 8 | 12 | 0 |
| X | 9838358 | 2 | 10 | 17 | 0 |
| X | 9840568 | 2 | 10 | 13 | 0 |
| X | 9841356 | 2 | 11 | 6 | 0 |
| X | 9841373 | 2 | 5 | 20 | 0 |
| X | 9841549 | 2 | 17 | 9 | 0 |
| X | 9843558 | 2 | 17 | 12 | 0 |
| X | 9845952 | 2 | 4 | 17 | 0 |
| X | 9847689 | 2 | 9 | 18 | 0 |
| X | 9848006 | 2 | 18 | 26 | 0 |
| X | 9863246 | 2 | 5 | 11 | 0 |
| X | 9875378 | 2 | 9 | 8 | 0 |
| X | 9876161 | 2 | 9 | 6 | 0 |
| X | 9879430 | 2 | 4 | 9 | 0 |
| X | 9881252 | 2 | 6 | 14 | 0 |
| X | 9888438 | 2 | 9 | 4 | 0 |
| X | 9889544 | 2 | 19 | 6 | 0 |
| X | 9905272 | 2 | 15 | 10 | 0 |
| X | 9905765 | 2 | 23 | 6 | 0 |
| X | 9908579 | 2 | 18 | 7 | 0 |
| X | 9909480 | 2 | 6 | 11 | 0 |
| X | 9911480 | 2 | 10 | 5 | 0 |
| X | 9916899 | 2 | 4 | 11 | 0 |
| X | 9926594 | 2 | 15 | 6 | 0 |
| X | 9926813 | 2 | 25 | 8 | 0 |
| X | 9987792 | 2 | 16 | 11 | 0 |
| X | 9988438 | 2 | 14 | 4 | 0 |
| X | 9998394 | 2 | 10 | 5 | 0 |
| X | 10000981 | 2 | 9 | 7 | 0 |
| X | 10005853 | 2 | 10 | 12 | 0 |
| X | 10015874 | 2 | 8 | 6 | 0 |
| X | 10023477 | 2 | 6 | 5 | 0 |
| X | 10027816 | 2 | 13 | 6 | 0 |
| X | 10030683 | 2 | 9 | 12 | 0 |
| X | 10041037 | 2 | 8 | 12 | 0 |
| X | 10058842 | 2 | 9 | 6 | 0 |
| X | 10063093 | 2 | 7 | 6 | 0 |
| X | 10065610 | 2 | 8 | 12 | 0 |
| X | 10069501 | 2 | 9 | 8 | 0 |
| X | 10070897 | 2 | 6 | 10 | 0 |
| X | 10071573 | 2 | 12 | 7 | 0 |
| X | 10072613 | 2 | 5 | 5 | 0 |
| X | 10074680 | 2 | 17 | 5 | 0 |
| X | 10082052 | 2 | 5 | 20 | 0 |
| X | 10082372 | 2 | 12 | 4 | 0 |
| X | 10083535 | 2 | 18 | 19 | 0 |
| X | 10089806 | 2 | 23 | 17 | 0 |
| X | 10111164 | 2 | 11 | 6 | 0 |
| X | 10112075 | 2 | 11 | 11 | 0 |
| X | 10117604 | 2 | 15 | 16 | 0 |
| X | 10122673 | 2 | 13 | 6 | 0 |
| X | 10132705 | 2 | 20 | 16 | 0 |
| X | 10141137 | 2 | 12 | 4 | 0 |
| X | 10147020 | 2 | 6 | 13 | 0 |
| X | 10152293 | 2 | 9 | 4 | 0 |
| X | 10152296 | 2 | 9 | 4 | 0 |
| X | 10181586 | 2 | 8 | 28 | 0 |
| X | 10204994 | 2 | 6 | 7 | 0 |
| X | 10236036 | 2 | 7 | 6 | 0 |
| X | 10236807 | 2 | 9 | 26 | 0 |
| X | 10238306 | 2 | 15 | 6 | 0 |
| X | 10265108 | 2 | 12 | 4 | 0 |
| X | 10278075 | 2 | 5 | 5 | 0 |
| X | 10278665 | 2 | 14 | 9 | 0 |
| X | 10286224 | 2 | 10 | 5 | 0 |
| X | 10302867 | 2 | 5 | 10 | 0 |
| X | 10314936 | 2 | 16 | 10 | 0 |
| X | 10330164 | 2 | 18 | 13 | 0 |
| X | 10332087 | 2 | 8 | 12 | 0 |
| X | 10332929 | 2 | 9 | 8 | 0 |
| X | 10340722 | 2 | 16 | 10 | 0 |
| X | 10356035 | 2 | 4 | 12 | 0 |
| X | 10369817 | 2 | 16 | 5 | 0 |
| X | 10383814 | 2 | 6 | 8 | 0 |
| X | 10388269 | 2 | 12 | 12 | 0 |
| X | 10388770 | 2 | 4 | 16 | 0 |
| X | 10392926 | 2 | 4 | 10 | 0 |
| X | 10397161 | 2 | 5 | 9 | 0 |
| X | 10426425 | 2 | 8 | 9 | 0 |
| X | 10431347 | 2 | 8 | 6 | 0 |
| X | 10680623 | 2 | 5 | 8 | 0 |
| X | 10716136 | 2 | 9 | 7 | 0 |
| X | 10737801 | 2 | 9 | 16 | 0 |
| X | 10740595 | 2 | 15 | 12 | 0 |
| X | 10746115 | 2 | 6 | 5 | 0 |
| X | 10749655 | 2 | 7 | 12 | 0 |
| X | 10770635 | 2 | 5 | 10 | 0 |
| X | 10888442 | 2 | 9 | 4 | 0 |
| X | 10892563 | 2 | 6 | 11 | 0 |
| X | 10955764 | 2 | 17 | 8 | 0 |
| X | 10982983 | 2 | 12 | 4 | 0 |
| X | 10987860 | 2 | 7 | 5 | 0 |
| X | 11044494 | 2 | 4 | 9 | 0 |
| X | 11137383 | 2 | 6 | 7 | 0 |
| X | 11144492 | 2 | 5 | 15 | 0 |
| X | 11158856 | 2 | 5 | 15 | 0 |
| X | 11199530 | 2 | 14 | 4 | 0 |
| X | 11202501 | 2 | 5 | 9 | 0 |
| X | 11253688 | 2 | 6 | 8 | 0 |
| X | 11304781 | 2 | 5 | 22 | 0 |
| X | 11306294 | 2 | 4 | 12 | 0 |
| X | 11352230 | 2 | 4 | 19 | 0 |
| X | 11362630 | 2 | 5 | 7 | 0 |
| X | 11567143 | 2 | 7 | 8 | 0 |
| X | 11578704 | 2 | 4 | 9 | 0 |
| X | 11619172 | 2 | 5 | 7 | 0 |
| X | 11651075 | 2 | 7 | 7 | 0 |
| X | 11658407 | 2 | 7 | 10 | 0 |
| X | 11701807 | 2 | 6 | 5 | 0 |
| X | 11703914 | 2 | 5 | 17 | 0 |
| X | 11744343 | 2 | 5 | 9 | 0 |
| X | 11769268 | 2 | 7 | 8 | 0 |
| X | 11775906 | 2 | 26 | 8 | 0 |
| X | 11779923 | 2 | 6 | 8 | 0 |
| X | 11795880 | 2 | 10 | 9 | 0 |
| X | 11855391 | 2 | 6 | 13 | 0 |
| X | 11867221 | 2 | 8 | 15 | 0 |
| X | 11876563 | 2 | 9 | 4 | 0 |
| X | 11894088 | 2 | 12 | 7 | 0 |
| X | 11896933 | 2 | 14 | 6 | 0 |
| X | 11919270 | 2 | 5 | 6 | 0 |
| X | 11938650 | 2 | 6 | 16 | 0 |
| X | 11943205 | 2 | 4 | 9 | 0 |
| X | 11944820 | 2 | 16 | 9 | 0 |
| X | 12026698 | 2 | 5 | 6 | 0 |
| X | 12096619 | 2 | 4 | 10 | 0 |
| X | 12141978 | 2 | 4 | 11 | 0 |
| X | 12207947 | 2 | 4 | 9 | 0 |
| X | 12239920 | 2 | 9 | 5 | 0 |
| X | 12269545 | 2 | 8 | 5 | 0 |
| X | 12313053 | 2 | 6 | 9 | 0 |
| X | 12458821 | 2 | 9 | 5 | 0 |
| X | 12556147 | 2 | 6 | 5 | 0 |
| X | 14603097 | 2 | 5 | 12 | 0 |
| X | 14707769 | 2 | 5 | 8 | 0 |
| X | 14749593 | 2 | 5 | 5 | 0 |
| X | 15048299 | 2 | 5 | 10 | 0 |
| X | 15059866 | 2 | 11 | 5 | 0 |
| X | 15102053 | 2 | 4 | 13 | 0 |
| X | 15114387 | 2 | 20 | 8 | 0 |
| X | 15137423 | 2 | 11 | 11 | 0 |
| X | 15142288 | 2 | 7 | 6 | 0 |
| X | 15162266 | 2 | 5 | 6 | 0 |
| X | 15164905 | 2 | 5 | 8 | 0 |
| X | 15164922 | 2 | 17 | 13 | 0 |
| X | 15167273 | 2 | 15 | 6 | 0 |
| X | 15175759 | 2 | 19 | 8 | 0 |
| X | 15183258 | 2 | 9 | 6 | 0 |
| X | 15185642 | 2 | 33 | 3 | 0 |
| X | 15191031 | 2 | 5 | 6 | 0 |
| X | 15193675 | 2 | 5 | 7 | 0 |
| X | 15212362 | 2 | 12 | 6 | 0 |
| X | 15217619 | 2 | 9 | 10 | 0 |
| X | 15218834 | 2 | 7 | 11 | 0 |
| X | 15227376 | 2 | 11 | 4 | 0 |
| X | 15227465 | 2 | 9 | 9 | 0 |
| X | 15240498 | 2 | 5 | 14 | 0 |
| X | 15249994 | 2 | 6 | 5 | 0 |
| X | 15252580 | 2 | 6 | 15 | 0 |
| X | 15262518 | 2 | 14 | 10 | 0 |
| X | 15266913 | 2 | 23 | 6 | 0 |
| X | 15271468 | 2 | 9 | 13 | 0 |
| X | 15274752 | 2 | 6 | 16 | 0 |
| X | 15275079 | 2 | 29 | 12 | 0 |
| X | 15286060 | 2 | 5 | 6 | 0 |
| X | 15289847 | 2 | 7 | 25 | 0 |
| X | 15297213 | 2 | 5 | 7 | 0 |
| X | 15298064 | 2 | 9 | 27 | 0 |
| X | 15298998 | 2 | 10 | 10 | 0 |
| X | 15319300 | 2 | 11 | 11 | 0 |
| X | 15363208 | 2 | 6 | 26 | 0 |
| X | 15384676 | 2 | 6 | 8 | 0 |
| X | 15385493 | 2 | 5 | 16 | 0 |
| X | 15386715 | 2 | 8 | 14 | 0 |
| X | 15394459 | 2 | 23 | 7 | 0 |
| X | 15491447 | 2 | 5 | 5 | 0 |
| X | 15542905 | 2 | 21 | 7 | 0 |
| X | 15998712 | 2 | 5 | 9 | 0 |
| X | 15998900 | 2 | 10 | 11 | 0 |
| X | 16002074 | 2 | 9 | 14 | 0 |
| X | 16002291 | 2 | 16 | 9 | 0 |
| X | 16006378 | 2 | 7 | 18 | 0 |
| X | 16006430 | 2 | 10 | 11 | 0 |
| X | 16018665 | 2 | 11 | 4 | 0 |
| X | 16029434 | 2 | 10 | 19 | 0 |
| X | 16029737 | 2 | 13 | 5 | 0 |
| X | 16033735 | 2 | 5 | 21 | 0 |
| X | 16034287 | 2 | 4 | 11 | 0 |
| X | 16037960 | 2 | 9 | 8 | 0 |
| X | 16038269 | 2 | 7 | 8 | 0 |
| X | 16046104 | 2 | 15 | 8 | 0 |
| X | 16055780 | 2 | 9 | 10 | 0 |
| X | 16068000 | 2 | 22 | 7 | 0 |
| X | 16069855 | 2 | 8 | 5 | 0 |
| X | 16070101 | 2 | 7 | 9 | 0 |
| X | 16128675 | 2 | 7 | 11 | 0 |
| X | 16132243 | 2 | 6 | 25 | 0 |
| X | 16132391 | 2 | 25 | 5 | 0 |
| X | 16132434 | 2 | 9 | 5 | 0 |
| X | 16135831 | 2 | 22 | 15 | 0 |
| X | 16152971 | 2 | 8 | 5 | 0 |
| X | 16154920 | 2 | 18 | 15 | 0 |
| X | 16155871 | 2 | 12 | 20 | 0 |
| X | 16156065 | 2 | 5 | 7 | 0 |
| X | 16162179 | 2 | 8 | 7 | 0 |
| X | 16166309 | 2 | 7 | 5 | 0 |
| X | 16171818 | 2 | 5 | 9 | 0 |
| X | 16172123 | 2 | 6 | 10 | 0 |
| X | 16172257 | 2 | 6 | 13 | 0 |
| X | 16175381 | 2 | 12 | 6 | 0 |
| X | 16176114 | 2 | 12 | 7 | 0 |
| X | 16176702 | 2 | 21 | 14 | 0 |
| X | 16178011 | 2 | 17 | 14 | 0 |
| X | 16179159 | 2 | 20 | 7 | 0 |
| X | 16181757 | 2 | 13 | 19 | 0 |
| X | 16182549 | 2 | 17 | 11 | 0 |
| X | 16186737 | 2 | 22 | 5 | 0 |
| X | 16186749 | 2 | 24 | 6 | 0 |
| X | 16186872 | 2 | 15 | 4 | 0 |
| X | 16192279 | 2 | 5 | 10 | 0 |
| X | 16192480 | 2 | 7 | 6 | 0 |
| X | 16194422 | 2 | 20 | 19 | 0 |
| X | 16200073 | 2 | 15 | 4 | 0 |
| X | 16209492 | 2 | 19 | 12 | 0 |
| X | 16216333 | 2 | 6 | 9 | 0 |
| X | 16218915 | 2 | 14 | 4 | 0 |
| X | 16250606 | 2 | 16 | 18 | 0 |
| X | 16252195 | 2 | 7 | 6 | 0 |
| X | 16254698 | 2 | 14 | 5 | 0 |
| X | 16255696 | 2 | 4 | 9 | 0 |
| X | 16271594 | 2 | 9 | 21 | 0 |
| X | 16273210 | 2 | 5 | 5 | 0 |
| X | 16274474 | 2 | 19 | 13 | 0 |
| X | 16277263 | 2 | 6 | 15 | 0 |
| X | 16277633 | 2 | 9 | 18 | 0 |
| X | 16279558 | 2 | 9 | 4 | 0 |
| X | 16281667 | 2 | 19 | 9 | 0 |
| X | 16281701 | 2 | 17 | 4 | 0 |
| X | 16283592 | 2 | 23 | 7 | 0 |
| X | 16283607 | 2 | 26 | 6 | 0 |
| X | 16284978 | 2 | 10 | 6 | 0 |
| X | 16286191 | 2 | 18 | 15 | 0 |
| X | 16289601 | 2 | 12 | 8 | 0 |
| X | 16291254 | 2 | 16 | 5 | 0 |
| X | 16294455 | 2 | 21 | 6 | 0 |
| X | 16295584 | 2 | 4 | 13 | 0 |
| X | 16300561 | 2 | 10 | 4 | 0 |
| X | 16320818 | 2 | 10 | 12 | 0 |
| X | 16321083 | 2 | 22 | 13 | 0 |
| X | 16321227 | 2 | 5 | 7 | 0 |
| X | 16321537 | 2 | 24 | 8 | 0 |
| X | 16333870 | 2 | 25 | 14 | 0 |
| X | 16350747 | 2 | 15 | 7 | 0 |
| X | 16355360 | 2 | 11 | 16 | 0 |
| X | 16359591 | 2 | 6 | 15 | 0 |
| X | 16359983 | 2 | 14 | 4 | 0 |
| X | 16362187 | 2 | 5 | 8 | 0 |
| X | 16362524 | 2 | 10 | 4 | 0 |
| X | 16368881 | 2 | 15 | 5 | 0 |
| X | 16372407 | 2 | 8 | 11 | 0 |
| X | 16372414 | 2 | 11 | 9 | 0 |
| X | 16375173 | 2 | 15 | 8 | 0 |
| X | 16401340 | 2 | 14 | 4 | 0 |
| X | 16404418 | 2 | 7 | 16 | 0 |
| X | 16408533 | 2 | 5 | 6 | 0 |
| X | 16410228 | 2 | 4 | 11 | 0 |
| X | 16411828 | 2 | 14 | 6 | 0 |
| X | 16416940 | 2 | 16 | 7 | 0 |
| X | 16418262 | 2 | 15 | 5 | 0 |
| X | 16425056 | 2 | 10 | 4 | 0 |
| X | 16427438 | 2 | 10 | 4 | 0 |
| X | 16427526 | 2 | 30 | 18 | 0 |
| X | 16430791 | 2 | 12 | 5 | 0 |
| X | 16433488 | 2 | 26 | 4 | 0 |
| X | 16433725 | 2 | 12 | 6 | 0 |
| X | 16435014 | 2 | 5 | 14 | 0 |
| X | 16435032 | 2 | 5 | 11 | 0 |
| X | 16435076 | 2 | 8 | 10 | 0 |
| X | 16436554 | 2 | 5 | 8 | 0 |
| X | 16448975 | 2 | 9 | 4 | 0 |
| X | 16451133 | 2 | 5 | 16 | 0 |
| X | 16462918 | 2 | 5 | 23 | 0 |
| X | 16468122 | 2 | 8 | 8 | 0 |
| X | 16485730 | 2 | 4 | 19 | 0 |
| X | 16538838 | 2 | 11 | 7 | 0 |
| X | 16549141 | 2 | 14 | 4 | 0 |
| X | 16566506 | 2 | 5 | 5 | 0 |
| X | 16582796 | 2 | 9 | 6 | 0 |
| X | 16604413 | 2 | 6 | 13 | 0 |
| X | 16605276 | 2 | 9 | 5 | 0 |
| X | 16608976 | 2 | 12 | 10 | 0 |
| X | 16622941 | 2 | 8 | 5 | 0 |
| X | 16622977 | 2 | 17 | 14 | 0 |
| X | 16672209 | 2 | 5 | 5 | 0 |
| X | 16726960 | 2 | 6 | 8 | 0 |
| X | 16739180 | 2 | 5 | 6 | 0 |
| X | 16753076 | 2 | 6 | 8 | 0 |
| X | 16792956 | 2 | 13 | 7 | 0 |
| X | 16812259 | 2 | 12 | 4 | 0 |
| X | 16859496 | 2 | 5 | 7 | 0 |
| X | 16889240 | 2 | 9 | 4 | 0 |
| X | 16904322 | 2 | 15 | 5 | 0 |
| X | 16921852 | 2 | 9 | 24 | 0 |
| X | 16926585 | 2 | 5 | 5 | 0 |
| X | 16928738 | 2 | 6 | 6 | 0 |
| X | 16930783 | 2 | 12 | 5 | 0 |
| X | 16934497 | 2 | 5 | 6 | 0 |
| X | 16953310 | 2 | 6 | 8 | 0 |
| X | 16959502 | 2 | 7 | 5 | 0 |
| X | 16965515 | 2 | 11 | 5 | 0 |
| X | 16979428 | 2 | 7 | 11 | 0 |
| X | 16980644 | 2 | 13 | 4 | 0 |
| X | 16991806 | 2 | 5 | 21 | 0 |
| X | 17004950 | 2 | 21 | 5 | 0 |
| X | 17005004 | 2 | 32 | 5 | 0 |
| X | 17016975 | 2 | 16 | 16 | 0 |
| X | 17022025 | 2 | 8 | 16 | 0 |
| X | 17024729 | 2 | 19 | 8 | 0 |
| X | 17042412 | 2 | 4 | 20 | 0 |
| X | 17047577 | 2 | 7 | 6 | 0 |
| X | 17061564 | 2 | 7 | 13 | 0 |
| X | 17062351 | 2 | 5 | 5 | 0 |
| X | 17078216 | 2 | 4 | 12 | 0 |
| X | 17081163 | 2 | 4 | 24 | 0 |
| X | 17082461 | 2 | 4 | 12 | 0 |
| X | 17082767 | 2 | 4 | 9 | 0 |
| X | 17085423 | 2 | 17 | 15 | 0 |
| X | 17088091 | 2 | 20 | 26 | 0 |
| X | 17089161 | 2 | 6 | 19 | 0 |
| X | 17094652 | 2 | 14 | 23 | 0 |
| X | 17094823 | 2 | 10 | 21 | 0 |
| X | 17095202 | 2 | 6 | 16 | 0 |
| X | 17095568 | 2 | 7 | 23 | 0 |
| X | 17096042 | 2 | 23 | 4 | 0 |
| X | 17096400 | 2 | 21 | 10 | 0 |
| X | 17098192 | 2 | 5 | 17 | 0 |
| X | 17100250 | 2 | 30 | 12 | 0 |
| X | 17100856 | 2 | 22 | 25 | 0 |
| X | 17101287 | 2 | 22 | 7 | 0 |
| X | 17102819 | 2 | 19 | 16 | 0 |
| X | 17109980 | 2 | 8 | 17 | 0 |
| X | 17158513 | 2 | 5 | 6 | 0 |
| X | 17636400 | 2 | 12 | 4 | 0 |
| X | 17636679 | 2 | 5 | 6 | 0 |
| X | 17639668 | 2 | 6 | 6 | 0 |
| X | 17641428 | 2 | 10 | 18 | 0 |
| X | 17641527 | 2 | 5 | 8 | 0 |
| X | 17644403 | 2 | 13 | 13 | 0 |
| X | 17649368 | 2 | 6 | 16 | 0 |
| X | 17652125 | 2 | 7 | 16 | 0 |
| X | 17655012 | 2 | 11 | 18 | 0 |
| X | 17655648 | 2 | 6 | 6 | 0 |
| X | 17664714 | 2 | 10 | 14 | 0 |
| X | 17673130 | 2 | 17 | 10 | 0 |
| X | 17674802 | 2 | 10 | 5 | 0 |
| X | 17674825 | 2 | 17 | 7 | 0 |
| X | 17676368 | 2 | 4 | 24 | 0 |
| X | 17676434 | 2 | 6 | 29 | 0 |
| X | 17676603 | 2 | 8 | 7 | 0 |
| X | 17677841 | 2 | 31 | 17 | 0 |
| X | 17678091 | 2 | 9 | 12 | 0 |
| X | 17682467 | 2 | 5 | 15 | 0 |
| X | 17697385 | 2 | 12 | 9 | 0 |
| X | 17706408 | 2 | 11 | 9 | 0 |
| X | 17706533 | 2 | 4 | 14 | 0 |
| X | 17708428 | 2 | 5 | 19 | 0 |
| X | 17708528 | 2 | 12 | 21 | 0 |
| X | 17710188 | 2 | 5 | 17 | 0 |
| X | 17710855 | 2 | 6 | 17 | 0 |
| X | 17711563 | 2 | 16 | 4 | 0 |
| X | 17713491 | 2 | 12 | 4 | 0 |
| X | 17713682 | 2 | 11 | 30 | 0 |
| X | 17718102 | 2 | 17 | 16 | 0 |
| X | 17721688 | 2 | 5 | 8 | 0 |
| X | 17722207 | 2 | 4 | 12 | 0 |
| X | 17722416 | 2 | 13 | 27 | 0 |
| X | 17724209 | 2 | 20 | 5 | 0 |
| X | 17726373 | 2 | 11 | 4 | 0 |
| X | 17730959 | 2 | 11 | 30 | 0 |
| X | 17731739 | 2 | 5 | 12 | 0 |
| X | 17731996 | 2 | 10 | 20 | 0 |
| X | 17732231 | 2 | 17 | 16 | 0 |
| X | 17732439 | 2 | 3 | 31 | 0 |
| X | 17732879 | 2 | 10 | 29 | 0 |
| X | 17733107 | 2 | 7 | 34 | 0 |
| X | 17733480 | 2 | 10 | 27 | 0 |
| X | 17733686 | 2 | 7 | 8 | 0 |
| X | 17737524 | 2 | 10 | 21 | 0 |
| X | 17737821 | 2 | 12 | 18 | 0 |
| X | 17738079 | 2 | 8 | 22 | 0 |
| X | 17739190 | 2 | 17 | 17 | 0 |
| X | 17739355 | 2 | 5 | 11 | 0 |
| X | 17755922 | 2 | 13 | 28 | 0 |
| X | 17766984 | 2 | 8 | 9 | 0 |
| X | 17767124 | 2 | 11 | 6 | 0 |
| X | 17772322 | 2 | 6 | 7 | 0 |
| X | 17772371 | 2 | 17 | 11 | 0 |
| X | 17775562 | 2 | 5 | 8 | 0 |
| X | 17775580 | 2 | 5 | 12 | 0 |
| X | 17778512 | 2 | 8 | 27 | 0 |
| X | 17778651 | 2 | 9 | 16 | 0 |
| X | 17779090 | 2 | 4 | 16 | 0 |
| X | 17781894 | 2 | 4 | 15 | 0 |
| X | 17782517 | 2 | 6 | 5 | 0 |
| X | 17800468 | 2 | 5 | 10 | 0 |
| X | 17802202 | 2 | 17 | 6 | 0 |
| X | 17804003 | 2 | 17 | 10 | 0 |
| X | 17804917 | 2 | 28 | 10 | 0 |
| X | 17805346 | 2 | 12 | 6 | 0 |
| X | 17807636 | 2 | 11 | 29 | 0 |
| X | 17808883 | 2 | 9 | 10 | 0 |
| X | 17810353 | 2 | 16 | 11 | 0 |
| X | 17813729 | 2 | 10 | 4 | 0 |
| X | 17814606 | 2 | 4 | 20 | 0 |
| X | 17815257 | 2 | 9 | 19 | 0 |
| X | 17816014 | 2 | 5 | 10 | 0 |
| X | 17816210 | 2 | 9 | 29 | 0 |
| X | 17816295 | 2 | 3 | 34 | 0 |
| X | 17816310 | 2 | 5 | 40 | 0 |
| X | 17826062 | 2 | 11 | 19 | 0 |
| X | 17826604 | 2 | 13 | 30 | 0 |
| X | 17829535 | 2 | 24 | 12 | 0 |
| X | 17831219 | 2 | 16 | 8 | 0 |
| X | 17832304 | 2 | 11 | 6 | 0 |
| X | 17874481 | 2 | 22 | 8 | 0 |
| X | 17876772 | 2 | 8 | 7 | 0 |
| X | 17883037 | 2 | 9 | 4 | 0 |
| X | 17885434 | 2 | 4 | 15 | 0 |
| X | 17886121 | 2 | 11 | 8 | 0 |
| X | 17888697 | 2 | 10 | 8 | 0 |
| X | 17898322 | 2 | 5 | 13 | 0 |
| X | 17898904 | 2 | 4 | 15 | 0 |
| X | 17898989 | 2 | 6 | 9 | 0 |
| X | 17899205 | 2 | 5 | 21 | 0 |
| X | 17900308 | 2 | 20 | 12 | 0 |
| X | 17900860 | 2 | 5 | 12 | 0 |
| X | 17900911 | 2 | 16 | 12 | 0 |
| X | 17903833 | 2 | 22 | 5 | 0 |
| X | 17907350 | 2 | 14 | 6 | 0 |
| X | 17918621 | 2 | 12 | 22 | 0 |
| X | 17922656 | 2 | 18 | 6 | 0 |
| X | 17923611 | 2 | 6 | 5 | 0 |
| X | 17925370 | 2 | 6 | 15 | 0 |
| X | 17925814 | 2 | 7 | 17 | 0 |
| X | 17926015 | 2 | 4 | 9 | 0 |
| X | 17926245 | 2 | 9 | 5 | 0 |
| X | 17931066 | 2 | 8 | 10 | 0 |
| X | 17934851 | 2 | 21 | 8 | 0 |
| X | 17939285 | 2 | 14 | 7 | 0 |
| X | 17939779 | 2 | 10 | 8 | 0 |
| X | 17941362 | 2 | 7 | 14 | 0 |
| X | 17943675 | 2 | 6 | 6 | 0 |
| X | 17945957 | 2 | 9 | 7 | 0 |
| X | 17947151 | 2 | 8 | 9 | 0 |
| X | 17960449 | 2 | 17 | 7 | 0 |
| X | 17960968 | 2 | 34 | 16 | 0 |
| X | 17961016 | 2 | 7 | 6 | 0 |
| X | 17962268 | 2 | 22 | 4 | 0 |
| X | 17962458 | 2 | 19 | 4 | 0 |
| X | 19098515 | 2 | 14 | 16 | 0 |
| X | 19099578 | 2 | 9 | 6 | 0 |
| X | 19099892 | 2 | 16 | 12 | 0 |
| X | 19101747 | 2 | 9 | 4 | 0 |
| X | 19103397 | 2 | 5 | 12 | 0 |
| X | 19112930 | 2 | 7 | 6 | 0 |
| X | 19113908 | 2 | 10 | 8 | 0 |
| X | 19123008 | 2 | 9 | 4 | 0 |
| X | 19141251 | 2 | 8 | 5 | 0 |
| X | 19152137 | 2 | 14 | 8 | 0 |
| X | 19152203 | 2 | 13 | 5 | 0 |
| X | 19158485 | 2 | 6 | 6 | 0 |
| X | 19159009 | 2 | 7 | 9 | 0 |
| X | 19161962 | 2 | 6 | 7 | 0 |
| X | 19162084 | 2 | 23 | 6 | 0 |
| X | 19166705 | 2 | 15 | 4 | 0 |
| X | 19172116 | 2 | 13 | 8 | 0 |
| X | 19180738 | 2 | 7 | 10 | 0 |
| X | 19185386 | 2 | 6 | 11 | 0 |
| X | 19186055 | 2 | 23 | 7 | 0 |
| X | 19189843 | 2 | 11 | 11 | 0 |
| X | 19190590 | 2 | 9 | 7 | 0 |
| X | 19211170 | 2 | 5 | 9 | 0 |
| X | 19211754 | 2 | 10 | 6 | 0 |
| X | 19213716 | 2 | 11 | 11 | 0 |
| X | 19222491 | 2 | 9 | 5 | 0 |
| X | 19223662 | 2 | 14 | 9 | 0 |
| X | 19224000 | 2 | 15 | 11 | 0 |
| X | 19226820 | 2 | 14 | 9 | 0 |
| X | 19228767 | 2 | 9 | 5 | 0 |
| X | 19231613 | 2 | 4 | 10 | 0 |
| X | 19231700 | 2 | 13 | 5 | 0 |
| X | 19234509 | 2 | 7 | 17 | 0 |
| X | 19249378 | 2 | 11 | 9 | 0 |
| X | 19252806 | 2 | 8 | 14 | 0 |
| X | 19302077 | 2 | 7 | 17 | 0 |
| X | 19313932 | 2 | 9 | 7 | 0 |
| X | 19325792 | 2 | 8 | 5 | 0 |
| X | 19325928 | 2 | 20 | 5 | 0 |
| X | 19354109 | 2 | 7 | 5 | 0 |
| X | 19368654 | 2 | 13 | 4 | 0 |
| X | 19370629 | 2 | 15 | 8 | 0 |
| X | 19371113 | 2 | 12 | 4 | 0 |
| X | 19381694 | 2 | 5 | 9 | 0 |
| X | 19397265 | 2 | 8 | 5 | 0 |
| X | 19397911 | 2 | 16 | 7 | 0 |
| X | 19400540 | 2 | 11 | 7 | 0 |
| X | 19419970 | 2 | 18 | 5 | 0 |
| X | 19434400 | 2 | 4 | 9 | 0 |
| X | 19436103 | 2 | 5 | 6 | 0 |
| X | 19436351 | 2 | 17 | 5 | 0 |
| X | 19437144 | 2 | 8 | 9 | 0 |
| X | 19439240 | 2 | 7 | 23 | 0 |
| X | 19440316 | 2 | 5 | 8 | 0 |
| X | 19442061 | 2 | 13 | 4 | 0 |
| X | 19442836 | 2 | 14 | 4 | 0 |
| X | 19444826 | 2 | 10 | 8 | 0 |
| X | 19446914 | 2 | 6 | 5 | 0 |
| X | 19481114 | 2 | 10 | 4 | 0 |
| X | 19481347 | 2 | 17 | 14 | 0 |
| X | 19481737 | 2 | 10 | 23 | 0 |
| X | 19482442 | 2 | 13 | 12 | 0 |
| X | 19487612 | 2 | 33 | 3 | 0 |
| X | 19489379 | 2 | 11 | 16 | 0 |
| X | 19506965 | 2 | 13 | 8 | 0 |
| X | 19507198 | 2 | 8 | 8 | 0 |
| X | 19521071 | 2 | 14 | 4 | 0 |
| X | 19521507 | 2 | 10 | 4 | 0 |
| X | 19524041 | 2 | 10 | 7 | 0 |
| X | 19525279 | 2 | 20 | 13 | 0 |
| X | 19525366 | 2 | 20 | 10 | 0 |
| X | 19527115 | 2 | 11 | 5 | 0 |
| X | 19541947 | 2 | 12 | 4 | 0 |
| X | 20409822 | 2 | 12 | 4 | 0 |
| X | 20410673 | 2 | 7 | 7 | 0 |
| X | 20411650 | 2 | 8 | 8 | 0 |
| X | 20413082 | 2 | 8 | 14 | 0 |
| X | 20415096 | 2 | 12 | 5 | 0 |
| X | 20418369 | 2 | 12 | 9 | 0 |
| X | 20422073 | 2 | 10 | 4 | 0 |
| X | 20422741 | 2 | 10 | 5 | 0 |
| X | 20426494 | 2 | 14 | 10 | 0 |
| X | 20427526 | 2 | 12 | 5 | 0 |
| X | 20440076 | 2 | 6 | 14 | 0 |
| X | 20451590 | 2 | 5 | 5 | 0 |
| X | 20460126 | 2 | 5 | 6 | 0 |
| X | 20463504 | 2 | 12 | 4 | 0 |
| X | 20463909 | 2 | 5 | 6 | 0 |
| X | 20470332 | 2 | 17 | 9 | 0 |
| X | 20489152 | 2 | 7 | 15 | 0 |
| X | 20503402 | 2 | 8 | 8 | 0 |
| X | 20573899 | 2 | 12 | 7 | 0 |
| X | 20574018 | 2 | 10 | 8 | 0 |
| X | 20578070 | 2 | 11 | 6 | 0 |
| X | 20583921 | 2 | 20 | 8 | 0 |
| X | 20584893 | 2 | 24 | 5 | 0 |
| X | 20585127 | 2 | 17 | 9 | 0 |
| X | 20634672 | 2 | 10 | 5 | 0 |
| X | 20634956 | 2 | 7 | 7 | 0 |
| X | 20643155 | 2 | 4 | 16 | 0 |
| X | 20644399 | 2 | 7 | 10 | 0 |
| X | 20674612 | 2 | 4 | 10 | 0 |
| X | 20686174 | 2 | 17 | 12 | 0 |
| X | 20703252 | 2 | 5 | 9 | 0 |
| X | 20715600 | 2 | 19 | 10 | 0 |
| X | 20722902 | 2 | 14 | 10 | 0 |
| X | 20723988 | 2 | 13 | 8 | 0 |
| X | 20741783 | 2 | 11 | 8 | 0 |
| X | 20743149 | 2 | 13 | 5 | 0 |
| X | 20746322 | 2 | 14 | 8 | 0 |
| X | 20748655 | 2 | 5 | 12 | 0 |
| X | 20750151 | 2 | 6 | 8 | 0 |
| X | 20753738 | 2 | 7 | 7 | 0 |
| X | 20757373 | 2 | 5 | 5 | 0 |
| X | 20796458 | 2 | 4 | 15 | 0 |
| X | 20819537 | 2 | 6 | 5 | 0 |
| X | 20847281 | 2 | 8 | 5 | 0 |
| X | 20854993 | 2 | 12 | 7 | 0 |
| X | 20873814 | 2 | 13 | 5 | 0 |
| X | 20898829 | 2 | 8 | 13 | 0 |
| X | 20919300 | 2 | 4 | 21 | 0 |
| X | 20920968 | 2 | 5 | 13 | 0 |
| X | 20938290 | 2 | 10 | 14 | 0 |
| X | 20951061 | 2 | 7 | 25 | 0 |
| X | 20971255 | 2 | 10 | 16 | 0 |
| X | 20973497 | 2 | 5 | 5 | 0 |
| X | 21110602 | 2 | 11 | 15 | 0 |
| X | 21157311 | 2 | 6 | 22 | 0 |
| X | 21225985 | 2 | 4 | 14 | 0 |
| X | 21226096 | 2 | 7 | 9 | 0 |

**Supplemental Table 2:**
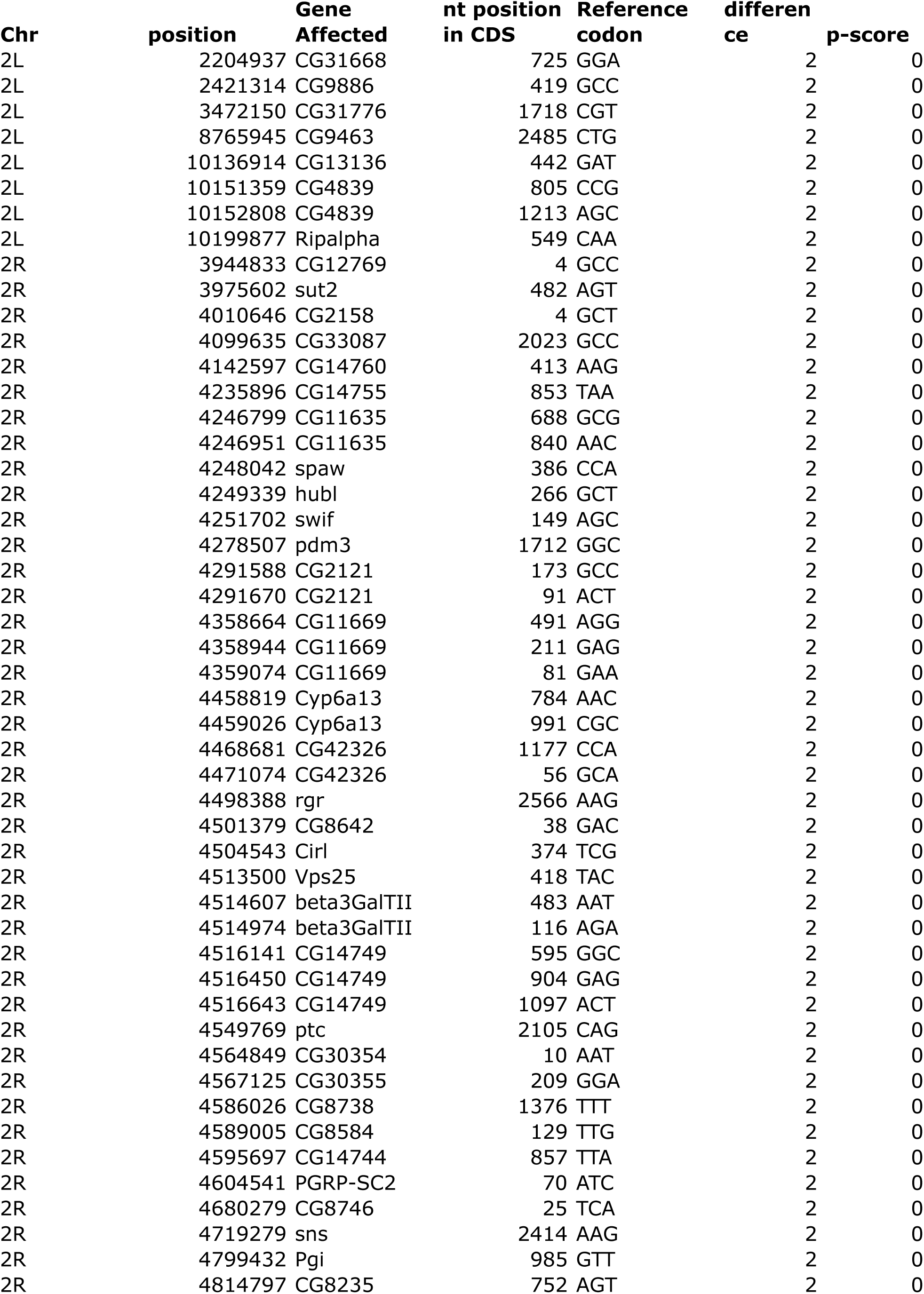
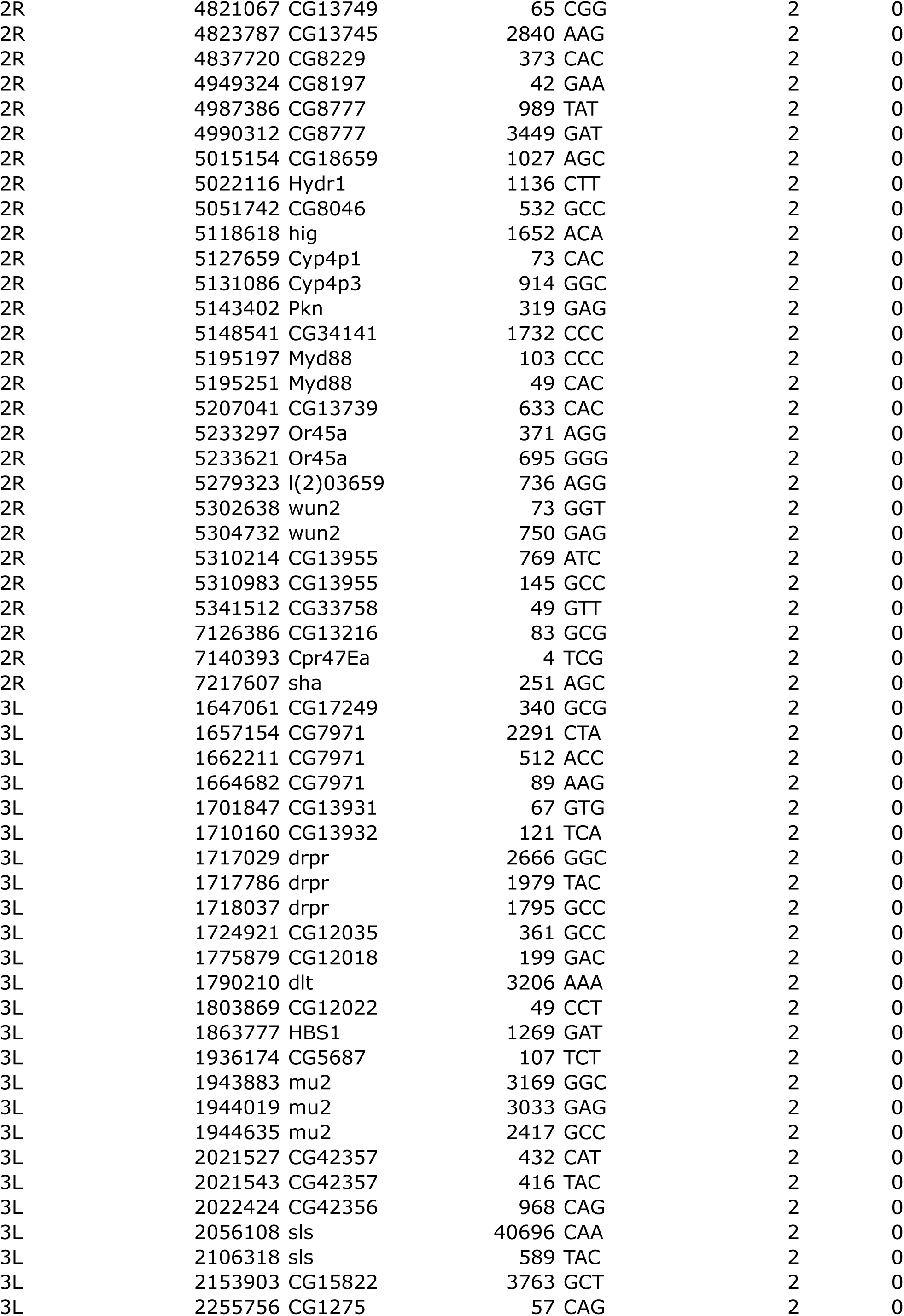
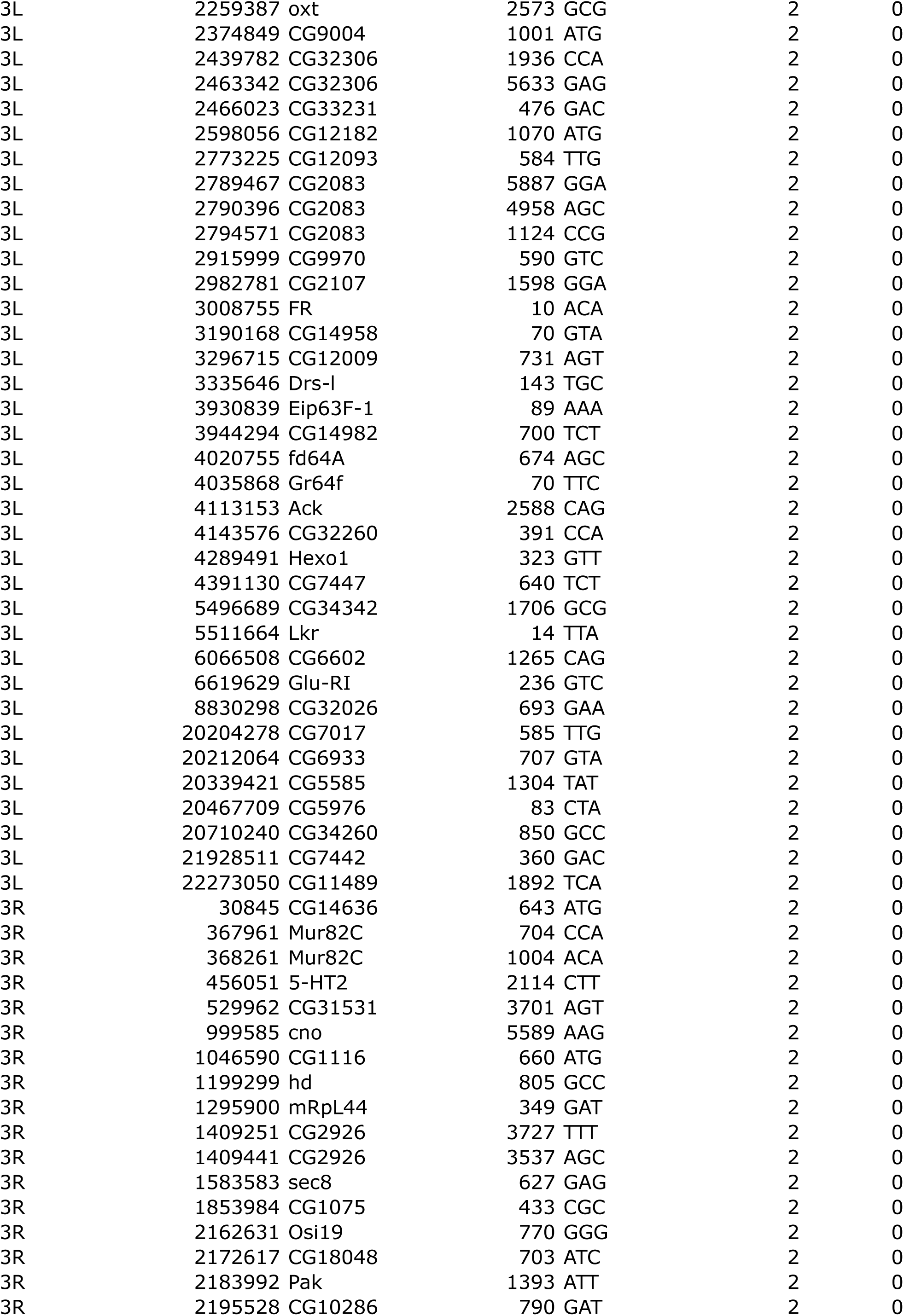
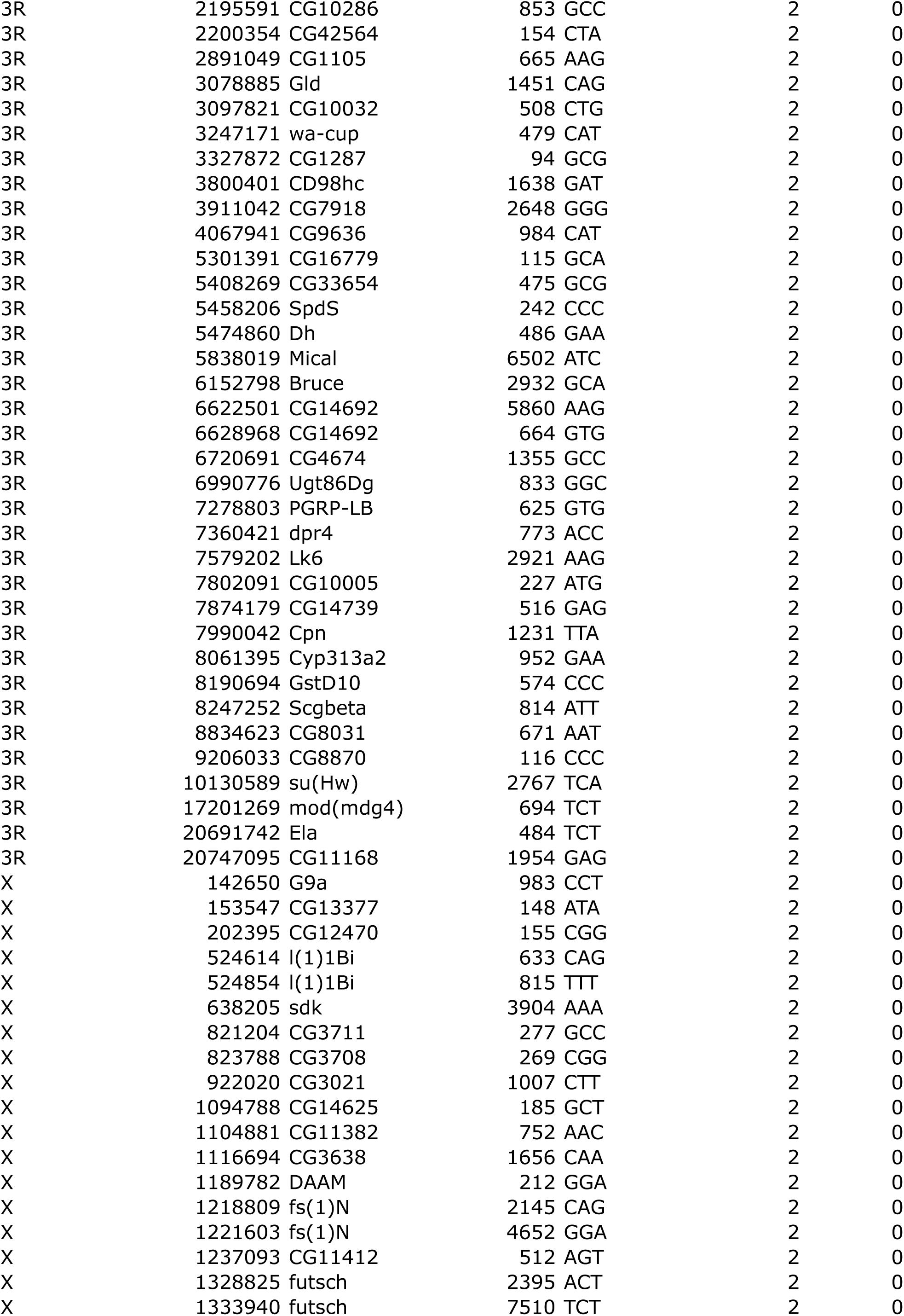
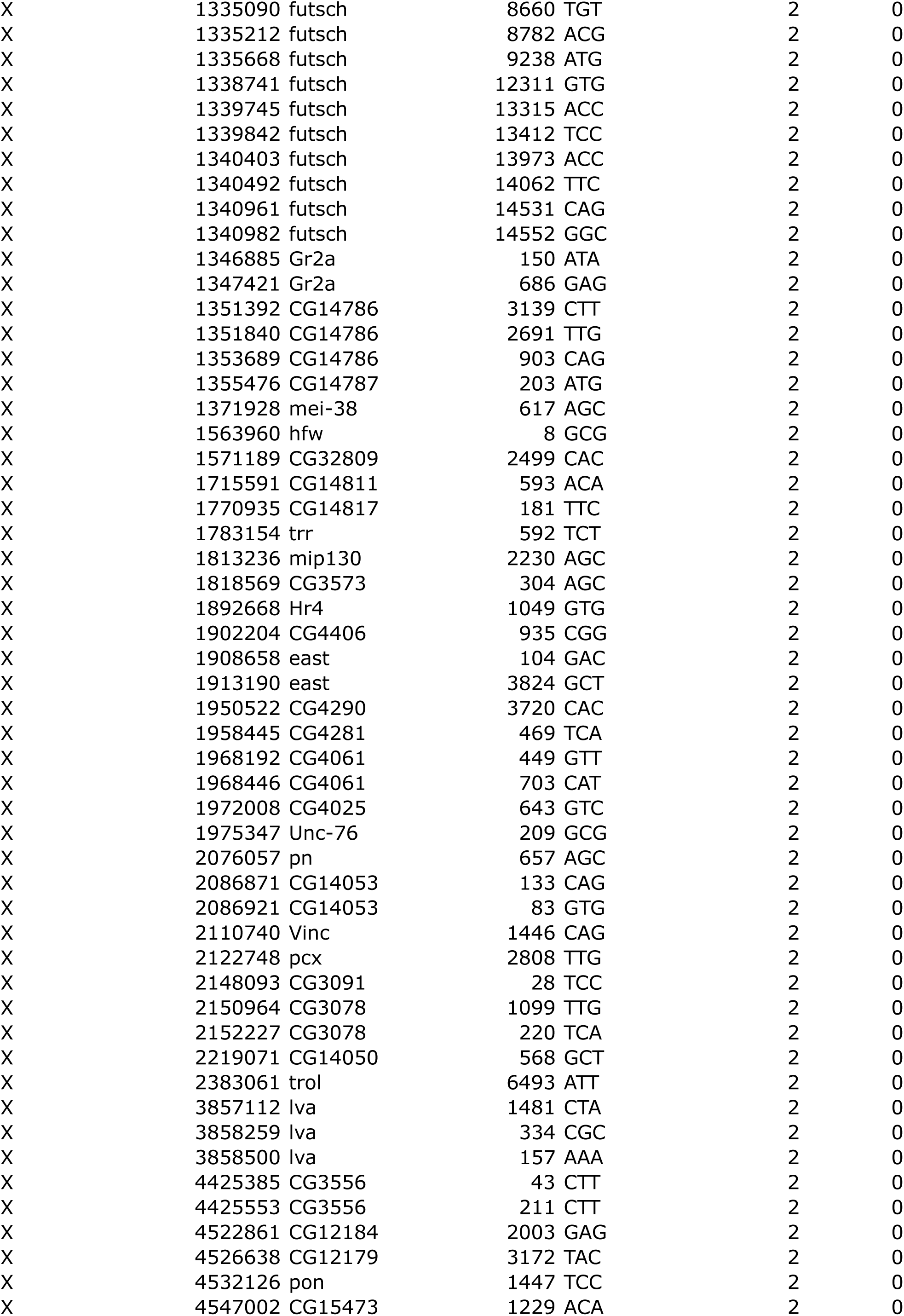
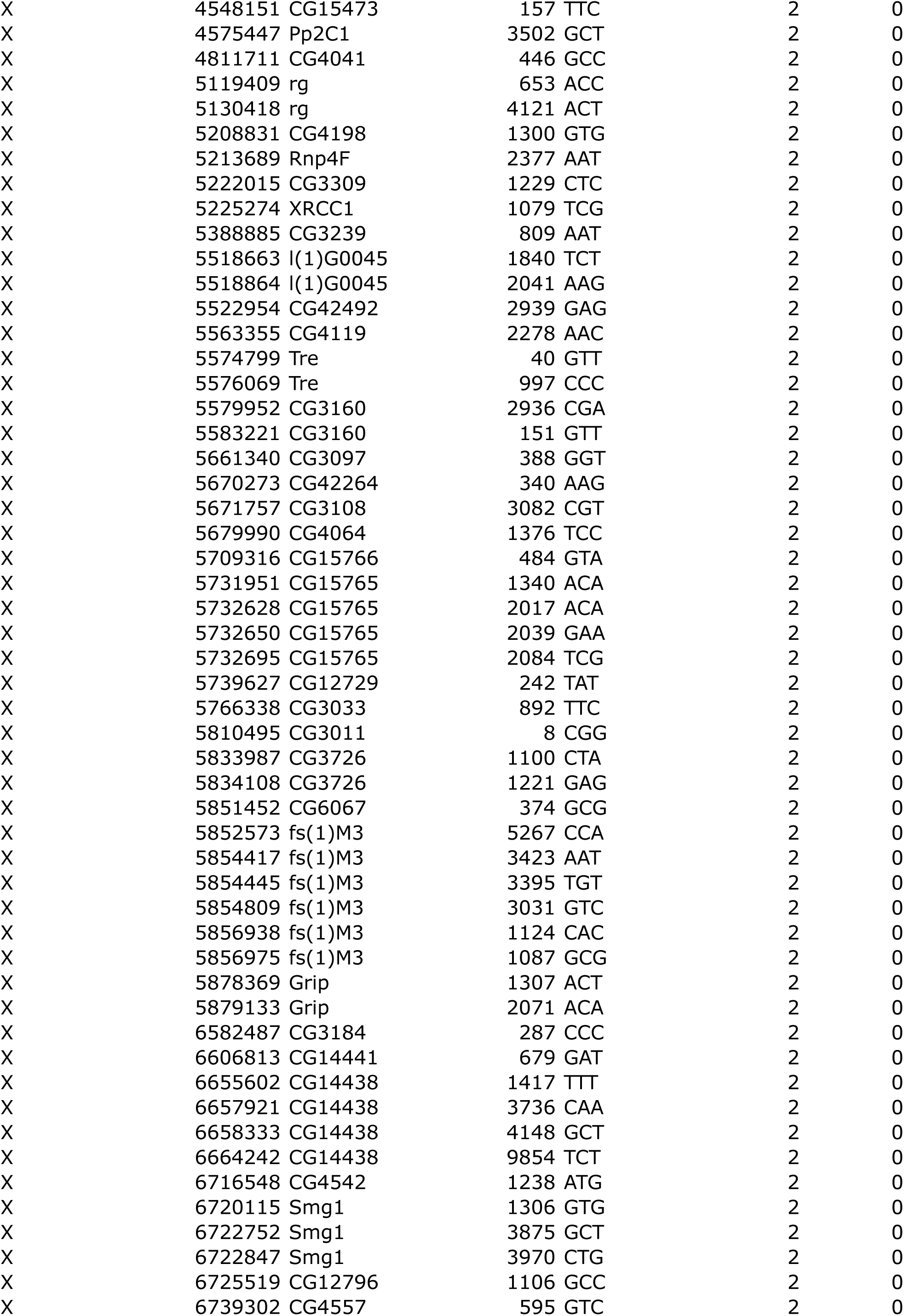
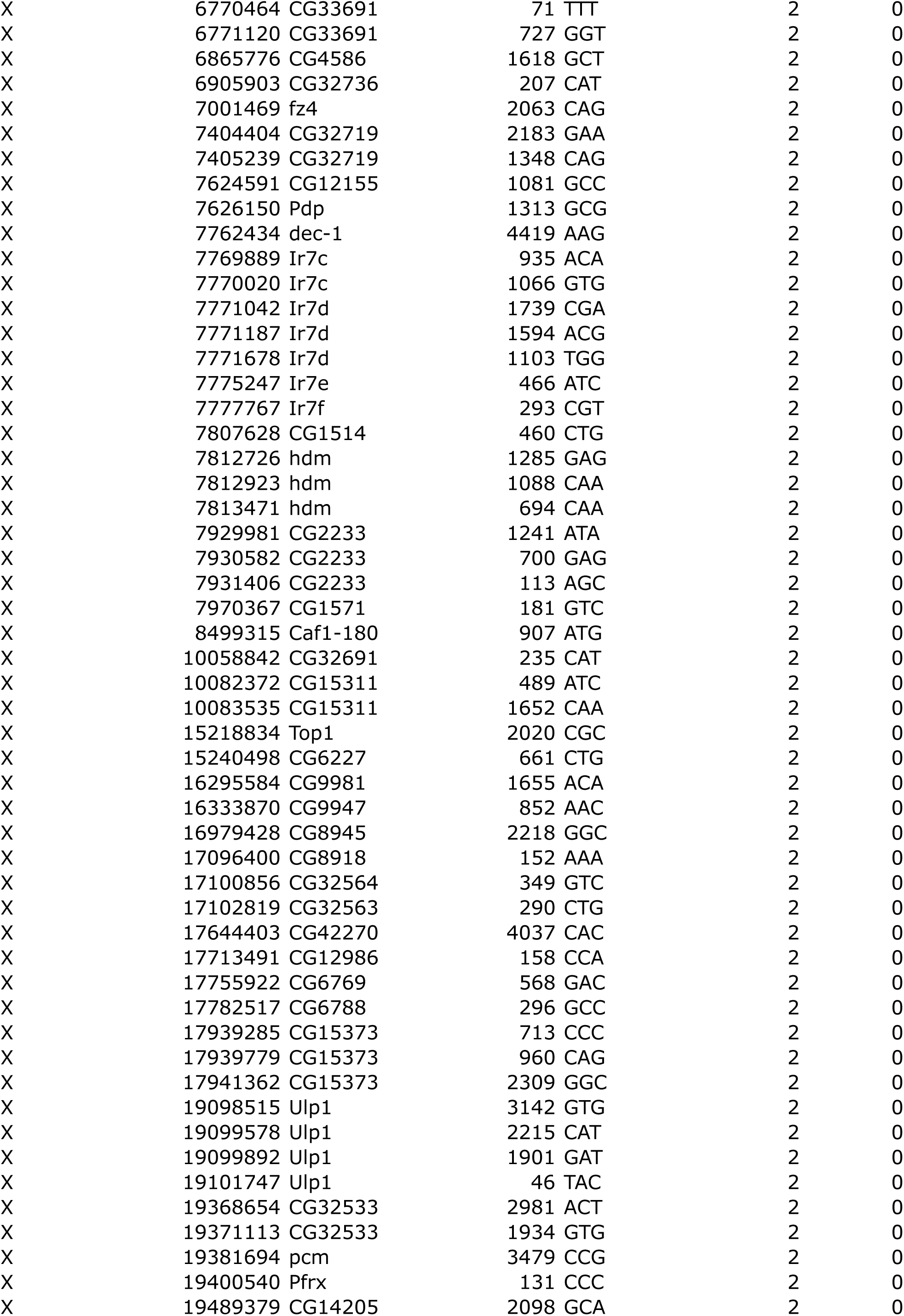
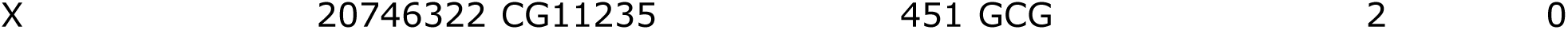
Differential non-synonymous SNPs.

